# High-Yield Recovery of Reactive Nitrogen as Cyanophycin by Engineering *Acinetobacter baylyi* ADP1 under Wastewater-Relevant Conditions

**DOI:** 10.64898/2026.06.25.733799

**Authors:** Kevin Fitzgerald, Keith Tyo

## Abstract

Municipal wastewater constitutes a major reservoir of unutilized reactive nitrogen, representing a significant opportunity for biological valorization. The biopolymer cyanophycin is promising as a means of nitrogen capture and recovery, but current production strategies are not optimized for the physicochemical constraints of municipal wastewater systems. Here, we engineered the naturally competent soil bacterium *Acinetobacter baylyi* ADP1 ISx to synthesize cyanophycin from carbon and nitrogen sources prevalent in municipal wastewater and over a range of wastewater-relevant temperatures. To overcome the recurring problem of arginine availability limiting cyanophycin synthesis, we engineered an arginine-producing strain (AP1) which accumulated cyanophycin when grown on acetate and ammonium (19% CDW), nitrate (9% CDW), or urea (29% CDW) and without arginine supplementation. During this work, we observed that conditions associated with reduced cell fitness correlated with increased intracellular cyanophycin content. As temperature strongly influences cell growth but cannot be realistically modulated in wastewater contexts, we investigated the potential of induced fructose-auxotrophy to modulate cell growth independently from temperature. This intervention, accomplished with a single knockout (Δ*gap*), expanded the effective range of cyanophycin accumulation from 12 °C up to 30 °C. Collectively, these results establish the relevance of arginine-producing strains for cyanophycin biosynthesis and position *A. baylyi* as a promising chassis for continued development under real-world wastewater conditions.

## Introduction

Municipal wastewater represents a substantial yet underutilized reservoir of low-cost nutrients, including an estimated sixteen million tons of nitrogen each year^1^. In this context biological nutrient removal (BNR) remains one of the most common means of removing nitrogen from wastewater and serves as one of the most widely implemented large-scale applications of biotechnology^2^. These strategies take advantage of the innate capacity of microorganisms to capture and process dilute aqueous nitrogen species while requiring minimal external inputs. While robust and well-studied, the final outcome of this transformation is dinitrogen gas, an inert product with limited utility and commercial potential. The development of wastewater treatment technologies that not only retain the innate nitrogen-concentrating features of biological systems but also support the production and purification of value-added products represents a promising opportunity to transform the ubiquitous wastewater treatment plant into a next-generation biorefinery.

The intracellular nitrogen storage polymer cyanophycin represents one possible route toward this goal. Cyanophycin is synthesized non-ribosomally by cyanophycin synthetase (CphA1) and consists of a polyaspartic acid backbone linked through standard peptide bonds, with arginine or lysine residues attached to the Asp β-carboxyl group via isopeptide bonds^3^. The incorporation of these basic amino acids imparts a zwitterionic character to the polymer, promoting the formation of insoluble intracellular granules during cell growth and enabling its selective recovery from biomass by modulating pH. Wild-type cyanophycin producers also typically produce a cyanophycinase (*cphI* or *cphB*) which depolymerizes cyanophycin into dipeptides to facilitate their eventual reuse. Cyanophycin has several proposed applications, including as a source of polyaspartic acid for industrial descaling or water softening, as a provider of dipeptides with potential use in human nutrition^4, 5^, and as replacement for the lysine HCl commonly used in livestock feed^6, 7^.

Cyanophycin-mediated nitrogen recovery has been demonstrated previously in multiple waste-associated contexts: cyanobacterial cultures have been shown to accumulate cyanophycin when cultivated on synthetic urine media^8^, *Escherichia coli* has been engineered to produce cyanophycin from mock manure hydrolysates^9^, and activated sludge-derived consortia have been enriched to produce cyanophycin, achieving titers equivalent to 10% of total biomass^10^. These observations highlight the feasibility of employing cyanophycin as a biological reservoir for nitrogen recovered from waste streams. Beyond these application-oriented demonstrations, strain engineering efforts have largely proven effective at enhancing cyanophycin production. Two such landmark studies achieved per-cell cyanophycin titers of 53% cell dry weight (CDW) in *Ralstonia eutropha*^11^ and 21% CDW in *E. coli* at the 500 L scale^12^. Despite these achievements, reliance on high-value carbon sources (e.g. gluconate, fructose) or amino acid supplementation (e.g. casamino acids, arginine) remain perennial challenges in the field, limiting the suitability of existing strains for wastewater applications.

Contemporary BNR typically occurs during secondary treatment once large particulates have been removed during primary treatment. Although the composition of municipal wastewater varies, process integration at this stage creates two broad requirements for a microbial chassis suitable for cyanophycin-mediated nitrogen recovery. First, the chassis should be able to catabolize volatile fatty acids (VFAs) such as acetate, propionate and butyrate, which form during anaerobic digestion and are often added to supply carbon during denitrification^5, 6^. Second, the chassis should be able to assimilate the nitrogen species removed during contemporary biological interventions, including ammonium, nitrite, and nitrate^15^.

*Acinetobacter baylyi* ADP1, formerly *A. calcoaceticus*, is a naturally competent soil bacterium whose metabolic characteristics are well-suited for cyanophycin-based nitrogen recovery from wastewater. In addition to its compatibility with acetate consumption, genome annotation indicates the presence of a complete methylcitrate cycle, suggesting the capacity to metabolize propionate. Furthermore, *A. baylyi* possesses genes associated with the uptake of ammonium, nitrate, and urea, supporting its ability to remove and utilize the primary nitrogen species present in municipal wastewater. Previous work has demonstrated the recovery of cyanophycin up to 46% CDW, however, this was done under conditions not amenable to robust production in municipal wastewater - most notably strict phosphate limitation (<50 μM) and abundant arginine supplementation (>60 mM)^16, 17^.

In this study we describe the engineering of *A. baylyi* ADP1 for cyanophycin production using carbon and nitrogen sources as well as cultivation temperatures reflecting those found in municipal wastewater. We constructed strains that synthesize cyanophycin from acetate together with ammonium, nitrate, or urea, and reached a maximum intracellular content of 29% CDW without arginine supplementation and in the presence of abundant phosphate. In addition, we examined how growth physiology relates to polymer accumulation and used this information to generate a fructose auxotroph that sustains cyanophycin production across a range of cultivation temperatures. Together, this work constitutes a promising step towards the recovery of nitrogen from municipal wastewater.

## 2. Materials & Methods

### 2.1 Strains & Media

All cyanophycin production experiments were conducted in *A. baylyi* ADP1 ISx, generously provided by Prof. Jeffrey Barrick^18^. Routine culturing was performed in LB Miller medium, supplemented with 25 μg/mL kanamycin monosulfate or 50 μg/mL spectinomycin dihydrochloride pentahydrate when appropriate. Cyanophycin cultivation utilized a mineral salts medium (MSM) formulated previously^19^. The original formulation included 75 mM L-arginine, 10 mM (NH4)2SO4, 20 mM KCl, and 0.8 mM MgSO4. A 100x stock of trace metals was prepared by dissolution 8.2 g/L ethylenediaminetetraacetic acid, followed by 600 mg/L FeCl3, MnCl2·4H2O, ZnCl2, CuSO4·5H2O, CaCl2·2H2O, and NaCl with pH adjustments back to ∼7 between each addition to prevent precipitation. The trace metals were added to MSM such that each metal salt was present at 6 mg/L. Buffering was achieved initially using 50 mM 3-(N-morpholino)propanesulfonic acid (MOPS). In later experiments, buffering was instead achieved using 33.9 g/L Na2HPO4 and 15 g/L KH2PO4 (70mM PO4^3-^ total) - which also served as source of phosphate. Medium components were individually adjusted to approximately pH 7, resulting in MSM with an initial pH of 7.0-7.2. Specific modifications to carbon and nitrogen sources are described in the experimental procedures.

### 2.2 Cloning

A description of the modifications used in this study is provided below in **Table 1**. A list of select DNA sequences and primers used in this study are provided in **Table S1** and **Table S2**, respectively. PCR reactions for cloning and colony screening were performed using PrimeSTAR Max DNA Polymerase (Takara; San Jose, CA). For plasmid assembly, homology of ∼40 bp was introduced via PCR between fragments and Gibson Assembly was carried out using Gibson 2x Master Mix or NEBuilder HiFi DNA assembly Master Mix (New England Biolabs; Ipswich, MA). Gibson ligation products were transformed into chemically competent *Escherichia coli* DH5α.

**Table 1.** Genetic features investigated in this study.

| Feature | Description |
| --- | --- |
| *cphA* | Native *A. baylyi* cyanophycin synthetase. Knockouts are deficient in cyanophycin synthesis. |
| *cphI* | Native *A. baylyi* cyanophycin degrading enzyme (cyanophycinase). Often knocked out along with *cphA*. |
| *argR* | Arginine-sensitive transcription factor. Knockout putatively derepresses arginine biosynthesis prevents arginine catabolism. |
| *argB* | Acetylglutamate kinase, the second enzyme in *A. baylyi* arginine biosynthesis. Later replaced with *E. coli* homolog *argB<sub>Ec</sub>*. |
| *ACIAD3341* | Putative ornithine decarboxylase. |
| *fbsJ* | Putative ornithine decarboxylase. |
| *recA* | Recombinase A, responsible for homologous recombination in *A. baylyi*. Knockout hypothesized to enhance plasmid stability. |
| *mCherry* | Fluorescent red protein. Used to control for impact of heterologous protein expression on growth and measured product signals. |
| *AbcphA1* | Cyanophycin synthetase from *A. baylyi*. |
| *6308cphA1* | Cyanophycin synthetase from the cyanobacteria *Synechocystis* sp. 6308. |
| *TmcphA1* | Cyanophycin synthetase from the heterotroph *Tatumella morbirosei*. |

**Table 2.**
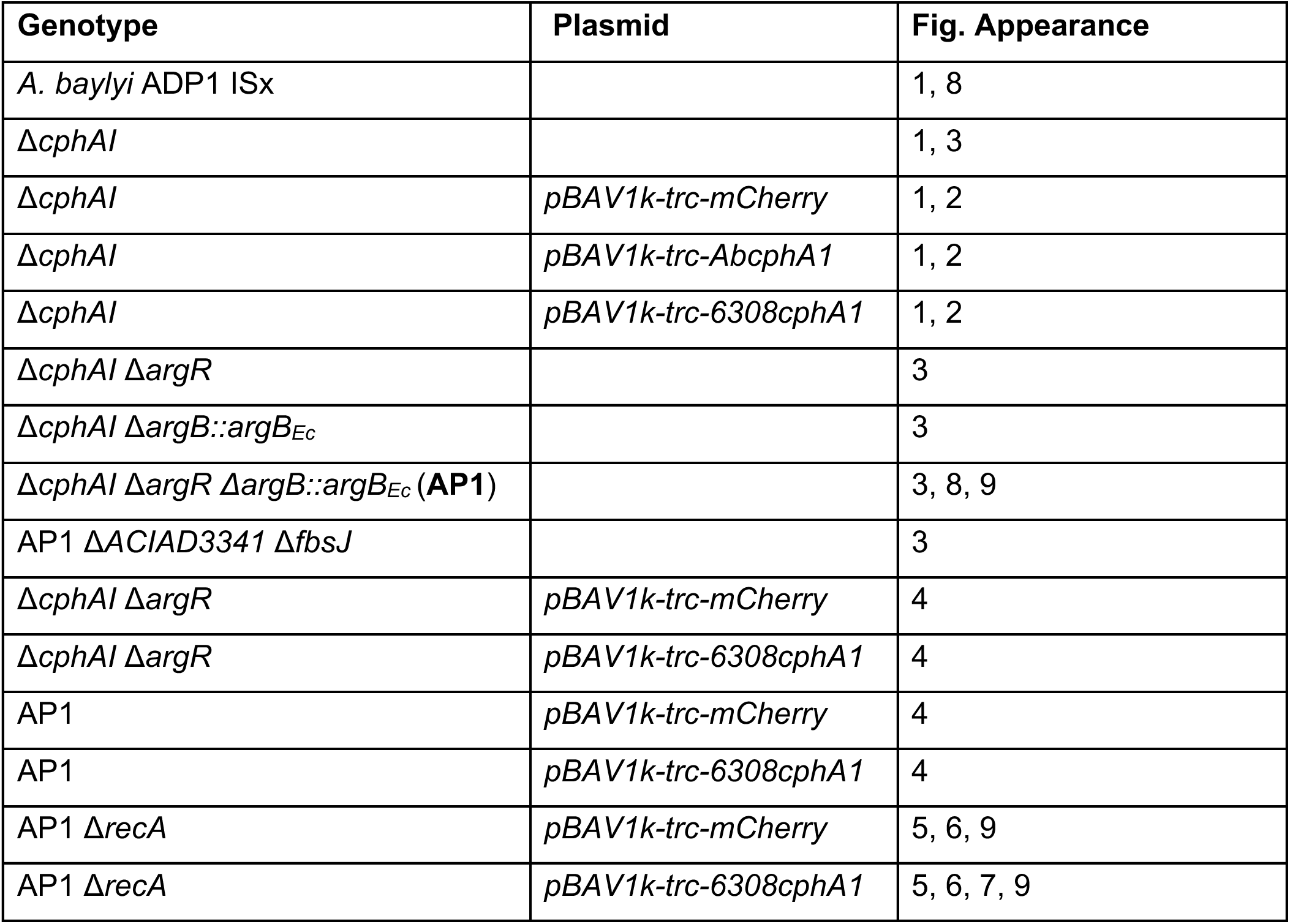
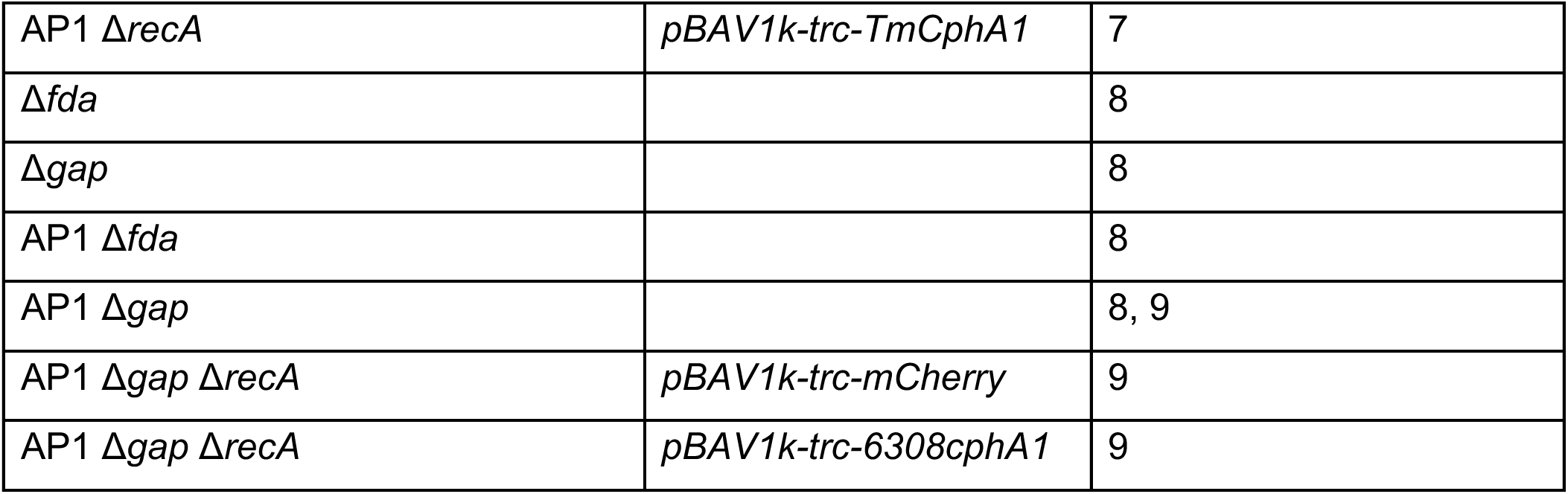
Strains used in this study.

Genomic knockouts and knock-ins in of *A. baylyi* were constructed as described previously^20^. Briefly, knockout constructs consisted of a *kanR-lacI-Trc-mCherry-rrnB* cassette flanked by 500 bp homology arms. For transformation, 70 μL of overnight culture was diluted into 1 mL of fresh LB, mixed with 100-200 ng of insertion cassette, and incubated with shaking for 3 h before selection on LB-kanamycin plates. Scarless integration fragments were similarly constructed with 500 bp homology arms upstream and downstream of the target site. Transformations were performed with 500-1000 ng of both the scarless integration fragment and the counterselection plasmid *RSF1010-T5-sgRNA(KanR)-TDK-Cas9*, followed by 6 h of incubation and selection on LB-spectinomycin plates. All constructs were validated by colony PCR and sequencing. To ensure the stability of plasmids carrying *kanR*, knockouts of the recombinase *recA* were generated via targeted integration of the spectinomycin resistance gene, *specR*.

### 2.3 Cyanophycin Cultivation under Varying Conditions

Individual colonies were inoculated into either 25 mL LB medium in 125 mL unbaffled Erlenmeyer flasks or 50 mL LB in 250 mL unbaffled Erlenmeyer flasks and grown overnight in biological triplicate. To generate sufficient biomass for experiments involving Δ*recA* mutants, cultures were scaled to 200 mL in 1 L flasks and allowed to continue growing for an additional 24 h when necessary. Unless otherwise noted, all cultures were incubated at 30 °C with shaking at 250 rpm. Cultures containing phosphate were directly inoculated to an OD600 of 0.05 for glutamate-based media or 0.2 for acetate-based media. Induction was performed with 1 mM isopropyl β-D-1-thiogalactopyranoside (IPTG) at a target OD600 of 0.4-0.6. As will be described in subsequent sections, cultivation conditions were frequently modified to accommodate specific production-phase interventions; however, they are summarized briefly here. Phosphate-limited cultures were induced after a fixed 6 h incubation. Cultures subjected to temperature shifts were pre-equilibrated to the target temperature for 10 min prior to induction. Chloramphenicol was added concurrently with IPTG, and Δ*gap* strains were supplemented with fructose as described in section 3.9. Where indicated, culture pH was manually adjusted with either HCl or NaOH at 24 h intervals.

### 2.4 Biomass Harvesting and Cell Dry Weight Determination

Cell cultures were transferred to 50 mL conical tubes and centrifuged (4 °C, 5,000 x g) for at least 10 min. Cell pellets were washed with once with chilled nanopure water and resuspended with fresh nanopure water at volume equal to one-fifth or one-tenth the initial culture volume. When necessary, samples were stored at -20 °C after these preparatory steps were completed. Cell dry weight was determined by dehydrating a fixed volume of this slurry at 80 °C for at least 24 h.

### 2.5 Cyanophycin Extraction for Sakaguchi Quantification and SDS-PAGE Analysis

Harvested and washed cell slurries were acidified to a final concentration of 0.1 M HCl and gently agitated at room temperature for 1 h. Following incubation, the slurry was centrifuged (17,000 x g, 2 min) and the “extracted” supernatant collected for Sakaguchi quantification or SDS-PAGE analysis. Where indicated, the insoluble pellet remaining after acid extraction was resuspended in the original volume of water and loaded as “unextracted” SDS-PAGE samples.

### 2.6 Cyanophycin Extraction for Gravimetric Quantification and SDS-PAGE Analysis

Using OD600 values and a g CDW/L per OD600 conversion factor of 0.532, harvested and washed cell slurries were diluted to an estimated concentration of 10 mg CDW/mL, acidified to a final concentration of 0.3 M HCl, and boiled at 95 °C in capped HPLC vials for 30 min. Following incubation, samples were centrifuged (17,000 x g, 2 min), and the resulting supernatant was collected as the “extracted” cyanophycin fraction. Where indicated, the insoluble pellet remaining after acid extraction was resuspended in the original volume of water and loaded as the “unextracted” fraction for SDS-PAGE analysis.

To approximate the amount of base required for rough neutralization to pH 6-8, a larger-volume acid extract was titrated with NaOH via pH probe. The smaller-volume acidic supernatants were then neutralized using a proportional volume of 1 M NaOH and subsequently adjusted more precisely by adding 200 μL of 1 M Tris-HCl (pH 7.2) per 1 mL of the initial extracted slurry. Neutralized samples were incubated overnight at 4 °C, and the precipitated material was recovered by centrifugation as the “insoluble phase.” The remaining supernatant was mixed with two volumes of -80 °C ethanol, incubated at -20 °C for ≥1 h, and centrifuged (5,000 × g, 15 min or 17,000 × g, 5 min). The resulting solids were collected as the “soluble phase”. Each fraction was freeze-dried (0.04 mPa, 24 h) prior to weighing. To account for weight loss due to dehydration of the microcentrifuge tubes during freeze-drying, 2-4 empty tubes were desiccated alongside the samples, and their average mass loss was added back to the measured sample weights.

For SDS-PAGE analysis, each fraction from a given strain was resuspended in 0.1 M HCl to a target concentration of 10 mg CDW/mL, calculated using the measured CDW values rather than the OD600-derived estimates used earlier. For example, if analysis revealed that 1 mL of the original biomass slurry contained 9.3 mg CDW, all fractions derived from that sample (unextracted, insoluble phase, soluble phase) were each resuspended in 930 μL of 0.1 M HCl, even though the individual fraction masses were each necessarily less than 9.3 mg. This normalization allows direct comparison of relative cyanophycin concentrations between fractions and with whole-cell lysates, which were loaded at 50 mg cell wet weight/mL, approximately equivalent to 10 mg CDW/mL.

### 2.7 Whole Cell Lysate Preparation and SDS-PAGE Analysis

Whole-cell lysates were prepared by pelleting 1-2 mL of cell culture by centrifugation (5,000 × g for 10 min or 17,000 × g for 2 min) and resuspending the pellets in Takara xTractor Buffer to an approximate concentration of 50 mg cell wet weight/mL. Suspensions were incubated at room temperature for 10 min. Aliquots (20 μL) of each sample type (whole-cell lysates, acid extracts, cyanophycin fractions, etc.) were combined with 20 μl of 2x Laemmli buffer containing β-mercaptoethanol (Millipore Sigma; Burlington, MA), heated at 95 °C for 10 min in PCR tubes, and loaded onto precast 11.5% SDS-PAGE gels (Bio-Rad; Des Plaines, IL). Electrophoresis was performed at 120 V for 60 min. Gels were stained with InstantBlue Coomassie Protein Stain (Abcam; Cambridge, UK) for at least 60 min prior to imaging.

### 2.8 Arginine Determination via Sakaguchi Reagent

The Sakaguchi reagent was used to quantify free arginine, arginine in native host proteins, and arginine in cyanophycin. Reagent preparation and assay setup were adapted from previously described methods with minor modifications^21^. Reagent A was prepared by dissolving 300 mg KI in 100 mL nanopure water. Reagent B was prepared by dissolving 2 g potassium sodium tartrate in 100 mL of 5 M KOH. Separately, 100 mg 2,4-dichloro-1-naphthol was dissolved in 180 mL ethanol and slowly added to the tartrate solution under stirring. The direct addition of molecular-grade NaOCl (9.33 mL, 5%) to Reagent B was discontinued due to elevated background absorbance. Reagent C was prepared by diluting 5% molecular-grade NaOCl tenfold with nanopure water. For each batch of Reagent B prepared, an optimal volume of Reagent C was determined by titration using 100 μg/mL L-arginine standards to achieve maximum signal intensity after a 10 min incubation. For analysis, 33 μL of Reagent A, 33 μL of sample, and 100 μL of Reagent B were combined in a 96-well plate. Samples were incubated for 1 h prior to Reagent C addition to allow peptide hydrolysis; for free arginine quantification, a 10 min incubation was used. Absorbance at 520 nm was measured 10 min after addition of Reagent C. Arginine standard reactions (0-100 μg/mL L-arginine) were prepared using either nanopure water or 0.1 N HCl to match the matrix of each sample and included on every assay plate.

### 2.9 Plate Reader Measurements

Optical density and fluorescence were measured in 96-well plates using a BioTek Synergy H1 microplate reader (Agilent; Santa Clara, CA). Optical density was measured at 600 nm on 300 μL samples diluted to below 1.0. *mCherry* fluorescence was measured using excitation and emission wavelengths of 585 nm and 615 nm, respectively, under maximum gain. Sakaguchi reactions (approximately 200 μL total volume) were analyzed at 520 nm.

### 2.10 HPLC Quantification of Acetate & Nitrate

Samples were clarified by centrifugation (5,000 x g for 10 min or 17,000 x g for 2 min) followed by filtration through 0.2 μM cellulose acetate membranes. Separation was carried out on a Bio-Rad Aminex HPX-87H column (300 x 7.8 mm) using 0.007 M H2SO4 as the mobile phase at a flow rate of 0.6 mL/min. An Agilent 2000 Series HPLC system equipped with diode array detection (DAD) at 210 nm was used for quantification. Acetate was measured without dilution (retention time: 15.2 min) using sodium acetate standards ranging from 1.69 mM to 169 mM. Nitrate samples were diluted 333-fold and quantified using potassium nitrate standards ranging from 150 μM to 600 μM (retention time: 6.1 min). Measurements of initial acetate and nitrate concentrations were performed in technical duplicate.

### 2.11 Ammonium Quantification

Ammonium concentrations were determined using Ammonia TNTplus vial tests (Hach; Loveland, CO) and a Hach DR3900 spectrophotometer. Clarified culture supernatants were diluted 50-fold to fall within the manufacturer’s specified detection range (2-47 mg/L NH4-N). Samples were incubated for 15 min at room temperature before measurement. Initial ammonium concentrations were measured in technical duplicate.

### 2.12 Flux Balance Analysis (FBA)

FBA was conducted using a previously compiled genome-scale model^22^ with minor modifications^23^. Fructose import and primary catabolism were absent in this model, so reactions representing fructose transport into the cytosol and a lumped reaction converting fructose to fructose-1,6-bisphosphate at the cost of 2 ATP molecules were incorporated. The finalized model is provided as a supplementary file. Simulations were performed using a fixed medium composition, with uptake rates of 1,000 mmol/h/g CDW for sulfate, molecular oxygen, inorganic phosphate, iron (II), sodium, and water.

## 3. Results & Discussion

### 3.1 Recombinant Cyanophycin Synthase Enables Cyanophycin Synthesis in Phosphate-Replete Conditions

Both qualitative and quantitative methods were employed to evaluate in vivo cyanophycin production. Qualitative analysis was performed using SDS-PAGE electrophoresis, which, when loading a fixed quantity of biomass per lane, provides a de facto illustration of specific cyanophycin content. Quantitative analysis was performed by extracting intracellular cyanophycin with HCl and measuring arginine in the liquid phase using the colorimetric Sakaguchi reagent. To account for arginine not derived from cyanophycin, cyanophycin-negative control strains were routinely cultured in parallel and subjected to the same extraction and quantification steps. Because these negative controls consistently yielded nonzero arginine values, this signal in *cphA1*-expressing strains is more accurately referred to as the *cyanophycin-enriched fraction* rather than purified cyanophycin. Since arginine is a principal component of cyanophycin, the difference in specific arginine content of these fractions was used as a proxy for cyanophycin concentration. Using these methods, we sought to reproduce previously reported effects of media composition on cyanophycin accumulation in *A. baylyi*.

Phosphate limitation constitutes one of the two previously cited requirements for robust cyanophycin production in wild type *A. baylyi* ADP1^16^. To verify this phenotype, cyanophycin accumulation was assessed in phosphate-limited (0 mM) and phosphate-replete (70 mM) mineral salts medium (MSM). Consistent with prior findings, phosphate limitation resulted in a clear qualitative increase in cyanophycin accumulation (**Fig. 1A**) and a corresponding quantitative increase in the arginine content of the cyanophycin-enriched fraction (+4.7% CDW; **Fig. 1B**).

**Figure 1.**
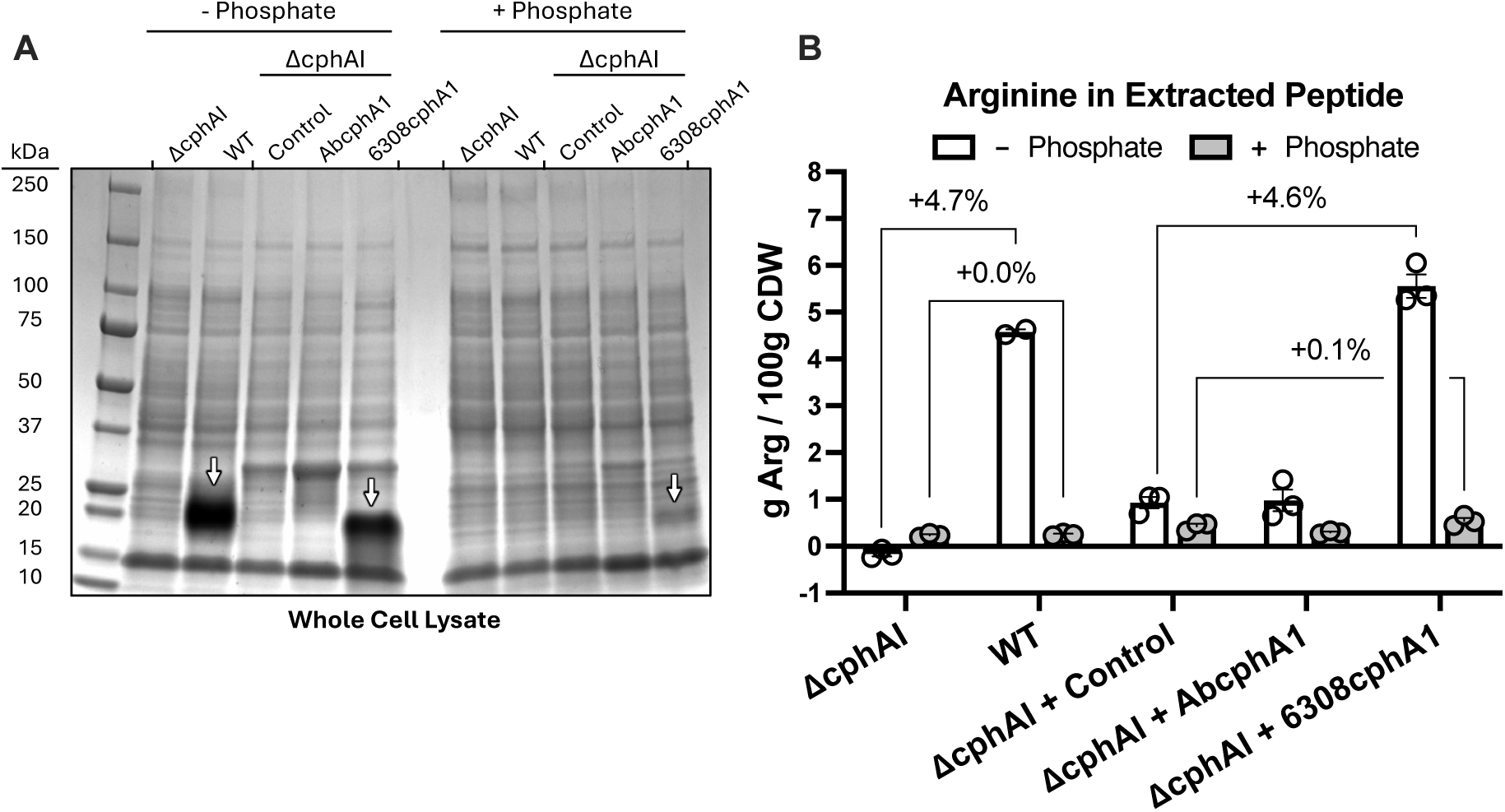
Recombinant expression of cyanophycin synthetase from *Synechocystis* sp. 6308 (6308cphA1) enables cyanophycin accumulation in phosphate-replete medium. **(A)** SDS-PAGE of whole-cell lysates from cultures grown in MSM containing 75 mM arginine, 10 mM ammonium sulfate, 50 mM MOPS, and either 0 mM (-Phosphate) or 70 mM (+Phosphate) total phosphate. To differentiate cyanophycin from non-cyanophycin components, *mCherry* was expressed as a control protein for comparison with strains heterologously expressing *cphA1*. Cyanophycin is indicated by white arrows. **(B)** Biomass-normalized arginine content of the samples shown in (A), quantified using the Sakaguchi reagent following incubation with 0.1 M HCl. After LB preculture, strains were resuspended and incubated in phosphate-free medium overnight, resuspended to an initial OD600 of 0.2 the following day, induced with 1 mM IPTG after 6 h, and harvested after ∼48 h at 30 °C. Error bars represent the SEM for n = 3 biological replicates, with the exception of the WT, -phosphate condition (n = 2) owing to sample loss.

A previous study identified the phosphate-responsive transcription factor *phoB* as a possible cause of this phosphate sensitivity, placing regulation of cyanophycin synthesis at the transcriptional level^17^. To test if transcriptional regulation of cyanophycin synthetase expression mediated the inverse relationship between cyanophycin production and phosphate concentration, the native *AbcphA1* and *cphI* genes were deleted at their native loci, and *AbcphA1* was reintroduced under the inducible *trc* promoter on a *pBAV1k* backbone. Induction of *AbcphA1*, however, resulted in impaired cyanophycin accumulation even under phosphate limitation, precluding interpretation of phosphate-replete results. To assess whether the effect was specific to the native enzyme, a second strain expressing a heterologous *cphA1* from *Synechocystis sp.* 6308 (*6308cphA1*) was constructed. Under phosphate limitation, this strain exhibited a comparable enrichment in arginine content (+4.6% CDW) relative to the wild type. In contrast to the wild type, however, it qualitatively produced cyanophycin under phosphate-replete conditions.

### 3.2 Arginine Remains a Key Bottleneck in Cyanophycin Synthesis

In addition to phosphate limitation, the supplementation of arginine as either the sole^16^ or secondary^17^ carbon source has been shown to be essential for robust cyanophycin production in *A. baylyi*. To test the stringency of this requirement in phosphate-replete conditions, a Δ*cphAI* strain expressing *6308cphA1* was cultivated using glutamate, the amino acid precursor to arginine, with or without arginine supplementation. In these conditions, the strain was only able to accumulate cyanophycin when arginine was supplemented (**Fig. 2**). Compared to the previous experiment, the inclusion of glutamate in phosphate-replete MSM produced a larger quantitative increase in arginine enrichment between strains expressing *6308cphA1* and the control (+2.1% CDW vs. +0.1% CDW).

**Figure 2.**
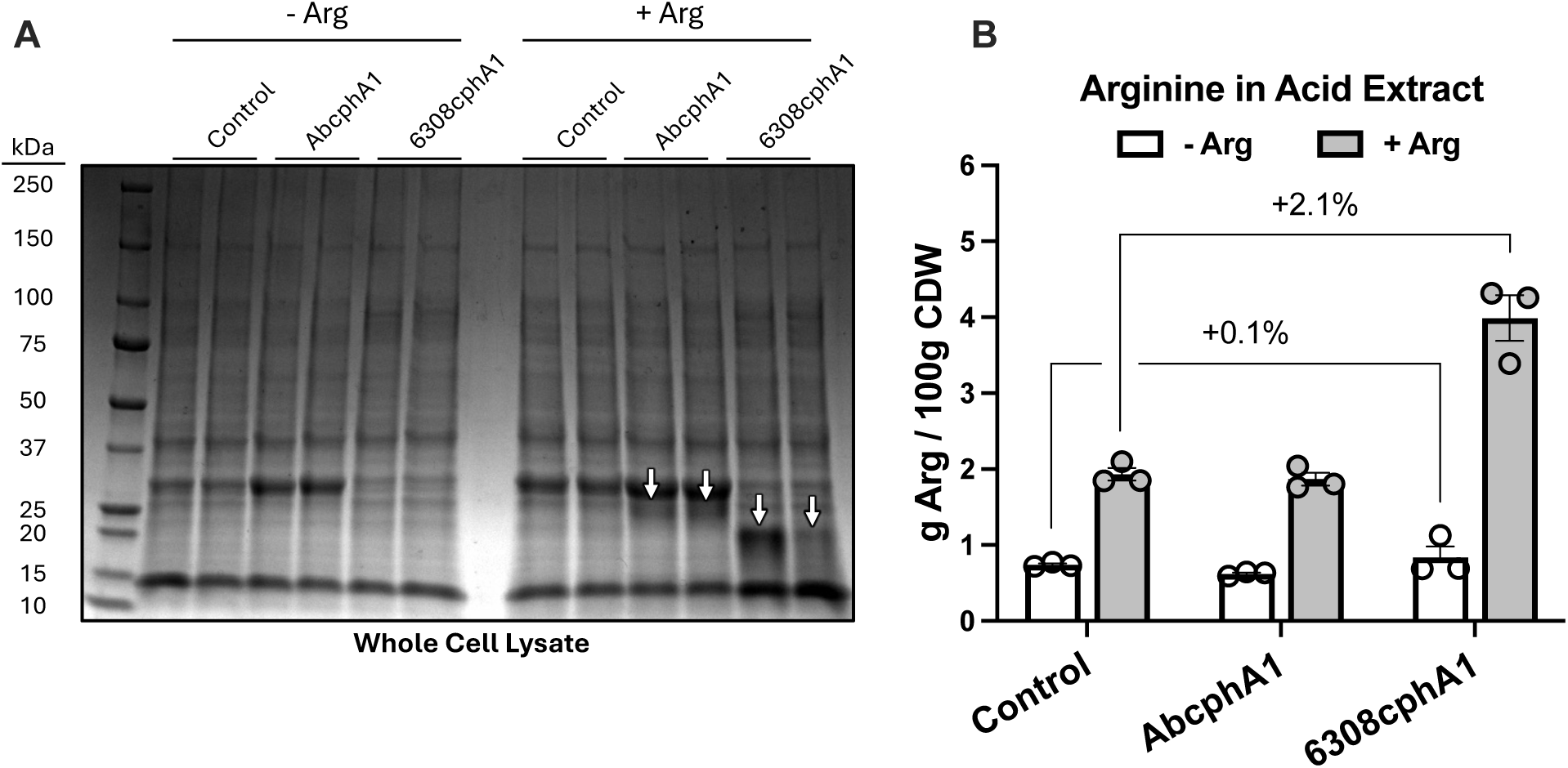
Arginine supplementation is essential for cyanophycin synthesis. **(A)** SDS-PAGE of whole-cell lysates from cultures grown in phosphate-replete MSM containing 60 mM sodium glutamate with or without 60 mM arginine. All strains carried the Δ*cphAI* genotype and expressed the indicated protein from plasmid *pBAV1k* under the *trc* promoter. *mCherry* was expressed as the control protein. **(B)** Biomass-normalized arginine content of the samples shown in panel A, quantified using the Sakaguchi reagent in technical triplicate following incubation with 0.1 M HCl. Cultures were grown for 54 h at 30 °C prior to harvest. Error bars represent the SEM for n = 3 biological replicates. An SDS-PAGE illustrating the effectiveness of cyanophycin extraction prior to Sakaguchi analysis is provided in **Fig. S1**.

### 3.3 *argR* and *argB* Regulate Arginine Biosynthesis in *A. baylyi*

Expression of *6308cphA1* in *A. baylyi* enabled cyanophycin accumulation under phosphate-replete conditions, but only when arginine was supplemented to the medium. Because supplying a major constituent of cyanophycin conflicts with our goal of recovering nitrogen from low-value “waste” substrates, we sought to eliminate this requirement by enhancing endogenous arginine biosynthesis.

Guided by prior studies, we first targeted the arginine-responsive transcriptional repressor *argR*, whose deletion has been reported to increase cyanophycin production in other systems^17^. Although the previous study examined Δ*argR* under phosphate-limited conditions, we found that the Δ*argR* mutant in our system also produced visibly higher cyanophycin levels than the wild type under phosphate-replete conditions without arginine supplementation (**Fig. S2**). However, the relative cyanophycin titer in the arginine-free condition appeared well below that achieved with arginine supplementation, indicating that arginine availability continued to limit polymer formation and that additional modifications to the arginine biosynthetic pathway (**Fig. 3A**) were required.

**Figure 3.**
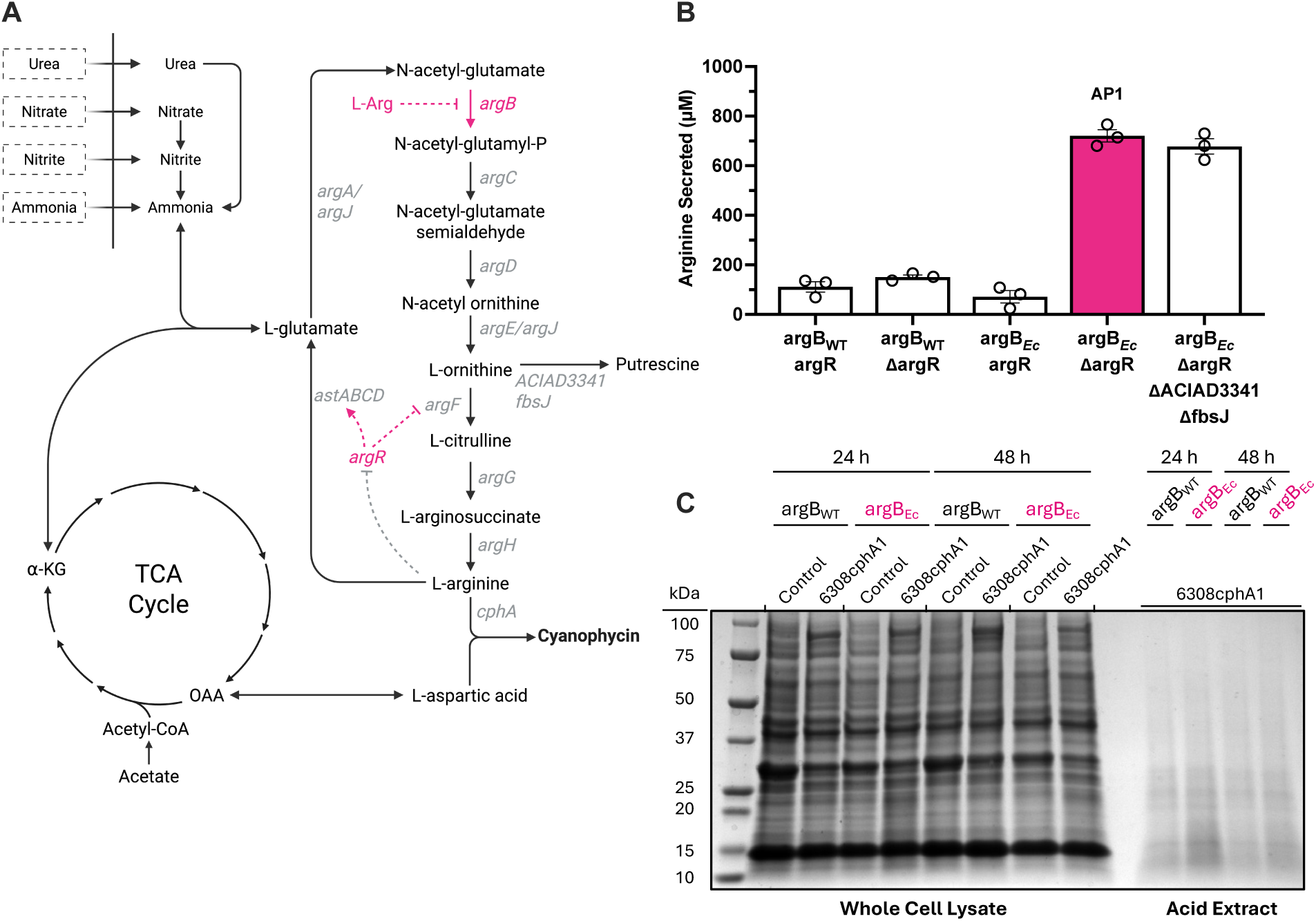
*argR* and *argB* regulate flux of arginine biosynthesis in *A. baylyi* ADP1. **(A)** Schematic of putative arginine biosynthesis pathways and regulatory nodes in *A. baylyi* ADP1. **(B)** Arginine production of engineered strains cultured in MSM with 60 mM glutamate at 30 °C for 72 h. *mCherry* was expressed as the control protein. Titers were quantified using the Sakaguchi assay in technical duplicate. Error bars represent SEM for n = 3 biological replicates. **(C)** SDS-PAGE of whole-cell lysates or acid extracts from cultures grown in phosphate-replete MSM containing 60 mM sodium glutamate. All strains carried the Δ*cphAI* Δ*argR* genotype. Cultures were harvested after ∼48 h at 30 °C.

Because cyanophycin accumulation reflects the combined action of both arginine synthesis and its diversion into polymer formation, we sought to more directly evaluate the effects of genetic modifications on arginine biosynthesis. To do so, we temporarily shifted from measuring intracellular cyanophycin to measuring arginine titers in culture supernatants as a proxy for biosynthetic flux. These assays were conducted in a Δ*cphAI* background to prevent biosynthesized arginine from being incorporated into cyanophycin, which could otherwise lead to underestimation of strain performance.

In bacteria such as *Escherichia coli* and *Corynebacterium glutamicum*, full derepression of arginine biosynthesis requires a secondary modification in addition to *argR* deletion, often targeting the first or second enzyme of the pathway to relieve allosteric inhibition^24^. Since ArgJ, the first enzyme in *A. baylyi*, is reportedly not feedback inhibited by arginine, we hypothesized that ArgB mediated the residual regulation. Replacement of wild type *argB* with an arginine-resistant homolog from *E. coli* (*argBEc*)^25^ confirmed this hypothesis: only the strain with both Δ*argR* and Δ*argB::argBEc* mutations displayed an enhanced capacity for arginine secretion, accumulating 721 ± 25 μM Arg (126 ± 4 mg/L) in the medium after 72 h (Fig. 3B). Deletion of putative ornithine decarboxylases *fbsJ* and *ACIAD3341* did not further enhance secretion, despite reports of a similar strategy improving arginine yields in *E. coli*^26^. The resulting Δ*cphAI* Δ*argR* Δ*argB::argBEc* strain, designated AP1 (Arginine Producer), was therefore selected for use in subsequent studies.

When *trc*-driven *6308cphA1* expression was reintroduced into AP1 via *pBAV1k*, however, the strain unexpectedly failed to visibly accumulate cyanophycin (**Fig. 3C**). This result prompted further investigation into alternative factors required for cyanophycin production.

### 3.4 Increased Arginine Synthesis Improves Cyanophycin Production at 12 °C

Previous work in *E. coli* indicated that cultivation at 12 °C enhances cyanophycin accumulation relative to 30 °C, particularly when expressing *6308cphA1*^9^. Guided by this observation, AP1 strains expressing either *6308cphA1* or the control protein *mCherry* were grown in phosphate-replete MSM containing acetate or glutamate without arginine supplementation. Cultures were first grown at 30 °C to an OD600 of 0.4-0.6, then shifted to 12 °C and induced 10 min later. Under these conditions, AP1 qualitatively accumulated more cyanophycin on both carbon sources compared to a non-arginine-secreting control (**Fig. 4A,B**). The potential mechanism by which growth at 12 °C enhances cyanophycin production is discussed further in section 3.7.

**Figure 4.**
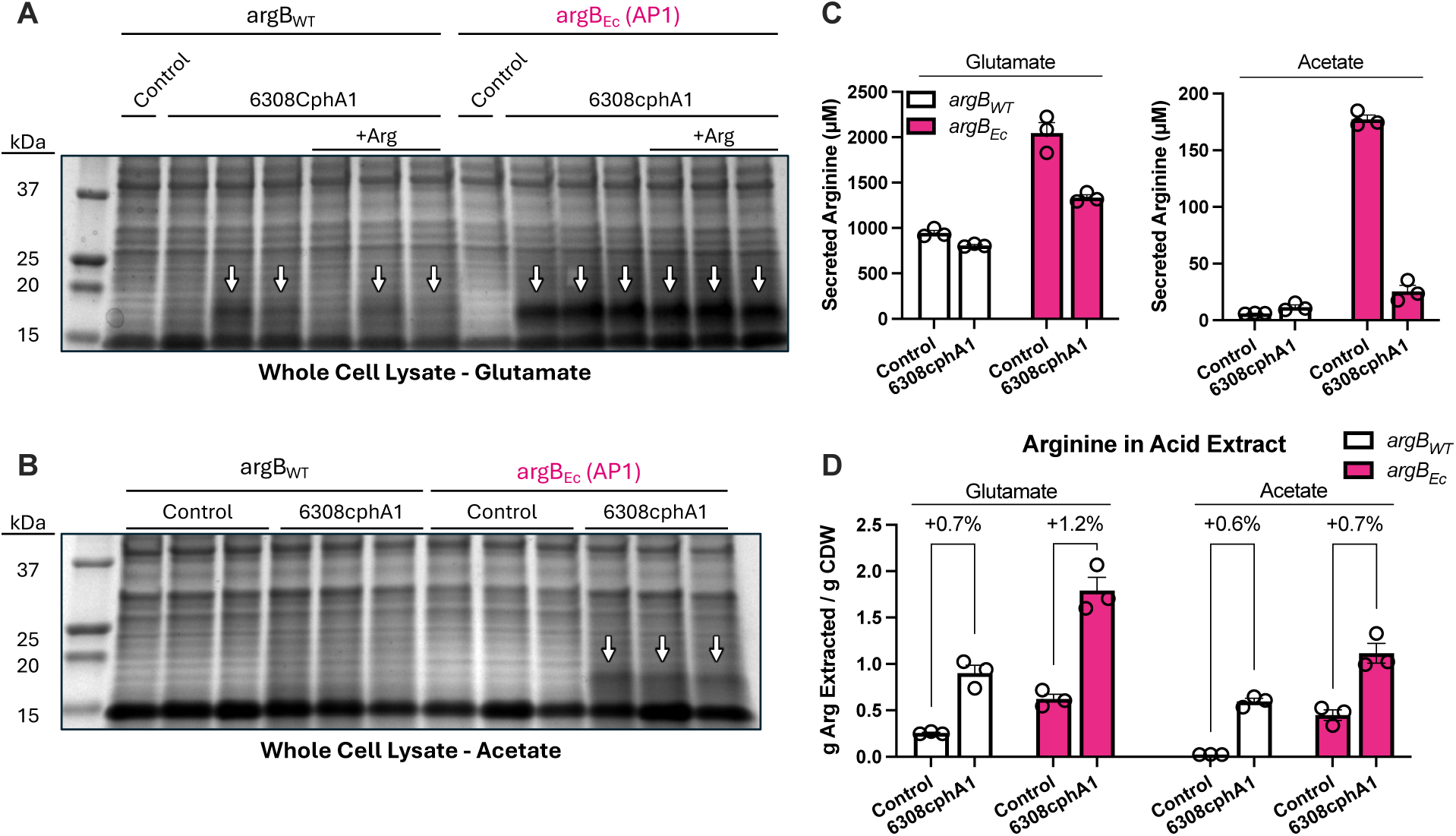
Arginine-producing strain AP1 enhances cyanophycin production without arginine supplementation at 12 °C. SDS-PAGE of whole-cell lysates from cultures grown in phosphate-replete MSM containing **(A)** 60 mM sodium glutamate (with 570 µM arginine added where indicated) or **(B)** 50 mM sodium acetate. All strains carried the Δ*cphAI* Δ*argR* genotype. *mCherry* was expressed as the control protein. Cultures were harvested 56 h (glutamate) or 23 h (acetate) post-inoculation. **(C)** Arginine titer in the medium at the time of harvest. **(D)** Biomass-normalized arginine content of the samples shown in panels A and B, quantified using the Sakaguchi reagent following at least 1 h incubation in 0.1 M HCl. Differences in specific arginine content are used as a proxy for cyanophycin concentration. All Sakaguchi measurements were performed in technical triplicate. Error bars represent SEM for n = 3 biological replicates.

To assess the extent to which biosynthesized arginine was incorporated into cyanophycin, extracellular arginine concentrations were measured at the end of the experiment (**Fig. 4C**). Assuming that both *mCherry*- and *6308cphA1*-expressing strains synthesized comparable amounts of arginine, increased cyanophycin production would be expected to correspond with lower extracellular arginine titers. Consistent with this expectation, AP1 expressing *mCherry* exhibited higher arginine titers than AP1 expressing *6308cphA1* for both carbon sources. Notably, the arginine titers of *6308cphA1*-expressing strains were similar to those of the non-secreting strains, particularly in the acetate-grown condition. These results suggest that cyanophycin synthesis in AP1 may remain limited by arginine availability such that further enhancements in arginine biosynthesis could translate to greater cyanophycin accumulation. However, this interpretation is tempered by the lack of a detectable difference in cyanophycin titer between *6308cphA1*-expressing strains grown on glutamate with or without exogenous arginine supplementation (**Fig. 4A**).

Independent of this interpretation, these results support the hypothesis that enhanced arginine biosynthesis correlates with increased cyanophycin production in the absence of exogenous arginine supplementation. This finding suggests that introducing *cphA1* into a strain already engineered for arginine overproduction could be a promising strategy for future cyanophycin cultivation, particularly given the low arginine titers of AP1 (<1 mM) compared to those of previously-engineered species (>500 mM^27^). This discrepancy, however, may reflect differences in cultivation conditions, medium composition, or the apparent absence of a dedicated arginine exporter in *A. baylyi*, in which case extracellular arginine titers may not accurately reflect raw biosynthetic arginine flux.

The qualitative enhancement in cyanophycin production by AP1 strains was reflected quantitatively in the glutamate condition, where the arginine enrichment in the acid-extract increased from +0.7% to +1.2% CDW (**Fig. 4D**). In contrast, arginine enrichment in acid extracts from acetate-derived cultures remained largely unchanged (+0.6% vs. +0.7%), despite apparent increases in cyanophycin observed via SDS-PAGE (**Fig. 4B**). We hypothesized that this discrepancy arose from incomplete extraction, with a significant portion of cyanophycin remaining in the biomass. Further analysis confirmed that, despite prior validation of the extraction protocol (**Fig. S1**), a substantial amount of cyanophycin was not recovered (**Fig. S3**). Accordingly, we next investigated our extraction procedure to facilitate complete cyanophycin removal in subsequent experiments.

### 3.5 Optimizing Cyanophycin Extraction

To improve cyanophycin recovery for our updated bioprocess, we systematically evaluated key parameters of the acid-based extraction procedure, including biomass concentration - which was previously not fixed between experiments - extraction duration, number of sequential incubations, acid concentration, and extraction temperature. Parameter ranges were guided by two prior studies that reported biomass loadings from 2-10 mg CDW/mL^16^ up to approximately 100 mg CDW/mL^28^, achieving near-complete extraction in either a single incubation or after three sequential incubations, respectively. Here, an incubation is defined as a step in which a fixed biomass concentration is treated with acid for a defined period, followed by centrifugation and collection of the supernatant for analysis. When additional incubations were performed, the pellet was resuspended in fresh acid and the process repeated. Using these bounds, we tested biomass concentrations of 2, 10, and 20 mg CDW/mL over 1, 2, or 3 incubations lasting 1, 3, or 6 h each (**Fig. 5A**).

**Figure 5.**
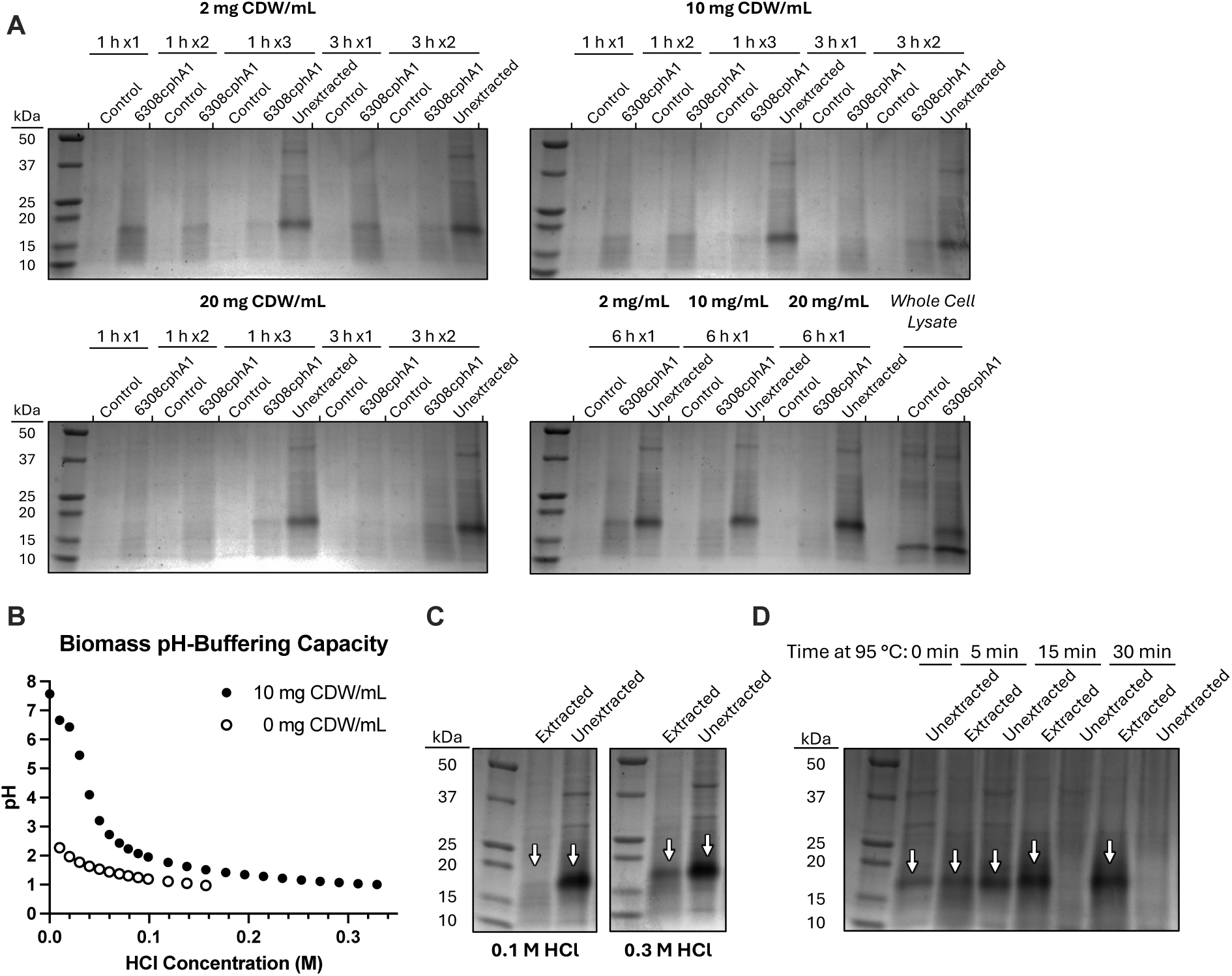
Boiling required to extract a recalcitrant fraction of cyanophycin from biomass. **(A)** Acid extraction parameters, including biomass concentration in the slurry (2, 10, 20 mg CDW/mL), extraction duration (1 h, 3 h, 6 h), and number of sequential extractions (1x, 2x, 3x), were varied and their impacts visualized via SDS-PAGE. A sample labelled “1 h x3”, for instance, underwent three one-hour incubations in a fresh volume of HCl each time and the supernatant from the third incubation is shown. To facilitate qualitative comparisons between samples of differing biomass concentrations, 10 and 20 mg CDW/mL samples were diluted to 2 mg CDW/mL prior to gel loading. *mCherry* served as the control protein. **(B)** Titration curve of 10 mg CDW/L cyanophycin-containing biomass with 10 M HCl. **(C)** SDS-PAGE showing the effect of HCl concentration on cyanophycin extraction using a 10 mg CDW/mL slurry. **(D)** SDS-PAGE showing the effect of boiling time on cyanophycin extraction using a 10 mg CDW/mL slurry with 0.3 M HCl. Cyanophycin is indicated by white arrows. Biomass samples were derived from a strain with the AP1 Δ*recA* genotype cultured in phosphate-replete MSM supplemented with 50 mM acetate and 50 mM ammonium sulfate. Cultivation conditions are identical to those detailed in Fig. 6.

Given the biomass concentrations tested, one would expect the lowest concentration (2 mg CDW/mL) to most effectively facilitate complete extraction, as the likelihood of approaching the cyanophycin solubility limit in the liquid phase is theoretically reduced. Similar to **Fig. S3**, however, a substantial amount of cyanophycin remained unextracted in 2 mg CDW/mL samples under even the most rigorous treatment regimens (1 h x3, 3 h x2, and 6 h x1). Although there appears to be a slight negative correlation between biomass concentration and extraction efficiency, these findings indicate that differences in biomass loading were unlikely to be responsible for the differential outcomes observed in **Fig. S1 and Fig. S3**. This conclusion is further supported by estimates of biomass concentrations in these experiments; using final OD600 and a conversion factor of 0.532 g CDW/L per OD600 the biomass loading was ∼30 mg CDW/mL for the successful extraction in **Fig. S1** and ∼11 mg CDW/mL for the unsuccessful extraction in **Fig. S3** - the opposite of what would be expected if biomass concentration were the critical variable. Nevertheless, a separate observation made during routine sample handling suggested a potentially related factor.

During SDS-PAGE sample preparation, we routinely observed that the Laemmli loading dye, which contains the pH-sensitive indicator bromophenol blue, turned yellow when mixed with acidified samples. For the 20 mg CDW/mL samples, however, the color change appeared only after multiple incubations, suggesting a possible buffering effect from the biomass (**Fig. S4**). This finding is consistent with the 1 h incubation results, where, unlike the 2 or 10 mg CDW/mL loadings, the third 1 h incubation of the 20 mg CDW/mL samples extracted more cyanophycin than the first or second. To quantify the extent of this buffering effect, we titrated 10 mg CDW/mL suspensions against water and found that approximately three times more HCl was required to reach pH 1 (**Fig. 5B**). Under the 0.1 M HCl conditions used in the initial extraction, the pH reached only ∼2. This finding is particularly relevant to high-throughput, low-volume workflows using fixed acid volumes too small for verification via pH probe. Increasing the acid concentration to 0.3 M HCl improved recovery but did not completely solubilize the recalcitrant fraction (**Fig. 5C**). Finally, we tested heat-assisted extraction by incubating 10 mg CDW/mL cell suspensions at 95 °C. Cyanophycin bands in the cell pellet disappeared completely after 15 min, and we extended the treatment to 30 min for routine use (**Fig. 5D**).

While previously-unfixed parameters such as biomass concentration and incubation pH had some effect on extraction efficiency, the ultimate requirement for boiling suggests that the differential outcomes in **Fig. S1** and **Fig. S3** were likely driven by differences in genotype (Δ*cphA1* versus AP1), carbon source (glutamate versus acetate), or cultivation temperature (12 °C versus 30 °C). Previous work has shown that similar process variables can alter cyanophycin composition, molecular weight, and solubility^29^. Because these properties likely influence granule morphology (e.g. volume-to-surface-area ratio), it follows that innate “extractability” may also be modulated by such process variables. These findings highlight a promising direction for future work aimed at elucidating how cellular and environmental factors shape the physical properties of cyanophycin granules and their compatibility with downstream recovery processes.

### 3.6 Cyanophycin Production from Ammonium, Nitrate, & Urea

Equipped with a more robust extraction protocol, we next evaluated the capacity of strain AP1 to grow and produce cyanophycin from carbon and nitrogen sources relevant to municipal wastewater, including acetate and propionate as carbon sources and ammonium, nitrate, nitrite, and urea as nitrogen sources. To ensure accurate quantification and maintain plasmid stability during these tests, two specific adjustments were made. First, the Sakaguchi-based assay was replaced with a gravimetric method to prevent free arginine produced by AP1 from contributing to apparent cyanophycin levels in the extracts. Briefly, acid extracts were neutralized to approximately pH 7.2, and the resulting precipitate was collected, dried, and weighed as the “insoluble” fraction. The remaining supernatant was then treated with two volumes of chilled ethanol, and the resulting precipitate was collected, dried, and weighed as the “soluble” fraction (Materials & Methods, 2.6). Like the previous quantification method, the amount of collected material was compared against that collected from a cyanophycin-negative control, and the difference was attributed to cyanophycin. Second, the recombinase gene *recA* was deleted to minimize the potential loss of plasmid-borne *6308cphA1* during cultivation, a phenomenon previously reported for heterologous enzyme expression in *A. baylyi* ADP1^30^.

Growth was demonstrated using acetate and propionate as carbon sources (**Fig. S5**), but for simplicity, acetate alone was used in all subsequent experiments. While nitrite did not support cell growth as sole nitrogen source, the other nitrogen species enabled both growth and the production of cyanophycin, comprising 18.9 ± 3.8% CDW from ammonium, 8.7 ± 3.8% CDW from nitrate, and 29.0 ± 3.3% CDW from urea (**Fig. 6A**). Following quantification, each fraction was resuspended and analyzed via SDS-PAGE for visualization (**Fig. 6B**). Acetate consumption was quantified under all three conditions, and inorganic nitrogen consumption was measured for ammonium and nitrate. Notably, the strain expressing *6308cphA1* consumed 2.9-fold (SEM ± 0.3) more nitrogen per unit biomass and 3.3-fold (SEM ± 0.1) more nitrogen per unit acetate than the control strain expressing *mCherry*, consistent with the value proposition of this process: the simultaneous removal of nitrogen for water quality purposes and production of a value-added product. This trend did not hold in the nitrate condition, however, where strains expressing *6308cphA1* consumed 2.4-fold (SEM ± 0.8) *less* nitrogen per unit biomass and 1.5-fold (SEM ± 0.4) *less* nitrogen per unit acetate than the control strain expressing *mCherry*. This result may be the consequence of nitrate disproportionally stressing nitrogen metabolism and overall cell viability in the cyanophycin producers: control strains grew to a final optical density 2.2-fold (SEM ± 0.2) greater than the cyanophycin-producing strains. In agreement with this hypothesis, nitrate supported the smallest per-cell cyanophycin titers of the three nitrogen sources tested. These results position waste streams rich in ammonium and urea as the most amenable to cyanophycin-mediated nitrogen recovery, with future work likely needed to optimize recovery from nitrate-rich sources.

**Figure 6.**
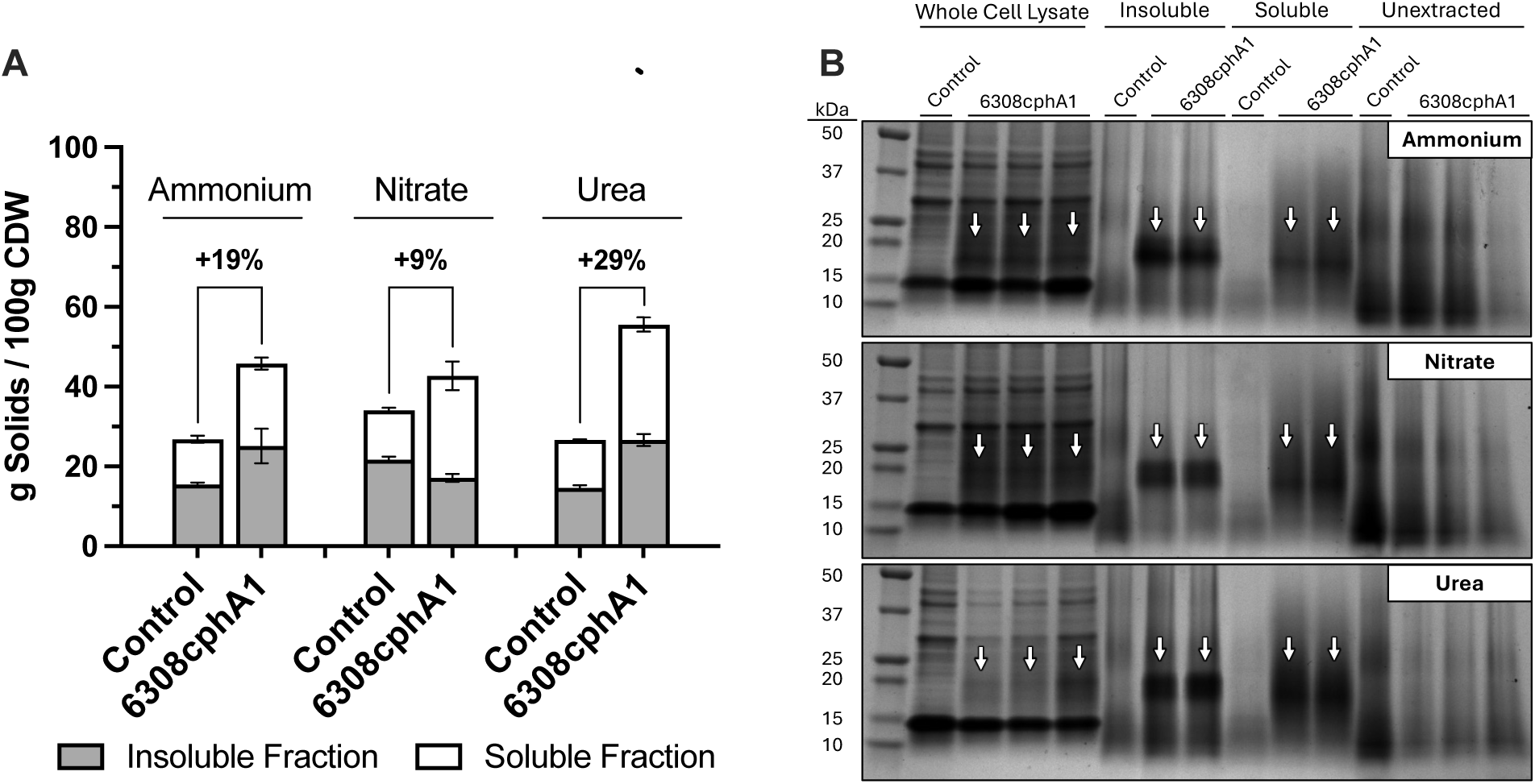
*A. baylyi* AP1 Δ*recA* can accumulate cyanophycin from simple carbon and reactive nitrogen sources. **(A)** Per-cell titers of cyanophycin-enriched fractions of biomass collected using the optimized acid extraction method and precipitation-based purification regimen. The difference between the sums of the soluble and insoluble fractions between Control- and *6308cphA1*-expressing strains is attributed to cyanophycin. *mCherry* served as control protein. **(B)** SDS-PAGE showing each fraction collected during the extraction and purification process. Cyanophycin is denoted by white arrows. All strains were cultivated in phosphate-replete MSM medium containing 50 mM acetate and either 50 mM ammonium sulfate, 100 mM potassium nitrate, or 50 mM urea. Cultures were moved to 12 °C 10 min before induction, the pH was adjusted to 7.00 ± 0.05 after 24 hours, and biomass was harvested 48 h post-inoculation.

### 3.7 Suboptimal Growth Conditions Correlate with Elevated Per-Cell Cyanophycin Titers

While these results demonstrate cyanophycin production 1) under phosphate-replete conditions, 2) without arginine supplementation, 3) using acetate as the sole carbon source, and 4) using wastewater-relevant nitrogen sources, the utility of this proposed bioprocess is severely limited by the 12 °C growth requirement. Wastewater temperatures vary seasonally and geographically, reaching temperatures as low as 6 °C during Wisconsin winters^31^ and up to 29 °C during Florida summers^32^. This temperature variability poses challenges for even contemporary BNR processes, due to the large, cost-prohibitive amounts of energy required to actively control influent temperature^33^. Consequently, similar constraints should be anticipated for cyanophycin-based nutrient recovery approaches.

We hypothesized that cultivation at 12 °C enhances cyanophycin accumulation at least in part by disproportionally slowing cell division relative to cyanophycin synthesis, effectively limiting losses in per-cell titer caused biomass dilution. To test this, AP1 Δ*recA* was exposed to three growth-rate limiting conditions: incubation with the ribosome-stalling antibiotic chloramphenicol^16^ (**Fig. 7A**), incubation at temperatures above and below the optimum (**Fig. 7B**), and culturing at elevated pH values >7.0 (**Fig. 7C**). All interventions reproduced the targeted phenotype to varying degrees. Increasing concentrations of chloramphenicol enhanced per-cell cyanophycin titers up to an apparent maximum of 5 µg/mL at 30 °C. Growth at 12 °C resulted in the highest accumulation for *6308cphA1*, whereas 37 °C was the only temperature at which *TmcphA1* (derived from the heterotroph *Tatumella morbirosei*) exhibited visible cyanophycin accumulation. Elevated pH (8.2 and 8.4) likewise enhanced cyanophycin production by *6308cphA1* at 30 °C. These findings are further supported by previous employments of phosphate-starvation^16^ - a condition also associated with impaired growth - to promote cyanophycin synthesis (**Fig. 1**). While moderate interventions improved cyanophycin accumulation, we found that overly aggressive conditions reduced titers, presumably by excessively compromising cellular homeostasis (e.g., 20 µg/mL chloramphenicol, cultivation at 40 °C). This suggests that a relative optimum exists for each intervention in which cell growth is slowed while key functionalities regarding precursor synthesis, *cphA1* expression, and cyanophycin polymerization are at least partially maintained.

**Figure 7.**
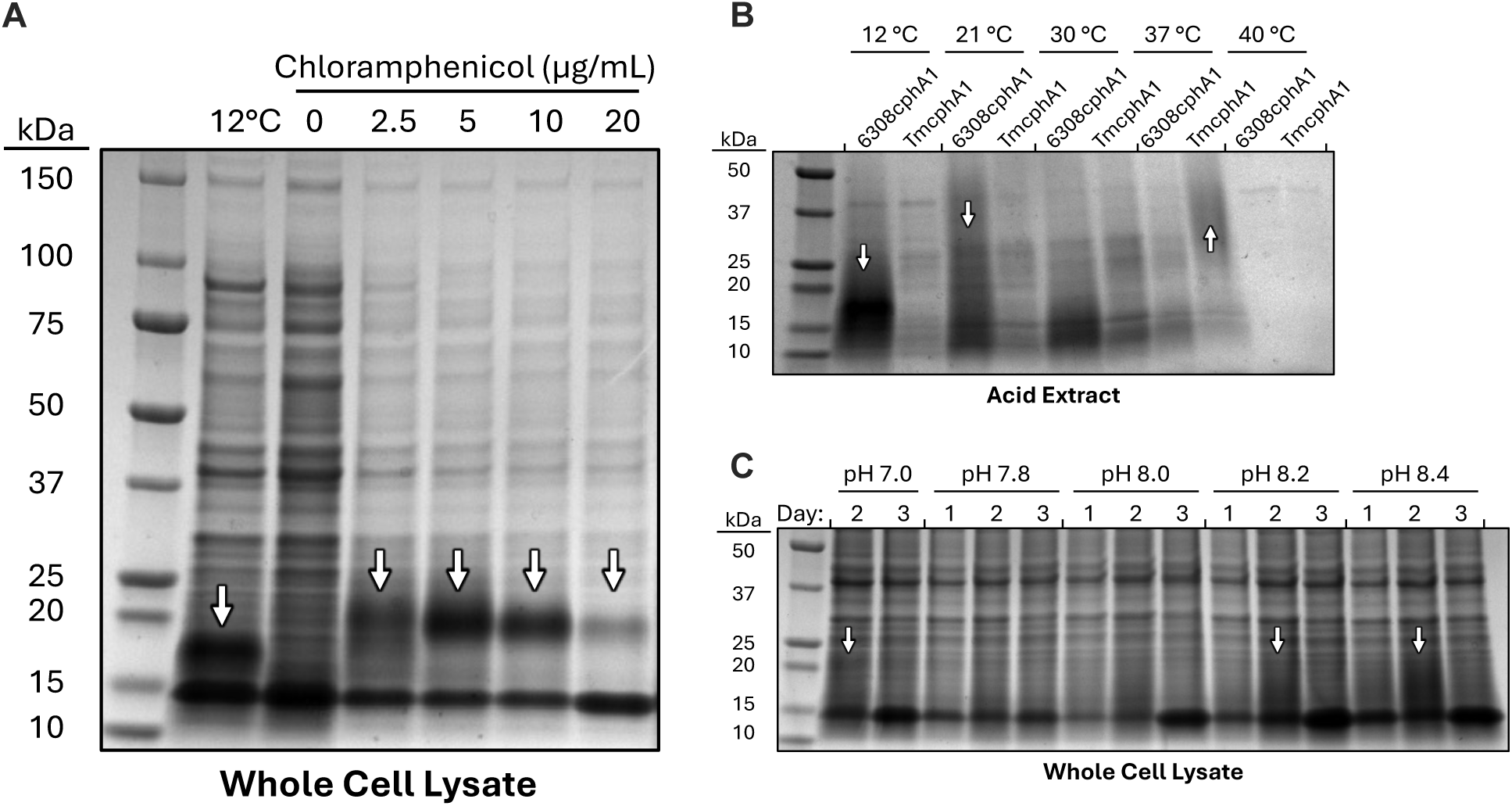
Conditions that reduce cell fitness are associated with enhanced cyanophycin production. SDS-PAGE showing the impact of different interventions on cyanophycin accumulation, including **(A)** addition of the antibiotic chloramphenicol, **(B)** cultivation at sub-optimal temperatures, and **(C)** cultivation at sub-optimal pH. All experiments were performed using AP1 Δ*recA* in phosphate-replete medium supplemented with 60 mM glutamate and 20 mM ammonium sulfate. Unless indicated otherwise, all strains expressed *6308cphA1* as described previously. Each intervention was applied once cultures reached a target OD600 of 0.4-0.6, immediately followed by addition of 1 mM IPTG. Samples were collected (A) ∼24 h post-inoculation, (B) ∼48 h post-inoculation, and (C) approximately every ∼24 h for 3 days. For (C), the pH was manually adjusted back to the setpoint every 24 h.

Although a more parsimonious explanation for the mechanisms underlying these observations may exist, these results demonstrate the utility of the dilution-from-growth hypothesis, particularly in predicting that temperature-independent interventions can enhance cyanophycin accumulation by AP1 Δ*recA* at temperatures above 12 °C. Despite this, these experiments do not provide a direct solution for improving compatibility with real-world deployment; aside from pH modulation, which had the most modest effect, interventions such as chloramphenicol supplementation are unlikely to be feasible in municipal wastewater applications for practical reasons.

### 3.8 Ä*gap* Mutation Induces Fructose Auxotrophy and Enables Control Over Cell Growth for Wastewater Applications

To selectively attenuate cell growth in municipal wastewater contexts, we explored the creation of an auxotrophic *A. baylyi* strain capable of growth only in the presence of acetate and a secondary carbon source. In this design, depletion of the secondary carbon source would prevent further growth. For practical feasibility, the secondary carbon source was required to be 1) relatively inexpensive and accessible, 2) largely absent from municipal wastewater, and 3) needed in minimal quantities relative to acetate.

Fructose meets the first two criteria; it can be sourced from high-fructose corn syrup in large volumes at relatively low-cost and is not considered a dominant source of carbon in municipal wastewater. Furthermore, previous work has demonstrated that fructose auxotrophy can be induced with only a single knockout (Δ*fda*)^22^. To estimate the amount of acetate that could be consumed per unit of fructose, we applied flux balance analysis. The results (**Table S3**) indicated that when grown on acetate and fructose, a Δ*fda* mutant would require a maximum of 6.9% of total carbon from fructose, whereas a Δ*gap* mutant, targeting the enzyme immediately downstream of *fda* in glycolysis, would require a maximum of 9.1% (**Fig. 8A**). These values compare very favorably with a previous acetate-gluconate partitioning strategy in *A. baylyi*, in which 51.3% of total carbon was derived from gluconate^34^, thus satisfying the third criterion.

**Figure 8.**
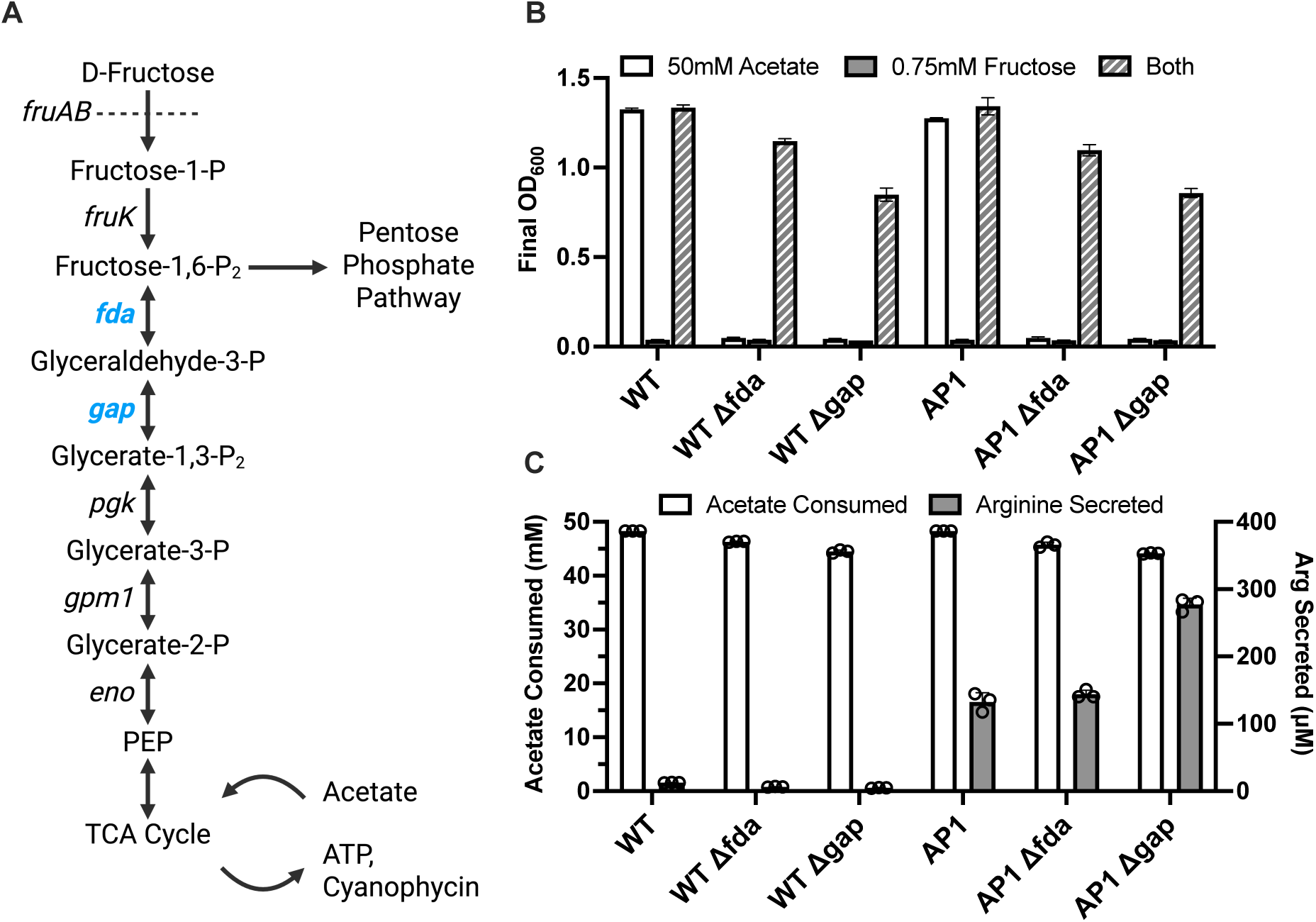
Δ*gap* mutation in AP1 correlates with the expected growth-stalled phenotype and enhanced arginine secretion. **(A)** Schematic for fructose catabolism in *A. baylyi* with knockout targets *fda* and *gap* highlighted in blue. **(B)** Final optical densities of Δ*fda* and Δ*gap* mutants grown on 50 mM acetate, 0.75 mM D-fructose, or both 50 mM acetate and 0.75 mM D-fructose at 30 °C after approximately 36 h post-inoculation. **(C)** Acetate uptake and arginine secretion for mutants supplemented with both 50 mM acetate and 0.75 mM fructose. Cultures were grown in 5 mL phosphate-replete MSM in 14 mL culture tubes. Samples were clarified by centrifugation at 17,000 × g and filtered through a 0.2 μm membrane. Arginine titers were quantified using the Sakaguchi assay in technical triplicate. Error bars represent SEM for n = 3 biological replicates.

Both Δ*fda* and Δ*gap* strains were cultured in MSM containing acetate, fructose, or both. As expected, neither strain grew unless both carbon sources were present (**Fig. 8B**). Based on acetate consumption data (**Fig. 8C**) and assuming complete catabolism of the supplemented fructose, carbon derived from fructose accounted for 4.6 ± 0.0% (WT) and 4.7 ± 0.0% (AP1) of total carbon for Δ*fda* mutants, and 4.8 ± 0.0% (WT) and 4.9 ± 0.0% (AP1) total carbon for Δ*gap* mutants. These values are consistent with the upper bounds calculated previously via flux balance analysis. Interestingly, AP1 Δ*gap* but not AP1 Δ*fda* exhibited improved extracellular arginine titers, motivating its selection for subsequent experiments.

### 3.9 Fructose Auxotrophy Enhances Per-Cell Cyanophycin Titers at 30 °C

We next sought to optimize the fructose feeding regimen for AP1 Δ*gap* to better support downstream cyanophycin production. Previous attempts to produce cyanophycin in AP1 grown on acetate and ammonium showed that arginine secretion by AP1 dropped to levels typical of non-secreting strains when *6308cphA1* was expressed (**Fig. 4C**), suggesting that arginine availability still limited cyanophycin formation even in AP1. We therefore hypothesized that the cultivation strategy maximizing the specific arginine secretion rate would be most likely to enhance specific cyanophycin titers at 30 °C. To test this hypothesis, we implemented a two-phase cultivation strategy in which the fructose feeding frequency was varied to identify the value that maximized arginine productivity.

In the first 24 h (Phase I), cultures were again grown in MSM containing 50 mM acetate and 0.75 mM fructose. Consistent with previous observations, AP1 Δ*gap* reached a lower optical density than AP1 but secreted more than twice the amount of arginine while consuming a similar quantity of acetate (**Fig. 9A**). After 24 h, AP1 Δ*gap* cultures were supplemented with an additional 100 mM acetate and subdivided for Phase II, during which distinct fructose feeding regimens were tested over the next 24 h (**Fig. 9B**). The control received no additional fructose, while the experimental groups received 0.4 mM total fructose administered in either one, two, or four regularly-spaced doses. Interestingly, the control culture continued to consume acetate and maintained specific arginine secretion rate comparable to Phase I, despite a continuous decline in optical density (**Fig. S6**). Cultures fed 0.1 mM fructose four times exhibited a 30% increase in specific arginine secretion rate compared to cultures given a single 0.4 mM fructose pulse. We therefore adopted this multi-pulse regimen to evaluate the effect of the Δ*gap* mutation on cyanophycin production at 12 °C and 30 °C (**Fig. 9C**).

**Figure 9.**
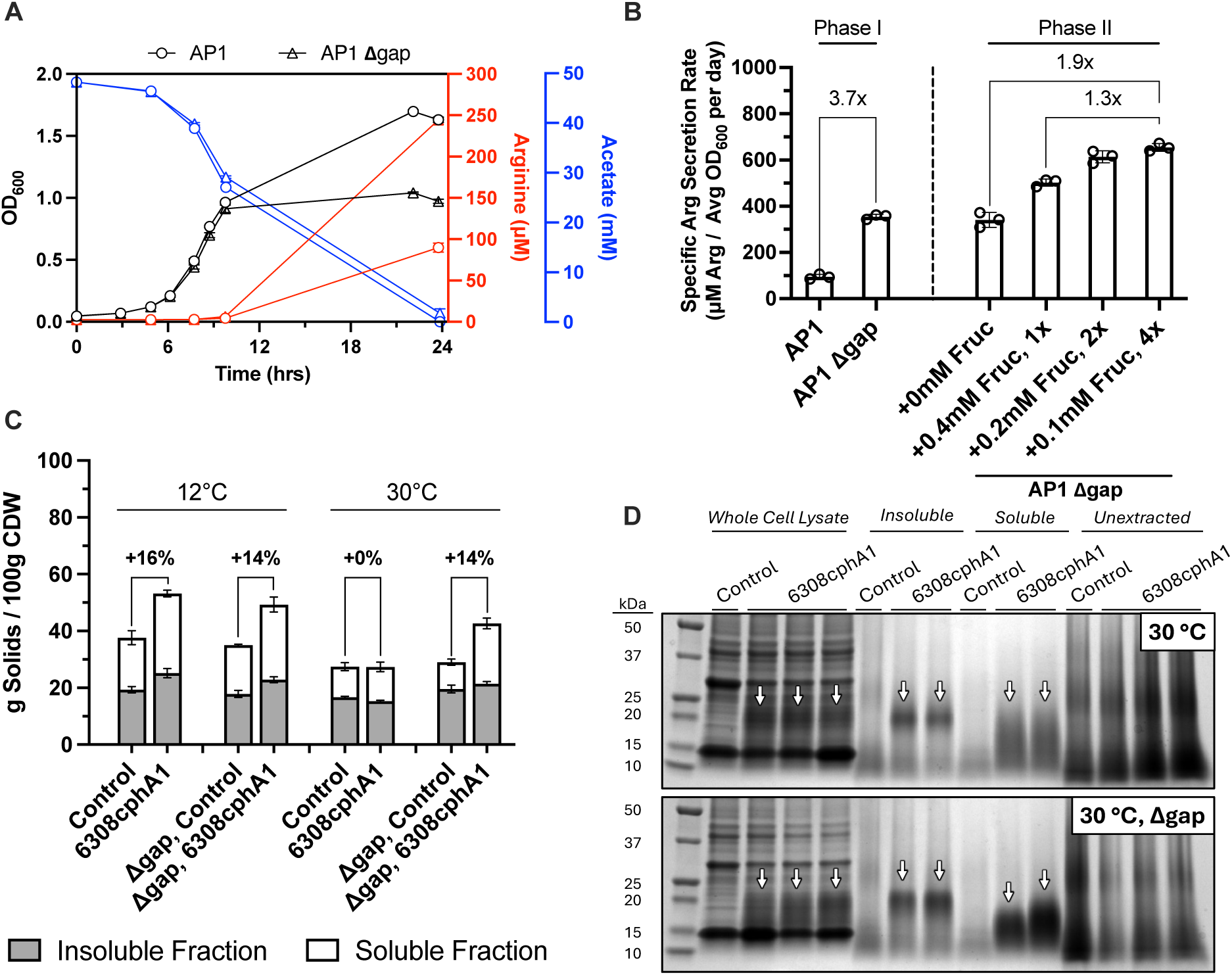
Δ*gap* mutation enhances cyanophycin production in AP1 Δ*recA* at 30 °C. **(A)** OD600, arginine secretion, and acetate uptake for AP1 and AP1 Δ*gap* over 24 h (Phase I). **(B)** Specific arginine production by AP1 Δ*gap* under different fructose feeding regimens over an additional 24 h (Phase II). Only arginine produced during this second 24 h period was used for calculations, and average OD600 was computed from measurements taken every 6 h. Phase I cultures were grown in 100 mL phosphate-replete MSM in biological triplicate, and 5 mL was transferred to 14 mL culture tubes for each Phase II condition. OD600, arginine secretion, and acetate uptake for Phase II provided in **Fig S6**. **(C)** Per-cell titers of cyanophycin-enriched biomass fractions. Differences between the sums of soluble and insoluble fractions in *mCherry*- and *6308cphA1*-expressing strains are attributed to cyanophycin. **(D)** SDS-PAGE of each fraction collected during the extraction and purification process. Cyanophycin is indicated by white arrows. All strains carried the AP1 Δ*recA* genotype and were grown in phosphate-replete MSM initially containing 50 mM sodium acetate, 0.75 mM D-fructose, and 50 mM ammonium sulfate. At 24 h, an additional 100 mM sodium acetate was added, culture pH was adjusted to 7.00 ± 0.05, and 0.1 mM D-fructose was added every 6 h. Biomass was harvested 48 h post-inoculation. Medium samples were clarified by centrifugation at 17,000 × g and filtered through a 0.2 μm membrane. Arginine titers were quantified using the Sakaguchi assay in technical triplicate. Error bars represent SEM for n = 3 biological replicates.

At 12 °C, the introduction of Δ*gap* had minimal effect on per-cell cyanophycin titers, producing a slight decrease in measured cyanophycin accumulation (16% CDW, *gap*; 14% CDW, Δ*gap*). In contrast, at 30 °C, significant quantities of cyanophycin were measured only in the Δ*gap* mutant (0% CDW, *gap*; 14% CDW, Δ*gap*). Furthermore, this amount mirrors that measured in the 12 °C, Δ*gap* condition, consistent with our goal of uniform cyanophycin production across wastewater-relevant temperatures. While the appearance of visible cyanophycin bands in the 30 °C, *gap* sample appear to disagree with a calculated 0% cyanophycin titer, the visible increase in the soluble fraction for Δ*gap* versus *gap* samples is consistent across both quantitative and qualitative methods and supports the broader notion that Δ*gap* qualitatively enhances cyanophycin production at 30 °C. This finding also suggests that the quantitative approach employed likely underestimates the actual contribution of cyanophycin to CDW.

Curiously, in this experiment the Δ*gap* mutation did not facilitate the >2-fold enhancement in arginine titers observed previously (**Fig. S7** versus **Fig. 8C**, **Fig. 9A**). For control strains not producing cyanophycin, we expected Δ*gap* strains to produce more arginine than their *gap* counterparts. At 12 °C, however, arginine titers were comparable with or without the Δ*gap* mutation (**Fig. S7**). After normalizing for acetate consumption to account for significant differences in growth between *gap* and Δ*gap* strains, this trend also held at 30 °C. Despite this discrepancy, at 30 °C Δ*gap* but not *gap* reproduced the previously observed pattern whereby *6308cphA1*-expressing strains secreted less arginine than the cyanophycin-negative control. Collectively, these observations are consistent with the hypothesis that the Δ*gap* mutation enhances cyanophycin accumulation by increasing the fraction of arginine routed into cyanophycin rather than by increasing the total arginine pool. This suggests that cyanophycin accumulation may now instead be limited by the capacity of expressed *cphA1* to incorporate the available arginine, an obstacle that could potentially be addressed by increasing *cphA1* expression. This is supported by the ability of an engineered *R. eutropha* strain to reach a record 53% CDW cyanophycin using only a plasmid addiction system^11^, a modification likely intended to enhance *cphA1* expression alone.

The inconsistency in Δ*gap*-mediated arginine secretion across experiments raises questions concerning the underlying mechanism. One possibility is that differences in innate growth rate drive differences in arginine secretion, as a mutant growth-limited by fructose uptake might be able to divert a larger fraction of total carbon to arginine synthesis versus biomass prior to carbon depletion. However, this explanation is inconsistent with the data, as AP1 *gap* and AP1 Δ*gap* exhibited identical growth trajectories during exponential phase growth (**Fig. 9A**). A second hypothesis is that Δ*gap* promotes premature cell lysis, releasing a disproportionate amount of free arginine or arginine-containing proteins into the growth medium and skewing Sakaguchi measurements. In argument against this theory, the highest arginine titers occurred under conditions where bulk cell death was not observed (0.1 mM fructose, **Fig. S6**). The arginine-enhancing effect also appeared genotype-specific, as Δ*gap* and Δ*fda* mutations in the wild-type background (i.e. not AP1) did not increase measured arginine titers (**Fig. 8C**). The tendency of AP1 Δ*gap* to consume acetate during regimes of apparent cell death (0 mM fructose, **Fig. S6**), however, suggests that the cell lysis hypothesis may not be mutually exclusive with alternative explanations for the enhanced arginine secretion.

Taken together, the data support a model in which Δ*gap* strains function largely as intended: upon fructose depletion, cells continue metabolizing acetate without net growth, with periodic low-level fructose supplementation reactivating metabolism and enhancing product formation. More broadly, this work demonstrates that a single knockout (Δ*gap*) can facilitate growth-production carbon partitioning, requiring minimal fructose to enable acetate consumption while maintaining metabolic activity even after bulk growth has ceased. Critically, these traits highlight the potential of Δ*gap* to support growth-partition strategies for other natural products in *A. baylyi* derived from TCA cycle intermediates, such as wax esters.

### 3.10 Limitations

Our efforts to engineer *A. baylyi* culminated in a strain capable of producing cyanophycin from wastewater-relevant carbon and nitrogen sources and across a range of temperatures representative of municipal wastewater treatment plants. While we believe this work represents a promising first step towards the capture and recovery of nitrogen in municipal wastewater, the heterogeneity of process variables in wastewater treatment cannot be understated, and we would like to explicitly address key limitations arising from our experimental design choices.

To simplify the interpretation of experimental results and ensure adequate biomass for purification and analysis, the concentrations of carbon and nitrogen used in our media were supplied at much higher levels than and in ratios not representative of municipal wastewater entering secondary treatment. In wastewater contexts, biological oxygen demand (BOD) is commonly used as a proxy for carbon concentration, describing the oxygen required to biologically catabolize the organic material present. Complete oxidation of one mole of acetate requires two moles of O2, so the range of carbon used in this study (50-100 mM sodium acetate) is equivalent to a theoretical BOD range of 3,200-6,400 mg O2/L. Our use of 50 mM ammonium sulfate (100 mM ammonium ions) translates to 1,400 mg N/L. While a description of “typical” conditions at a “typical” wastewater treatment plant is complicated by the heterogeneity inherent to these processes, the nutrient concentrations in our media clearly exceed reported values from primary effluent/secondary influent by over an order of magnitude: BOD ranges span 60-450 mg O2/L and total nitrogen ranges span 15-100 mg N/L^35, 36^. Because substrate import slows and bacterial metabolism tends to restructure in dilute conditions, strain performance in high-nutrient lab media may not translate to performance in a low-nutrient real wastewater setting.

Our media also differed from real wastewater in its ratio of available carbon to nitrogen. The BOD:N ratio of the media used in this study ranged from approximately 2.3-4.6 g O2/g N, at or below the estimated minimum of 4-6 g O2/g N required for conventional nitrification-denitrification^37^. This is consistent with experimental observations, as cell growth was typically limited by acetate availability. As reduced nitrogen availability relative to carbon could constrain flux into arginine and aspartate biosynthesis and thereby limit cyanophycin production, future experiments should test higher C:N ratios more representative of secondary treatment influent.

Additional considerations include the presence of compounds and microbes in real wastewater that were not represented in our media. While we demonstrated cyanophycin production from acetate and confirmed growth on propionate, the effects of other volatile fatty acids or mixed carbon conditions remain untested. In real wastewater, combinations of multiple carbon and nitrogen sources may have unanticipated effects on cellular regulation and metabolism. Similarly, native microbial consortia could compete with or otherwise interact with engineered cyanophycin-producing strains. In this regard, questions regarding the impact of initial engineered seeding ratios on the relative abundance and longevity of engineered strains in consortia remain relevant, along with considerations of long-term genetic stability and biocontainment. Collectively, these considerations highlight the value of testing engineered strains using real wastewater samples in future research efforts.

## 4. Conclusion

This work serves as an example of use-inspired basic research directed at the advancement of a novel nitrogen recovery process in municipal wastewater treatment. To this end, our efforts culminated in the implementation of two groups of genomic modifications: the deregulation of native arginine biosynthesis in *A. baylyi* and the creation of a fructose auxotroph. Each of these changes addressed a critical gap between contemporary cyanophycin production methods and projected constraints for integration with municipal wastewater treatment.

We first constructed an arginine production strain AP1 (Δ*argR* Δ*argB::argBEc*) to address the economically infeasible constraint of exogenous arginine supplementation, enabling per-cell cyanophycin titers of 29% CDW from acetate and urea alone. The success of this demonstration opens an exciting new realm of inquiry for cyanophycin production efforts, suggesting that integration of cyanophycin synthetase into existing arginine-producing strains may yield similar results with minimal effort. Next, based on our observations that cyanophycin production was broadly enhanced across sub-optimal growth conditions (e.g. phosphate limitation, low growth temperature, antibiotics) we hypothesized these conditions enhanced cyanophycin production by limiting dilution to cell growth and increasing the relative carbon flux to cyanophycin synthesis. To test this theory, we constructed fructose-auxotrophs, effectively implanting a new control lever for cell growth orthogonal to unalterable wastewater process variables like temperature. One such auxotroph (Δ*gap*) was integrated into our cyanophycin producing strain and enabled similar per-cell cyanophycin titers at both 12 °C - our previous best production temperature - and 30 °, the upper bound for wastewater temperatures and the conventional incubation temperature for *A. baylyi*. Constructed with a single knockout, this auxotroph required exceptionally small quantities of fructose to facilitate cell growth (< 5% total carbon; ∼1 mmol fructose per 57 mmol acetate). Furthermore, the location of the knockout in mid-glycolysis positions it as an attractive means for implementing growth-production partitioning in *A. baylyi* for other TCA-derived natural products. While this study’s findings are tempered by nontrivial differences between the media employed here and real-world wastewater, this work represents a meaningful step towards societal nitrogen circularity and the use of municipal wastewater as a new locus for industrial biotechnology.

## CRediT authorship contribution statement

**Kevin Fitzgerald:** Conceptualization, Methodology, Investigation, Data curation, Formal analysis, Writing - original draft, Visualization. **Keith Tyo**: Supervision, Writing - review & editing.

## Supporting information

ADP1 GEM with fructose catabolism

## Acknowledgements

We would like to thank Professor Martin Shmeing (McGill University) for providing the codon optimized sequence for *TmcphA1* employed in this study. We would also like to acknowledge the US National Science Foundation’s support via ECO-CBET Award #2033793 as well as the Graduate Student Research Fellowship.

