## Supplementary Information for "High-Yield Recovery of Reactive Nitrogen as Cyanophycin by Engineering *Acinetobacter baylyi* ADP1 under Wastewater-Relevant Conditions"

Kevin Fitzgerald, Keith Tyo*

Department of Chemical and Biological Engineering, Northwestern University, Evanston, IL, USA

*Corresponding author:

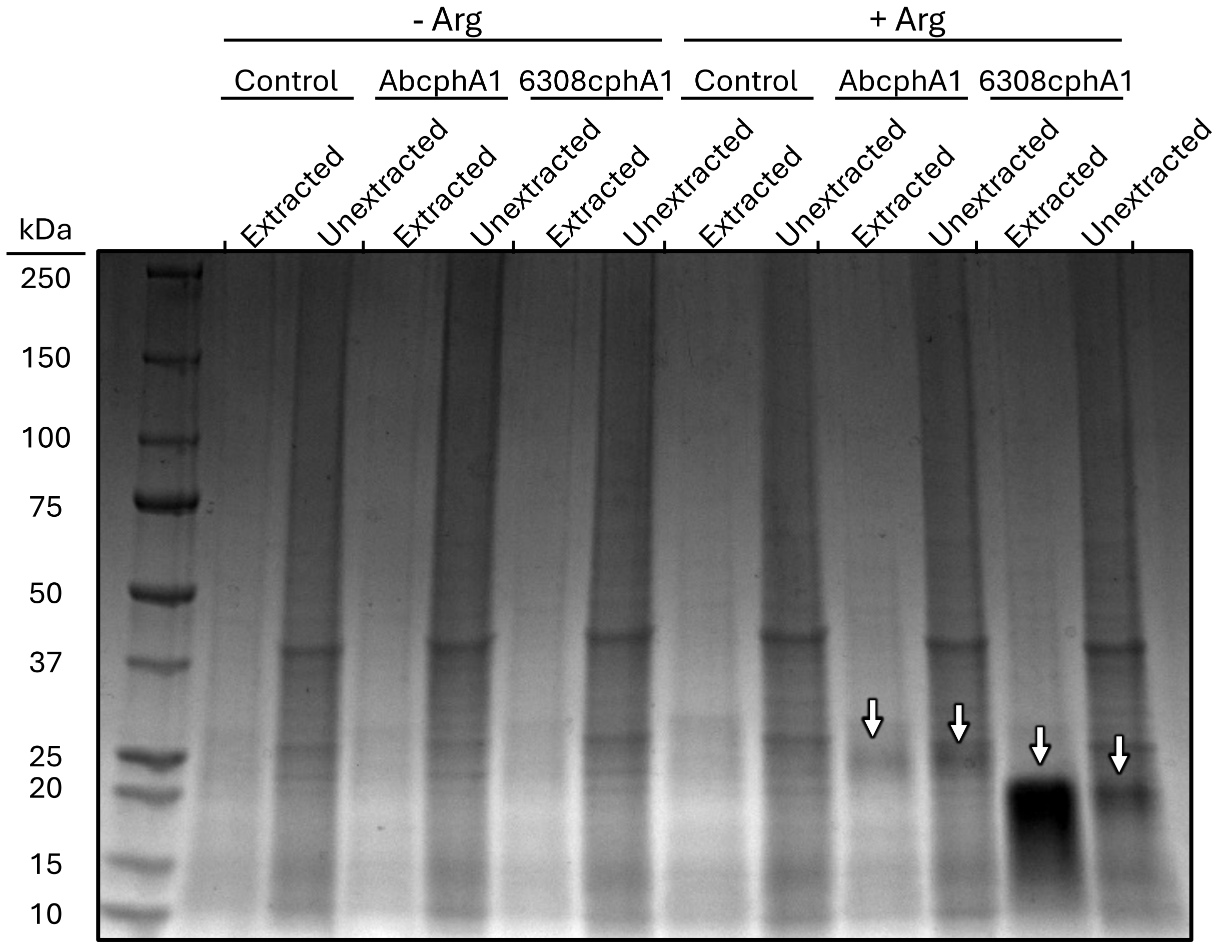

**Figure S1. Efficiency of acid extraction using 0.1 M HCl.** SDS-PAGE gel depicting the effectiveness of acid incubation with removal of intracellular biomass in the experiment outlines in Fig. 2. Following incubation with 0.1 M HCl for at least 1 h, samples were centrifuged and the liquid phase was prepared and loaded as the “extracted” fraction. The remaining solids were resuspended in an equivalent volume of nanopure water, prepared, and loaded as the “unextracted” fraction.

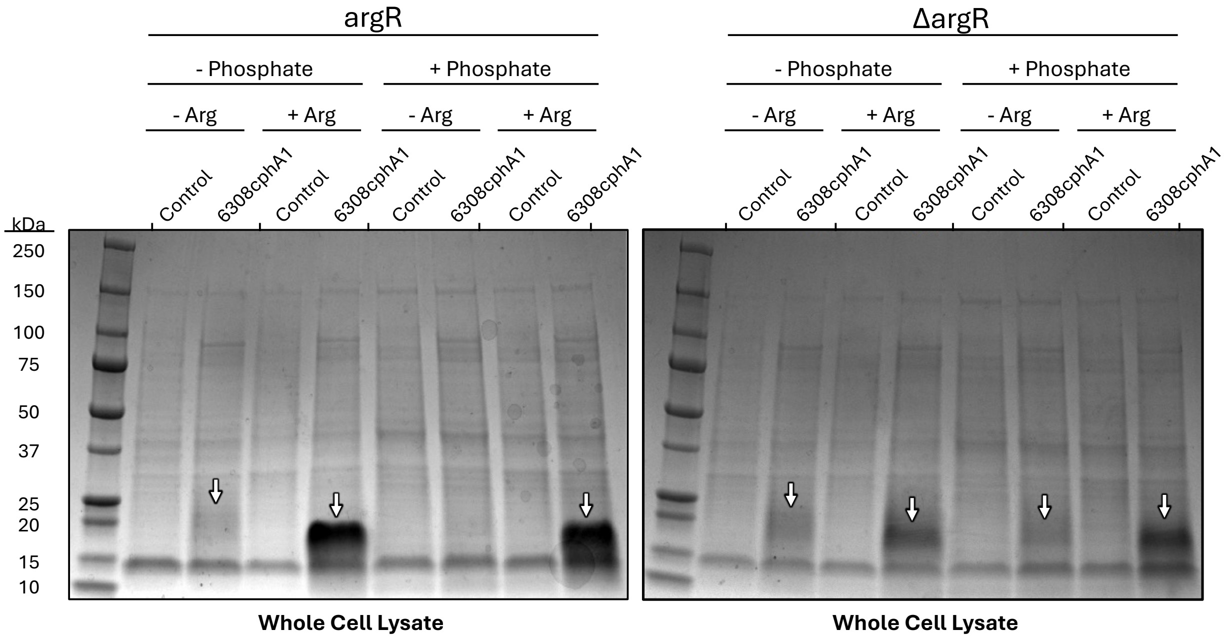

**Figure S2. *argR* enhances cyanophycin accumulation in phosphate replete conditions without arginine supplementation.** SDS-PAGE gel showing the effect of Δ*argR* on cyanophycin accumulation under different medium conditions, specifically phosphate concentration (-Phosphate, 50 μM; +Phosphate, 70 mM) and arginine supplementation (-Arg, 0 mM; +Arg, 60 mM). Cyanophycin bands are indicated by white arrows. Cultures were grown overnight in LB, chilled at 4 °C for 10 min, and precultured in phosphate-limited medium with 1 mM IPTG for 3 h. Cells were then resuspended in the final experimental MSM containing 60 mM glutamate and incubated in 5 mL aliquots in a 24-well plate with 10mL wells.

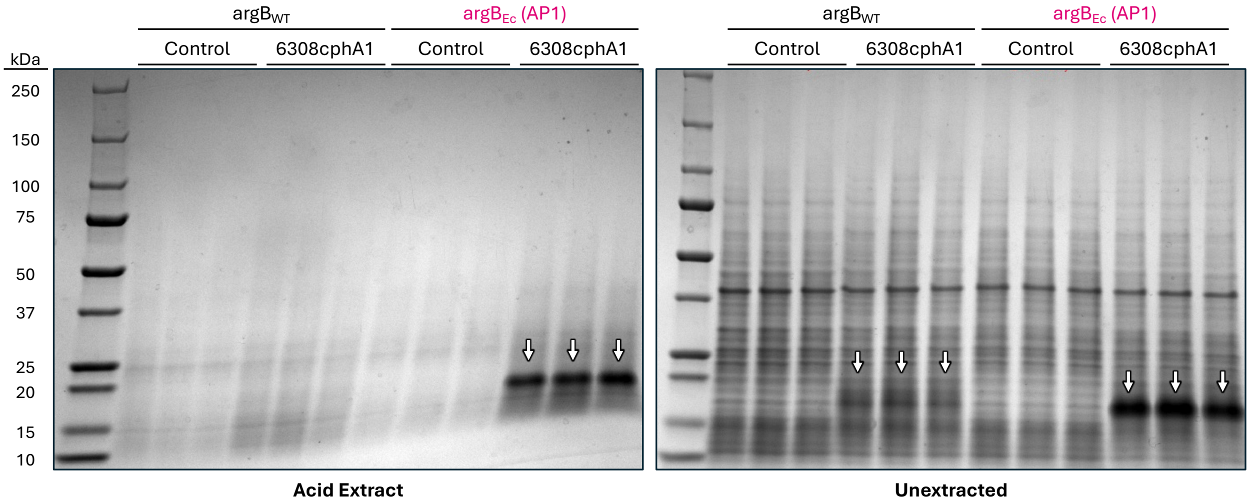

**Figure S3. Acid extraction using 0.1 M HCl insufficient for complete cyanophycin recovery from AP1 samples.** SDS-PAGE gel depicting the effectiveness of acid extraction in recovering cyanophycin from biomass in the experiment described in Fig. 4B (acetate condition). Following incubation with 0.1 M HCl for at least 1 h, samples were centrifuged and the liquid phase was prepared and loaded as the “extracted” fraction. The remaining solids were resuspended in an equivalent volume of nanopure water, prepared, and loaded as the “unextracted” fraction.

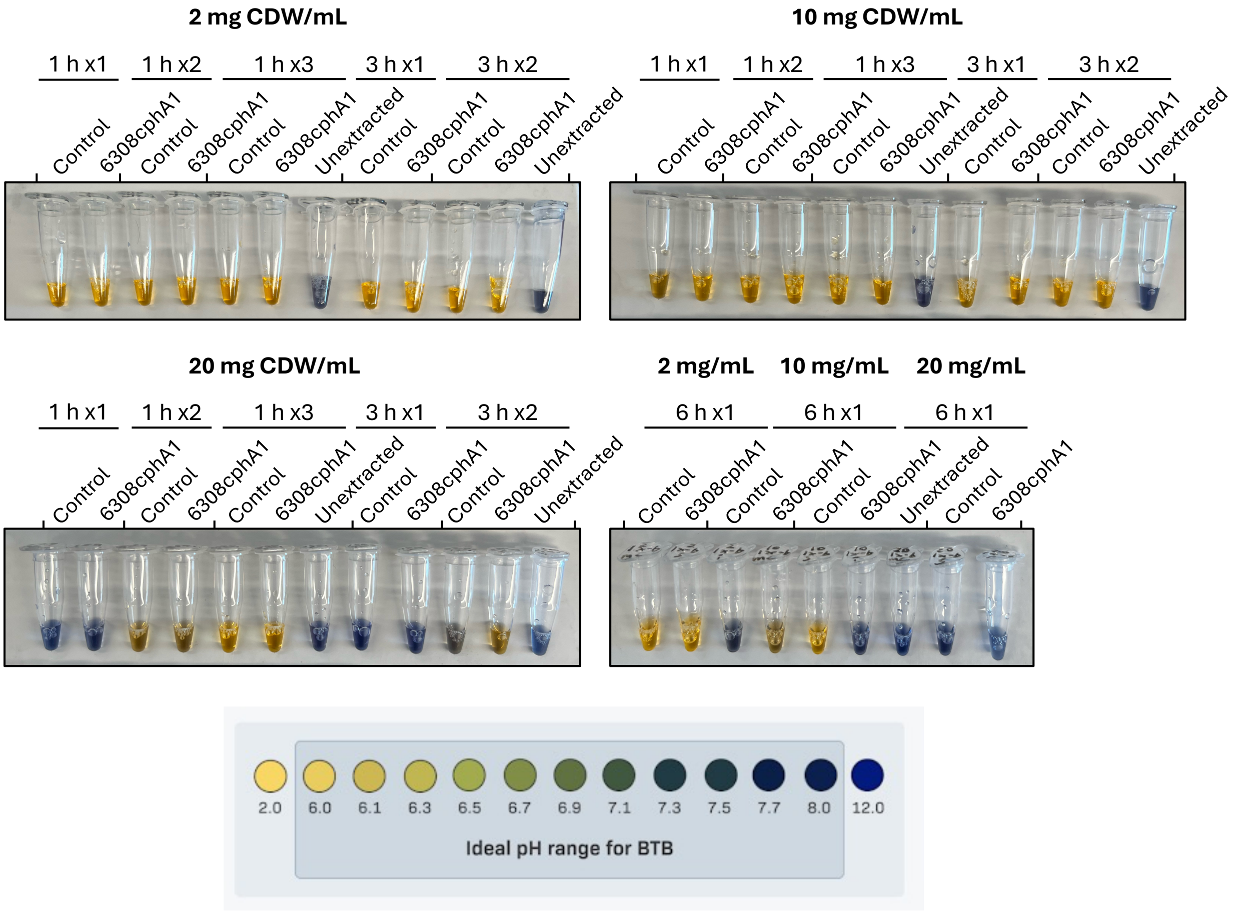

**Figure S4. pH-sensitive bromophenol blue indicates biomass-mediated buffering effect at elevated higher biomass loadings.** Images of SDS-PAGE samples prior to loading in gel depicted in Fig. 5A. Samples at higher biomass concentrations are blue initially and turn yellow after additional acid extractions. Given the identical amount of HCl added to each sample, this is consistent with biomass-mediated pH buffering. The color scale image was adapted from reference[35].

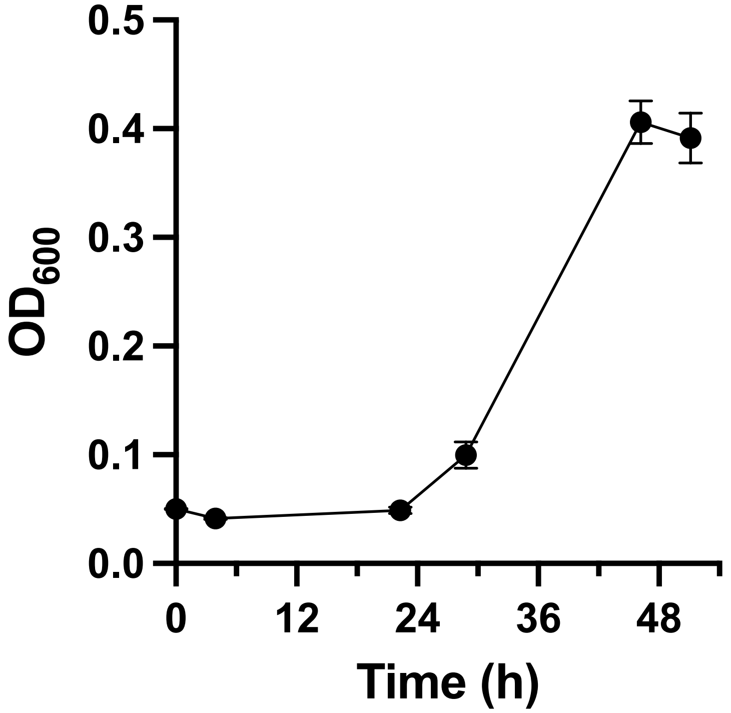

**Figure S5.** ***A. baylyi*** **is capable of utilizing propionate as sole carbon source.**

Growth curve of *A. baylyi* Δ*acr1* (preventing wax ester synthesis) grown on MSM containing 10 mM sodium propionate as the sole carbon source. Strains were precultured in LB medium overnight and inoculated to a starting $OD_{600}$ of 0.05 the following day. MSM medium was identical to that used previously with the exception that 18.8 mM ammonium chloride replaced 50 mM ammonium sulfate as the nitrogen source. Error bars represent SEM for n = 3 biological replicates.

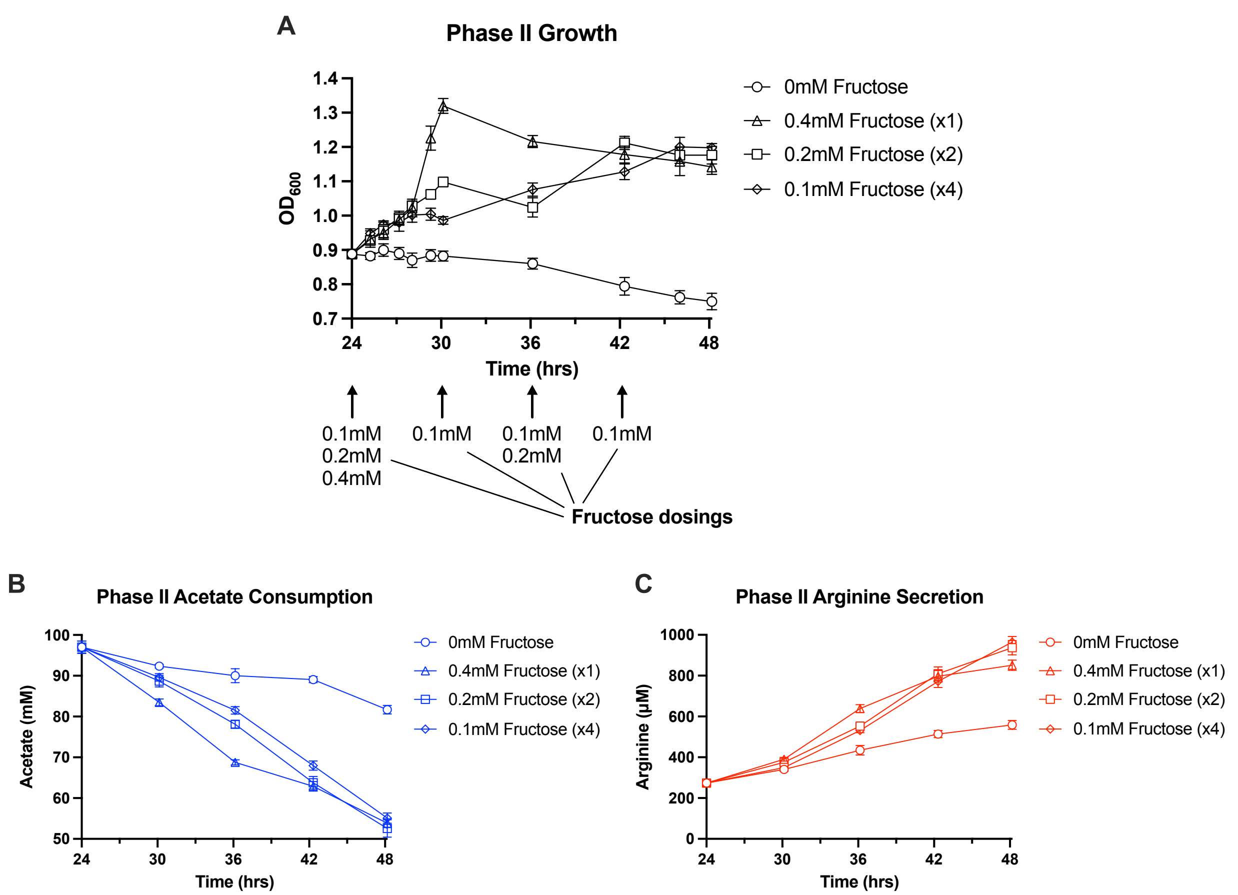

**Figure S6. AP1 Δgap differentially consumes acetate and secretes arginine based on the amount and timing of fructose additions.** (A) $OD_{600}$, fructose dosing, (B) acetate consumption, and (C) arginine secretion data for AP1 Δ*gap* strains during hours 24-48 (Phase II) of the experiment described in Fig. 3.9 A, B. Arginine titers were quantified using the Sakaguchi assay in technical triplicate. Error bars represent SEM for n = 3 biological replicates.

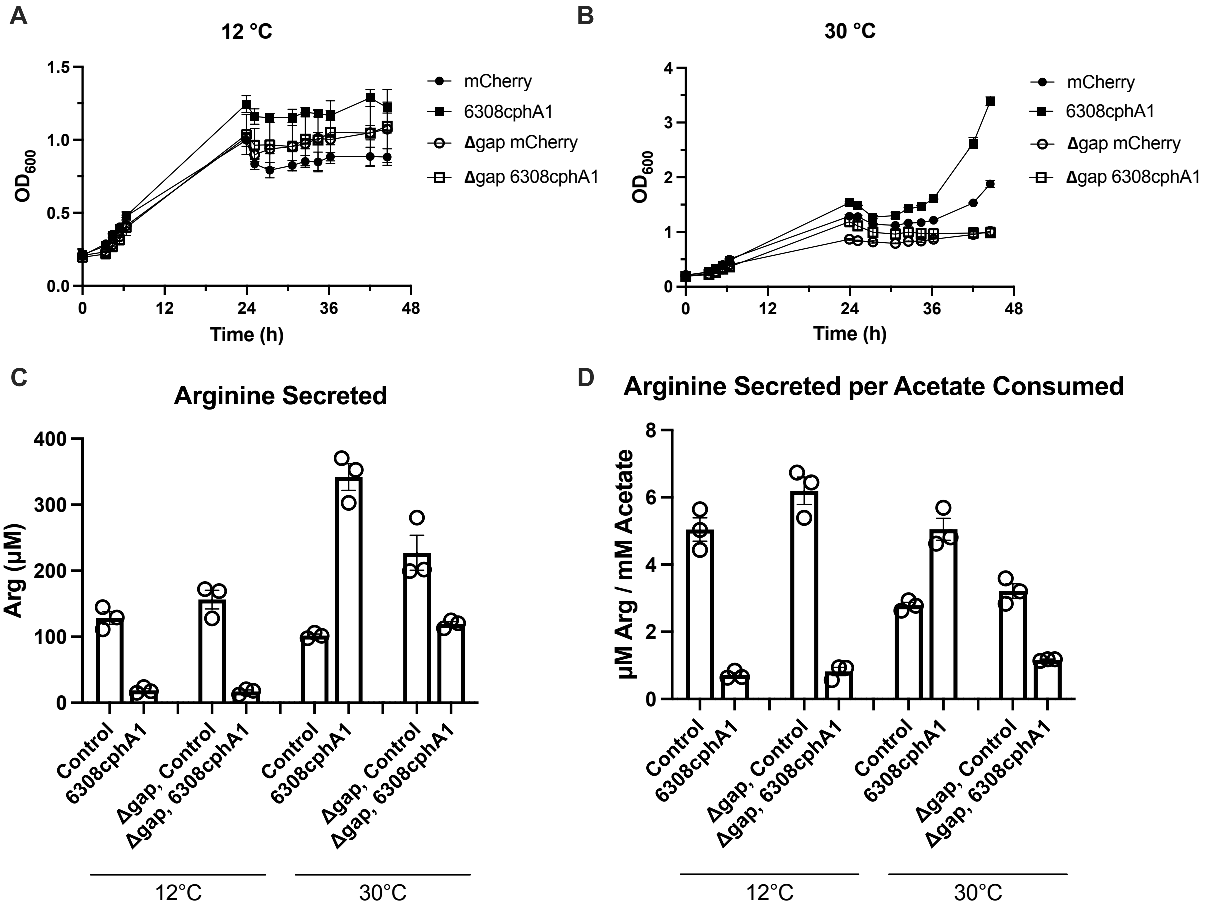

**Figure S7. AP1 and AP1 Δgap display similar arginine-secreting phenotype at 12 °C.** $OD_{600}$ data for cultures grown at **(A)** 12 °C and **(B)** 30 °C in the experiment described in Fig. 9C, D. Supplementation of 100 mM sodium acetate at 24 h appears to temporarily inhibit growth, a response different from that elicited in previous experiments (Fig. S6). Arginine secretion **(C)** and arginine-secretion normalized by acetate-consumption **(D)** data for the experiment described in Fig. 9C, D are also provided. Arginine titers were quantified using the Sakaguchi assay in technical triplicate. Error bars represent SEM for n = 3 biological replicates.

### Table S3. Flux balance analysis of glycolytic gene deletions predicted to confer fructose auxotrophy.

| | | Biomass Flux Across Media Compositions | | | Uptake Flux for Coutilization | | |
|---|---|---|---|---|---|---|---|
| **Genotype** | **Accession #** | **Fructose: 5** | **Acetate: 500** | **Coutilization** | **Fructose** | **Acetate** | **% C from Fructose:** |
| WT | - | 0.46 | 7.62 | 7.85 | 5.00 | 333.05 | - |
| **Δ**fda | ACIAD1925 | 0.00 | 0.00 | 5.42 | 5.00 | 202.92 | 6.88 |
| **Δ**gap | ACIAD2565 | 0.41 | 0.00 | 4.04 | 5.00 | 150.70 | 9.05 |
| **Δ**pgk | ACIAD1927 | 0.46 | 7.62 | 7.85 | 5.00 | 333.05 | - |
| **Δ**gpm1 | ACIAD0256 | 0.04 | 0.00 | 2.17 | 5.00 | 68.82 | 17.90 |
| **Δ**eno | ACIAD2001 | 0.04 | 0.00 | 2.17 | 5.00 | 68.82 | 17.90 |
| **Δ**fda **Δ**gap | - | 0.00 | 0.00 | 0.00 | - | - | - |

Carbon source availability was simulated by setting environmental acetate flux to 500 mmol/h/g CDW and fructose flux to 5 mmol/h/g CDW. The amounts of each consumed were measured and compared to calculate the percent of total carbon derived from fructose. The simulation featuring both Δ*fda* and Δ*gap* mutations revealed both negligible biomass formation and negligible consumption of either substrate. See methods for details of FBA analysis. Flux in units of mmol/h/g CDW.
