## Supplementary material for "High-Yield Recovery of Reactive Nitrogen as Cyanophycin by Engineering *Acinetobacter baylyi* ADP1 under Wastewater-Relevant Conditions": ADP1 GEM with fructose catabolism: ADP1_GEM-L3V1_fbc_cleaned_gpr_annotated_v2.yaml

```
{
"metabolites":[
{
"id":"cpd01727_C",
"name":"Pimelate",
"compartment":"C",
"annotation":{
"smiles":"O=C([O-])CCCCCC(=O)[O-]",
"smiles_sanitized":"O=C(O)CCCCCC(=O)O"
}
},
{
"id":"cpd00001_P",
"name":"H2O",
"compartment":"P",
"annotation":{
"smiles":"O",
"smiles_sanitized":"O"
}
},
{
"id":"cpd02069_C",
"name":"3-Phosphonooxypyruvate",
"compartment":"C",
"annotation":{
"smiles":"O=C([O-])C(=O)COP(=O)([O-])[O-]",
"smiles_sanitized":"O=C(O)C(=O)COP(=O)(O)O"
}
},
{
"id":"cpd02799_C",
"name":"Methylisocitrate",
"compartment":"C",
"annotation":{
"smiles":"C[C@](O)(C(=O)[O-])[C@H](CC(=O)[O-])C(=O)[O-]",
"smiles_sanitized":"CC(O)(C(=O)O)C(CC(=O)O)C(=O)O"
}
},
{
"id":"cpd02597_C",
"name":"cis-2-Methylaconitate",
"compartment":"C",
"annotation":{
"smiles":"C/C(C(=O)[O-])=C(\\CC(=O)[O-])C(=O)[O-]",
"smiles_sanitized":"CC(C(=O)O)=C(CC(=O)O)C(=O)O"
}
},
{
"id":"cpd00153_E",
"name":"Benzoate",
"compartment":"E",
"annotation":{
"smiles":"O=C([O-])c1ccccc1",
"smiles_sanitized":"O=C(O)c1ccccc1"
}
},
{
"id":"cpd00037_C",
"name":"UDP-N-acetylglucosamine",
"compartment":"C",
"annotation":{
"smiles":"CC(=O)N[C@H]1[C@@H](OP(=O)([O-])OP(=O)([O-])OC[C@H]2O[C@@H](n3ccc(=O)[nH]c3=O)[C@H](O)[C@@H]2O)O[C@H](CO)[C@@H](O)[C@@H]1O",
"smiles_sanitized":"CC(=O)NC1C(OP(=O)(O)OP(=O)(O)OCC2OC(n3ccc(=O)[nH]c3=O)C(O)C2O)OC(CO)C(O)C1O"
}
},
{
"id":"cpd00013_C",
"name":"NH3",
"compartment":"C",
"annotation":{
"smiles":"[NH4+]",
"smiles_sanitized":"N"
}
},
{
"id":"cpd00007_E",
"name":"O2",
"compartment":"E",
"annotation":{
"smiles":"O=O",
"smiles_sanitized":"O=O"
}
},
{
"id":"cpd00170_P",
"name":"Gluconolactone",
"compartment":"P",
"annotation":{
"smiles":"O=C1O[C@H](CO)[C@@H](O)[C@H](O)[C@H]1O",
"smiles_sanitized":"O=C1OC(CO)C(O)C(O)C1O"
}
},
{
"id":"cpd07333_E",
"name":"Dihydrocaffeic acid",
"compartment":"E",
"annotation":{
"smiles":"O=C([O-])CCc1ccc(O)c(O)c1",
"smiles_sanitized":"O=C(O)CCc1ccc(O)c(O)c1"
}
},
{
"id":"cpd00033_E",
"name":"Glycine",
"compartment":"E",
"annotation":{
"smiles":"[NH3+]CC(=O)[O-]",
"smiles_sanitized":"NCC(=O)O"
}
},
{
"id":"cpd00029_C",
"name":"Acetate",
"compartment":"C",
"annotation":{
"smiles":"CC(=O)[O-]",
"smiles_sanitized":"CC(=O)O"
}
},
{
"id":"cpd00026_C",
"name":"UDP-glucose",
"compartment":"C",
"annotation":{
"smiles":"O=c1ccn([C@@H]2O[C@H](COP(=O)([O-])OP(=O)([O-])O[C@H]3O[C@H](CO)[C@@H](O)[C@H](O)[C@H]3O)[C@@H](O)[C@H]2O)c(=O)[nH]1",
"smiles_sanitized":"O=c1ccn(C2OC(COP(=O)(O)OP(=O)(O)OC3OC(CO)C(O)C(O)C3O)C(O)C2O)c(=O)[nH]1"
}
},
{
"id":"cpd00135_C",
"name":"Homocysteine",
"compartment":"C",
"annotation":{
"smiles":"[NH3+][C@@H](CCS)C(=O)[O-]",
"smiles_sanitized":"NC(CCS)C(=O)O"
}
},
{
"id":"cpd00971_E",
"name":"Na+",
"compartment":"E",
"annotation":{
"smiles":"[Na+]",
"smiles_sanitized":"[Na+]"
}
},
{
"id":"cpd16605_C",
"name":"9-carboxy-3-hydroxynonanoyl-coa",
"compartment":"C",
"annotation":{
"smiles":"",
"smiles_sanitized":""
}
},
{
"id":"cpd02552_C",
"name":"n-acetylglutamyl-phosphate",
"compartment":"C",
"annotation":{
"smiles":"CC(=O)N[C@@H](CCC(=O)OP(=O)([O-])[O-])C(=O)[O-]",
"smiles_sanitized":"CC(=O)NC(CCC(=O)OP(=O)(O)O)C(=O)O"
}
},
{
"id":"cpd01311_C",
"name":"Dethiobiotin",
"compartment":"C",
"annotation":{
"smiles":"CC1NC(=O)NC1CCCCCC(=O)[O-]",
"smiles_sanitized":"CC1NC(=O)NC1CCCCCC(=O)O"
}
},
{
"id":"cpd00019_C",
"name":"S-Adenosyl-homocysteine",
"compartment":"C",
"annotation":{
"smiles":"Nc1ncnc2c1ncn2[C@@H]1O[C@H](CSCC[C@H]([NH3+])C(=O)[O-])[C@@H](O)[C@H]1O",
"smiles_sanitized":"Nc1ncnc2c1ncn2C1OC(CSCCC(N)C(=O)O)C(O)C1O"
}
},
{
"id":"cpd00043_C",
"name":"UDP-galactose",
"compartment":"C",
"annotation":{
"smiles":"O=c1ccn([C@@H]2O[C@H](COP(=O)([O-])OP(=O)([O-])OC3O[C@H](CO)[C@H](O)[C@H](O)[C@H]3O)[C@@H](O)[C@H]2O)c(=O)[nH]1",
"smiles_sanitized":"O=c1ccn(C2OC(COP(=O)(O)OP(=O)(O)OC3OC(CO)C(O)C(O)C3O)C(O)C2O)c(=O)[nH]1"
}
},
{
"id":"cpd16607_C",
"name":"9-carboxynonanoyl-coa",
"compartment":"C",
"annotation":{
"smiles":"",
"smiles_sanitized":""
}
},
{
"id":"cpd00809_C",
"name":"O-Phospho-L-homoserine",
"compartment":"C",
"annotation":{
"smiles":"[NH3+][C@@H](CCOP(=O)([O-])[O-])C(=O)[O-]",
"smiles_sanitized":"NC(CCOP(=O)(O)O)C(=O)O"
}
},
{
"id":"cpd00156_C",
"name":"L-Valine",
"compartment":"C",
"annotation":{
"smiles":"CC(C)[C@H]([NH3+])C(=O)[O-]",
"smiles_sanitized":"CC(C)C(N)C(=O)O"
}
},
{
"id":"cpd00412_E",
"name":"Deoxyuridine",
"compartment":"E",
"annotation":{
"smiles":"O=c1ccn([C@H]2C[C@H](O)[C@@H](CO)O2)c(=O)[nH]1",
"smiles_sanitized":"O=c1ccn(C2CC(O)C(CO)O2)c(=O)[nH]1"
}
},
{
"id":"cpd00599_C",
"name":"SALC",
"compartment":"C",
"annotation":{
"smiles":"O=C([O-])c1ccccc1O",
"smiles_sanitized":"O=C(O)c1ccccc1O"
}
},
{
"id":"cpd02775_C",
"name":"4-Amino-5-phosphomethyl-2-methylpyrimidine",
"compartment":"C",
"annotation":{
"smiles":"Cc1nc(N)c(COP(=O)([O-])[O-])c[nH+]1",
"smiles_sanitized":"Cc1ncc(COP(=O)(O)O)c(N)n1"
}
},
{
"id":"cpd01471_C",
"name":"Arylsulfate",
"compartment":"C",
"annotation":{
"smiles":"O=S(=O)([O-])Oc1ccccc1",
"smiles_sanitized":"O=S(=O)(O)Oc1ccccc1"
}
},
{
"id":"cpd00153_P",
"name":"Benzoate",
"compartment":"P",
"annotation":{
"smiles":"O=C([O-])c1ccccc1",
"smiles_sanitized":"O=C(O)c1ccccc1"
}
},
{
"id":"cpd00209_C",
"name":"Nitrate",
"compartment":"C",
"annotation":{
"smiles":"O=[N+]([O-])[O-]",
"smiles_sanitized":"O=[N+]([O-])[O-]"
}
},
{
"id":"cpd00248_E",
"name":"Quinate",
"compartment":"E",
"annotation":{
"smiles":"O=C([O-])[C@]1(O)C[C@@H](O)[C@@H](O)[C@H](O)C1",
"smiles_sanitized":"O=C(O)C1(O)CC(O)C(O)C(O)C1"
}
},
{
"id":"cpd00201_C",
"name":"10-Formyltetrahydrofolate",
"compartment":"C",
"annotation":{
"smiles":"Nc1nc2c(c(=O)[nH]1)N[C@@H](CN(C=O)c1ccc(C(=O)N[C@@H](CCC(=O)[O-])C(=O)[O-])cc1)CN2",
"smiles_sanitized":"Nc1nc2c(c(=O)[nH]1)NC(CN(C=O)c1ccc(C(=O)NC(CCC(=O)O)C(=O)O)cc1)CN2"
}
},
{
"id":"cpd00270_C",
"name":"Caffeoyl-CoA",
"compartment":"C",
"annotation":{
"smiles":"CC(C)(COP(=O)([O-])OP(=O)([O-])OC[C@H]1O[C@@H](n2cnc3c(N)ncnc32)[C@H](O)[C@@H]1OP(=O)([O-])[O-])[C@@H](O)C(=O)NCCC(=O)NCCSC(=O)/C=C/c1ccc(O)c(O)c1",
"smiles_sanitized":"CC(C)(COP(=O)(O)OP(=O)(O)OCC1OC(n2cnc3c(N)ncnc32)C(O)C1OP(=O)(O)O)C(O)C(=O)NCCC(=O)NCCSC(=O)C=Cc1ccc(O)c(O)c1"
}
},
{
"id":"cpd02546_C",
"name":"CMP-KDO",
"compartment":"C",
"annotation":{
"smiles":"Nc1ccn([C@@H]2O[C@H](COP(=O)([O-])O[C@]3(C(=O)[O-])C[C@@H](O)[C@@H](O)[C@@H]([C@H](O)CO)O3)[C@@H](O)[C@H]2O)c(=O)n1",
"smiles_sanitized":"Nc1ccn(C2OC(COP(=O)(O)OC3(C(=O)O)CC(O)C(O)C(C(O)CO)O3)C(O)C2O)c(=O)n1"
}
},
{
"id":"cpd00001_C",
"name":"H2O",
"compartment":"C",
"annotation":{
"smiles":"O",
"smiles_sanitized":"O"
}
},
{
"id":"cpd00506_C",
"name":"gamma-Glutamylcysteine",
"compartment":"C",
"annotation":{
"smiles":"[NH3+][C@@H](CCC(=O)N[C@@H](CS)C(=O)[O-])C(=O)[O-]",
"smiles_sanitized":"NC(CCC(=O)NC(CS)C(=O)O)C(=O)O"
}
},
{
"id":"cpd00103_C",
"name":"PRPP",
"compartment":"C",
"annotation":{
"smiles":"O=P([O-])([O-])OC[C@H]1O[C@H](OP(=O)([O-])OP(=O)([O-])O)[C@H](O)[C@@H]1O",
"smiles_sanitized":"O=P(O)(O)OCC1OC(OP(=O)(O)OP(=O)(O)O)C(O)C1O"
}
},
{
"id":"cpd03128_C",
"name":"Decanoyl-CoA",
"compartment":"C",
"annotation":{
"smiles":"CCCCCCCCCC(=O)SCCNC(=O)CCNC(=O)[C@H](O)C(C)(C)COP(=O)([O-])OP(=O)([O-])OC[C@H]1O[C@@H](n2cnc3c(N)ncnc32)[C@H](O)[C@@H]1OP(=O)([O-])[O-]",
"smiles_sanitized":"CCCCCCCCCC(=O)SCCNC(=O)CCNC(=O)C(O)C(C)(C)COP(=O)(O)OP(=O)(O)OCC1OC(n2cnc3c(N)ncnc32)C(O)C1OP(=O)(O)O"
}
},
{
"id":"cpd00383_E",
"name":"Shikimate",
"compartment":"E",
"annotation":{
"smiles":"O=C([O-])C1=C[C@@H](O)[C@@H](O)[C@H](O)C1",
"smiles_sanitized":"O=C(O)C1=CC(O)C(O)C(O)C1"
}
},
{
"id":"ACYL_ACP_TAG_C",
"name":"ACYL-ACP_TAG",
"compartment":"C"
},
{
"id":"cpd00515_C",
"name":"5-Dehydro-4-deoxy-D-glucarate",
"compartment":"C",
"annotation":{
"smiles":"O=C([O-])C(=O)C[C@H](O)[C@@H](O)C(=O)[O-]",
"smiles_sanitized":"O=C(O)C(=O)CC(O)C(O)C(=O)O"
}
},
{
"id":"cpd00051_E",
"name":"L-Arginine",
"compartment":"E",
"annotation":{
"smiles":"NC(=[NH2+])NCCC[C@H]([NH3+])C(=O)[O-]",
"smiles_sanitized":"N=C(N)NCCCC(N)C(=O)O"
}
},
{
"id":"cpd00227_E",
"name":"L-Homoserine",
"compartment":"E",
"annotation":{
"smiles":"[NH3+][C@@H](CCO)C(=O)[O-]",
"smiles_sanitized":"NC(CCO)C(=O)O"
}
},
{
"id":"cpd02843_C",
"name":"D-erythro-imidazol-glycerol-phosphate",
"compartment":"C",
"annotation":{
"smiles":"O=P([O-])([O-])OC[C@@H](O)[C@@H](O)c1c[nH]cn1",
"smiles_sanitized":"O=P(O)(O)OCC(O)C(O)c1c[nH]cn1"
}
},
{
"id":"L_1_PHOSPHATIDYL_GLYCEROL_P_C",
"name":"generic l-1-phosphatidyl-glycerol-phosphate",
"compartment":"C"
},
{
"id":"cpd00048_E",
"name":"Sulfate",
"compartment":"E",
"annotation":{
"smiles":"O=S(=O)([O-])[O-]",
"smiles_sanitized":"O=S(=O)(O)O"
}
},
{
"id":"cpd00241_C",
"name":"dGTP",
"compartment":"C",
"annotation":{
"smiles":"Nc1nc2c(ncn2[C@H]2C[C@H](O)[C@@H](COP(=O)([O-])OP(=O)([O-])OP(=O)([O-])O)O2)c(=O)[nH]1",
"smiles_sanitized":"Nc1nc2c(ncn2C2CC(O)C(COP(=O)(O)OP(=O)(O)OP(=O)(O)O)O2)c(=O)[nH]1"
}
},
{
"id":"C6_0_3HYDROXYACP_C",
"name":"3-hydroxyhexanoyl-acp",
"compartment":"C"
},
{
"id":"cpd02886_C",
"name":"UDP-3-O-(beta-hydroxymyristoyl)-N-acetylglucosamine",
"compartment":"C",
"annotation":{
"smiles":"CCCCCCCCCCC[C@@H](O)CC(=O)O[C@H]1[C@H](O)[C@@H](CO)O[C@H](OP(=O)([O-])OP(=O)([O-])OC[C@H]2O[C@@H](n3ccc(=O)[nH]c3=O)[C@H](O)[C@@H]2O)[C@@H]1NC(C)=O",
"smiles_sanitized":"CCCCCCCCCCCC(O)CC(=O)OC1C(O)C(CO)OC(OP(=O)(O)OP(=O)(O)OCC2OC(n3ccc(=O)[nH]c3=O)C(O)C2O)C1NC(C)=O"
}
},
{
"id":"cpd00146_C",
"name":"Carbamoylphosphate",
"compartment":"C",
"annotation":{
"smiles":"NC(=O)OP(=O)([O-])[O-]",
"smiles_sanitized":"NC(=O)OP(=O)(O)O"
}
},
{
"id":"cpd00810_C",
"name":"Orotidylic acid",
"compartment":"C",
"annotation":{
"smiles":"O=C([O-])c1cc(=O)[nH]c(=O)n1[C@@H]1O[C@H](COP(=O)([O-])[O-])[C@@H](O)[C@H]1O",
"smiles_sanitized":"O=C(O)c1cc(=O)[nH]c(=O)n1C1OC(COP(=O)(O)O)C(O)C1O"
}
},
{
"id":"cpd00104_C",
"name":"BIOT",
"compartment":"C",
"annotation":{
"smiles":"O=C([O-])CCCC[C@@H]1SC[C@@H]2NC(=O)N[C@H]12",
"smiles_sanitized":"O=C(O)CCCCC1SCC2NC(=O)NC21"
}
},
{
"id":"cpd00221_C",
"name":"D-Lactate",
"compartment":"C",
"annotation":{
"smiles":"C[C@@H](O)C(=O)[O-]",
"smiles_sanitized":"CC(O)C(=O)O"
}
},
{
"id":"3_OHLAUROYL_ACP_C",
"name":"3-hydroxydodecanoyl-acp",
"compartment":"C"
},
{
"id":"cpd00363_E",
"name":"Ethanol",
"compartment":"E",
"annotation":{
"smiles":"CCO",
"smiles_sanitized":"CCO"
}
},
{
"id":"cpd00226_C",
"name":"HYXN",
"compartment":"C",
"annotation":{
"smiles":"O=c1[nH]cnc2[nH]cnc12",
"smiles_sanitized":"O=c1[nH]cnc2[nH]cnc12"
}
},
{
"id":"cpd00550_C",
"name":"D-Serine",
"compartment":"C",
"annotation":{
"smiles":"[NH3+][C@H](CO)C(=O)[O-]",
"smiles_sanitized":"NC(CO)C(=O)O"
}
},
{
"id":"cpd00142_E",
"name":"Acetoacetate",
"compartment":"E",
"annotation":{
"smiles":"CC(=O)CC(=O)[O-]",
"smiles_sanitized":"CC(=O)CC(=O)O"
}
},
{
"id":"cpd00357_C",
"name":"TTP",
"compartment":"C",
"annotation":{
"smiles":"Cc1cn([C@H]2C[C@H](O)[C@@H](COP(=O)([O-])OP(=O)([O-])OP(=O)([O-])O)O2)c(=O)[nH]c1=O",
"smiles_sanitized":"Cc1cn(C2CC(O)C(COP(=O)(O)OP(=O)(O)OP(=O)(O)O)O2)c(=O)[nH]c1=O"
}
},
{
"id":"cpd00236_C",
"name":"D-Erythrose4-phosphate",
"compartment":"C",
"annotation":{
"smiles":"O=C[C@H](O)[C@H](O)COP(=O)([O-])[O-]",
"smiles_sanitized":"O=CC(O)C(O)COP(=O)(O)O"
}
},
{
"id":"cpd02701_C",
"name":"S-Adenosyl-4-methylthio-2-oxobutanoate",
"compartment":"C",
"annotation":{
"smiles":"C[S+](CCC(=O)C(=O)[O-])C[C@H]1O[C@@H](n2cnc3c(N)ncnc32)[C@H](O)[C@@H]1O",
"smiles_sanitized":"C[S+](CCC(=O)C(=O)O)CC1OC(n2cnc3c(N)ncnc32)C(O)C1O"
}
},
{
"id":"cpd31963_C",
"name":"(2E,9Z)-octadecenoyl-CoA",
"compartment":"C",
"annotation":{
"smiles":"CCCCCCCC/C=C\\CCCCC/C=C/C(=O)SCCNC(=O)CCNC(=O)[C@H](O)C(C)(C)COP(=O)([O-])OP(=O)([O-])OC[C@H]1O[C@@H](n2cnc3c(N)ncnc32)[C@H](O)[C@@H]1OP(=O)([O-])[O-]",
"smiles_sanitized":"CCCCCCCCC=CCCCCCC=CC(=O)SCCNC(=O)CCNC(=O)C(O)C(C)(C)COP(=O)(O)OP(=O)(O)OCC1OC(n2cnc3c(N)ncnc32)C(O)C1OP(=O)(O)O"
}
},
{
"id":"cpd00220_C",
"name":"Riboflavin",
"compartment":"C",
"annotation":{
"smiles":"Cc1cc2nc3c(=O)[n-]c(=O)nc-3n(C[C@H](O)[C@H](O)[C@H](O)CO)c2cc1C",
"smiles_sanitized":"Cc1cc2nc3c(=O)[nH]c(=O)nc-3n(CC(O)C(O)C(O)CO)c2cc1C"
}
},
{
"id":"cpd00764_C",
"name":"7-8-Diaminononanoate",
"compartment":"C",
"annotation":{
"smiles":"CC([NH3+])C([NH3+])CCCCCC(=O)[O-]",
"smiles_sanitized":"CC(N)C(N)CCCCCC(=O)O"
}
},
{
"id":"cpd03526_C",
"name":"N-Succinyl-L-glutamate 5-semialdehyde",
"compartment":"C",
"annotation":{
"smiles":"O=CCC[C@H](NC(=O)CCC(=O)[O-])C(=O)[O-]",
"smiles_sanitized":"O=CCCC(NC(=O)CCC(=O)O)C(=O)O"
}
},
{
"id":"cpd00521_C",
"name":"dTDP-4-oxo-6-deoxy-D-glucose",
"compartment":"C",
"annotation":{
"smiles":"Cc1cn([C@H]2C[C@H](O)[C@@H](COP(=O)([O-])OP(=O)([O-])O[C@H]3O[C@H](C)C(=O)[C@H](O)[C@H]3O)O2)c(=O)[nH]c1=O",
"smiles_sanitized":"Cc1cn(C2CC(O)C(COP(=O)(O)OP(=O)(O)OC3OC(C)C(=O)C(O)C3O)O2)c(=O)[nH]c1=O"
}
},
{
"id":"cpd00638_C",
"name":"Deamido-NAD",
"compartment":"C",
"annotation":{
"smiles":"Nc1ncnc2c1ncn2[C@@H]1O[C@H](COP(=O)([O-])OP(=O)([O-])OC[C@H]2O[C@@H]([n+]3cccc(C(=O)[O-])c3)[C@H](O)[C@@H]2O)[C@@H](O)[C@H]1O",
"smiles_sanitized":"Nc1ncnc2c1ncn2C1OC(COP(=O)(O)OP(=O)(O)OCC2OC([n+]3cccc(C(=O)O)c3)C(O)C2O)C(O)C1O"
}
},
{
"id":"cpd00652_C",
"name":"D-Mucic acid",
"compartment":"C",
"annotation":{
"smiles":"O=C([O-])[C@@H](O)[C@H](O)[C@H](O)[C@@H](O)C(=O)[O-]",
"smiles_sanitized":"O=C(O)C(O)C(O)C(O)C(O)C(=O)O"
}
},
{
"id":"cpd00097_P",
"name":"Pyrroloquinoline-quinone",
"compartment":"P",
"annotation":{
"smiles":"O=C([O-])c1cc(C(=O)[O-])c2c(n1)C(=O)C(=O)c1cc(C(=O)[O-])[nH]c1-2",
"smiles_sanitized":"O=C(O)c1cc(C(=O)O)c2c(n1)C(=O)C(=O)c1cc(C(=O)O)[nH]c1-2"
}
},
{
"id":"cpd08289_C",
"name":"1-deoxy-D-xylulose5-phosphate",
"compartment":"C",
"annotation":{
"smiles":"CC(=O)[C@@H](O)[C@H](O)COP(=O)([O-])[O-]",
"smiles_sanitized":"CC(=O)C(O)C(O)COP(=O)(O)O"
}
},
{
"id":"cpd00184_E",
"name":"Thymidine",
"compartment":"E",
"annotation":{
"smiles":"Cc1cn([C@H]2C[C@H](O)[C@@H](CO)O2)c(=O)[nH]c1=O",
"smiles_sanitized":"Cc1cn(C2CC(O)C(CO)O2)c(=O)[nH]c1=O"
}
},
{
"id":"cpd01034_E",
"name":"Toluol",
"compartment":"E",
"annotation":{
"smiles":"Cc1ccccc1",
"smiles_sanitized":"Cc1ccccc1"
}
},
{
"id":"cpd00145_C",
"name":"Hydroxypyruvate",
"compartment":"C",
"annotation":{
"smiles":"O=C([O-])C(=O)CO",
"smiles_sanitized":"O=C(O)C(=O)CO"
}
},
{
"id":"C14_1_C5__3HYDROXYACP_C",
"name":"cis-3-hydroxytetradec-4-enoyl-acp",
"compartment":"C"
},
{
"id":"cpd00742_C",
"name":"Allophanate",
"compartment":"C",
"annotation":{
"smiles":"NC(=O)NC(=O)[O-]",
"smiles_sanitized":"NC(=O)NC(=O)O"
}
},
{
"id":"cpd11420_C",
"name":"trdox",
"compartment":"C",
"annotation":{
"smiles":"*[NH2+][C@H]1CSSC[C@@H](C(*)=O)NC(=O)[C@H](*)NC(=O)[C@H](*)NC1=O",
"smiles_sanitized":"*NC1CSSCC(C(*)=O)NC(=O)C(*)NC(=O)C(*)NC1=O"
}
},
{
"id":"cpd00178_C",
"name":"Acetone",
"compartment":"C",
"annotation":{
"smiles":"CC(C)=O",
"smiles_sanitized":"CC(C)=O"
}
},
{
"id":"cpd15378_C",
"name":"4-Hydroxy-benzylalcohol",
"compartment":"C",
"annotation":{
"smiles":"OCc1ccc(O)cc1",
"smiles_sanitized":"OCc1ccc(O)cc1"
}
},
{
"id":"cpd00098_E",
"name":"Choline",
"compartment":"E",
"annotation":{
"smiles":"C[N+](C)(C)CCO",
"smiles_sanitized":"C[N+](C)(C)CCO"
}
},
{
"id":"cpd02851_C",
"name":"AICAR",
"compartment":"C",
"annotation":{
"smiles":"NC(=O)c1ncn([C@@H]2O[C@H](COP(=O)([O-])[O-])[C@@H](O)[C@H]2O)c1N",
"smiles_sanitized":"NC(=O)c1ncn(C2OC(COP(=O)(O)O)C(O)C2O)c1N"
}
},
{
"id":"cpd03709_C",
"name":"Salicylaldehyde",
"compartment":"C",
"annotation":{
"smiles":"O=Cc1ccccc1O",
"smiles_sanitized":"O=Cc1ccccc1O"
}
},
{
"id":"cpd00938_C",
"name":"4-Carboxymuconolactone",
"compartment":"C",
"annotation":{
"smiles":"O=C([O-])C[C@]1(C(=O)[O-])C=CC(=O)O1",
"smiles_sanitized":"O=C(O)CC1(C(=O)O)C=CC(=O)O1"
}
},
{
"id":"C8_0_ACP_C",
"name":"octanoyl-acp",
"compartment":"C"
},
{
"id":"cpd16654_C",
"name":"tricarballylate",
"compartment":"C",
"annotation":{
"smiles":"O=C([O-])CC(CC(=O)[O-])C(=O)[O-]",
"smiles_sanitized":"O=C(O)CC(CC(=O)O)C(=O)O"
}
},
{
"id":"cpd07333_C",
"name":"Dihydrocaffeic acid",
"compartment":"C",
"annotation":{
"smiles":"O=C([O-])CCc1ccc(O)c(O)c1",
"smiles_sanitized":"O=C(O)CCc1ccc(O)c(O)c1"
}
},
{
"id":"cpd01620_C",
"name":"Precorrin 2",
"compartment":"C",
"annotation":{
"smiles":"C[C@@]1(CC(=O)[O-])/C2=C/c3[nH]c(c(CCC(=O)[O-])c3CC(=O)[O-])Cc3[nH]c(c(CC(=O)[O-])c3CCC(=O)[O-])/C=C3\\N/C(=C\\C(=[NH+]2)[C@H]1CCC(=O)[O-])[C@@](C)(CC(=O)[O-])[C@@H]3CCC(=O)[O-]",
"smiles_sanitized":"CC1(CC(=O)O)C2=Cc3[nH]c(c(CCC(=O)O)c3CC(=O)O)Cc3[nH]c(c(CC(=O)O)c3CCC(=O)O)C=C3NC(=CC(=N2)C1CCC(=O)O)C(C)(CC(=O)O)C3CCC(=O)O"
}
},
{
"id":"cpd16666_C",
"name":"propane aldoxime",
"compartment":"C",
"annotation":{
"smiles":"",
"smiles_sanitized":""
}
},
{
"id":"cpd00290_C",
"name":"D-fructose-1,6-bisphosphate",
"compartment":"C",
"annotation":{
"smiles":"O=P([O-])([O-])OC[C@H]1OC(O)(COP(=O)([O-])[O-])[C@@H](O)[C@@H]1O",
"smiles_sanitized":"O=P(O)(O)OCC1OC(O)(COP(=O)(O)O)C(O)C1O"
}
},
{
"id":"cpd00533_C",
"name":"dCDP",
"compartment":"C",
"annotation":{
"smiles":"Nc1ccn([C@H]2C[C@H](O)[C@@H](COP(=O)([O-])OP(=O)([O-])O)O2)c(=O)n1",
"smiles_sanitized":"Nc1ccn(C2CC(O)C(COP(=O)(O)OP(=O)(O)O)O2)c(=O)n1"
}
},
{
"id":"cpd00924_C",
"name":"AppppA",
"compartment":"C",
"annotation":{
"smiles":"Nc1ncnc2c1ncn2[C@@H]1O[C@H](COP(=O)([O-])OP(=O)([O-])OP(=O)([O-])OP(=O)([O-])OC[C@H]2O[C@@H](n3cnc4c(N)ncnc43)[C@H](O)[C@@H]2O)[C@@H](O)[C@H]1O",
"smiles_sanitized":"Nc1ncnc2c1ncn2C1OC(COP(=O)(O)OP(=O)(O)OP(=O)(O)OP(=O)(O)OCC2OC(n3cnc4c(N)ncnc43)C(O)C2O)C(O)C1O"
}
},
{
"id":"cpd02465_C",
"name":"tetrahydrodipicolinate",
"compartment":"C",
"annotation":{
"smiles":"O=C([O-])C1=N[C@H](C(=O)[O-])CCC1",
"smiles_sanitized":"O=C(O)C1=NC(C(=O)O)CCC1"
}
},
{
"id":"cpd00561_C",
"name":"Vanillin",
"compartment":"C",
"annotation":{
"smiles":"COc1cc(C=O)ccc1O",
"smiles_sanitized":"COc1cc(C=O)ccc1O"
}
},
{
"id":"ACYL_OH_WAX_C",
"name":"ACYL-OH_WAX",
"compartment":"C"
},
{
"id":"cpd00221_E",
"name":"D-Lactate",
"compartment":"E",
"annotation":{
"smiles":"C[C@@H](O)C(=O)[O-]",
"smiles_sanitized":"CC(O)C(=O)O"
}
},
{
"id":"cpd15352_C",
"name":"2-Demethylmenaquinone 8",
"compartment":"C",
"annotation":{
"smiles":"CC(C)=CCCC(\\C)=C/CCC(\\C)=C/CCC(\\C)=C/CCC(\\C)=C/CCC(\\C)=C/CCC(\\C)=C/CCC(\\C)=C/CC1(/C=C(C2(/C=C\\C=C/C(/C(/O)=1)=2))/O)",
"smiles_sanitized":"CC(C)=CCCC(C)=CCCC(C)=CCCC(C)=CCCC(C)=CCCC(C)=CCCC(C)=CCCC(C)=CCC1=CC(=O)c2ccccc2C1=O"
}
},
{
"id":"TRP_TRNAS_C",
"name":"tryptophan trna",
"compartment":"C"
},
{
"id":"CHARGED_HIS_TRNAS_C",
"name":"charged histidine trna",
"compartment":"C"
},
{
"id":"cpd00222_E",
"name":"GLCN",
"compartment":"E",
"annotation":{
"smiles":"O=C([O-])[C@H](O)[C@@H](O)[C@H](O)[C@H](O)CO",
"smiles_sanitized":"O=C(O)C(O)C(O)C(O)C(O)CO"
}
},
{
"id":"PRO_TRNAS_C",
"name":"proline trna",
"compartment":"C"
},
{
"id":"cpd00052_C",
"name":"CTP",
"compartment":"C",
"annotation":{
"smiles":"Nc1ccn([C@@H]2O[C@H](COP(=O)([O-])OP(=O)([O-])OP(=O)([O-])O)[C@@H](O)[C@H]2O)c(=O)n1",
"smiles_sanitized":"Nc1ccn(C2OC(COP(=O)(O)OP(=O)(O)OP(=O)(O)O)C(O)C2O)c(=O)n1"
}
},
{
"id":"cpd00053_C",
"name":"L-Glutamine",
"compartment":"C",
"annotation":{
"smiles":"NC(=O)CC[C@H]([NH3+])C(=O)[O-]",
"smiles_sanitized":"NC(=O)CCC(N)C(=O)O"
}
},
{
"id":"cpd16649_C",
"name":"trans-7-carboxyhept-2-enoyl-coa",
"compartment":"C",
"annotation":{
"smiles":"",
"smiles_sanitized":""
}
},
{
"id":"cpd00932_C",
"name":"5-O--1-Carboxyvinyl-3-phosphoshikimate",
"compartment":"C",
"annotation":{
"smiles":"C=C(O[C@@H]1CC(C(=O)[O-])=C[C@@H](OP(=O)([O-])[O-])[C@H]1O)C(=O)[O-]",
"smiles_sanitized":"C=C(OC1CC(C(=O)O)=CC(OP(=O)(O)O)C1O)C(=O)O"
}
},
{
"id":"cpd00048_C",
"name":"Sulfate",
"compartment":"C",
"annotation":{
"smiles":"O=S(=O)([O-])[O-]",
"smiles_sanitized":"O=S(=O)(O)O"
}
},
{
"id":"cpd00571_E",
"name":"Glucarate",
"compartment":"E",
"annotation":{
"smiles":"O=C([O-])[C@@H](O)[C@@H](O)[C@H](O)[C@@H](O)C(=O)[O-]",
"smiles_sanitized":"O=C(O)C(O)C(O)C(O)C(O)C(=O)O"
}
},
{
"id":"cpd00327_C",
"name":"strcoa",
"compartment":"C",
"annotation":{
"smiles":"CCCCCCCCCCCCCCCCCC(=O)SCCNC(=O)CCNC(=O)[C@H](O)C(C)(C)COP(=O)([O-])OP(=O)([O-])OC[C@H]1O[C@@H](n2cnc3c(N)ncnc32)[C@H](O)[C@@H]1OP(=O)([O-])[O-]",
"smiles_sanitized":"CCCCCCCCCCCCCCCCCC(=O)SCCNC(=O)CCNC(=O)C(O)C(C)(C)COP(=O)(O)OP(=O)(O)OCC1OC(n2cnc3c(N)ncnc32)C(O)C1OP(=O)(O)O"
}
},
{
"id":"cpd00033_C",
"name":"Glycine",
"compartment":"C",
"annotation":{
"smiles":"[NH3+]CC(=O)[O-]",
"smiles_sanitized":"NCC(=O)O"
}
},
{
"id":"LIPOYL_ACP_C",
"name":"lipoyl-acp",
"compartment":"C"
},
{
"id":"cpd00020_E",
"name":"Pyruvate",
"compartment":"E",
"annotation":{
"smiles":"CC(=O)C(=O)[O-]",
"smiles_sanitized":"CC(=O)C(=O)O"
}
},
{
"id":"cpd00161_E",
"name":"L-Threonine",
"compartment":"E",
"annotation":{
"smiles":"C[C@@H](O)[C@H]([NH3+])C(=O)[O-]",
"smiles_sanitized":"CC(O)C(N)C(=O)O"
}
},
{
"id":"C12_0_2ENOYLACP_C",
"name":"trans-dodec-2-enoyl-acp",
"compartment":"C"
},
{
"id":"cpd00023_C",
"name":"L-Glutamate",
"compartment":"C",
"annotation":{
"smiles":"[NH3+][C@@H](CCC(=O)[O-])C(=O)[O-]",
"smiles_sanitized":"NC(CCC(=O)O)C(=O)O"
}
},
{
"id":"ACYL_SN_GLYCEROL_3P_TAG_C",
"name":"ACYL-SN-GLYCEROL-3P_TAG",
"compartment":"C"
},
{
"id":"C8_0_3OXOACP_C",
"name":"3-oxooctanoyl-acp",
"compartment":"C"
},
{
"id":"C4_0_ACP_C",
"name":"butanoyl-acp",
"compartment":"C"
},
{
"id":"cpd11579_C",
"name":"ethanesulfonate",
"compartment":"C",
"annotation":{
"smiles":"CCS(=O)(=O)[O-]",
"smiles_sanitized":"CCS(=O)(=O)O"
}
},
{
"id":"cpd03125_C",
"name":"(2E)-Hexenoyl-CoA",
"compartment":"C",
"annotation":{
"smiles":"CCC/C=C/C(=O)SCCNC(=O)CCNC(=O)[C@H](O)C(C)(C)COP(=O)([O-])OP(=O)([O-])OC[C@H]1O[C@@H](n2cnc3c(N)ncnc32)[C@H](O)[C@@H]1OP(=O)([O-])[O-]",
"smiles_sanitized":"CCCC=CC(=O)SCCNC(=O)CCNC(=O)C(O)C(C)(C)COP(=O)(O)OP(=O)(O)OCC1OC(n2cnc3c(N)ncnc32)C(O)C1OP(=O)(O)O"
}
},
{
"id":"C16_1_C7__3OXOACP_C",
"name":"cis-3-oxohexadec-7-enoyl-acp",
"compartment":"C"
},
{
"id":"cpd16627_C",
"name":"cis-hexadec-7-enoyl-coa",
"compartment":"C",
"annotation":{
"smiles":"",
"smiles_sanitized":""
}
},
{
"id":"cpd00198_C",
"name":"D-Xylulose5-phosphate",
"compartment":"C",
"annotation":{
"smiles":"O=C(CO)[C@@H](O)[C@H](O)COP(=O)([O-])[O-]",
"smiles_sanitized":"O=C(CO)C(O)C(O)COP(=O)(O)O"
}
},
{
"id":"C12_1_C3__ACP_C",
"name":"cis-dodec-3-enoyl-acp",
"compartment":"C"
},
{
"id":"cpd00321_C",
"name":"Feruloyl-CoA",
"compartment":"C",
"annotation":{
"smiles":"COc1cc(/C=C/C(=O)SCCNC(=O)CCNC(=O)[C@H](O)C(C)(C)COP(=O)([O-])OP(=O)([O-])OC[C@H]2O[C@@H](n3cnc4c(N)ncnc43)[C@H](O)[C@@H]2OP(=O)([O-])[O-])ccc1O",
"smiles_sanitized":"COc1cc(C=CC(=O)SCCNC(=O)CCNC(=O)C(O)C(C)(C)COP(=O)(O)OP(=O)(O)OCC2OC(n3cnc4c(N)ncnc43)C(O)C2OP(=O)(O)O)ccc1O"
}
},
{
"id":"cpd02095_C",
"name":"L-2,4-Diaminobutyrate",
"compartment":"C",
"annotation":{
"smiles":"[NH3+]CC[C@H]([NH3+])C(=O)[O-]",
"smiles_sanitized":"NCCC(N)C(=O)O"
}
},
{
"id":"cpd16639_C",
"name":"cis-octadec-9-enol",
"compartment":"C",
"annotation":{
"smiles":"",
"smiles_sanitized":""
}
},
{
"id":"cpd01088_C",
"name":"1-Octanal",
"compartment":"C",
"annotation":{
"smiles":"CCCCCCCC=O",
"smiles_sanitized":"CCCCCCCC=O"
}
},
{
"id":"cpd32253_C",
"name":"3-oxo-(9Z)-octadecenoyl-CoA",
"compartment":"C",
"annotation":{
"smiles":"CCCCCCCC/C=C\\CCCCCC(=O)CC(=O)SCCNC(=O)CCNC(=O)[C@H](O)C(C)(C)COP(=O)([O-])OP(=O)([O-])OC[C@H]1O[C@@H](n2cnc3c(N)ncnc32)[C@H](O)[C@@H]1OP(=O)([O-])[O-]",
"smiles_sanitized":"CCCCCCCCC=CCCCCCC(=O)CC(=O)SCCNC(=O)CCNC(=O)C(O)C(C)(C)COP(=O)(O)OP(=O)(O)OCC1OC(n2cnc3c(N)ncnc32)C(O)C1OP(=O)(O)O"
}
},
{
"id":"C18_0_3HYDROXYACP_C",
"name":"3-hydroxyoctadecanoyl-acp",
"compartment":"C"
},
{
"id":"cpd00182_E",
"name":"Adenosine",
"compartment":"E",
"annotation":{
"smiles":"Nc1ncnc2c1ncn2[C@@H]1O[C@H](CO)[C@@H](O)[C@H]1O",
"smiles_sanitized":"Nc1ncnc2c1ncn2C1OC(CO)C(O)C1O"
}
},
{
"id":"cpd01501_C",
"name":"2-Methylcitrate",
"compartment":"C",
"annotation":{
"smiles":"C[C@H](C(=O)[O-])[C@](O)(CC(=O)[O-])C(=O)[O-]",
"smiles_sanitized":"CC(C(=O)O)C(O)(CC(=O)O)C(=O)O"
}
},
{
"id":"cpd03117_C",
"name":"3-Oxododecanoyl-CoA",
"compartment":"C",
"annotation":{
"smiles":"CCCCCCCCCC(=O)CC(=O)SCCNC(=O)CCNC(=O)[C@H](O)C(C)(C)COP(=O)([O-])OP(=O)([O-])OC[C@H]1O[C@@H](n2cnc3c(N)ncnc32)[C@H](O)[C@@H]1OP(=O)([O-])[O-]",
"smiles_sanitized":"CCCCCCCCCC(=O)CC(=O)SCCNC(=O)CCNC(=O)C(O)C(C)(C)COP(=O)(O)OP(=O)(O)OCC1OC(n2cnc3c(N)ncnc32)C(O)C1OP(=O)(O)O"
}
},
{
"id":"cpd00094_C",
"name":"2-Oxobutyrate",
"compartment":"C",
"annotation":{
"smiles":"CCC(=O)C(=O)[O-]",
"smiles_sanitized":"CCC(=O)C(=O)O"
}
},
{
"id":"cpd00383_C",
"name":"Shikimate",
"compartment":"C",
"annotation":{
"smiles":"O=C([O-])C1=C[C@@H](O)[C@@H](O)[C@H](O)C1",
"smiles_sanitized":"O=C(O)C1=CC(O)C(O)C(O)C1"
}
},
{
"id":"TRIACYLGLYCEROL_W_C",
"name":"TRIACYLGLYCEROL_W",
"compartment":"C"
},
{
"id":"cpd00986_P",
"name":"PQQH2",
"compartment":"P",
"annotation":{
"smiles":"O=C([O-])c1cc(C(=O)[O-])c2c(n1)c(O)c(O)c1cc(C(=O)[O-])[nH]c12",
"smiles_sanitized":"O=C(O)c1cc(C(=O)O)c2c(n1)c(O)c(O)c1cc(C(=O)O)[nH]c12"
}
},
{
"id":"cpd00105_C",
"name":"D-Ribose",
"compartment":"C",
"annotation":{
"smiles":"OC[C@H]1OC(O)[C@H](O)[C@@H]1O",
"smiles_sanitized":"OCC1OC(O)C(O)C1O"
}
},
{
"id":"cpd00881_E",
"name":"Caffeate",
"compartment":"E",
"annotation":{
"smiles":"O=C([O-])/C=C/c1ccc(O)c(O)c1",
"smiles_sanitized":"O=C(O)C=Cc1ccc(O)c(O)c1"
}
},
{
"id":"cpd00050_C",
"name":"FMN",
"compartment":"C",
"annotation":{
"smiles":"Cc1cc2nc3c(=O)[n-]c(=O)nc-3n(C[C@H](O)[C@H](O)[C@H](O)COP(=O)([O-])[O-])c2cc1C",
"smiles_sanitized":"Cc1cc2nc3c(=O)[nH]c(=O)nc-3n(CC(O)C(O)C(O)COP(=O)(O)O)c2cc1C"
}
},
{
"id":"PL_W_C",
"name":"PL_W",
"compartment":"C"
},
{
"id":"cpd00249_C",
"name":"Uridine",
"compartment":"C",
"annotation":{
"smiles":"O=c1ccn([C@@H]2O[C@H](CO)[C@@H](O)[C@H]2O)c(=O)[nH]1",
"smiles_sanitized":"O=c1ccn(C2OC(CO)C(O)C2O)c(=O)[nH]1"
}
},
{
"id":"cpd00286_C",
"name":"Undecaprenylphosphate",
"compartment":"C",
"annotation":{
"smiles":"CC(C)=CCC/C(C)=C/CC/C(C)=C/CC/C(C)=C/CC/C(C)=C/CC/C(C)=C/CC/C(C)=C/CC/C(C)=C/CC/C(C)=C/CC/C(C)=C/CC/C(C)=C/COP(=O)([O-])[O-]",
"smiles_sanitized":"CC(C)=CCCC(C)=CCCC(C)=CCCC(C)=CCCC(C)=CCCC(C)=CCCC(C)=CCCC(C)=CCCC(C)=CCCC(C)=CCCC(C)=CCOP(=O)(O)O"
}
},
{
"id":"C16_0_ACP_C",
"name":"hexadecanoyl-acp",
"compartment":"C"
},
{
"id":"cpd00331_C",
"name":"cis-Aconitate",
"compartment":"C",
"annotation":{
"smiles":"O=C([O-])/C=C(/CC(=O)[O-])C(=O)[O-]",
"smiles_sanitized":"O=C(O)C=C(CC(=O)O)C(=O)O"
}
},
{
"id":"PEPTIDO_W_E",
"name":"PEPTIDO_W",
"compartment":"E"
},
{
"id":"cpd00092_C",
"name":"Uracil",
"compartment":"C",
"annotation":{
"smiles":"O=c1cc[nH]c(=O)[nH]1",
"smiles_sanitized":"O=c1cc[nH]c(=O)[nH]1"
}
},
{
"id":"cpd00309_E",
"name":"XAN",
"compartment":"E",
"annotation":{
"smiles":"O=c1[nH]c(=O)c2[nH]cnc2[nH]1",
"smiles_sanitized":"O=c1[nH]c(=O)c2[nH]cnc2[nH]1"
}
},
{
"id":"cpd16628_C",
"name":"cis-hexadec-7-enol",
"compartment":"C",
"annotation":{
"smiles":"",
"smiles_sanitized":""
}
},
{
"id":"cpd00095_C",
"name":"Glycerone-phosphate",
"compartment":"C",
"annotation":{
"smiles":"O=C(CO)COP(=O)([O-])[O-]",
"smiles_sanitized":"O=C(CO)COP(=O)(O)O"
}
},
{
"id":"GLT_TRNAS_C",
"name":"glutamate trna",
"compartment":"C"
},
{
"id":"cpd11421_C",
"name":"trdrd",
"compartment":"C",
"annotation":{
"smiles":"*[NH2+][C@@H](CS)C(=O)N[C@@H](*)C(=O)N[C@@H](*)C(=O)N[C@@H](CS)C(*)=O",
"smiles_sanitized":"*NC(CS)C(=O)NC(*)C(=O)NC(*)C(=O)NC(CS)C(*)=O"
}
},
{
"id":"cpd00842_C",
"name":"(S)-3-Hydroxybutyryl-CoA",
"compartment":"C",
"annotation":{
"smiles":"C[C@H](O)CC(=O)SCCNC(=O)CCNC(=O)[C@H](O)C(C)(C)COP(=O)([O-])OP(=O)([O-])OC[C@H]1O[C@@H](n2cnc3c(N)ncnc32)[C@H](O)[C@@H]1OP(=O)([O-])[O-]",
"smiles_sanitized":"CC(O)CC(=O)SCCNC(=O)CCNC(=O)C(O)C(C)(C)COP(=O)(O)OP(=O)(O)OCC1OC(n2cnc3c(N)ncnc32)C(O)C1OP(=O)(O)O"
}
},
{
"id":"cpd00136_C",
"name":"4-Hydroxybenzoate",
"compartment":"C",
"annotation":{
"smiles":"O=C([O-])c1ccc(O)cc1",
"smiles_sanitized":"O=C(O)c1ccc(O)cc1"
}
},
{
"id":"C14_1_C5__2ENOYLACP_C",
"name":"trans,cis-tetradec-2,4-enoyl-acp",
"compartment":"C"
},
{
"id":"cpd00340_C",
"name":"2,5-Dioxopentanoate",
"compartment":"C",
"annotation":{
"smiles":"O=CCCC(=O)C(=O)[O-]",
"smiles_sanitized":"O=CCCC(=O)C(=O)O"
}
},
{
"id":"cpd00130_E",
"name":"L-Malate",
"compartment":"E",
"annotation":{
"smiles":"O=C([O-])C[C@H](O)C(=O)[O-]",
"smiles_sanitized":"O=C(O)CC(O)C(=O)O"
}
},
{
"id":"cpd00007_C",
"name":"O2",
"compartment":"C",
"annotation":{
"smiles":"O=O",
"smiles_sanitized":"O=O"
}
},
{
"id":"cpd00616_C",
"name":"Pretyrosine",
"compartment":"C",
"annotation":{
"smiles":"[NH3+][C@@H](CC1(C(=O)[O-])C=CC(O)C=C1)C(=O)[O-]",
"smiles_sanitized":"NC(CC1(C(=O)O)C=CC(O)C=C1)C(=O)O"
}
},
{
"id":"cpd00800_C",
"name":"8-Amino-7-oxononanoate",
"compartment":"C",
"annotation":{
"smiles":"CC([NH3+])C(=O)CCCCCC(=O)[O-]",
"smiles_sanitized":"CC(N)C(=O)CCCCCC(=O)O"
}
},
{
"id":"C12_0_3OXOACP_C",
"name":"3-oxododecanoyl-acp",
"compartment":"C"
},
{
"id":"cpd00070_C",
"name":"Malonyl-CoA",
"compartment":"C",
"annotation":{
"smiles":"CC(C)(COP(=O)([O-])OP(=O)([O-])OC[C@H]1O[C@@H](n2cnc3c(N)ncnc32)[C@H](O)[C@@H]1OP(=O)([O-])[O-])[C@@H](O)C(=O)NCCC(=O)NCCSC(=O)CC(=O)[O-]",
"smiles_sanitized":"CC(C)(COP(=O)(O)OP(=O)(O)OCC1OC(n2cnc3c(N)ncnc32)C(O)C1OP(=O)(O)O)C(O)C(=O)NCCC(=O)NCCSC(=O)CC(=O)O"
}
},
{
"id":"C10_0_3OXOACP_C",
"name":"3-oxodecanoyl-acp",
"compartment":"C"
},
{
"id":"cpd00115_C",
"name":"dATP",
"compartment":"C",
"annotation":{
"smiles":"Nc1ncnc2c1ncn2[C@H]1C[C@H](O)[C@@H](COP(=O)([O-])OP(=O)([O-])OP(=O)([O-])O)O1",
"smiles_sanitized":"Nc1ncnc2c1ncn2C1CC(O)C(COP(=O)(O)OP(=O)(O)OP(=O)(O)O)O1"
}
},
{
"id":"cpd00355_C",
"name":"Nicotinamide ribonucleotide",
"compartment":"C",
"annotation":{
"smiles":"NC(=O)c1ccc[n+]([C@@H]2O[C@H](COP(=O)([O-])[O-])[C@@H](O)[C@H]2O)c1",
"smiles_sanitized":"NC(=O)c1ccc[n+](C2OC(COP(=O)(O)O)C(O)C2O)c1"
}
},
{
"id":"cpd00203_C",
"name":"1,3-Bisphospho-D-glycerate",
"compartment":"C",
"annotation":{
"smiles":"O=C(OP(=O)([O-])[O-])[C@H](O)COP(=O)([O-])[O-]",
"smiles_sanitized":"O=C(OP(=O)(O)O)C(O)COP(=O)(O)O"
}
},
{
"id":"cpd00141_C",
"name":"Propionate",
"compartment":"C",
"annotation":{
"smiles":"CCC(=O)[O-]",
"smiles_sanitized":"CCC(=O)O"
}
},
{
"id":"UDP_POLYSACCH_LPS_C",
"name":"UDP-POLYSACCH_LPS",
"compartment":"C"
},
{
"id":"cpd02894_C",
"name":"4-Amino-2-methyl-5-diphosphomethylpyrimidine",
"compartment":"C",
"annotation":{
"smiles":"Cc1nc(N)c(COP(=O)([O-])OP(=O)([O-])[O-])c[nH+]1",
"smiles_sanitized":"Cc1ncc(COP(=O)(O)OP(=O)(O)O)c(N)n1"
}
},
{
"id":"cpd00120_C",
"name":"Butyryl-CoA",
"compartment":"C",
"annotation":{
"smiles":"CCCC(=O)SCCNC(=O)CCNC(=O)[C@H](O)C(C)(C)COP(=O)([O-])OP(=O)([O-])OC[C@@H]1O[C@H](n2cnc3c(N)ncnc32)C(O)C1OP(=O)([O-])[O-]",
"smiles_sanitized":"CCCC(=O)SCCNC(=O)CCNC(=O)C(O)C(C)(C)COP(=O)(O)OP(=O)(O)OCC1OC(n2cnc3c(N)ncnc32)C(O)C1OP(=O)(O)O"
}
},
{
"id":"cpd00738_C",
"name":"phosphoserine",
"compartment":"C",
"annotation":{
"smiles":"[NH3+][C@@H](COP(=O)([O-])[O-])C(=O)[O-]",
"smiles_sanitized":"NC(COP(=O)(O)O)C(=O)O"
}
},
{
"id":"CHARGED_LYS_TRNAS_C",
"name":"charged lysine trna",
"compartment":"C"
},
{
"id":"cpd00119_C",
"name":"L-Histidine",
"compartment":"C",
"annotation":{
"smiles":"[NH3+][C@@H](Cc1c[nH]cn1)C(=O)[O-]",
"smiles_sanitized":"NC(Cc1c[nH]cn1)C(=O)O"
}
},
{
"id":"cpd00202_C",
"name":"DMAPP",
"compartment":"C",
"annotation":{
"smiles":"CC(C)=CCOP(=O)([O-])OP(=O)([O-])O",
"smiles_sanitized":"CC(C)=CCOP(=O)(O)OP(=O)(O)O"
}
},
{
"id":"cpd00719_C",
"name":"GDP-D-mannuronate",
"compartment":"C",
"annotation":{
"smiles":"Nc1nc2c(ncn2C2O[C@H](COP(=O)([O-])OP(=O)([O-])O[C@H]3O[C@H](C(=O)[O-])[C@@H](O)[C@H](O)[C@@H]3O)[C@@H](O)[C@H]2O)c(=O)[nH]1",
"smiles_sanitized":"Nc1nc2c(ncn2C2OC(COP(=O)(O)OP(=O)(O)OC3OC(C(=O)O)C(O)C(O)C3O)C(O)C2O)c(=O)[nH]1"
}
},
{
"id":"cpd00061_C",
"name":"Phosphoenolpyruvate",
"compartment":"C",
"annotation":{
"smiles":"C=C(OP(=O)([O-])[O-])C(=O)[O-]",
"smiles_sanitized":"C=C(OP(=O)(O)O)C(=O)O"
}
},
{
"id":"PHE_TRNAS_C",
"name":"phenylalanine trna",
"compartment":"C"
},
{
"id":"cpd00075_C",
"name":"Nitrite",
"compartment":"C",
"annotation":{
"smiles":"O=NO",
"smiles_sanitized":"O=NO"
}
},
{
"id":"cpd00248_P",
"name":"Quinate",
"compartment":"P",
"annotation":{
"smiles":"O=C([O-])[C@]1(O)C[C@@H](O)[C@@H](O)[C@H](O)C1",
"smiles_sanitized":"O=C(O)C1(O)CC(O)C(O)C(O)C1"
}
},
{
"id":"cpd01507_C",
"name":"3-Oxoadipyl-CoA",
"compartment":"C",
"annotation":{
"smiles":"CC(C)(COP(=O)([O-])OP(=O)([O-])OC[C@H]1O[C@@H](n2cnc3c(N)ncnc32)[C@H](O)[C@@H]1OP(=O)([O-])[O-])[C@@H](O)C(=O)NCCC(=O)NCCSC(=O)CC(=O)CCC(=O)[O-]",
"smiles_sanitized":"CC(C)(COP(=O)(O)OP(=O)(O)OCC1OC(n2cnc3c(N)ncnc32)C(O)C1OP(=O)(O)O)C(O)C(=O)NCCC(=O)NCCSC(=O)CC(=O)CCC(=O)O"
}
},
{
"id":"cpd16638_C",
"name":"cis-octadec-9-enoyl-coa",
"compartment":"C",
"annotation":{
"smiles":"CCCCCCCC/C=C\\CCCCCCCC(=O)SCCNC(=O)CCNC(=O)[C@H](O)C(C)(C)COP(=O)([O-])OP(=O)([O-])OC[C@H]1O[C@@H](n2cnc3c(N)ncnc32)[C@H](O)[C@@H]1OP(=O)([O-])[O-]",
"smiles_sanitized":"CCCCCCCCC=CCCCCCCCC(=O)SCCNC(=O)CCNC(=O)C(O)C(C)(C)COP(=O)(O)OP(=O)(O)OCC1OC(n2cnc3c(N)ncnc32)C(O)C1OP(=O)(O)O"
}
},
{
"id":"cpd03129_C",
"name":"(2E)-Decenoyl-CoA",
"compartment":"C",
"annotation":{
"smiles":"CCCCCCC/C=C/C(=O)SCCNC(=O)CCNC(=O)[C@H](O)C(C)(C)COP(=O)([O-])OP(=O)([O-])OC[C@H]1O[C@@H](n2cnc3c(N)ncnc32)[C@H](O)[C@@H]1OP(=O)([O-])[O-]",
"smiles_sanitized":"CCCCCCCC=CC(=O)SCCNC(=O)CCNC(=O)C(O)C(C)(C)COP(=O)(O)OP(=O)(O)OCC1OC(n2cnc3c(N)ncnc32)C(O)C1OP(=O)(O)O"
}
},
{
"id":"HIS_TRNAS_C",
"name":"histidine trna",
"compartment":"C"
},
{
"id":"cpd16507_C",
"name":"Protocatechualdehyde",
"compartment":"C",
"annotation":{
"smiles":"O=Cc1ccc(O)c(O)c1",
"smiles_sanitized":"O=Cc1ccc(O)c(O)c1"
}
},
{
"id":"cpd33217_C",
"name":"3-(3,4-dihydroxyphenyl)-3-hydroxypropanoate",
"compartment":"C",
"annotation":{
"smiles":"CC(C)(COP(=O)([O-])OP(=O)([O-])OC[C@H]1O[C@@H](n2cnc3c(N)ncnc32)[C@H](O)[C@@H]1OP(=O)([O-])[O-])[C@@H](O)C(=O)NCCC(=O)NCCSC(=O)CC(O)c1ccc(O)c(O)c1",
"smiles_sanitized":"CC(C)(COP(=O)(O)OP(=O)(O)OCC1OC(n2cnc3c(N)ncnc32)C(O)C1OP(=O)(O)O)C(O)C(=O)NCCC(=O)NCCSC(=O)CC(O)c1ccc(O)c(O)c1"
}
},
{
"id":"cpd00998_E",
"name":"Octane",
"compartment":"E",
"annotation":{
"smiles":"CCCCCCCC",
"smiles_sanitized":"CCCCCCCC"
}
},
{
"id":"cpd00169_C",
"name":"3-Phosphoglycerate",
"compartment":"C",
"annotation":{
"smiles":"O=C([O-])[C@H](O)COP(=O)([O-])[O-]",
"smiles_sanitized":"O=C(O)C(O)COP(=O)(O)O"
}
},
{
"id":"cpd23287_C",
"name":"3-(4-hydroxyphenyl)-3-hydroxy-propanoyl-CoA",
"compartment":"C",
"annotation":{
"smiles":"CC(C)(COP(=O)([O-])OP(=O)([O-])OC[C@H]1O[C@@H](n2cnc3c(N)ncnc32)[C@H](O)[C@@H]1OP(=O)([O-])[O-])[C@@H](O)C(=O)NCCC(=O)NCCSC(=O)C[C@H](O)c1ccc(O)cc1",
"smiles_sanitized":"CC(C)(COP(=O)(O)OP(=O)(O)OCC1OC(n2cnc3c(N)ncnc32)C(O)C1OP(=O)(O)O)C(O)C(=O)NCCC(=O)NCCSC(=O)CC(O)c1ccc(O)cc1"
}
},
{
"id":"cpd00222_C",
"name":"GLCN",
"compartment":"C",
"annotation":{
"smiles":"O=C([O-])[C@H](O)[C@@H](O)[C@H](O)[C@H](O)CO",
"smiles_sanitized":"O=C(O)C(O)C(O)C(O)C(O)CO"
}
},
{
"id":"cpd00300_C",
"name":"Urate",
"compartment":"C",
"annotation":{
"smiles":"O=c1[nH]c(=O)c2[nH]c(=O)[nH]c2[nH]1",
"smiles_sanitized":"O=c1[nH]c(=O)c2[nH]c(=O)[nH]c2[nH]1"
}
},
{
"id":"C14_1_C5__3OXOACP_C",
"name":"cis-3-oxotetradec-4-enoyl-acp",
"compartment":"C"
},
{
"id":"cpd02884_C",
"name":"FAICAR",
"compartment":"C",
"annotation":{
"smiles":"NC(=O)c1ncn([C@@H]2O[C@H](COP(=O)([O-])[O-])[C@@H](O)[C@H]2O)c1NC=O",
"smiles_sanitized":"NC(=O)c1ncn(C2OC(COP(=O)(O)O)C(O)C2O)c1NC=O"
}
},
{
"id":"cpd00143_C",
"name":"Phenylpyruvate",
"compartment":"C",
"annotation":{
"smiles":"O=C([O-])C(=O)Cc1ccccc1",
"smiles_sanitized":"O=C(O)C(=O)Cc1ccccc1"
}
},
{
"id":"cpd00654_C",
"name":"Deoxycytidine",
"compartment":"C",
"annotation":{
"smiles":"Nc1ccn([C@H]2C[C@H](O)[C@@H](CO)O2)c(=O)n1",
"smiles_sanitized":"Nc1ccn(C2CC(O)C(CO)O2)c(=O)n1"
}
},
{
"id":"cpd00025_C",
"name":"H2O2",
"compartment":"C",
"annotation":{
"smiles":"OO",
"smiles_sanitized":"OO"
}
},
{
"id":"cpd00101_C",
"name":"ribose-5-phosphate",
"compartment":"C",
"annotation":{
"smiles":"O=P([O-])([O-])OC[C@H]1OC(O)[C@H](O)[C@@H]1O",
"smiles_sanitized":"O=P(O)(O)OCC1OC(O)C(O)C1O"
}
},
{
"id":"cpd00211_E",
"name":"Butyrate",
"compartment":"E",
"annotation":{
"smiles":"CCCC(=O)[O-]",
"smiles_sanitized":"CCCC(=O)O"
}
},
{
"id":"cpd00197_C",
"name":"Protocatechuate",
"compartment":"C",
"annotation":{
"smiles":"O=C([O-])c1ccc(O)c(O)c1",
"smiles_sanitized":"O=C(O)c1ccc(O)c(O)c1"
}
},
{
"id":"cpd02993_C",
"name":"Lipid IV(A)",
"compartment":"C",
"annotation":{
"smiles":"CCCCCCCCCCC[C@@H](O)CC(=O)N[C@H]1[C@@H](OP(=O)([O-])[O-])O[C@H](CO[C@@H]2O[C@H](CO)[C@@H](OP(=O)([O-])[O-])[C@H](OC(=O)C[C@H](O)CCCCCCCCCCC)[C@H]2NC(=O)C[C@H](O)CCCCCCCCCCC)[C@@H](O)[C@@H]1OC(=O)C[C@H](O)CCCCCCCCCCC",
"smiles_sanitized":"CCCCCCCCCCCC(O)CC(=O)NC1C(OP(=O)(O)O)OC(COC2OC(CO)C(OP(=O)(O)O)C(OC(=O)CC(O)CCCCCCCCCCC)C2NC(=O)CC(O)CCCCCCCCCCC)C(O)C1OC(=O)CC(O)CCCCCCCCCCC"
}
},
{
"id":"cpd00014_C",
"name":"UDP",
"compartment":"C",
"annotation":{
"smiles":"O=c1ccn([C@@H]2O[C@H](COP(=O)([O-])OP(=O)([O-])O)[C@@H](O)[C@H]2O)c(=O)[nH]1",
"smiles_sanitized":"O=c1ccn(C2OC(COP(=O)(O)OP(=O)(O)O)C(O)C2O)c(=O)[nH]1"
}
},
{
"id":"ACETYL_ACP_C",
"name":"acetyl-acp",
"compartment":"C"
},
{
"id":"cpd00654_E",
"name":"Deoxycytidine",
"compartment":"E",
"annotation":{
"smiles":"Nc1ccn([C@H]2C[C@H](O)[C@@H](CO)O2)c(=O)n1",
"smiles_sanitized":"Nc1ccn(C2CC(O)C(CO)O2)c(=O)n1"
}
},
{
"id":"ARG_TRNAS_C",
"name":"arginine trna",
"compartment":"C"
},
{
"id":"cpd00793_C",
"name":"Thiamine phosphate",
"compartment":"C",
"annotation":{
"smiles":"Cc1nc(N)c(C[n+]2csc(CCOP(=O)([O-])[O-])c2C)c[nH+]1",
"smiles_sanitized":"Cc1ncc(C[n+]2csc(CCOP(=O)(O)O)c2C)c(N)n1"
}
},
{
"id":"cpd01107_E",
"name":"Decanoate",
"compartment":"E",
"annotation":{
"smiles":"CCCCCCCCCC(=O)[O-]",
"smiles_sanitized":"CCCCCCCCCC(=O)O"
}
},
{
"id":"L_PHOSPHATIDATE_TAG_C",
"name":"L-PHOSPHATIDATE_TAG",
"compartment":"C"
},
{
"id":"cpd16647_C",
"name":"trans-3-oxohex-4-enoyl-coa",
"compartment":"C",
"annotation":{
"smiles":"",
"smiles_sanitized":""
}
},
{
"id":"ACYL_SN_GLYCEROL_3P_PL_C",
"name":"ACYL-SN-GLYCEROL-3P_PL",
"compartment":"C"
},
{
"id":"cpd00055_C",
"name":"Formaldehyde",
"compartment":"C",
"annotation":{
"smiles":"C=O",
"smiles_sanitized":"C=O"
}
},
{
"id":"cpd03446_C",
"name":"2-Octaprenyl-6-methoxyphenol",
"compartment":"C",
"annotation":{
"smiles":"COc1cccc(C/C=C(\\C)CC/C=C(\\C)CC/C=C(\\C)CC/C=C(\\C)CC/C=C(\\C)CC/C=C(\\C)CC/C=C(\\C)CCC=C(C)C)c1O",
"smiles_sanitized":"COc1cccc(CC=C(C)CCC=C(C)CCC=C(C)CCC=C(C)CCC=C(C)CCC=C(C)CCC=C(C)CCC=C(C)C)c1O"
}
},
{
"id":"cpd01850_C",
"name":"L-Sorbose 1P",
"compartment":"C",
"annotation":{
"smiles":"O=C(COP(=O)([O-])[O-])[C@@H](O)[C@H](O)[C@@H](O)CO",
"smiles_sanitized":"O=C(COP(=O)(O)O)C(O)C(O)C(O)CO"
}
},
{
"id":"cpd00834_C",
"name":"Phosphopantetheine",
"compartment":"C",
"annotation":{
"smiles":"CC(C)(COP(=O)([O-])[O-])[C@@H](O)C(=O)NCCC(=O)NCCS",
"smiles_sanitized":"CC(C)(COP(=O)(O)O)C(O)C(=O)NCCC(=O)NCCS"
}
},
{
"id":"cpd00794_C",
"name":"TRHL",
"compartment":"C",
"annotation":{
"smiles":"OC[C@H]1O[C@H](O[C@H]2O[C@H](CO)[C@@H](O)[C@H](O)[C@H]2O)[C@H](O)[C@@H](O)[C@@H]1O",
"smiles_sanitized":"OCC1OC(OC2OC(CO)C(O)C(O)C2O)C(O)C(O)C1O"
}
},
{
"id":"cpd00428_C",
"name":"2-Oxopropanal",
"compartment":"C",
"annotation":{
"smiles":"CC(=O)C=O",
"smiles_sanitized":"CC(=O)C=O"
}
},
{
"id":"cpd14936_C",
"name":"3-Hydroxystearoyl-CoA",
"compartment":"C",
"annotation":{
"smiles":"CCCCCCCCCCCCCCCC(O)CC(=O)SCCNC(=O)CCNC(=O)[C@H](O)C(C)(C)COP(=O)([O-])OP(=O)([O-])OC[C@H]1O[C@@H](n2cnc3c(N)ncnc32)[C@H](O)[C@@H]1OP(=O)([O-])[O-]",
"smiles_sanitized":"CCCCCCCCCCCCCCCC(O)CC(=O)SCCNC(=O)CCNC(=O)C(O)C(C)(C)COP(=O)(O)OP(=O)(O)OCC1OC(n2cnc3c(N)ncnc32)C(O)C1OP(=O)(O)O"
}
},
{
"id":"cpd02201_C",
"name":"4-phosphopantothenate",
"compartment":"C",
"annotation":{
"smiles":"CC(C)(COP(=O)([O-])[O-])[C@@H](O)C(=O)NCCC(=O)[O-]",
"smiles_sanitized":"CC(C)(COP(=O)(O)O)C(O)C(=O)NCCC(=O)O"
}
},
{
"id":"cpd00141_E",
"name":"Propionate",
"compartment":"E",
"annotation":{
"smiles":"CCC(=O)[O-]",
"smiles_sanitized":"CCC(=O)O"
}
},
{
"id":"cpd01630_E",
"name":"cis,cis-Muconate",
"compartment":"E",
"annotation":{
"smiles":"O=C([O-])/C=C\\C=C/C(=O)[O-]",
"smiles_sanitized":"O=C(O)C=CC=CC(=O)O"
}
},
{
"id":"cpd00134_C",
"name":"Palmitoyl-CoA",
"compartment":"C",
"annotation":{
"smiles":"CCCCCCCCCCCCCCCC(=O)SCCNC(=O)CCNC(=O)[C@H](O)C(C)(C)COP(=O)([O-])OP(=O)([O-])OC[C@H]1O[C@@H](n2cnc3c(N)ncnc32)[C@H](O)[C@@H]1OP(=O)([O-])[O-]",
"smiles_sanitized":"CCCCCCCCCCCCCCCC(=O)SCCNC(=O)CCNC(=O)C(O)C(C)(C)COP(=O)(O)OP(=O)(O)OCC1OC(n2cnc3c(N)ncnc32)C(O)C1OP(=O)(O)O"
}
},
{
"id":"AFATTYACID_TAG_C",
"name":"AFATTYACID_TAG",
"compartment":"C"
},
{
"id":"cpd16657_E",
"name":"ethylsalicylate",
"compartment":"E",
"annotation":{
"smiles":"",
"smiles_sanitized":""
}
},
{
"id":"cpd00182_C",
"name":"Adenosine",
"compartment":"C",
"annotation":{
"smiles":"Nc1ncnc2c1ncn2[C@@H]1O[C@H](CO)[C@@H](O)[C@H]1O",
"smiles_sanitized":"Nc1ncnc2c1ncn2C1OC(CO)C(O)C1O"
}
},
{
"id":"cpd00355_E",
"name":"Nicotinamide ribonucleotide",
"compartment":"E",
"annotation":{
"smiles":"NC(=O)c1ccc[n+]([C@@H]2O[C@H](COP(=O)([O-])[O-])[C@@H](O)[C@H]2O)c1",
"smiles_sanitized":"NC(=O)c1ccc[n+](C2OC(COP(=O)(O)O)C(O)C2O)c1"
}
},
{
"id":"CHARGED_ASP_TRNAS_C",
"name":"charged aspartate trna",
"compartment":"C"
},
{
"id":"cpd00162_E",
"name":"Aminoethanol",
"compartment":"E",
"annotation":{
"smiles":"[NH3+]CCO",
"smiles_sanitized":"NCCO"
}
},
{
"id":"cpd00597_E",
"name":"Valerate",
"compartment":"E",
"annotation":{
"smiles":"CCCCC(=O)[O-]",
"smiles_sanitized":"CCCCC(=O)O"
}
},
{
"id":"cpd00581_C",
"name":"Urocanate",
"compartment":"C",
"annotation":{
"smiles":"O=C([O-])/C=C/c1c[nH]cn1",
"smiles_sanitized":"O=C(O)C=Cc1c[nH]cn1"
}
},
{
"id":"cpd01982_C",
"name":"5-Phosphoribosylamine",
"compartment":"C",
"annotation":{
"smiles":"[NH3+][C@@H]1O[C@H](COP(=O)([O-])[O-])[C@@H](O)[C@H]1O",
"smiles_sanitized":"NC1OC(COP(=O)(O)O)C(O)C1O"
}
},
{
"id":"cpd01947_C",
"name":"BDOH",
"compartment":"C",
"annotation":{
"smiles":"C[C@@H](O)[C@@H](C)O",
"smiles_sanitized":"CC(O)C(C)O"
}
},
{
"id":"cpd00246_C",
"name":"Inosine",
"compartment":"C",
"annotation":{
"smiles":"O=c1[nH]cnc2c1ncn2[C@@H]1O[C@H](CO)[C@@H](O)[C@H]1O",
"smiles_sanitized":"O=c1[nH]cnc2c1ncn2C1OC(CO)C(O)C1O"
}
},
{
"id":"cpd00447_C",
"name":"Betaine aldehyde",
"compartment":"C",
"annotation":{
"smiles":"C[N+](C)(C)CC=O",
"smiles_sanitized":"C[N+](C)(C)CC=O"
}
},
{
"id":"cpd00830_C",
"name":"4-Hydroxy-2-oxoglutarate",
"compartment":"C",
"annotation":{
"smiles":"O=C([O-])C(=O)CC(O)C(=O)[O-]",
"smiles_sanitized":"O=C(O)C(=O)CC(O)C(=O)O"
}
},
{
"id":"cpd00229_C",
"name":"Glycolaldehyde",
"compartment":"C",
"annotation":{
"smiles":"O=CCO",
"smiles_sanitized":"O=CCO"
}
},
{
"id":"CHARGED_MET_TRNAS_C",
"name":"charged methionine trna",
"compartment":"C"
},
{
"id":"cpd00277_E",
"name":"Deoxyguanosine",
"compartment":"E",
"annotation":{
"smiles":"Nc1nc2c(ncn2[C@H]2C[C@H](O)[C@@H](CO)O2)c(=O)[nH]1",
"smiles_sanitized":"Nc1nc2c(ncn2C2CC(O)C(CO)O2)c(=O)[nH]1"
}
},
{
"id":"cpd00773_C",
"name":"UDP-MurNAc",
"compartment":"C",
"annotation":{
"smiles":"CC(=O)N[C@H]1[C@@H](OP(=O)([O-])OP(=O)([O-])OC[C@H]2O[C@@H](n3ccc(=O)[nH]c3=O)[C@H](O)[C@@H]2O)O[C@H](CO)[C@@H](O)[C@@H]1O[C@H](C)C(=O)[O-]",
"smiles_sanitized":"CC(=O)NC1C(OP(=O)(O)OP(=O)(O)OCC2OC(n3ccc(=O)[nH]c3=O)C(O)C2O)OC(CO)C(O)C1OC(C)C(=O)O"
}
},
{
"id":"cpd02498_C",
"name":"2,3-Dihydroxy-isovalerate",
"compartment":"C",
"annotation":{
"smiles":"CC(C)(O)C(O)C(=O)[O-]",
"smiles_sanitized":"CC(C)(O)C(O)C(=O)O"
}
},
{
"id":"cpd02611_C",
"name":"N-Acetyl-D-glucosamine1-phosphate",
"compartment":"C",
"annotation":{
"smiles":"CC(=O)N[C@@H]1C(OP(=O)([O-])[O-])O[C@H](CO)[C@@H](O)[C@@H]1O",
"smiles_sanitized":"CC(=O)NC1C(OP(=O)(O)O)OC(CO)C(O)C1O"
}
},
{
"id":"cpd03470_C",
"name":"Iminoaspartate",
"compartment":"C",
"annotation":{
"smiles":"[NH2+]=C(CC(=O)[O-])C(=O)[O-]",
"smiles_sanitized":"N=C(CC(=O)O)C(=O)O"
}
},
{
"id":"cpd00626_C",
"name":"dTDPglucose",
"compartment":"C",
"annotation":{
"smiles":"Cc1cn([C@H]2C[C@H](O)[C@@H](COP(=O)([O-])OP(=O)([O-])O[C@H]3O[C@H](CO)[C@@H](O)[C@H](O)[C@H]3O)O2)c(=O)[nH]c1=O",
"smiles_sanitized":"Cc1cn(C2CC(O)C(COP(=O)(O)OP(=O)(O)OC3OC(CO)C(O)C(O)C3O)O2)c(=O)[nH]c1=O"
}
},
{
"id":"C18_0_3OXOACP_C",
"name":"3-oxooctadecanoyl-acp",
"compartment":"C"
},
{
"id":"cpd23920_C",
"name":"(R)-3-hydroxyvaleryl-CoA",
"compartment":"C",
"annotation":{
"smiles":"CC[C@@H](O)CC(=O)SCCNC(=O)CCNC(=O)C(O)C(C)(C)COP(=O)([O-])OP(=O)([O-])OC[C@H]1O[C@@H](n2cnc3c(N)ncnc32)[C@H](O)[C@@H]1OP(=O)([O-])[O-]",
"smiles_sanitized":"CCC(O)CC(=O)SCCNC(=O)CCNC(=O)C(O)C(C)(C)COP(=O)(O)OP(=O)(O)OCC1OC(n2cnc3c(N)ncnc32)C(O)C1OP(=O)(O)O"
}
},
{
"id":"cpd02536_C",
"name":"Muconolactone",
"compartment":"C",
"annotation":{
"smiles":"O=C([O-])CC1C=CC(=O)O1",
"smiles_sanitized":"O=C(O)CC1C=CC(=O)O1"
}
},
{
"id":"cpd00484_C",
"name":"p-Hydroxybenzaldehyde",
"compartment":"C",
"annotation":{
"smiles":"O=Cc1ccc(O)cc1",
"smiles_sanitized":"O=Cc1ccc(O)cc1"
}
},
{
"id":"cpd00637_C",
"name":"D-Methionine",
"compartment":"C",
"annotation":{
"smiles":"CSCC[C@@H]([NH3+])C(=O)[O-]",
"smiles_sanitized":"CSCCC(N)C(=O)O"
}
},
{
"id":"cpd00080_C",
"name":"Glycerol-3-phosphate",
"compartment":"C",
"annotation":{
"smiles":"O=P([O-])([O-])OC[C@H](O)CO",
"smiles_sanitized":"O=P(O)(O)OCC(O)CO"
}
},
{
"id":"cpd00890_C",
"name":"UDP-N-acetylmuramoyl-L-alanine",
"compartment":"C",
"annotation":{
"smiles":"CC(=O)N[C@H]1[C@@H](OP(=O)([O-])OP(=O)([O-])OC[C@H]2O[C@@H](n3ccc(=O)[nH]c3=O)[C@H](O)[C@@H]2O)O[C@H](CO)[C@@H](O)[C@@H]1O[C@H](C)C(=O)N[C@@H](C)C(=O)[O-]",
"smiles_sanitized":"CC(=O)NC1C(OP(=O)(O)OP(=O)(O)OCC2OC(n3ccc(=O)[nH]c3=O)C(O)C2O)OC(CO)C(O)C1OC(C)C(=O)NC(C)C(=O)O"
}
},
{
"id":"cpd00136_E",
"name":"4-Hydroxybenzoate",
"compartment":"E",
"annotation":{
"smiles":"O=C([O-])c1ccc(O)cc1",
"smiles_sanitized":"O=C(O)c1ccc(O)cc1"
}
},
{
"id":"cpd08449_C",
"name":"3-Amino-2-oxopropyl phosphate",
"compartment":"C",
"annotation":{
"smiles":"[NH3+]CC(=O)COP(=O)([O-])[O-]",
"smiles_sanitized":"NCC(=O)COP(=O)(O)O"
}
},
{
"id":"cpd00036_E",
"name":"Succinate",
"compartment":"E",
"annotation":{
"smiles":"O=C([O-])CCC(=O)[O-]",
"smiles_sanitized":"O=C(O)CCC(=O)O"
}
},
{
"id":"cpd00536_C",
"name":"Oleate",
"compartment":"C",
"annotation":{
"smiles":"CCCCCCCC/C=C\\CCCCCCCC(=O)[O-]",
"smiles_sanitized":"CCCCCCCCC=CCCCCCCCC(=O)O"
}
},
{
"id":"cpd01741_E",
"name":"ddca",
"compartment":"E",
"annotation":{
"smiles":"CCCCCCCCCCCC(=O)[O-]",
"smiles_sanitized":"CCCCCCCCCCCC(=O)O"
}
},
{
"id":"APS_C",
"name":"aps",
"compartment":"C"
},
{
"id":"cpd03279_E",
"name":"Deoxyinosine",
"compartment":"E",
"annotation":{
"smiles":"O=c1[nH]cnc2c1ncn2[C@H]1C[C@H](O)[C@@H](CO)O1",
"smiles_sanitized":"O=c1[nH]cnc2c1ncn2C1CC(O)C(CO)O1"
}
},
{
"id":"cpd00017_C",
"name":"S-Adenosyl-L-methionine",
"compartment":"C",
"annotation":{
"smiles":"C[S+](CC[C@H]([NH3+])C(=O)[O-])C[C@H]1O[C@@H](n2cnc3c(N)ncnc32)[C@H](O)[C@@H]1O",
"smiles_sanitized":"C[S+](CCC(N)C(=O)O)CC1OC(n2cnc3c(N)ncnc32)C(O)C1O"
}
},
{
"id":"cpd00859_C",
"name":"Pseudouridine 5'-phosphate",
"compartment":"C",
"annotation":{
"smiles":"O=c1[nH]cc([C@@H]2O[C@H](COP(=O)([O-])[O-])[C@@H](O)[C@H]2O)c(=O)[nH]1",
"smiles_sanitized":"O=c1[nH]cc(C2OC(COP(=O)(O)O)C(O)C2O)c(=O)[nH]1"
}
},
{
"id":"cpd00540_E",
"name":"BET",
"compartment":"E",
"annotation":{
"smiles":"C[N+](C)(C)CC(=O)[O-]",
"smiles_sanitized":"C[N+](C)(C)CC(=O)O"
}
},
{
"id":"cpd00071_C",
"name":"Acetaldehyde",
"compartment":"C",
"annotation":{
"smiles":"CC=O",
"smiles_sanitized":"CC=O"
}
},
{
"id":"cpd00562_C",
"name":"1-Octanol",
"compartment":"C",
"annotation":{
"smiles":"CCCCCCCCO",
"smiles_sanitized":"CCCCCCCCO"
}
},
{
"id":"cpd00069_C",
"name":"L-Tyrosine",
"compartment":"C",
"annotation":{
"smiles":"[NH3+][C@@H](Cc1ccc(O)cc1)C(=O)[O-]",
"smiles_sanitized":"NC(Cc1ccc(O)cc1)C(=O)O"
}
},
{
"id":"cpd00668_C",
"name":"ALCTT",
"compartment":"C",
"annotation":{
"smiles":"CC(=O)C(C)(O)C(=O)[O-]",
"smiles_sanitized":"CC(=O)C(C)(O)C(=O)O"
}
},
{
"id":"cpd00308_E",
"name":"Malonate",
"compartment":"E",
"annotation":{
"smiles":"O=C([O-])CC(=O)[O-]",
"smiles_sanitized":"O=C(O)CC(=O)O"
}
},
{
"id":"cpd00637_E",
"name":"D-Methionine",
"compartment":"E",
"annotation":{
"smiles":"CSCC[C@@H]([NH3+])C(=O)[O-]",
"smiles_sanitized":"CSCCC(N)C(=O)O"
}
},
{
"id":"cpd05193_C",
"name":"suberate",
"compartment":"C",
"annotation":{
"smiles":"O=C([O-])CCCCCCC(=O)[O-]",
"smiles_sanitized":"O=C(O)CCCCCCC(=O)O"
}
},
{
"id":"cpd03130_C",
"name":"(2E)-Octenoyl-CoA",
"compartment":"C",
"annotation":{
"smiles":"CCCCC/C=C/C(=O)SCCNC(=O)CCNC(=O)[C@H](O)C(C)(C)COP(=O)([O-])OP(=O)([O-])OC[C@H]1O[C@@H](n2cnc3c(N)ncnc32)[C@H](O)[C@@H]1OP(=O)([O-])[O-]",
"smiles_sanitized":"CCCCCC=CC(=O)SCCNC(=O)CCNC(=O)C(O)C(C)(C)COP(=O)(O)OP(=O)(O)OCC1OC(n2cnc3c(N)ncnc32)C(O)C1OP(=O)(O)O"
}
},
{
"id":"cpd00388_C",
"name":"Allantoate",
"compartment":"C",
"annotation":{
"smiles":"NC(=O)NC(NC(N)=O)C(=O)[O-]",
"smiles_sanitized":"NC(=O)NC(NC(N)=O)C(=O)O"
}
},
{
"id":"cpd00383_P",
"name":"Shikimate",
"compartment":"P",
"annotation":{
"smiles":"O=C([O-])C1=C[C@@H](O)[C@@H](O)[C@H](O)C1",
"smiles_sanitized":"O=C(O)C1=CC(O)C(O)C(O)C1"
}
},
{
"id":"cpd00363_C",
"name":"Ethanol",
"compartment":"C",
"annotation":{
"smiles":"CCO",
"smiles_sanitized":"CCO"
}
},
{
"id":"cpd00069_E",
"name":"L-Tyrosine",
"compartment":"E",
"annotation":{
"smiles":"[NH3+][C@@H](Cc1ccc(O)cc1)C(=O)[O-]",
"smiles_sanitized":"NC(Cc1ccc(O)cc1)C(=O)O"
}
},
{
"id":"cpd00214_E",
"name":"Palmitate",
"compartment":"E",
"annotation":{
"smiles":"CCCCCCCCCCCCCCCC(=O)[O-]",
"smiles_sanitized":"CCCCCCCCCCCCCCCC(=O)O"
}
},
{
"id":"cpd00106_C",
"name":"Fumarate",
"compartment":"C",
"annotation":{
"smiles":"O=C([O-])/C=C/C(=O)[O-]",
"smiles_sanitized":"O=C(O)C=CC(=O)O"
}
},
{
"id":"cpd00133_C",
"name":"Nicotinamide",
"compartment":"C",
"annotation":{
"smiles":"NC(=O)c1cccnc1",
"smiles_sanitized":"NC(=O)c1cccnc1"
}
},
{
"id":"cpd00077_C",
"name":"Catechol",
"compartment":"C",
"annotation":{
"smiles":"Oc1ccccc1O",
"smiles_sanitized":"Oc1ccccc1O"
}
},
{
"id":"cpd01947_E",
"name":"BDOH",
"compartment":"E",
"annotation":{
"smiles":"C[C@@H](O)[C@@H](C)O",
"smiles_sanitized":"CC(O)C(C)O"
}
},
{
"id":"cpd02030_C",
"name":"3-phosphoshikimate",
"compartment":"C",
"annotation":{
"smiles":"O=C([O-])C1=C[C@@H](OP(=O)([O-])[O-])[C@@H](O)[C@H](O)C1",
"smiles_sanitized":"O=C(O)C1=CC(OP(=O)(O)O)C(O)C(O)C1"
}
},
{
"id":"ACYL_COA_WAX_C",
"name":"ACYL-COA_WAX",
"compartment":"C"
},
{
"id":"cpd00571_C",
"name":"Glucarate",
"compartment":"C",
"annotation":{
"smiles":"O=C([O-])[C@@H](O)[C@@H](O)[C@H](O)[C@@H](O)C(=O)[O-]",
"smiles_sanitized":"O=C(O)C(O)C(O)C(O)C(O)C(=O)O"
}
},
{
"id":"cpd00196_C",
"name":"Acetylphosphate",
"compartment":"C",
"annotation":{
"smiles":"CC(=O)OP(=O)([O-])[O-]",
"smiles_sanitized":"CC(=O)OP(=O)(O)O"
}
},
{
"id":"cpd01873_E",
"name":"2-Aminobutan-4-olide",
"compartment":"E",
"annotation":{
"smiles":"[NH3+]C1CCOC1=O",
"smiles_sanitized":"NC1CCOC1=O"
}
},
{
"id":"cpd16613_C",
"name":"cis-3-hydroxytetradec-4-enoyl-coa",
"compartment":"C",
"annotation":{
"smiles":"",
"smiles_sanitized":""
}
},
{
"id":"cpd00358_C",
"name":"dUTP",
"compartment":"C",
"annotation":{
"smiles":"O=c1ccn([C@H]2C[C@H](O)[C@@H](COP(=O)([O-])OP(=O)([O-])OP(=O)([O-])O)O2)c(=O)[nH]1",
"smiles_sanitized":"O=c1ccn(C2CC(O)C(COP(=O)(O)OP(=O)(O)OP(=O)(O)O)O2)c(=O)[nH]1"
}
},
{
"id":"cpd00525_C",
"name":"UDP-N-acetylmuramoyl-L-alanyl-D-glutamate",
"compartment":"C",
"annotation":{
"smiles":"CC(=O)N[C@H]1[C@@H](OP(=O)([O-])OP(=O)([O-])OC[C@H]2O[C@@H](n3ccc(=O)[nH]c3=O)[C@H](O)[C@@H]2O)O[C@H](CO)[C@@H](O)[C@@H]1O[C@H](C)C(=O)N[C@@H](C)C(=O)N[C@H](CCC(=O)[O-])C(=O)[O-]",
"smiles_sanitized":"CC(=O)NC1C(OP(=O)(O)OP(=O)(O)OCC2OC(n3ccc(=O)[nH]c3=O)C(O)C2O)OC(CO)C(O)C1OC(C)C(=O)NC(C)C(=O)NC(CCC(=O)O)C(=O)O"
}
},
{
"id":"cpd00065_E",
"name":"L-Tryptophan",
"compartment":"E",
"annotation":{
"smiles":"[NH3+][C@@H](Cc1c[nH]c2ccccc12)C(=O)[O-]",
"smiles_sanitized":"NC(Cc1c[nH]c2ccccc12)C(=O)O"
}
},
{
"id":"cpd02535_C",
"name":"2,3-Dihydroxy-3-methylvalerate",
"compartment":"C",
"annotation":{
"smiles":"CC[C@@](C)(O)[C@@H](O)C(=O)[O-]",
"smiles_sanitized":"CCC(C)(O)C(O)C(=O)O"
}
},
{
"id":"cpd00288_C",
"name":"D-Glucosamine phosphate",
"compartment":"C",
"annotation":{
"smiles":"[NH3+][C@H]1C(O)O[C@H](COP(=O)([O-])[O-])[C@@H](O)[C@@H]1O",
"smiles_sanitized":"NC1C(O)OC(COP(=O)(O)O)C(O)C1O"
}
},
{
"id":"cpd00266_C",
"name":"Carnitine",
"compartment":"C",
"annotation":{
"smiles":"C[N+](C)(C)C[C@H](O)CC(=O)[O-]",
"smiles_sanitized":"C[N+](C)(C)CC(O)CC(=O)O"
}
},
{
"id":"cpd08210_C",
"name":"ADC",
"compartment":"C",
"annotation":{
"smiles":"C=C(O[C@@H]1C=C(C(=O)[O-])C=C[C@H]1[NH3+])C(=O)[O-]",
"smiles_sanitized":"C=C(OC1C=C(C(=O)O)C=CC1N)C(=O)O"
}
},
{
"id":"cpd04103_C",
"name":"3-Hydroxypimelyl-CoA",
"compartment":"C",
"annotation":{
"smiles":"CC(C)(COP(=O)([O-])OP(=O)([O-])OC[C@H]1O[C@@H](n2cnc3c(N)ncnc32)[C@H](O)[C@@H]1OP(=O)([O-])[O-])[C@@H](O)C(=O)NCCC(=O)NCCSC(=O)CC(O)CCCC(=O)[O-]",
"smiles_sanitized":"CC(C)(COP(=O)(O)OP(=O)(O)OCC1OC(n2cnc3c(N)ncnc32)C(O)C1OP(=O)(O)O)C(O)C(=O)NCCC(=O)NCCSC(=O)CC(O)CCCC(=O)O"
}
},
{
"id":"cpd03113_C",
"name":"(S)-3-Hydroxyhexadecanoyl-CoA",
"compartment":"C",
"annotation":{
"smiles":"CCCCCCCCCCCCC[C@H](O)CC(=O)SCCNC(=O)CCNC(=O)C(O)C(C)(C)COP(=O)([O-])OP(=O)([O-])OC[C@H]1O[C@@H](n2cnc3c(N)ncnc32)[C@H](O)[C@@H]1OP(=O)([O-])[O-]",
"smiles_sanitized":"CCCCCCCCCCCCCC(O)CC(=O)SCCNC(=O)CCNC(=O)C(O)C(C)(C)COP(=O)(O)OP(=O)(O)OCC1OC(n2cnc3c(N)ncnc32)C(O)C1OP(=O)(O)O"
}
},
{
"id":"cpd00210_C",
"name":"Taurine",
"compartment":"C",
"annotation":{
"smiles":"[NH3+]CCS(=O)(=O)[O-]",
"smiles_sanitized":"NCCS(=O)(=O)O"
}
},
{
"id":"cpd16654_E",
"name":"tricarballylate",
"compartment":"E",
"annotation":{
"smiles":"O=C([O-])CC(CC(=O)[O-])C(=O)[O-]",
"smiles_sanitized":"O=C(O)CC(CC(=O)O)C(=O)O"
}
},
{
"id":"cpd04111_C",
"name":"6-Carboxyhex-2-enoyl-CoA",
"compartment":"C",
"annotation":{
"smiles":"CC(C)(COP(=O)([O-])OP(=O)([O-])OC[C@H]1O[C@@H](n2cnc3c(N)ncnc32)[C@H](O)[C@@H]1OP(=O)([O-])[O-])[C@@H](O)C(=O)NCCC(=O)NCCSC(=O)/C=C/CCCC(=O)[O-]",
"smiles_sanitized":"CC(C)(COP(=O)(O)OP(=O)(O)OCC1OC(n2cnc3c(N)ncnc32)C(O)C1OP(=O)(O)O)C(O)C(=O)NCCC(=O)NCCSC(=O)C=CCCCC(=O)O"
}
},
{
"id":"cpd00945_C",
"name":"2-Dehydro-3-deoxy-D-galactonate 6-phosphate",
"compartment":"C",
"annotation":{
"smiles":"O=C([O-])C(=O)C[C@@H](O)[C@H](O)COP(=O)([O-])[O-]",
"smiles_sanitized":"O=C(O)C(=O)CC(O)C(O)COP(=O)(O)O"
}
},
{
"id":"C6_0_ACP_C",
"name":"hexanoyl-acp",
"compartment":"C"
},
{
"id":"C16_1_C7__ACP_C",
"name":"cis-hexadec-7-enoyl-acp",
"compartment":"C"
},
{
"id":"cpd11225_C",
"name":"3-4-dihydroxy-2-butanone4-phosphate",
"compartment":"C",
"annotation":{
"smiles":"CC(=O)[C@H](O)COP(=O)([O-])[O-]",
"smiles_sanitized":"CC(=O)C(O)COP(=O)(O)O"
}
},
{
"id":"cpd00281_C",
"name":"GABA",
"compartment":"C",
"annotation":{
"smiles":"[NH3+]CCCC(=O)[O-]",
"smiles_sanitized":"NCCCC(=O)O"
}
},
{
"id":"cpd00307_E",
"name":"Cytosine",
"compartment":"E",
"annotation":{
"smiles":"Nc1cc[nH]c(=O)n1",
"smiles_sanitized":"Nc1cc[nH]c(=O)n1"
}
},
{
"id":"cpd00062_C",
"name":"UTP",
"compartment":"C",
"annotation":{
"smiles":"O=c1ccn([C@@H]2O[C@H](COP(=O)([O-])OP(=O)([O-])OP(=O)([O-])O)[C@@H](O)[C@H]2O)c(=O)[nH]1",
"smiles_sanitized":"O=c1ccn(C2OC(COP(=O)(O)OP(=O)(O)OP(=O)(O)O)C(O)C2O)c(=O)[nH]1"
}
},
{
"id":"cpd00130_C",
"name":"L-Malate",
"compartment":"C",
"annotation":{
"smiles":"O=C([O-])C[C@H](O)C(=O)[O-]",
"smiles_sanitized":"O=C(O)CC(O)C(=O)O"
}
},
{
"id":"cpd16615_C",
"name":"cis-3-oxotetradec-4-enoyl-coa",
"compartment":"C",
"annotation":{
"smiles":"",
"smiles_sanitized":""
}
},
{
"id":"cpd00113_C",
"name":"Isopentenyldiphosphate",
"compartment":"C",
"annotation":{
"smiles":"C=C(C)CCOP(=O)([O-])OP(=O)([O-])O",
"smiles_sanitized":"C=C(C)CCOP(=O)(O)OP(=O)(O)O"
}
},
{
"id":"cpd03121_C",
"name":"3-Oxooctanoyl-CoA",
"compartment":"C",
"annotation":{
"smiles":"CCCCCC(=O)CC(=O)SCCNC(=O)CCNC(=O)[C@H](O)C(C)(C)COP(=O)([O-])OP(=O)([O-])OC[C@H]1O[C@@H](n2cnc3c(N)ncnc32)[C@H](O)[C@@H]1OP(=O)([O-])[O-]",
"smiles_sanitized":"CCCCCC(=O)CC(=O)SCCNC(=O)CCNC(=O)C(O)C(C)(C)COP(=O)(O)OP(=O)(O)OCC1OC(n2cnc3c(N)ncnc32)C(O)C1OP(=O)(O)O"
}
},
{
"id":"cpd02979_C",
"name":"phosphoribosylformiminoaicar-phosphate",
"compartment":"C",
"annotation":{
"smiles":"NC(=O)c1ncn([C@@H]2O[C@H](COP(=O)([O-])[O-])[C@@H](O)[C@H]2O)c1/N=C/N[C@@H]1O[C@H](COP(=O)([O-])[O-])[C@@H](O)[C@H]1O",
"smiles_sanitized":"NC(=O)c1ncn(C2OC(COP(=O)(O)O)C(O)C2O)c1N=CNC1OC(COP(=O)(O)O)C(O)C1O"
}
},
{
"id":"cpd12801_C",
"name":"Cardiolipin",
"compartment":"C",
"annotation":{
"smiles":"*C(=O)OC[C@H](COP(=O)([O-])OCC(O)COP(=O)([O-])OC[C@@H](COC(*)=O)OC(*)=O)OC(*)=O",
"smiles_sanitized":"*C(=O)OCC(COP(=O)(O)OCC(O)COP(=O)(O)OCC(COC(*)=O)OC(*)=O)OC(*)=O"
}
},
{
"id":"cpd00073_C",
"name":"Urea",
"compartment":"C",
"annotation":{
"smiles":"NC(N)=O",
"smiles_sanitized":"NC(N)=O"
}
},
{
"id":"LPS_W_C",
"name":"LPS_W",
"compartment":"C"
},
{
"id":"cpd00168_C",
"name":"2,3-Dihydroxybenzoate",
"compartment":"C",
"annotation":{
"smiles":"O=C([O-])c1cccc(O)c1O",
"smiles_sanitized":"O=C(O)c1cccc(O)c1O"
}
},
{
"id":"cpd27812_C",
"name":"propanamide",
"compartment":"C",
"annotation":{
"smiles":"CCC(N)=O",
"smiles_sanitized":"CCC(N)=O"
}
},
{
"id":"cpd03124_C",
"name":"Hexanoyl-CoA",
"compartment":"C",
"annotation":{
"smiles":"CCCCCC(=O)SCCNC(=O)CCNC(=O)[C@H](O)C(C)(C)COP(=O)([O-])OP(=O)([O-])OC[C@H]1O[C@@H](n2cnc3c(N)ncnc32)[C@H](O)[C@@H]1OP(=O)([O-])[O-]",
"smiles_sanitized":"CCCCCC(=O)SCCNC(=O)CCNC(=O)C(O)C(C)(C)COP(=O)(O)OP(=O)(O)OCC1OC(n2cnc3c(N)ncnc32)C(O)C1OP(=O)(O)O"
}
},
{
"id":"cpd00881_C",
"name":"Caffeate",
"compartment":"C",
"annotation":{
"smiles":"O=C([O-])/C=C/c1ccc(O)c(O)c1",
"smiles_sanitized":"O=C(O)C=Cc1ccc(O)c(O)c1"
}
},
{
"id":"C18_1_C9__3OXOACP_C",
"name":"cis-3-oxooctadec-9-enoyl-acp",
"compartment":"C"
},
{
"id":"cpd02227_C",
"name":"S-Ribosylhomocysteine",
"compartment":"C",
"annotation":{
"smiles":"[NH3+][C@@H](CCSC[C@H]1O[C@@H](O)[C@H](O)[C@@H]1O)C(=O)[O-]",
"smiles_sanitized":"NC(CCSCC1OC(O)C(O)C1O)C(=O)O"
}
},
{
"id":"cpd08301_C",
"name":"2-C-methyl-D-erythritol2-4-cyclodiphosphate",
"compartment":"C",
"annotation":{
"smiles":"C[C@@]1(CO)OP(=O)([O-])OP(=O)([O-])OC[C@H]1O",
"smiles_sanitized":"CC1(CO)OP(=O)(O)OP(=O)(O)OCC1O"
}
},
{
"id":"cpd01977_C",
"name":"4-Phospho-L-aspartate",
"compartment":"C",
"annotation":{
"smiles":"[NH3+][C@@H](CC(=O)OP(=O)([O-])[O-])C(=O)[O-]",
"smiles_sanitized":"NC(CC(=O)OP(=O)(O)O)C(=O)O"
}
},
{
"id":"cpd00002_C",
"name":"ATP",
"compartment":"C",
"annotation":{
"smiles":"Nc1ncnc2c1ncn2[C@@H]1O[C@H](COP(=O)([O-])OP(=O)([O-])OP(=O)([O-])O)[C@@H](O)[C@H]1O",
"smiles_sanitized":"Nc1ncnc2c1ncn2C1OC(COP(=O)(O)OP(=O)(O)OP(=O)(O)O)C(O)C1O"
}
},
{
"id":"cpd00184_C",
"name":"Thymidine",
"compartment":"C",
"annotation":{
"smiles":"Cc1cn([C@H]2C[C@H](O)[C@@H](CO)O2)c(=O)[nH]c1=O",
"smiles_sanitized":"Cc1cn(C2CC(O)C(CO)O2)c(=O)[nH]c1=O"
}
},
{
"id":"cpd01716_C",
"name":"3-Dehydroshikimate",
"compartment":"C",
"annotation":{
"smiles":"O=C([O-])C1=CC(=O)[C@@H](O)[C@H](O)C1",
"smiles_sanitized":"O=C(O)C1=CC(=O)C(O)C(O)C1"
}
},
{
"id":"cpd00597_C",
"name":"Valerate",
"compartment":"C",
"annotation":{
"smiles":"CCCCC(=O)[O-]",
"smiles_sanitized":"CCCCC(=O)O"
}
},
{
"id":"cpd00067_P",
"name":"H+",
"compartment":"P",
"annotation":{
"smiles":"[H+]",
"smiles_sanitized":"[H+]"
}
},
{
"id":"cpd02120_C",
"name":"Dihydrodipicolinate",
"compartment":"C",
"annotation":{
"smiles":"O=C([O-])C1=N[C@H](C(=O)[O-])CC=C1",
"smiles_sanitized":"O=C(O)C1=NC(C(=O)O)CC=C1"
}
},
{
"id":"cpd03116_C",
"name":"(S)-3-Hydroxydodecanoyl-CoA",
"compartment":"C",
"annotation":{
"smiles":"CCCCCCCCC[C@H](O)CC(=O)SCCNC(=O)CCNC(=O)[C@H](O)C(C)(C)COP(=O)([O-])OP(=O)([O-])OC[C@H]1O[C@@H](n2cnc3c(N)ncnc32)[C@H](O)[C@@H]1OP(=O)([O-])[O-]",
"smiles_sanitized":"CCCCCCCCCC(O)CC(=O)SCCNC(=O)CCNC(=O)C(O)C(C)(C)COP(=O)(O)OP(=O)(O)OCC1OC(n2cnc3c(N)ncnc32)C(O)C1OP(=O)(O)O"
}
},
{
"id":"cpd01270_C",
"name":"FMNH2",
"compartment":"C",
"annotation":{
"smiles":"Cc1cc2c(cc1C)N(C[C@H](O)[C@H](O)[C@H](O)COP(=O)([O-])[O-])c1[nH]c(=O)[nH]c(=O)c1N2",
"smiles_sanitized":"Cc1cc2c(cc1C)N(CC(O)C(O)C(O)COP(=O)(O)O)c1[nH]c(=O)[nH]c(=O)c1N2"
}
},
{
"id":"cpd00054_C",
"name":"L-Serine",
"compartment":"C",
"annotation":{
"smiles":"[NH3+][C@@H](CO)C(=O)[O-]",
"smiles_sanitized":"NC(CO)C(=O)O"
}
},
{
"id":"C16_0_3OXOACP_C",
"name":"3-oxohexadecanoyl-acp",
"compartment":"C"
},
{
"id":"cpd02724_C",
"name":"N-Succinyl-L-2-amino-6-oxopimelate",
"compartment":"C",
"annotation":{
"smiles":"O=C([O-])CCC(=O)N[C@@H](CCCC(=O)C(=O)[O-])C(=O)[O-]",
"smiles_sanitized":"O=C(O)CCC(=O)NC(CCCC(=O)C(=O)O)C(=O)O"
}
},
{
"id":"cpd00222_P",
"name":"GLCN",
"compartment":"P",
"annotation":{
"smiles":"O=C([O-])[C@H](O)[C@@H](O)[C@H](O)[C@H](O)CO",
"smiles_sanitized":"O=C(O)C(O)C(O)C(O)C(O)CO"
}
},
{
"id":"CHARGED_TYR_TRNAS_C",
"name":"charged tyrosine trna",
"compartment":"C"
},
{
"id":"cpd00117_C",
"name":"D-Alanine",
"compartment":"C",
"annotation":{
"smiles":"C[C@@H]([NH3+])C(=O)[O-]",
"smiles_sanitized":"CC(N)C(=O)O"
}
},
{
"id":"cpd00599_E",
"name":"SALC",
"compartment":"E",
"annotation":{
"smiles":"O=C([O-])c1ccccc1O",
"smiles_sanitized":"O=C(O)c1ccccc1O"
}
},
{
"id":"cpd00516_C",
"name":"meso-2,6-Diaminopimelate",
"compartment":"C",
"annotation":{
"smiles":"[NH3+][C@@H](CCC[C@@H]([NH3+])C(=O)[O-])C(=O)[O-]",
"smiles_sanitized":"NC(CCCC(N)C(=O)O)C(=O)O"
}
},
{
"id":"cpd02961_C",
"name":"Dihydroneopterin",
"compartment":"C",
"annotation":{
"smiles":"Nc1nc2c(c(=O)[nH]1)N=C([C@H](O)[C@H](O)CO)CN2",
"smiles_sanitized":"Nc1nc2c(c(=O)[nH]1)N=C(C(O)C(O)CO)CN2"
}
},
{
"id":"cpd00132_C",
"name":"L-Asparagine",
"compartment":"C",
"annotation":{
"smiles":"NC(=O)C[C@H]([NH3+])C(=O)[O-]",
"smiles_sanitized":"NC(=O)CC(N)C(=O)O"
}
},
{
"id":"cpd16617_C",
"name":"cis-tetradec-4-enoyl-coa",
"compartment":"C",
"annotation":{
"smiles":"",
"smiles_sanitized":""
}
},
{
"id":"cpd00127_C",
"name":"Phenol",
"compartment":"C",
"annotation":{
"smiles":"Oc1ccccc1",
"smiles_sanitized":"Oc1ccccc1"
}
},
{
"id":"cpd02730_C",
"name":"3-Deoxy-D-manno-octulosonate8-phosphate",
"compartment":"C",
"annotation":{
"smiles":"O=C([O-])[C@@]1(O)C[C@@H](O)[C@@H](O)[C@@H]([C@H](O)COP(=O)([O-])[O-])O1",
"smiles_sanitized":"O=C(O)C1(O)CC(O)C(O)C(C(O)COP(=O)(O)O)O1"
}
},
{
"id":"cpd00726_C",
"name":"1,3-Propanediamine",
"compartment":"C",
"annotation":{
"smiles":"[NH3+]CCC[NH3+]",
"smiles_sanitized":"NCCCN"
}
},
{
"id":"cpd00041_E",
"name":"L-Aspartate",
"compartment":"E",
"annotation":{
"smiles":"[NH3+][C@@H](CC(=O)[O-])C(=O)[O-]",
"smiles_sanitized":"NC(CC(=O)O)C(=O)O"
}
},
{
"id":"cpd00630_C",
"name":"3-Oxoadipate",
"compartment":"C",
"annotation":{
"smiles":"O=C([O-])CCC(=O)CC(=O)[O-]",
"smiles_sanitized":"O=C(O)CCC(=O)CC(=O)O"
}
},
{
"id":"cpd03525_C",
"name":"N-Succinyl-L-glutamate",
"compartment":"C",
"annotation":{
"smiles":"O=C([O-])CCC(=O)N[C@@H](CCC(=O)[O-])C(=O)[O-]",
"smiles_sanitized":"O=C(O)CCC(=O)NC(CCC(=O)O)C(=O)O"
}
},
{
"id":"cpd00038_C",
"name":"GTP",
"compartment":"C",
"annotation":{
"smiles":"Nc1nc2c(ncn2[C@@H]2O[C@H](COP(=O)([O-])OP(=O)([O-])OP(=O)([O-])O)[C@@H](O)[C@H]2O)c(=O)[nH]1",
"smiles_sanitized":"Nc1nc2c(ncn2C2OC(COP(=O)(O)OP(=O)(O)OP(=O)(O)O)C(O)C2O)c(=O)[nH]1"
}
},
{
"id":"cpd00114_C",
"name":"IMP",
"compartment":"C",
"annotation":{
"smiles":"O=c1[nH]cnc2c1ncn2[C@@H]1O[C@H](COP(=O)([O-])[O-])[C@@H](O)[C@H]1O",
"smiles_sanitized":"O=c1[nH]cnc2c1ncn2C1OC(COP(=O)(O)O)C(O)C1O"
}
},
{
"id":"cpd01777_C",
"name":"Phosphoribosyl-AMP",
"compartment":"C",
"annotation":{
"smiles":"N=c1c2ncn(C3O[C@H](COP(=O)([O-])[O-])[C@@H](O)[C@H]3O)c2ncn1[C@@H]1O[C@H](COP(=O)([O-])[O-])[C@@H](O)[C@H]1O",
"smiles_sanitized":"N=c1c2ncn(C3OC(COP(=O)(O)O)C(O)C3O)c2ncn1C1OC(COP(=O)(O)O)C(O)C1O"
}
},
{
"id":"C6_0_2ENOYLACP_C",
"name":"trans-hex-2-enoyl-acp",
"compartment":"C"
},
{
"id":"cpd00227_C",
"name":"L-Homoserine",
"compartment":"C",
"annotation":{
"smiles":"[NH3+][C@@H](CCO)C(=O)[O-]",
"smiles_sanitized":"NC(CCO)C(=O)O"
}
},
{
"id":"cpd00142_C",
"name":"Acetoacetate",
"compartment":"C",
"annotation":{
"smiles":"CC(=O)CC(=O)[O-]",
"smiles_sanitized":"CC(=O)CC(=O)O"
}
},
{
"id":"cpd00954_C",
"name":"6-hydroxymethyl dihydropterin",
"compartment":"C",
"annotation":{
"smiles":"Nc1nc(O)c2c(n1)NCC(CO)=N2",
"smiles_sanitized":"Nc1nc(O)c2c(n1)NCC(CO)=N2"
}
},
{
"id":"cpd00027_P",
"name":"D-Glucose",
"compartment":"P",
"annotation":{
"smiles":"OC[C@H]1OC(O)[C@H](O)[C@@H](O)[C@@H]1O",
"smiles_sanitized":"OCC1OC(O)C(O)C(O)C1O"
}
},
{
"id":"cpd01733_C",
"name":"Ferulaldehyde",
"compartment":"C",
"annotation":{
"smiles":"COc1cc(/C=C/C=O)ccc1O",
"smiles_sanitized":"COc1cc(C=CC=O)ccc1O"
}
},
{
"id":"cpd00024_E",
"name":"2-Oxoglutarate",
"compartment":"E",
"annotation":{
"smiles":"O=C([O-])CCC(=O)C(=O)[O-]",
"smiles_sanitized":"O=C(O)CCC(=O)C(=O)O"
}
},
{
"id":"cpd04122_C",
"name":"Aminoacetaldehyde",
"compartment":"C",
"annotation":{
"smiles":"[NH3+]CC=O",
"smiles_sanitized":"NCC=O"
}
},
{
"id":"cpd00078_C",
"name":"Succinyl-CoA",
"compartment":"C",
"annotation":{
"smiles":"CC(C)(COP(=O)([O-])OP(=O)([O-])OC[C@H]1O[C@@H](n2cnc3c(N)ncnc32)[C@H](O)[C@@H]1OP(=O)([O-])[O-])[C@@H](O)C(=O)NCCC(=O)NCCSC(=O)CCC(=O)[O-]",
"smiles_sanitized":"CC(C)(COP(=O)(O)OP(=O)(O)OCC1OC(n2cnc3c(N)ncnc32)C(O)C1OP(=O)(O)O)C(O)C(=O)NCCC(=O)NCCSC(=O)CCC(=O)O"
}
},
{
"id":"cpd02656_C",
"name":"6-7-Dimethyl-8--1-D-ribityllumazine",
"compartment":"C",
"annotation":{
"smiles":"Cc1nc2c(=O)[n-]c(=O)nc-2n(C[C@H](O)[C@H](O)[C@H](O)CO)c1C",
"smiles_sanitized":"Cc1nc2c(=O)[nH]c(=O)nc-2n(CC(O)C(O)C(O)CO)c1C"
}
},
{
"id":"cpd00159_C",
"name":"L-Lactate",
"compartment":"C",
"annotation":{
"smiles":"C[C@H](O)C(=O)[O-]",
"smiles_sanitized":"CC(O)C(=O)O"
}
},
{
"id":"cpd00282_C",
"name":"S-Dihydroorotate",
"compartment":"C",
"annotation":{
"smiles":"O=C1C[C@@H](C(=O)[O-])NC(=O)N1",
"smiles_sanitized":"O=C1CC(C(=O)O)NC(=O)N1"
}
},
{
"id":"cpd25799_C",
"name":"MK-4",
"compartment":"C",
"annotation":{
"smiles":"CC(C)=CCC/C(C)=C/CC/C(C)=C/CC/C(C)=C/CC1=C(C)C(=O)c2ccccc2C1=O",
"smiles_sanitized":"CC(C)=CCCC(C)=CCCC(C)=CCCC(C)=CCC1=C(C)C(=O)c2ccccc2C1=O"
}
},
{
"id":"cpd01775_C",
"name":"Phosphoribosyl-ATP",
"compartment":"C",
"annotation":{
"smiles":"N=c1c2ncn(C3O[C@H](COP(=O)([O-])OP(=O)([O-])OP(=O)([O-])O)[C@@H](O)[C@H]3O)c2ncn1[C@@H]1O[C@H](COP(=O)([O-])[O-])[C@@H](O)[C@H]1O",
"smiles_sanitized":"N=c1c2ncn(C3OC(COP(=O)(O)OP(=O)(O)OP(=O)(O)O)C(O)C3O)c2ncn1C1OC(COP(=O)(O)O)C(O)C1O"
}
},
{
"id":"cpd02229_C",
"name":"Bactoprenyl diphosphate",
"compartment":"C",
"annotation":{
"smiles":"CC(C)=CCC/C(C)=C/CC/C(C)=C/CC/C(C)=C\\CC/C(C)=C\\CC/C(C)=C\\CC/C(C)=C\\CC/C(C)=C\\CC/C(C)=C\\CC/C(C)=C\\CC/C(C)=C\\COP(=O)([O-])OP(=O)([O-])O",
"smiles_sanitized":"CC(C)=CCCC(C)=CCCC(C)=CCCC(C)=CCCC(C)=CCCC(C)=CCCC(C)=CCCC(C)=CCCC(C)=CCCC(C)=CCCC(C)=CCOP(=O)(O)OP(=O)(O)O"
}
},
{
"id":"C4_0_3HYDROXYACP_C",
"name":"3-hydroxybutanoyl-acp",
"compartment":"C"
},
{
"id":"cpd02893_C",
"name":"5'-Phosphoribosyl-4-carboxy-5-aminoimidazole",
"compartment":"C",
"annotation":{
"smiles":"Nc1c(C(=O)[O-])[nH+]cn1[C@@H]1O[C@H](COP(=O)([O-])[O-])[C@@H](O)[C@H]1O",
"smiles_sanitized":"Nc1c(C(=O)O)ncn1C1OC(COP(=O)(O)O)C(O)C1O"
}
},
{
"id":"cpd00408_C",
"name":"Pantoate",
"compartment":"C",
"annotation":{
"smiles":"CC(C)(CO)[C@@H](O)C(=O)[O-]",
"smiles_sanitized":"CC(C)(CO)C(O)C(=O)O"
}
},
{
"id":"cpd02693_C",
"name":"3-Isopropylmalate",
"compartment":"C",
"annotation":{
"smiles":"CC(C)[C@H](C(=O)[O-])[C@@H](O)C(=O)[O-]",
"smiles_sanitized":"CC(C)C(C(=O)O)C(O)C(=O)O"
}
},
{
"id":"cpd02835_C",
"name":"UDP-2,3-bis(3-hydroxytetradecanoyl)glucosamine",
"compartment":"C",
"annotation":{
"smiles":"CCCCCCCCCCC[C@@H](O)CC(=O)N[C@H]1[C@@H](OP(=O)([O-])OP(=O)([O-])OC[C@H]2O[C@@H](n3ccc(=O)[nH]c3=O)[C@H](O)[C@@H]2O)O[C@H](CO)[C@@H](O)[C@@H]1OC(=O)C[C@H](O)CCCCCCCCCCC",
"smiles_sanitized":"CCCCCCCCCCCC(O)CC(=O)NC1C(OP(=O)(O)OP(=O)(O)OCC2OC(n3ccc(=O)[nH]c3=O)C(O)C2O)OC(CO)C(O)C1OC(=O)CC(O)CCCCCCCCCCC"
}
},
{
"id":"C14_0_3OXOACP_C",
"name":"3-oxotetradecanoyl-acp",
"compartment":"C"
},
{
"id":"CHARGED_PRO_TRNAS_C",
"name":"charged proline trna",
"compartment":"C"
},
{
"id":"cpd00183_C",
"name":"Sarcosine",
"compartment":"C",
"annotation":{
"smiles":"C[NH2+]CC(=O)[O-]",
"smiles_sanitized":"CNCC(=O)O"
}
},
{
"id":"cpd00956_C",
"name":"1-(2-carboxyphenylamino)-1-deoxyribulose 5-phosphate",
"compartment":"C",
"annotation":{
"smiles":"O=C([O-])c1ccccc1NCC(=O)[C@H](O)[C@H](O)COP(=O)([O-])[O-]",
"smiles_sanitized":"O=C(O)c1ccccc1NCC(=O)C(O)C(O)COP(=O)(O)O"
}
},
{
"id":"CHARGED_ASN_TRNAS_C",
"name":"charged asparagine trna",
"compartment":"C"
},
{
"id":"ACYL_ACP_PL_C",
"name":"ACYL-ACP_PL",
"compartment":"C"
},
{
"id":"cpd01873_C",
"name":"2-Aminobutan-4-olide",
"compartment":"C",
"annotation":{
"smiles":"[NH3+]C1CCOC1=O",
"smiles_sanitized":"NC1CCOC1=O"
}
},
{
"id":"cpd00311_E",
"name":"Guanosine",
"compartment":"E",
"annotation":{
"smiles":"Nc1nc2c(ncn2[C@@H]2O[C@H](CO)[C@@H](O)[C@H]2O)c(=O)[nH]1",
"smiles_sanitized":"Nc1nc2c(ncn2C2OC(CO)C(O)C2O)c(=O)[nH]1"
}
},
{
"id":"C16_0_2ENOYLACP_C",
"name":"trans-hexadec-2-enoyl-acp",
"compartment":"C"
},
{
"id":"cpd00084_C",
"name":"L-Cysteine",
"compartment":"C",
"annotation":{
"smiles":"[NH3+][C@@H](CS)C(=O)[O-]",
"smiles_sanitized":"NC(CS)C(=O)O"
}
},
{
"id":"cpd00930_C",
"name":"imidazole acetol-phosphate",
"compartment":"C",
"annotation":{
"smiles":"O=C(COP(=O)([O-])[O-])Cc1c[nH]cn1",
"smiles_sanitized":"O=C(COP(=O)(O)O)Cc1c[nH]cn1"
}
},
{
"id":"cpd00347_C",
"name":"5-10-Methenyltetrahydrofolate",
"compartment":"C",
"annotation":{
"smiles":"Nc1nc2c(c(=O)[nH]1)[N+]1=CN(c3ccc(C(=O)N[C@@H](CCC(=O)[O-])C(=O)[O-])cc3)C[C@H]1CN2",
"smiles_sanitized":"Nc1nc2c(c(=O)[nH]1)[N+]1=CN(c3ccc(C(=O)NC(CCC(=O)O)C(=O)O)cc3)CC1CN2"
}
},
{
"id":"cpd00557_C",
"name":"Siroheme",
"compartment":"C",
"annotation":{
"smiles":"C[C@@]1(CC(=O)[O-])/C2=C/c3c(CC(=O)[O-])c(CCC(=O)[O-])c4n3[Fe]N3/C(=C\\C5=N/C(=C\\4)C(CCC(=O)[O-])=C5CC(=O)[O-])[C@@H](CCC(=O)[O-])[C@](C)(CC(=O)[O-])/C3=C/C(=N2)[C@H]1CCC(=O)[O-]",
"smiles_sanitized":"CC1(CC(=O)O)C2=Cc3c(CC(=O)O)c(CCC(=O)O)c4n3[Fe]N3C(=CC5=NC(=C4)C(CCC(=O)O)=C5CC(=O)O)C(CCC(=O)O)C(C)(CC(=O)O)C3=CC(=N2)C1CCC(=O)O"
}
},
{
"id":"cpd16609_C",
"name":"cis-tetradec-4-enoate",
"compartment":"C",
"annotation":{
"smiles":"",
"smiles_sanitized":""
}
},
{
"id":"cpd00722_C",
"name":"O-Acetyl-L-serine",
"compartment":"C",
"annotation":{
"smiles":"CC(=O)OC[C@H]([NH3+])C(=O)[O-]",
"smiles_sanitized":"CC(=O)OCC(N)C(=O)O"
}
},
{
"id":"cpd00132_E",
"name":"L-Asparagine",
"compartment":"E",
"annotation":{
"smiles":"NC(=O)C[C@H]([NH3+])C(=O)[O-]",
"smiles_sanitized":"NC(=O)CC(N)C(=O)O"
}
},
{
"id":"cpd00477_C",
"name":"N-Acetyl-L-glutamate",
"compartment":"C",
"annotation":{
"smiles":"CC(=O)N[C@@H](CCC(=O)[O-])C(=O)[O-]",
"smiles_sanitized":"CC(=O)NC(CCC(=O)O)C(=O)O"
}
},
{
"id":"cpd15560_C",
"name":"Ubiquinone-8",
"compartment":"C",
"annotation":{
"smiles":"COC1=C(OC)C(=O)C(C/C=C(\\C)CC/C=C(\\C)CC/C=C(\\C)CC/C=C(\\C)CC/C=C(\\C)CC/C=C(\\C)CC/C=C(\\C)CCC=C(C)C)=C(C)C1=O",
"smiles_sanitized":"COC1=C(OC)C(=O)C(CC=C(C)CCC=C(C)CCC=C(C)CCC=C(C)CCC=C(C)CCC=C(C)CCC=C(C)CCC=C(C)C)=C(C)C1=O"
}
},
{
"id":"C6_1_C4__2ENOYLCOA_C",
"name":"trans,trans-hex-2,4-enoyl-coa",
"compartment":"C"
},
{
"id":"cpd00117_E",
"name":"D-Alanine",
"compartment":"E",
"annotation":{
"smiles":"C[C@@H]([NH3+])C(=O)[O-]",
"smiles_sanitized":"CC(N)C(=O)O"
}
},
{
"id":"cpd01080_E",
"name":"ocdca",
"compartment":"E",
"annotation":{
"smiles":"CCCCCCCCCCCCCCCCCC(=O)[O-]",
"smiles_sanitized":"CCCCCCCCCCCCCCCCCC(=O)O"
}
},
{
"id":"cpd00413_C",
"name":"Glutaryl-CoA",
"compartment":"C",
"annotation":{
"smiles":"CC(C)(COP(=O)([O-])OP(=O)([O-])OC[C@H]1O[C@@H](n2cnc3c(N)ncnc32)[C@H](O)[C@@H]1OP(=O)([O-])[O-])[C@@H](O)C(=O)NCCC(=O)NCCSC(=O)CCCC(=O)[O-]",
"smiles_sanitized":"CC(C)(COP(=O)(O)OP(=O)(O)OCC1OC(n2cnc3c(N)ncnc32)C(O)C1OP(=O)(O)O)C(O)C(=O)NCCC(=O)NCCSC(=O)CCCC(=O)O"
}
},
{
"id":"cpd00046_C",
"name":"CMP",
"compartment":"C",
"annotation":{
"smiles":"Nc1ccn([C@@H]2O[C@H](COP(=O)([O-])[O-])[C@@H](O)[C@H]2O)c(=O)n1",
"smiles_sanitized":"Nc1ccn(C2OC(COP(=O)(O)O)C(O)C2O)c(=O)n1"
}
},
{
"id":"cpd03091_C",
"name":"5'-Deoxyadenosine",
"compartment":"C",
"annotation":{
"smiles":"C[C@H]1O[C@@H](n2cnc3c(N)ncnc32)[C@H](O)[C@@H]1O",
"smiles_sanitized":"CC1OC(n2cnc3c(N)ncnc32)C(O)C1O"
}
},
{
"id":"cpd00508_C",
"name":"3MOP",
"compartment":"C",
"annotation":{
"smiles":"CC[C@H](C)C(=O)C(=O)[O-]",
"smiles_sanitized":"CCC(C)C(=O)C(=O)O"
}
},
{
"id":"cpd02375_C",
"name":"Adenylosuccinate",
"compartment":"C",
"annotation":{
"smiles":"O=C([O-])CC(Nc1ncnc2c1ncn2[C@@H]1O[C@H](COP(=O)([O-])[O-])[C@@H](O)[C@H]1O)C(=O)[O-]",
"smiles_sanitized":"O=C(O)CC(Nc1ncnc2c1ncn2C1OC(COP(=O)(O)O)C(O)C1O)C(=O)O"
}
},
{
"id":"C14_1_C5__ACP_C",
"name":"cis-tetradec-4-enoyl-acp",
"compartment":"C"
},
{
"id":"cpd00350_C",
"name":"Farnesyldiphosphate",
"compartment":"C",
"annotation":{
"smiles":"CC(C)=CCC/C(C)=C/CC/C(C)=C/COP(=O)([O-])OP(=O)([O-])O",
"smiles_sanitized":"CC(C)=CCCC(C)=CCCC(C)=CCOP(=O)(O)OP(=O)(O)O"
}
},
{
"id":"cpd00029_E",
"name":"Acetate",
"compartment":"E",
"annotation":{
"smiles":"CC(=O)[O-]",
"smiles_sanitized":"CC(=O)O"
}
},
{
"id":"cpd00379_E",
"name":"Glutarate",
"compartment":"E",
"annotation":{
"smiles":"O=C([O-])CCCC(=O)[O-]",
"smiles_sanitized":"O=C(O)CCCC(=O)O"
}
},
{
"id":"cpd02678_C",
"name":"N-Formyl-GAR",
"compartment":"C",
"annotation":{
"smiles":"O=CNCC(=O)N[C@@H]1O[C@H](COP(=O)([O-])[O-])[C@@H](O)[C@H]1O",
"smiles_sanitized":"O=CNCC(=O)NC1OC(COP(=O)(O)O)C(O)C1O"
}
},
{
"id":"cpd16650_C",
"name":"7-carboxy-3-hydroxyheptanoyl-coa",
"compartment":"C",
"annotation":{
"smiles":"",
"smiles_sanitized":""
}
},
{
"id":"cpd32368_C",
"name":"3-oxo-(7Z)-hexadecenoyl-CoA",
"compartment":"C",
"annotation":{
"smiles":"CCCCCCCC/C=C\\CCCC(=O)CC(=O)SCCNC(=O)CCNC(=O)[C@H](O)C(C)(C)COP(=O)([O-])OP(=O)([O-])OC[C@H]1O[C@@H](n2cnc3c(N)ncnc32)[C@H](O)[C@@H]1OP(=O)([O-])[O-]",
"smiles_sanitized":"CCCCCCCCC=CCCCC(=O)CC(=O)SCCNC(=O)CCNC(=O)C(O)C(C)(C)COP(=O)(O)OP(=O)(O)OCC1OC(n2cnc3c(N)ncnc32)C(O)C1OP(=O)(O)O"
}
},
{
"id":"cpd00223_C",
"name":"Glycerate",
"compartment":"C",
"annotation":{
"smiles":"O=C([O-])[C@H](O)CO",
"smiles_sanitized":"O=C(O)C(O)CO"
}
},
{
"id":"cpd00192_C",
"name":"4-Coumaroyl-CoA",
"compartment":"C",
"annotation":{
"smiles":"CC(C)(COP(=O)([O-])OP(=O)([O-])OC[C@H]1O[C@@H](n2cnc3c(N)ncnc32)[C@H](O)[C@@H]1OP(=O)([O-])[O-])[C@@H](O)C(=O)NCCC(=O)NCCSC(=O)/C=C/c1ccc(O)cc1",
"smiles_sanitized":"CC(C)(COP(=O)(O)OP(=O)(O)OCC1OC(n2cnc3c(N)ncnc32)C(O)C1OP(=O)(O)O)C(O)C(=O)NCCC(=O)NCCSC(=O)C=Cc1ccc(O)cc1"
}
},
{
"id":"cpd00106_E",
"name":"Fumarate",
"compartment":"E",
"annotation":{
"smiles":"O=C([O-])/C=C/C(=O)[O-]",
"smiles_sanitized":"O=C(O)C=CC(=O)O"
}
},
{
"id":"cpd00311_C",
"name":"Guanosine",
"compartment":"C",
"annotation":{
"smiles":"Nc1nc2c(ncn2[C@@H]2O[C@H](CO)[C@@H](O)[C@H]2O)c(=O)[nH]1",
"smiles_sanitized":"Nc1nc2c(ncn2C2OC(CO)C(O)C2O)c(=O)[nH]1"
}
},
{
"id":"cpd00266_E",
"name":"Carnitine",
"compartment":"E",
"annotation":{
"smiles":"C[N+](C)(C)C[C@H](O)CC(=O)[O-]",
"smiles_sanitized":"C[N+](C)(C)CC(O)CC(=O)O"
}
},
{
"id":"cpd03445_C",
"name":"2-Octaprenyl-6-hydroxyphenol",
"compartment":"C",
"annotation":{
"smiles":"CC(C)=CCC/C(C)=C/CC/C(C)=C/CC/C(C)=C/CC/C(C)=C/CC/C(C)=C/CC/C(C)=C/CC/C(C)=C/Cc1cccc(O)c1O",
"smiles_sanitized":"CC(C)=CCCC(C)=CCCC(C)=CCCC(C)=CCCC(C)=CCCC(C)=CCCC(C)=CCCC(C)=CCc1cccc(O)c1O"
}
},
{
"id":"cpd00817_C",
"name":"D-Arabinose5-phosphate",
"compartment":"C",
"annotation":{
"smiles":"O=C[C@@H](O)[C@H](O)[C@H](O)COP(=O)([O-])[O-]",
"smiles_sanitized":"O=CC(O)C(O)C(O)COP(=O)(O)O"
}
},
{
"id":"VAL_TRNAS_C",
"name":"valine trna",
"compartment":"C"
},
{
"id":"ACYL_ACP_LPS_C",
"name":"ACYL-ACP_LPS",
"compartment":"C"
},
{
"id":"CHARGED_ARG_TRNAS_C",
"name":"charged arginine trna",
"compartment":"C"
},
{
"id":"cpd00482_C",
"name":"2-Phospho-D-glycerate",
"compartment":"C",
"annotation":{
"smiles":"O=C([O-])[C@@H](CO)OP(=O)([O-])[O-]",
"smiles_sanitized":"O=C(O)C(CO)OP(=O)(O)O"
}
},
{
"id":"LEU_TRNAS_C",
"name":"leucine-trna",
"compartment":"C"
},
{
"id":"cpd02182_C",
"name":"(R)-S-Lactoylglutathione",
"compartment":"C",
"annotation":{
"smiles":"C[C@@H](O)C(=O)SC[C@H](NC(=O)CC[C@H]([NH3+])C(=O)[O-])C(=O)NCC(=O)[O-]",
"smiles_sanitized":"CC(O)C(=O)SCC(NC(=O)CCC(N)C(=O)O)C(=O)NCC(=O)O"
}
},
{
"id":"cpd00881_P",
"name":"Caffeate",
"compartment":"P",
"annotation":{
"smiles":"O=C([O-])/C=C/c1ccc(O)c(O)c1",
"smiles_sanitized":"O=C(O)C=Cc1ccc(O)c(O)c1"
}
},
{
"id":"C8_0_2ENOYLACP_C",
"name":"trans-octa-2-enoyl-acp",
"compartment":"C"
},
{
"id":"cpd00009_E",
"name":"Phosphate",
"compartment":"E",
"annotation":{
"smiles":"O=P([O-])([O-])O",
"smiles_sanitized":"O=P(O)(O)O"
}
},
{
"id":"cpd00343_C",
"name":"N-Carbamoyl-L-aspartate",
"compartment":"C",
"annotation":{
"smiles":"NC(=O)N[C@@H](CC(=O)[O-])C(=O)[O-]",
"smiles_sanitized":"NC(=O)NC(CC(=O)O)C(=O)O"
}
},
{
"id":"cpd00083_C",
"name":"GDP-mannose",
"compartment":"C",
"annotation":{
"smiles":"Nc1nc2c(ncn2C2O[C@H](COP(=O)([O-])OP(=O)([O-])O[C@H]3O[C@H](CO)[C@@H](O)[C@H](O)[C@@H]3O)[C@@H](O)[C@H]2O)c(=O)[nH]1",
"smiles_sanitized":"Nc1nc2c(ncn2C2OC(COP(=O)(O)OP(=O)(O)OC3OC(CO)C(O)C(O)C3O)C(O)C2O)c(=O)[nH]1"
}
},
{
"id":"L_1_PHOSPHATIDYL_SERINE_C",
"name":"generic l-1-phosphatidyl-serine",
"compartment":"C"
},
{
"id":"cpd00978_C",
"name":"dUDP",
"compartment":"C",
"annotation":{
"smiles":"O=c1ccn([C@H]2C[C@H](O)[C@@H](COP(=O)([O-])OP(=O)([O-])O)O2)c(=O)[nH]1",
"smiles_sanitized":"O=c1ccn(C2CC(O)C(COP(=O)(O)OP(=O)(O)O)O2)c(=O)[nH]1"
}
},
{
"id":"cpd08615_C",
"name":"1-Hydroxy-2-methyl-2-butenyl 4-diphosphate",
"compartment":"C",
"annotation":{
"smiles":"C/C(=C\\COP(=O)([O-])OP(=O)([O-])O)CO",
"smiles_sanitized":"CC(=CCOP(=O)(O)OP(=O)(O)O)CO"
}
},
{
"id":"cpd15561_C",
"name":"Ubiquinol-8",
"compartment":"C",
"annotation":{
"smiles":"COc1c(O)c(C)c(C/C=C(\\C)CC/C=C(\\C)CC/C=C(\\C)CC/C=C(\\C)CC/C=C(\\C)CC/C=C(\\C)CC/C=C(\\C)CCC=C(C)C)c(O)c1OC",
"smiles_sanitized":"COc1c(O)c(C)c(CC=C(C)CCC=C(C)CCC=C(C)CCC=C(C)CCC=C(C)CCC=C(C)CCC=C(C)CCC=C(C)C)c(O)c1OC"
}
},
{
"id":"cpd00246_E",
"name":"Inosine",
"compartment":"E",
"annotation":{
"smiles":"O=c1[nH]cnc2c1ncn2[C@@H]1O[C@H](CO)[C@@H](O)[C@H]1O",
"smiles_sanitized":"O=c1[nH]cnc2c1ncn2C1OC(CO)C(O)C1O"
}
},
{
"id":"cpd11677_C",
"name":"Triglyceride",
"compartment":"C",
"annotation":{
"smiles":"*C(=O)OCC(COC(*)=O)OC(*)=O",
"smiles_sanitized":"*C(=O)OCC(COC(*)=O)OC(*)=O"
}
},
{
"id":"cpd11579_E",
"name":"ethanesulfonate",
"compartment":"E",
"annotation":{
"smiles":"CCS(=O)(=O)[O-]",
"smiles_sanitized":"CCS(=O)(=O)O"
}
},
{
"id":"cpd02097_C",
"name":"L-Glutamyl 5-phosphate",
"compartment":"C",
"annotation":{
"smiles":"[NH3+][C@@H](CCC(=O)OP(=O)([O-])[O-])C(=O)[O-]",
"smiles_sanitized":"NC(CCC(=O)OP(=O)(O)O)C(=O)O"
}
},
{
"id":"cpd03426_C",
"name":"Sirohydrochlorin",
"compartment":"C",
"annotation":{
"smiles":"C[C@@]1(CC(=O)[O-])c2cc3[nH]c(cc4nc(cc5[nH]c(cc(n2)[C@H]1CCC(=O)[O-])[C@@](C)(CC(=O)[O-])[C@@H]5CCC(=O)[O-])C(CC(=O)[O-])=C4CCC(=O)[O-])c(CCC(=O)[O-])c3CC(=O)[O-]",
"smiles_sanitized":"CC1(CC(=O)O)c2cc3[nH]c(cc4nc(cc5[nH]c(cc(n2)C1CCC(=O)O)C(C)(CC(=O)O)C5CCC(=O)O)C(CC(=O)O)=C4CCC(=O)O)c(CCC(=O)O)c3CC(=O)O"
}
},
{
"id":"cpd02642_C",
"name":"N-5-phosphoribosyl-anthranilate",
"compartment":"C",
"annotation":{
"smiles":"O=C([O-])c1ccccc1N[C@@H]1O[C@H](COP(=O)([O-])[O-])[C@@H](O)[C@H]1O",
"smiles_sanitized":"O=C(O)c1ccccc1NC1OC(COP(=O)(O)O)C(O)C1O"
}
},
{
"id":"cpd32607_C",
"name":"(S)-3-hydroxy-(7Z)-hexadecenoyl-CoA",
"compartment":"C",
"annotation":{
"smiles":"CCCCCCCC/C=C\\CCC[C@H](O)CC(=O)SCCNC(=O)CCNC(=O)[C@H](O)C(C)(C)COP(=O)([O-])OP(=O)([O-])OC[C@H]1O[C@@H](n2cnc3c(N)ncnc32)[C@H](O)[C@@H]1OP(=O)([O-])[O-]",
"smiles_sanitized":"CCCCCCCCC=CCCCC(O)CC(=O)SCCNC(=O)CCNC(=O)C(O)C(C)(C)COP(=O)(O)OP(=O)(O)OCC1OC(n2cnc3c(N)ncnc32)C(O)C1OP(=O)(O)O"
}
},
{
"id":"cpd08287_C",
"name":"4--cytidine5-diphospho-2-C-methyl-D-erythritol",
"compartment":"C",
"annotation":{
"smiles":"C[C@](O)(CO)[C@H](O)COP(=O)([O-])OP(=O)([O-])OC[C@H]1O[C@@H](n2ccc(N)nc2=O)[C@H](O)[C@@H]1O",
"smiles_sanitized":"CC(O)(CO)C(O)COP(=O)(O)OP(=O)(O)OCC1OC(n2ccc(N)nc2=O)C(O)C1O"
}
},
{
"id":"cpd03607_C",
"name":"4-(Phosphonooxy)-threonine",
"compartment":"C",
"annotation":{
"smiles":"[NH3+][C@H](C(=O)[O-])[C@H](O)COP(=O)([O-])[O-]",
"smiles_sanitized":"NC(C(=O)O)C(O)COP(=O)(O)O"
}
},
{
"id":"cpd00424_C",
"name":"Cystathionine",
"compartment":"C",
"annotation":{
"smiles":"[NH3+][C@H](CSCC[C@@H]([NH3+])C(=O)[O-])C(=O)[O-]",
"smiles_sanitized":"NC(CCSCC(N)C(=O)O)C(=O)O"
}
},
{
"id":"cpd02210_C",
"name":"Indoleglycerol phosphate",
"compartment":"C",
"annotation":{
"smiles":"O=P([O-])([O-])OC[C@@H](O)[C@@H](O)c1c[nH]c2ccccc12",
"smiles_sanitized":"O=P(O)(O)OCC(O)C(O)c1c[nH]c2ccccc12"
}
},
{
"id":"cpd00660_C",
"name":"Pentanoyl-CoA",
"compartment":"C",
"annotation":{
"smiles":"CCCCC(=O)SCCNC(=O)CCNC(=O)[C@H](O)C(C)(C)COP(=O)([O-])OP(=O)([O-])OC[C@H]1O[C@@H](n2cnc3c(N)ncnc32)[C@H](O)[C@@H]1OP(=O)([O-])[O-]",
"smiles_sanitized":"CCCCC(=O)SCCNC(=O)CCNC(=O)C(O)C(C)(C)COP(=O)(O)OP(=O)(O)OCC1OC(n2cnc3c(N)ncnc32)C(O)C1OP(=O)(O)O"
}
},
{
"id":"cpd08288_C",
"name":"2-phospho-4--cytidine5-diphospho-2-C-methyl-D-erythritol",
"compartment":"C",
"annotation":{
"smiles":"C[C@@](CO)(OP(=O)([O-])[O-])[C@H](O)COP(=O)([O-])OP(=O)([O-])OC[C@H]1O[C@@H](n2ccc(N)nc2=O)[C@H](O)[C@@H]1O",
"smiles_sanitized":"CC(CO)(OP(=O)(O)O)C(O)COP(=O)(O)OP(=O)(O)OCC1OC(n2ccc(N)nc2=O)C(O)C1O"
}
},
{
"id":"cpd03002_C",
"name":"Lipid A disaccharide",
"compartment":"C",
"annotation":{
"smiles":"CCCCCCCCCCC[C@@H](O)CC(=O)N[C@H]1[C@H](OC[C@H]2O[C@H](OP(=O)([O-])[O-])[C@H](NC(=O)C[C@H](O)CCCCCCCCCCC)[C@@H](OC(=O)C[C@H](O)CCCCCCCCCCC)[C@@H]2O)O[C@H](CO)[C@@H](O)[C@@H]1OC(=O)C[C@H](O)CCCCCCCCCCC",
"smiles_sanitized":"CCCCCCCCCCCC(O)CC(=O)NC1C(OCC2OC(OP(=O)(O)O)C(NC(=O)CC(O)CCCCCCCCCCC)C(OC(=O)CC(O)CCCCCCCCCCC)C2O)OC(CO)C(O)C1OC(=O)CC(O)CCCCCCCCCCC"
}
},
{
"id":"cpd00212_E",
"name":"L-Sorbose",
"compartment":"E",
"annotation":{
"smiles":"OCC1(O)OC[C@H](O)[C@@H](O)[C@@H]1O",
"smiles_sanitized":"OCC1(O)OCC(O)C(O)C1O"
}
},
{
"id":"cpd00731_C",
"name":"Ala-Ala",
"compartment":"C",
"annotation":{
"smiles":"C[C@@H]([NH3+])C(=O)N[C@H](C)C(=O)[O-]",
"smiles_sanitized":"CC(N)C(=O)NC(C)C(=O)O"
}
},
{
"id":"cpd00211_C",
"name":"Butyrate",
"compartment":"C",
"annotation":{
"smiles":"CCCC(=O)[O-]",
"smiles_sanitized":"CCCC(=O)O"
}
},
{
"id":"cpd01080_C",
"name":"ocdca",
"compartment":"C",
"annotation":{
"smiles":"CCCCCCCCCCCCCCCCCC(=O)[O-]",
"smiles_sanitized":"CCCCCCCCCCCCCCCCCC(=O)O"
}
},
{
"id":"cpd00641_C",
"name":"L-Histidinol",
"compartment":"C",
"annotation":{
"smiles":"[NH3+][C@H](CO)Cc1c[nH]cn1",
"smiles_sanitized":"NC(CO)Cc1c[nH]cn1"
}
},
{
"id":"cpd00206_C",
"name":"dCMP",
"compartment":"C",
"annotation":{
"smiles":"Nc1ccn([C@H]2C[C@H](O)[C@@H](COP(=O)([O-])[O-])O2)c(=O)n1",
"smiles_sanitized":"Nc1ccn(C2CC(O)C(COP(=O)(O)O)O2)c(=O)n1"
}
},
{
"id":"cpd00023_E",
"name":"L-Glutamate",
"compartment":"E",
"annotation":{
"smiles":"[NH3+][C@@H](CCC(=O)[O-])C(=O)[O-]",
"smiles_sanitized":"NC(CCC(=O)O)C(=O)O"
}
},
{
"id":"cpd01887_C",
"name":"(3Z)-Dodecenoyl-CoA",
"compartment":"C",
"annotation":{
"smiles":"CCCCCCCC/C=C\\CC(=O)SCCNC(=O)CCNC(=O)C(O)C(C)(C)COP(=O)([O-])OP(=O)([O-])OCC1OC(n2cnc3c(N)ncnc32)C(O)C1OP(=O)([O-])[O-]",
"smiles_sanitized":"CCCCCCCCC=CCC(=O)SCCNC(=O)CCNC(=O)C(O)C(C)(C)COP(=O)(O)OP(=O)(O)OCC1OC(n2cnc3c(N)ncnc32)C(O)C1OP(=O)(O)O"
}
},
{
"id":"cpd01695_C",
"name":"Myristoyl-CoA",
"compartment":"C",
"annotation":{
"smiles":"CCCCCCCCCCCCCC(=O)SCCNC(=O)CCNC(=O)[C@H](O)C(C)(C)COP(=O)([O-])OP(=O)([O-])OC[C@H]1O[C@@H](n2cnc3c(N)ncnc32)[C@H](O)[C@@H]1OP(=O)([O-])[O-]",
"smiles_sanitized":"CCCCCCCCCCCCCC(=O)SCCNC(=O)CCNC(=O)C(O)C(C)(C)COP(=O)(O)OP(=O)(O)OCC1OC(n2cnc3c(N)ncnc32)C(O)C1OP(=O)(O)O"
}
},
{
"id":"cpd01260_C",
"name":"Lauroyl-CoA",
"compartment":"C",
"annotation":{
"smiles":"CCCCCCCCCCCC(=O)SCCNC(=O)CCNC(=O)[C@H](O)C(C)(C)COP(=O)([O-])OP(=O)([O-])OC[C@H]1O[C@@H](n2cnc3c(N)ncnc32)[C@H](O)[C@@H]1OP(=O)([O-])[O-]",
"smiles_sanitized":"CCCCCCCCCCCC(=O)SCCNC(=O)CCNC(=O)C(O)C(C)(C)COP(=O)(O)OP(=O)(O)OCC1OC(n2cnc3c(N)ncnc32)C(O)C1OP(=O)(O)O"
}
},
{
"id":"SER_TRNAS_C",
"name":"serine trna",
"compartment":"C"
},
{
"id":"cpd00712_C",
"name":"2-Dehydropantoate",
"compartment":"C",
"annotation":{
"smiles":"CC(C)(CO)C(=O)C(=O)[O-]",
"smiles_sanitized":"CC(C)(CO)C(=O)C(=O)O"
}
},
{
"id":"C18_0_ACP_C",
"name":"octadenoyl-acp",
"compartment":"C"
},
{
"id":"cpd02636_C",
"name":"4-Methyl-5--2-hydroxyethyl-thiazole",
"compartment":"C",
"annotation":{
"smiles":"Cc1ncsc1CCO",
"smiles_sanitized":"Cc1ncsc1CCO"
}
},
{
"id":"cpd00683_C",
"name":"Dihydropteroate",
"compartment":"C",
"annotation":{
"smiles":"Nc1nc2c(c(=O)[nH]1)N=C(CNc1ccc(C(=O)[O-])cc1)CN2",
"smiles_sanitized":"Nc1nc2c(c(=O)[nH]1)N=C(CNc1ccc(C(=O)O)cc1)CN2"
}
},
{
"id":"cpd00762_C",
"name":"4-Guanidinobutanoate",
"compartment":"C",
"annotation":{
"smiles":"NC(=[NH2+])NCCCC(=O)[O-]",
"smiles_sanitized":"N=C(N)NCCCC(=O)O"
}
},
{
"id":"cpd03521_C",
"name":"Dihydroneopterin phosphate",
"compartment":"C",
"annotation":{
"smiles":"Nc1nc2c(c(=O)[nH]1)N=C([C@H](O)[C@H](O)COP(=O)([O-])[O-])CN2",
"smiles_sanitized":"Nc1nc2c(c(=O)[nH]1)N=C(C(O)C(O)COP(=O)(O)O)CN2"
}
},
{
"id":"cpd00982_C",
"name":"FADH2",
"compartment":"C",
"annotation":{
"smiles":"Cc1cc2c(cc1C)N(C[C@H](O)[C@H](O)[C@H](O)COP(=O)([O-])OP(=O)([O-])OC[C@H]1O[C@@H](n3cnc4c(N)ncnc43)[C@H](O)[C@@H]1O)c1[nH]c(=O)[nH]c(=O)c1N2",
"smiles_sanitized":"Cc1cc2c(cc1C)N(CC(O)C(O)C(O)COP(=O)(O)OP(=O)(O)OCC1OC(n3cnc4c(N)ncnc43)C(O)C1O)c1[nH]c(=O)[nH]c(=O)c1N2"
}
},
{
"id":"cpd00161_C",
"name":"L-Threonine",
"compartment":"C",
"annotation":{
"smiles":"C[C@@H](O)[C@H]([NH3+])C(=O)[O-]",
"smiles_sanitized":"CC(O)C(N)C(=O)O"
}
},
{
"id":"cpd16604_C",
"name":"trans-9-carboxynon-2-enoyl-coa",
"compartment":"C",
"annotation":{
"smiles":"",
"smiles_sanitized":""
}
},
{
"id":"cpd00079_C",
"name":"D-glucose-6-phosphate",
"compartment":"C",
"annotation":{
"smiles":"O=P([O-])([O-])OC[C@H]1OC(O)[C@H](O)[C@@H](O)[C@@H]1O",
"smiles_sanitized":"O=P(O)(O)OCC1OC(O)C(O)C(O)C1O"
}
},
{
"id":"cpd01599_C",
"name":"Glutaconyl-1-CoA",
"compartment":"C",
"annotation":{
"smiles":"CC(C)(COP(=O)([O-])OP(=O)([O-])OC[C@H]1O[C@@H](n2cnc3c(N)ncnc32)[C@H](O)[C@@H]1OP(=O)([O-])[O-])[C@@H](O)C(=O)NCCC(=O)NCCSC(=O)/C=C/CC(=O)[O-]",
"smiles_sanitized":"CC(C)(COP(=O)(O)OP(=O)(O)OCC1OC(n2cnc3c(N)ncnc32)C(O)C1OP(=O)(O)O)C(O)C(=O)NCCC(=O)NCCSC(=O)C=CCC(=O)O"
}
},
{
"id":"cpd00087_C",
"name":"Tetrahydrofolate",
"compartment":"C",
"annotation":{
"smiles":"Nc1nc2c(c(=O)[nH]1)N[C@@H](CNc1ccc(C(=O)N[C@@H](CCC(=O)[O-])C(=O)[O-])cc1)CN2",
"smiles_sanitized":"Nc1nc2c(c(=O)[nH]1)NC(CNc1ccc(C(=O)NC(CCC(=O)O)C(=O)O)cc1)CN2"
}
},
{
"id":"cpd00367_E",
"name":"Cytidine",
"compartment":"E",
"annotation":{
"smiles":"Nc1ccn([C@@H]2O[C@H](CO)[C@@H](O)[C@H]2O)c(=O)n1",
"smiles_sanitized":"Nc1ccn(C2OC(CO)C(O)C2O)c(=O)n1"
}
},
{
"id":"CHARGED_GLN_TRNAS_C",
"name":"charged glutamine trna",
"compartment":"C"
},
{
"id":"cpd35448_C",
"name":"(5Z)-tetradec-5-enoyl-CoA",
"compartment":"C",
"annotation":{
"smiles":"CCCCCCCC/C=C\\CCCC(=O)SCCNC(=O)CCNC(=O)[C@H](O)C(C)(C)COP(=O)([O-])OP(=O)([O-])OC[C@H]1O[C@@H](n2cnc3c(N)ncnc32)[C@H](O)[C@@H]1OP(=O)([O-])[O-]",
"smiles_sanitized":"CCCCCCCCC=CCCCC(=O)SCCNC(=O)CCNC(=O)C(O)C(C)(C)COP(=O)(O)OP(=O)(O)OCC1OC(n2cnc3c(N)ncnc32)C(O)C1OP(=O)(O)O"
}
},
{
"id":"cpd34461_C",
"name":"(7Z)-hexadec-7-enoate",
"compartment":"C",
"annotation":{
"smiles":"CCCCCCCC/C=C\\CCCCCC(=O)[O-]",
"smiles_sanitized":"CCCCCCCCC=CCCCCCC(=O)O"
}
},
{
"id":"cpd00634_E",
"name":"Chlorogenate",
"compartment":"E",
"annotation":{
"smiles":"O=C(/C=C/c1ccc(O)c(O)c1)O[C@@H]1C[C@](O)(C(=O)[O-])C[C@@H](O)[C@H]1O",
"smiles_sanitized":"O=C(C=Cc1ccc(O)c(O)c1)OC1CC(O)(C(=O)O)CC(O)C1O"
}
},
{
"id":"cpd00207_C",
"name":"Guanine",
"compartment":"C",
"annotation":{
"smiles":"Nc1nc2[nH]cnc2c(=O)[nH]1",
"smiles_sanitized":"Nc1nc2[nH]cnc2c(=O)[nH]1"
}
},
{
"id":"cpd00345_C",
"name":"5-Methyltetrahydrofolate",
"compartment":"C",
"annotation":{
"smiles":"CN1c2c(nc(N)[nH]c2=O)NC[C@@H]1CNc1ccc(C(=O)N[C@@H](CCC(=O)[O-])C(=O)[O-])cc1",
"smiles_sanitized":"CN1c2c(nc(N)[nH]c2=O)NCC1CNc1ccc(C(=O)NC(CCC(=O)O)C(=O)O)cc1"
}
},
{
"id":"cpd04511_C",
"name":"4-Hydroxy-3-methoxyphenyl-beta-hydroxypropionyl-CoA",
"compartment":"C",
"annotation":{
"smiles":"COc1cc(C(O)CC(=O)SCCNC(=O)CCNC(=O)[C@H](O)C(C)(C)COP(=O)([O-])OP(=O)([O-])OC[C@H]2O[C@@H](n3cnc4c(N)ncnc43)[C@H](O)[C@@H]2OP(=O)([O-])[O-])ccc1O",
"smiles_sanitized":"COc1cc(C(O)CC(=O)SCCNC(=O)CCNC(=O)C(O)C(C)(C)COP(=O)(O)OP(=O)(O)OCC2OC(n3cnc4c(N)ncnc43)C(O)C2OP(=O)(O)O)ccc1O"
}
},
{
"id":"cpd00465_C",
"name":"Ureidoglycolate",
"compartment":"C",
"annotation":{
"smiles":"NC(=O)N[C@@H](O)C(=O)[O-]",
"smiles_sanitized":"NC(=O)NC(O)C(=O)O"
}
},
{
"id":"CHARGED_ALA_TRNAS_C",
"name":"charged alanine trna",
"compartment":"C"
},
{
"id":"cpd02921_C",
"name":"SAICAR",
"compartment":"C",
"annotation":{
"smiles":"Nc1c(C(=O)NC(CC(=O)[O-])C(=O)[O-])ncn1[C@@H]1O[C@H](COP(=O)([O-])[O-])[C@@H](O)[C@H]1O",
"smiles_sanitized":"Nc1c(C(=O)NC(CC(=O)O)C(=O)O)ncn1C1OC(COP(=O)(O)O)C(O)C1O"
}
},
{
"id":"cpd00040_C",
"name":"Glyoxalate",
"compartment":"C",
"annotation":{
"smiles":"O=CC(=O)[O-]",
"smiles_sanitized":"O=CC(=O)O"
}
},
{
"id":"cpd01741_C",
"name":"ddca",
"compartment":"C",
"annotation":{
"smiles":"CCCCCCCCCCCC(=O)[O-]",
"smiles_sanitized":"CCCCCCCCCCCC(=O)O"
}
},
{
"id":"cpd35321_C",
"name":"(2E,7Z)-hexadecenoyl-CoA",
"compartment":"C",
"annotation":{
"smiles":"CCCCCCCC/C=C\\CCC/C=C/C(=O)SCCNC(=O)CCNC(=O)[C@H](O)C(C)(C)COP(=O)([O-])OP(=O)([O-])OC[C@H]1O[C@@H](n2cnc3c(N)ncnc32)[C@H](O)[C@@H]1OP(=O)([O-])[O-]",
"smiles_sanitized":"CCCCCCCCC=CCCCC=CC(=O)SCCNC(=O)CCNC(=O)C(O)C(C)(C)COP(=O)(O)OP(=O)(O)OCC1OC(n2cnc3c(N)ncnc32)C(O)C1OP(=O)(O)O"
}
},
{
"id":"cpd00225_C",
"name":"Benzaldehyde",
"compartment":"C",
"annotation":{
"smiles":"O=Cc1ccccc1",
"smiles_sanitized":"O=Cc1ccccc1"
}
},
{
"id":"cpd04073_C",
"name":"Vanillate",
"compartment":"C",
"annotation":{
"smiles":"COc1cc(C(=O)[O-])ccc1O",
"smiles_sanitized":"COc1cc(C(=O)O)ccc1O"
}
},
{
"id":"cpd01059_C",
"name":"Ferulate",
"compartment":"C",
"annotation":{
"smiles":"COc1cc(/C=C/C(=O)[O-])ccc1O",
"smiles_sanitized":"COc1cc(C=CC(=O)O)ccc1O"
}
},
{
"id":"CHARGED_CYS_TRNAS_C",
"name":"charged cysteine trna",
"compartment":"C"
},
{
"id":"cpd00216_C",
"name":"Chorismate",
"compartment":"C",
"annotation":{
"smiles":"C=C(O[C@@H]1C=C(C(=O)[O-])C=C[C@H]1O)C(=O)[O-]",
"smiles_sanitized":"C=C(OC1C=C(C(=O)O)C=CC1O)C(=O)O"
}
},
{
"id":"cpd00342_C",
"name":"N-Acetylornithine",
"compartment":"C",
"annotation":{
"smiles":"CC(=O)N[C@@H](CCC[NH3+])C(=O)[O-]",
"smiles_sanitized":"CC(=O)NC(CCCN)C(=O)O"
}
},
{
"id":"cpd14935_C",
"name":"3-Oxostearoyl-CoA",
"compartment":"C",
"annotation":{
"smiles":"CCCCCCCCCCCCCCCC(=O)CC(=O)SCCNC(=O)CCNC(=O)[C@H](O)C(C)(C)COP(=O)([O-])OP(=O)([O-])OC[C@H]1O[C@@H](n2cnc3c(N)ncnc32)[C@H](O)[C@@H]1OP(=O)([O-])[O-]",
"smiles_sanitized":"CCCCCCCCCCCCCCCC(=O)CC(=O)SCCNC(=O)CCNC(=O)C(O)C(C)(C)COP(=O)(O)OP(=O)(O)OCC1OC(n2cnc3c(N)ncnc32)C(O)C1OP(=O)(O)O"
}
},
{
"id":"cpd14937_C",
"name":"(2E)-Octadecenoyl-CoA",
"compartment":"C",
"annotation":{
"smiles":"CCCCCCCCCCCCCCC/C=C/C(=O)SCCNC(=O)CCNC(=O)[C@H](O)C(C)(C)COP(=O)([O-])OP(=O)([O-])OC[C@H]1O[C@@H](n2cnc3c(N)ncnc32)[C@H](O)[C@@H]1OP(=O)([O-])[O-]",
"smiles_sanitized":"CCCCCCCCCCCCCCCC=CC(=O)SCCNC(=O)CCNC(=O)C(O)C(C)(C)COP(=O)(O)OP(=O)(O)OCC1OC(n2cnc3c(N)ncnc32)C(O)C1OP(=O)(O)O"
}
},
{
"id":"cpd00067_C",
"name":"H+",
"compartment":"C",
"annotation":{
"smiles":"[H+]",
"smiles_sanitized":"[H+]"
}
},
{
"id":"C18_1_C9__ACP_C",
"name":"cis-octadec-9-enoyl-acp",
"compartment":"C"
},
{
"id":"CHARGED_THR_TRNAS_C",
"name":"charged threonine trna",
"compartment":"C"
},
{
"id":"cpd01613_C",
"name":"Pent-2-enoyl-CoA",
"compartment":"C",
"annotation":{
"smiles":"CC/C=C/C(=O)SCCNC(=O)CCNC(=O)[C@H](O)C(C)(C)COP(=O)([O-])OP(=O)([O-])OC[C@H]1O[C@@H](n2cnc3c(N)ncnc32)[C@H](O)[C@@H]1OP(=O)([O-])[O-]",
"smiles_sanitized":"CCC=CC(=O)SCCNC(=O)CCNC(=O)C(O)C(C)(C)COP(=O)(O)OP(=O)(O)OCC1OC(n2cnc3c(N)ncnc32)C(O)C1OP(=O)(O)O"
}
},
{
"id":"cpd00111_C",
"name":"Oxidized glutathione",
"compartment":"C",
"annotation":{
"smiles":"[NH3+][C@@H](CCC(=O)N[C@@H](CSSC[C@H](NC(=O)CC[C@H]([NH3+])C(=O)[O-])C(=O)NCC(=O)[O-])C(=O)NCC(=O)[O-])C(=O)[O-]",
"smiles_sanitized":"NC(CCC(=O)NC(CSSCC(NC(=O)CCC(N)C(=O)O)C(=O)NCC(=O)O)C(=O)NCC(=O)O)C(=O)O"
}
},
{
"id":"cpd02930_C",
"name":"Lipid X",
"compartment":"C",
"annotation":{
"smiles":"CCCCCCCCCCC[C@@H](O)CC(=O)N[C@H]1[C@@H](OP(=O)([O-])[O-])O[C@H](CO)[C@@H](O)[C@@H]1OC(=O)C[C@H](O)CCCCCCCCCCC",
"smiles_sanitized":"CCCCCCCCCCCC(O)CC(=O)NC1C(OP(=O)(O)O)OC(CO)C(O)C1OC(=O)CC(O)CCCCCCCCCCC"
}
},
{
"id":"cpd00089_C",
"name":"Glucose-1-phosphate",
"compartment":"C",
"annotation":{
"smiles":"O=P([O-])([O-])O[C@H]1O[C@H](CO)[C@@H](O)[C@H](O)[C@H]1O",
"smiles_sanitized":"O=P(O)(O)OC1OC(CO)C(O)C(O)C1O"
}
},
{
"id":"C8_0_3HYDROXYACP_C",
"name":"3-hydroxyoctanoyl-acp",
"compartment":"C"
},
{
"id":"cpd00281_E",
"name":"GABA",
"compartment":"E",
"annotation":{
"smiles":"[NH3+]CCCC(=O)[O-]",
"smiles_sanitized":"NCCCC(=O)O"
}
},
{
"id":"cpd15394_C",
"name":"Allulose 6-phosphate",
"compartment":"C",
"annotation":{
"smiles":"O=C(CO)[C@@H](O)[C@@H](O)[C@@H](O)COP(=O)([O-])[O-]",
"smiles_sanitized":"O=C(CO)C(O)C(O)C(O)COP(=O)(O)O"
}
},
{
"id":"cpd00791_C",
"name":"ProtoporphyrinogenIX",
"compartment":"C",
"annotation":{
"smiles":"C=Cc1c2[nH]c(c1C)Cc1[nH]c(c(CCC(=O)[O-])c1C)Cc1[nH]c(c(C)c1CCC(=O)[O-])Cc1[nH]c(c(C)c1C=C)C2",
"smiles_sanitized":"C=Cc1c2[nH]c(c1C)Cc1[nH]c(c(CCC(=O)O)c1C)Cc1[nH]c(c(C)c1CCC(=O)O)Cc1[nH]c(c(C)c1C=C)C2"
}
},
{
"id":"cpd16606_C",
"name":"9-carboxy-3-oxononanoyl-coa",
"compartment":"C",
"annotation":{
"smiles":"",
"smiles_sanitized":""
}
},
{
"id":"cpd00186_C",
"name":"D-Glutamate",
"compartment":"C",
"annotation":{
"smiles":"[NH3+][C@H](CCC(=O)[O-])C(=O)[O-]",
"smiles_sanitized":"NC(CCC(=O)O)C(=O)O"
}
},
{
"id":"cpd02140_C",
"name":"AIR",
"compartment":"C",
"annotation":{
"smiles":"Nc1c[nH+]cn1[C@@H]1O[C@H](COP(=O)([O-])[O-])[C@@H](O)[C@H]1O",
"smiles_sanitized":"Nc1cncn1C1OC(COP(=O)(O)O)C(O)C1O"
}
},
{
"id":"cpd00171_C",
"name":"D-Ribulose5-phosphate",
"compartment":"C",
"annotation":{
"smiles":"O=C(CO)[C@H](O)[C@H](O)COP(=O)([O-])[O-]",
"smiles_sanitized":"O=C(CO)C(O)C(O)COP(=O)(O)O"
}
},
{
"id":"cpd03449_C",
"name":"2-Octaprenyl-3-methyl-5-hydroxy-6-methoxy-1,4-benzoquinone",
"compartment":"C",
"annotation":{
"smiles":"COc1c(O)c(O)c(C)c(C/C=C(\\C)CC/C=C(\\C)CC/C=C(\\C)CC/C=C(\\C)CC/C=C(\\C)CC/C=C(\\C)CC/C=C(\\C)CCC=C(C)C)c1O",
"smiles_sanitized":"COc1c(O)c(O)c(C)c(CC=C(C)CCC=C(C)CCC=C(C)CCC=C(C)CCC=C(C)CCC=C(C)CCC=C(C)CCC=C(C)C)c1O"
}
},
{
"id":"cpd05192_C",
"name":"sebacate",
"compartment":"C",
"annotation":{
"smiles":"O=C([O-])CCCCCCCCC(=O)[O-]",
"smiles_sanitized":"O=C(O)CCCCCCCCC(=O)O"
}
},
{
"id":"cpd00009_C",
"name":"Phosphate",
"compartment":"C",
"annotation":{
"smiles":"O=P([O-])([O-])O",
"smiles_sanitized":"O=P(O)(O)O"
}
},
{
"id":"cpd03444_C",
"name":"2-Octaprenylphenol",
"compartment":"C",
"annotation":{
"smiles":"CC(C)=CCC/C(C)=C/CC/C(C)=C/CC/C(C)=C/CC/C(C)=C/CC/C(C)=C/CC/C(C)=C/CC/C(C)=C/Cc1ccccc1O",
"smiles_sanitized":"CC(C)=CCCC(C)=CCCC(C)=CCCC(C)=CCCC(C)=CCCC(C)=CCCC(C)=CCCC(C)=CCc1ccccc1O"
}
},
{
"id":"cpd00356_C",
"name":"dCTP",
"compartment":"C",
"annotation":{
"smiles":"Nc1ccn([C@H]2C[C@H](O)[C@@H](COP(=O)([O-])OP(=O)([O-])OP(=O)([O-])O)O2)c(=O)n1",
"smiles_sanitized":"Nc1ccn(C2CC(O)C(COP(=O)(O)OP(=O)(O)OP(=O)(O)O)O2)c(=O)n1"
}
},
{
"id":"cpd00755_C",
"name":"Hydroxymethylbilane",
"compartment":"C",
"annotation":{
"smiles":"O=C([O-])CCc1c[nH]c(Cc2[nH]c(Cc3[nH]c(Cc4[nH]c(CO)c(CC(=O)[O-])c4CCC(=O)[O-])c(CC(=O)[O-])c3CCC(=O)[O-])c(CC(=O)[O-])c2CCC(=O)[O-])c1CC(=O)[O-]",
"smiles_sanitized":"O=C(O)CCc1c[nH]c(Cc2[nH]c(Cc3[nH]c(Cc4[nH]c(CO)c(CC(=O)O)c4CCC(=O)O)c(CC(=O)O)c3CCC(=O)O)c(CC(=O)O)c2CCC(=O)O)c1CC(=O)O"
}
},
{
"id":"cpd00296_C",
"name":"dGMP",
"compartment":"C",
"annotation":{
"smiles":"Nc1nc2c(ncn2[C@H]2C[C@H](O)[C@@H](COP(=O)([O-])[O-])O2)c(=O)[nH]1",
"smiles_sanitized":"Nc1nc2c(ncn2C2CC(O)C(COP(=O)(O)O)O2)c(=O)[nH]1"
}
},
{
"id":"cpd01113_E",
"name":"Hexanoate",
"compartment":"E",
"annotation":{
"smiles":"CCCCCC(=O)[O-]",
"smiles_sanitized":"CCCCCC(=O)O"
}
},
{
"id":"cpd00118_C",
"name":"Putrescine",
"compartment":"C",
"annotation":{
"smiles":"[NH3+]CCCC[NH3+]",
"smiles_sanitized":"NCCCCN"
}
},
{
"id":"cpd00162_C",
"name":"Aminoethanol",
"compartment":"C",
"annotation":{
"smiles":"[NH3+]CCO",
"smiles_sanitized":"NCCO"
}
},
{
"id":"cpd05192_E",
"name":"sebacate",
"compartment":"E",
"annotation":{
"smiles":"O=C([O-])CCCCCCCCC(=O)[O-]",
"smiles_sanitized":"O=C(O)CCCCCCCCC(=O)O"
}
},
{
"id":"cpd00320_E",
"name":"D-Aspartate",
"compartment":"E",
"annotation":{
"smiles":"[NH3+][C@H](CC(=O)[O-])C(=O)[O-]",
"smiles_sanitized":"NC(CC(=O)O)C(=O)O"
}
},
{
"id":"cpd00644_C",
"name":"PAN",
"compartment":"C",
"annotation":{
"smiles":"CC(C)(CO)[C@@H](O)C(=O)NCCC(=O)[O-]",
"smiles_sanitized":"CC(C)(CO)C(O)C(=O)NCCC(=O)O"
}
},
{
"id":"cpd00360_C",
"name":"Purine",
"compartment":"C",
"annotation":{
"smiles":"c1ncc2[nH]cnc2n1",
"smiles_sanitized":"c1ncc2[nH]cnc2n1"
}
},
{
"id":"L_1_PHOSPHATIDYL_ETHANOLAMINE_C",
"name":"generic l-1-phosphatidyl-ethanolamine",
"compartment":"C"
},
{
"id":"cpd00123_C",
"name":"3-Methyl-2-oxobutanoate",
"compartment":"C",
"annotation":{
"smiles":"CC(C)C(=O)C(=O)[O-]",
"smiles_sanitized":"CC(C)C(=O)C(=O)O"
}
},
{
"id":"cpd00634_P",
"name":"Chlorogenate",
"compartment":"P",
"annotation":{
"smiles":"O=C(/C=C/c1ccc(O)c(O)c1)O[C@@H]1C[C@](O)(C(=O)[O-])C[C@@H](O)[C@H]1O",
"smiles_sanitized":"O=C(C=Cc1ccc(O)c(O)c1)OC1CC(O)(C(=O)O)CC(O)C1O"
}
},
{
"id":"cpd00807_C",
"name":"L-histidinol-phosphate",
"compartment":"C",
"annotation":{
"smiles":"[NH3+][C@H](COP(=O)([O-])[O-])Cc1c[nH]cn1",
"smiles_sanitized":"NC(COP(=O)(O)O)Cc1c[nH]cn1"
}
},
{
"id":"cpd00011_E",
"name":"CO2",
"compartment":"E",
"annotation":{
"smiles":"O=C=O",
"smiles_sanitized":"O=C=O"
}
},
{
"id":"cpd03118_C",
"name":"(S)-Hydroxydecanoyl-CoA",
"compartment":"C",
"annotation":{
"smiles":"CCCCCCC[C@H](O)CC(=O)SCCNC(=O)CCNC(=O)[C@H](O)C(C)(C)COP(=O)([O-])OP(=O)([O-])OC[C@H]1O[C@@H](n2cnc3c(N)ncnc32)[C@H](O)[C@@H]1OP(=O)([O-])[O-]",
"smiles_sanitized":"CCCCCCCC(O)CC(=O)SCCNC(=O)CCNC(=O)C(O)C(C)(C)COP(=O)(O)OP(=O)(O)OCC1OC(n2cnc3c(N)ncnc32)C(O)C1OP(=O)(O)O"
}
},
{
"id":"cpd00100_E",
"name":"Glycerol",
"compartment":"E",
"annotation":{
"smiles":"OCC(O)CO",
"smiles_sanitized":"OCC(O)CO"
}
},
{
"id":"C14_0_2ENOYLCOA_C",
"name":"trans-tetradec-2-enoyl-acp",
"compartment":"C"
},
{
"id":"cpd02821_C",
"name":"1,2-Dihydroxycyclohexa-3,5-diene-1-carboxylate",
"compartment":"C",
"annotation":{
"smiles":"O=C([O-])C1(O)C=CC=CC1O",
"smiles_sanitized":"O=C(O)C1(O)C=CC=CC1O"
}
},
{
"id":"cpd00295_C",
"name":"dGDP",
"compartment":"C",
"annotation":{
"smiles":"Nc1nc2c(ncn2[C@H]2C[C@H](O)[C@@H](COP(=O)([O-])OP(=O)([O-])O)O2)c(=O)[nH]1",
"smiles_sanitized":"Nc1nc2c(ncn2C2CC(O)C(COP(=O)(O)OP(=O)(O)O)O2)c(=O)[nH]1"
}
},
{
"id":"cpd00129_C",
"name":"L-Proline",
"compartment":"C",
"annotation":{
"smiles":"O=C([O-])[C@@H]1CCC[NH2+]1",
"smiles_sanitized":"O=C(O)C1CCCN1"
}
},
{
"id":"cpd01352_C",
"name":"Undecaprenol",
"compartment":"C",
"annotation":{
"smiles":"CC/C=C(\\C)CC/C=C(\\C)CC/C=C(\\C)CC/C=C(\\C)CC/C=C(\\C)CC/C=C(\\C)CC/C=C(\\C)CC/C=C(\\C)CC/C=C(\\C)CC/C=C(\\C)CCC=C(C)C",
"smiles_sanitized":"CCC=C(C)CCC=C(C)CCC=C(C)CCC=C(C)CCC=C(C)CCC=C(C)CCC=C(C)CCC=C(C)CCC=C(C)CCC=C(C)CCC=C(C)C"
}
},
{
"id":"cpd00298_C",
"name":"dTMP",
"compartment":"C",
"annotation":{
"smiles":"Cc1cn([C@H]2C[C@H](O)[C@@H](COP(=O)([O-])[O-])O2)c(=O)[nH]c1=O",
"smiles_sanitized":"Cc1cn(C2CC(O)C(COP(=O)(O)O)O2)c(=O)[nH]c1=O"
}
},
{
"id":"cpd00004_C",
"name":"NADH",
"compartment":"C",
"annotation":{
"smiles":"NC(=O)C1=CN([C@@H]2O[C@H](COP(=O)([O-])OP(=O)([O-])OC[C@H]3O[C@@H](n4cnc5c(N)ncnc54)[C@H](O)[C@@H]3O)[C@@H](O)[C@H]2O)C=CC1",
"smiles_sanitized":"NC(=O)C1=CN(C2OC(COP(=O)(O)OP(=O)(O)OCC3OC(n4cnc5c(N)ncnc54)C(O)C3O)C(O)C2O)C=CC1"
}
},
{
"id":"cpd00091_C",
"name":"UMP",
"compartment":"C",
"annotation":{
"smiles":"O=c1ccn([C@@H]2O[C@H](COP(=O)([O-])[O-])[C@@H](O)[C@H]2O)c(=O)[nH]1",
"smiles_sanitized":"O=c1ccn(C2OC(COP(=O)(O)O)C(O)C2O)c(=O)[nH]1"
}
},
{
"id":"cpd00128_C",
"name":"Adenine",
"compartment":"C",
"annotation":{
"smiles":"Nc1ncnc2[nH]cnc12",
"smiles_sanitized":"Nc1ncnc2[nH]cnc12"
}
},
{
"id":"CDPDIACYLGLYCEROL_C",
"name":"generic cdp-diacylglycerol",
"compartment":"C"
},
{
"id":"cpd00308_C",
"name":"Malonate",
"compartment":"C",
"annotation":{
"smiles":"O=C([O-])CC(=O)[O-]",
"smiles_sanitized":"O=C(O)CC(=O)O"
}
},
{
"id":"cpd00604_C",
"name":"4-Coumarate",
"compartment":"C",
"annotation":{
"smiles":"O=C([O-])/C=C/c1ccc(O)cc1",
"smiles_sanitized":"O=C(O)C=Cc1ccc(O)cc1"
}
},
{
"id":"cpd00001_E",
"name":"H2O",
"compartment":"E",
"annotation":{
"smiles":"O",
"smiles_sanitized":"O"
}
},
{
"id":"cpd00367_C",
"name":"Cytidine",
"compartment":"C",
"annotation":{
"smiles":"Nc1ccn([C@@H]2O[C@H](CO)[C@@H](O)[C@H]2O)c(=O)n1",
"smiles_sanitized":"Nc1ccn(C2OC(CO)C(O)C2O)c(=O)n1"
}
},
{
"id":"cpd01553_C",
"name":"Saligenin",
"compartment":"C",
"annotation":{
"smiles":"OCc1ccccc1O",
"smiles_sanitized":"OCc1ccccc1O"
}
},
{
"id":"cpd00214_C",
"name":"Palmitate",
"compartment":"C",
"annotation":{
"smiles":"CCCCCCCCCCCCCCCC(=O)[O-]",
"smiles_sanitized":"CCCCCCCCCCCCCCCC(=O)O"
}
},
{
"id":"cpd25464_C",
"name":"propionitrile",
"compartment":"C",
"annotation":{
"smiles":"CCC#N",
"smiles_sanitized":"CCC#N"
}
},
{
"id":"cpd00876_C",
"name":"3-Hydroxyisobutyrate",
"compartment":"C",
"annotation":{
"smiles":"CC(CO)C(=O)[O-]",
"smiles_sanitized":"CC(CO)C(=O)O"
}
},
{
"id":"cpd00006_C",
"name":"NADP",
"compartment":"C",
"annotation":{
"smiles":"NC(=O)c1ccc[n+]([C@@H]2O[C@H](COP(=O)([O-])OP(=O)([O-])OC[C@H]3O[C@@H](n4cnc5c(N)ncnc54)[C@H](OP(=O)([O-])[O-])[C@@H]3O)[C@@H](O)[C@H]2O)c1",
"smiles_sanitized":"NC(=O)c1ccc[n+](C2OC(COP(=O)(O)OP(=O)(O)OCC3OC(n4cnc5c(N)ncnc54)C(OP(=O)(O)O)C3O)C(O)C2O)c1"
}
},
{
"id":"cpd16657_C",
"name":"ethylsalicylate",
"compartment":"C",
"annotation":{
"smiles":"",
"smiles_sanitized":""
}
},
{
"id":"cpd00875_C",
"name":"KDO",
"compartment":"C",
"annotation":{
"smiles":"O=C([O-])C(=O)C[C@@H](O)[C@@H](O)[C@H](O)[C@H](O)CO",
"smiles_sanitized":"O=C(O)C(=O)CC(O)C(O)C(O)C(O)CO"
}
},
{
"id":"cpd01006_C",
"name":"Aspirin",
"compartment":"C",
"annotation":{
"smiles":"CC(=O)Oc1ccccc1C(=O)[O-]",
"smiles_sanitized":"CC(=O)Oc1ccccc1C(=O)O"
}
},
{
"id":"cpd16658_C",
"name":"4-hydroxybenzylacetate",
"compartment":"C",
"annotation":{
"smiles":"",
"smiles_sanitized":""
}
},
{
"id":"cpd00689_C",
"name":"Porphobilinogen",
"compartment":"C",
"annotation":{
"smiles":"[NH3+]Cc1[nH]cc(CCC(=O)[O-])c1CC(=O)[O-]",
"smiles_sanitized":"NCc1[nH]cc(CCC(=O)O)c1CC(=O)O"
}
},
{
"id":"cpd35716_C",
"name":"3-(3,4-dihydroxyphenyl)propanoyl-CoA",
"compartment":"C",
"annotation":{
"smiles":"CC(C)(COP(=O)([O-])OP(=O)([O-])OC[C@H]1O[C@@H](n2cnc3c(N)ncnc32)[C@H](O)[C@@H]1OP(=O)([O-])[O-])[C@@H](O)C(=O)NCCC(=O)NCCSC(=O)CCc1ccc(O)c(O)c1",
"smiles_sanitized":"CC(C)(COP(=O)(O)OP(=O)(O)OCC1OC(n2cnc3c(N)ncnc32)C(O)C1OP(=O)(O)O)C(O)C(=O)NCCC(=O)NCCSC(=O)CCc1ccc(O)c(O)c1"
}
},
{
"id":"cpd00294_C",
"name":"dAMP",
"compartment":"C",
"annotation":{
"smiles":"Nc1ncnc2c1ncn2[C@H]1C[C@H](O)[C@@H](COP(=O)([O-])[O-])O1",
"smiles_sanitized":"Nc1ncnc2c1ncn2C1CC(O)C(COP(=O)(O)O)O1"
}
},
{
"id":"cpd00762_E",
"name":"4-Guanidinobutanoate",
"compartment":"E",
"annotation":{
"smiles":"NC(=[NH2+])NCCCC(=O)[O-]",
"smiles_sanitized":"N=C(N)NCCCC(=O)O"
}
},
{
"id":"cpd02160_C",
"name":"N2-Succinyl-L-ornithine",
"compartment":"C",
"annotation":{
"smiles":"[NH3+]CCC[C@H](NC(=O)CCC(=O)[O-])C(=O)[O-]",
"smiles_sanitized":"NCCCC(NC(=O)CCC(=O)O)C(=O)O"
}
},
{
"id":"cpd00153_C",
"name":"Benzoate",
"compartment":"C",
"annotation":{
"smiles":"O=C([O-])c1ccccc1",
"smiles_sanitized":"O=C(O)c1ccccc1"
}
},
{
"id":"cpd03642_E",
"name":"Adipate",
"compartment":"E",
"annotation":{
"smiles":"O=C([O-])CCCCC(=O)[O-]",
"smiles_sanitized":"O=C(O)CCCCC(=O)O"
}
},
{
"id":"cpd01217_C",
"name":"Xanthosine",
"compartment":"C",
"annotation":{
"smiles":"O=c1[nH]c(=O)c2ncn([C@@H]3O[C@H](CO)[C@@H](O)[C@H]3O)c2[nH]1",
"smiles_sanitized":"O=c1[nH]c(=O)c2ncn(C3OC(CO)C(O)C3O)c2[nH]1"
}
},
{
"id":"cpd00550_E",
"name":"L-Serine",
"compartment":"E",
"annotation":{
"smiles":"[NH3+][C@H](CO)C(=O)[O-]",
"smiles_sanitized":"NC(CO)C(=O)O"
}
},
{
"id":"cpd02932_C",
"name":"UDP-N-acetylmuramoyl-L-Ala-D-gamma-Glu-6-carboxy-L-Lys-(D-Ala)2",
"compartment":"C",
"annotation":{
"smiles":"CC(=O)N[C@H]1[C@@H](OP(=O)([O-])OP(=O)([O-])OC[C@H]2O[C@@H](n3ccc(=O)[nH]c3=O)[C@H](O)[C@@H]2O)O[C@H](CO)[C@@H](O)[C@@H]1O[C@H](C)C(=O)N[C@@H](C)C(=O)N[C@H](CCC(=O)N[C@@H](CCC[C@@H]([NH3+])C(=O)[O-])C(=O)N[C@H](C)C(=O)N[C@H](C)C(=O)[O-])C(=O)[O-]",
"smiles_sanitized":"CC(=O)NC1C(OP(=O)(O)OP(=O)(O)OCC2OC(n3ccc(=O)[nH]c3=O)C(O)C2O)OC(CO)C(O)C1OC(C)C(=O)NC(C)C(=O)NC(CCC(=O)NC(CCCC(N)C(=O)O)C(=O)NC(C)C(=O)NC(C)C(=O)O)C(=O)O"
}
},
{
"id":"cpd00021_C",
"name":"Fe",
"compartment":"C",
"annotation":{
"smiles":"[Fe]",
"smiles_sanitized":"[Fe]"
}
},
{
"id":"cpd00998_C",
"name":"Octane",
"compartment":"C",
"annotation":{
"smiles":"CCCCCCCC",
"smiles_sanitized":"CCCCCCCC"
}
},
{
"id":"cpd03494_C",
"name":"Undecaprenyl-diphospho-N-acetylmuramoyl-L-alanyl-D-glutamyl-meso-2-6-diaminopimeloyl-D-alanyl-D-alanine",
"compartment":"C",
"annotation":{
"smiles":"CC(=O)N[C@H]1[C@@H](OP(=O)([O-])OP(=O)([O-])OC/C=C(/C)CC/C=C(/C)CC/C=C(/C)CC/C=C(/C)CC/C=C(/C)CC/C=C(/C)CC/C=C(/C)CC/C=C(/C)CC/C=C(\\C)CC/C=C(\\C)CCC=C(C)C)O[C@H](CO)[C@@H](O)[C@@H]1O[C@H](C)C(=O)N[C@@H](C)C(=O)N[C@H](CCC(=O)N[C@@H](CCC[C@@H]([NH3+])C(=O)[O-])C(=O)N[C@H](C)C(=O)N[C@H](C)C(=O)[O-])C(=O)[O-]",
"smiles_sanitized":"CC(=O)NC1C(OP(=O)(O)OP(=O)(O)OCC=C(C)CCC=C(C)CCC=C(C)CCC=C(C)CCC=C(C)CCC=C(C)CCC=C(C)CCC=C(C)CCC=C(C)CCC=C(C)CCC=C(C)C)OC(CO)C(O)C1OC(C)C(=O)NC(C)C(=O)NC(CCC(=O)NC(CCCC(N)C(=O)O)C(=O)NC(C)C(=O)NC(C)C(=O)O)C(=O)O"
}
},
{
"id":"cpd02826_C",
"name":"5'-Phosphoribosylformylglycinamidine",
"compartment":"C",
"annotation":{
"smiles":"[NH2+]=C(CNC=O)N[C@@H]1O[C@H](COP(=O)([O-])[O-])[C@@H](O)[C@H]1O",
"smiles_sanitized":"N=C(CNC=O)NC1OC(COP(=O)(O)O)C(O)C1O"
}
},
{
"id":"cpd01646_C",
"name":"2-Isopropylmalate",
"compartment":"C",
"annotation":{
"smiles":"CC(C)[C@@](O)(CC(=O)[O-])C(=O)[O-]",
"smiles_sanitized":"CC(C)C(O)(CC(=O)O)C(=O)O"
}
},
{
"id":"cpd00421_C",
"name":"Triphosphate",
"compartment":"C",
"annotation":{
"smiles":"O=P([O-])([O-])OP(=O)([O-])OP(=O)([O-])O",
"smiles_sanitized":"O=P(O)(O)OP(=O)(O)OP(=O)(O)O"
}
},
{
"id":"cpd16648_C",
"name":"sorbate",
"compartment":"C",
"annotation":{
"smiles":"C/C=C/C=C/C(=O)[O-]",
"smiles_sanitized":"CC=CC=CC(=O)O"
}
},
{
"id":"3_OHACYL_ACP_LPS_C",
"name":"3-OHACYL-ACP_LPS",
"compartment":"C"
},
{
"id":"cpd00020_C",
"name":"Pyruvate",
"compartment":"C",
"annotation":{
"smiles":"CC(=O)C(=O)[O-]",
"smiles_sanitized":"CC(=O)C(=O)O"
}
},
{
"id":"cpd00274_C",
"name":"Citrulline",
"compartment":"C",
"annotation":{
"smiles":"NC(=O)NCCC[C@H]([NH3+])C(=O)[O-]",
"smiles_sanitized":"NC(=O)NCCCC(N)C(=O)O"
}
},
{
"id":"cpd00005_C",
"name":"NADPH",
"compartment":"C",
"annotation":{
"smiles":"NC(=O)C1=CN([C@@H]2O[C@H](COP(=O)([O-])OP(=O)([O-])OC[C@H]3O[C@@H](n4cnc5c(N)ncnc54)[C@H](OP(=O)([O-])[O-])[C@@H]3O)[C@@H](O)[C@H]2O)C=CC1",
"smiles_sanitized":"NC(=O)C1=CN(C2OC(COP(=O)(O)OP(=O)(O)OCC3OC(n4cnc5c(N)ncnc54)C(OP(=O)(O)O)C3O)C(O)C2O)C=CC1"
}
},
{
"id":"cpd00283_C",
"name":"Geranyldiphosphate",
"compartment":"C",
"annotation":{
"smiles":"CC(C)=CCC/C(C)=C/COP(=O)([O-])OP(=O)([O-])O",
"smiles_sanitized":"CC(C)=CCCC(C)=CCOP(=O)(O)OP(=O)(O)O"
}
},
{
"id":"cpd00931_C",
"name":"5-Amino-6--5-phosphoribosylaminouracil",
"compartment":"C",
"annotation":{
"smiles":"Nc1c(N[C@@H]2O[C@H](COP(=O)([O-])[O-])[C@@H](O)[C@H]2O)[nH]c(=O)[nH]c1=O",
"smiles_sanitized":"Nc1c(NC2OC(COP(=O)(O)O)C(O)C2O)[nH]c(=O)[nH]c1=O"
}
},
{
"id":"cpd01727_E",
"name":"Pimelate",
"compartment":"E",
"annotation":{
"smiles":"O=C([O-])CCCCCC(=O)[O-]",
"smiles_sanitized":"O=C(O)CCCCCC(=O)O"
}
},
{
"id":"cpd00013_E",
"name":"NH3",
"compartment":"E",
"annotation":{
"smiles":"[NH4+]",
"smiles_sanitized":"N"
}
},
{
"id":"cpd00763_C",
"name":"4-Maleylacetoacetate",
"compartment":"C",
"annotation":{
"smiles":"O=C([O-])/C=C\\C(=O)CC(=O)CC(=O)[O-]",
"smiles_sanitized":"O=C(O)C=CC(=O)CC(=O)CC(=O)O"
}
},
{
"id":"cpd01476_C",
"name":"Protoporphyrin",
"compartment":"C",
"annotation":{
"smiles":"C=CC1=C(C)c2cc3[nH]c(cc4nc(cc5[nH]c(cc1n2)c(C)c5CCC(=O)[O-])C(CCC(=O)[O-])=C4C)c(C)c3C=C",
"smiles_sanitized":"C=CC1=C(C)c2cc3[nH]c(cc4nc(cc5[nH]c(cc1n2)c(C)c5CCC(=O)O)C(CCC(=O)O)=C4C)c(C)c3C=C"
}
},
{
"id":"cpd01107_C",
"name":"Decanoate",
"compartment":"C",
"annotation":{
"smiles":"CCCCCCCCCC(=O)[O-]",
"smiles_sanitized":"CCCCCCCCCC(=O)O"
}
},
{
"id":"RNA_W_C",
"name":"RNA_W",
"compartment":"C"
},
{
"id":"cpd02654_C",
"name":"4-Methyl-5--2-phosphoethyl-thiazole",
"compartment":"C",
"annotation":{
"smiles":"Cc1ncsc1CCOP(=O)([O-])[O-]",
"smiles_sanitized":"Cc1ncsc1CCOP(=O)(O)O"
}
},
{
"id":"WAX_ESTER_C",
"name":"generic wax esters",
"compartment":"C"
},
{
"id":"cpd03586_C",
"name":"kdo2-lipid iva",
"compartment":"C",
"annotation":{
"smiles":"CCCCCCCCCCC[C@@H](O)CC(=O)N[C@H]1[C@@H](OP(=O)([O-])[O-])O[C@H](CO[C@@H]2O[C@H](CO[C@]3(C(=O)[O-])C[C@@H](O[C@]4(C(=O)[O-])C[C@@H](O)[C@@H](O)[C@@H]([C@H](O)CO)O4)[C@@H](O)[C@@H]([C@H](O)CO)O3)[C@@H](OP(=O)([O-])[O-])[C@H](OC(=O)C[C@H](O)CCCCCCCCCCC)[C@H]2NC(=O)C[C@H](O)CCCCCCCCCCC)[C@@H](O)[C@@H]1OC(=O)C[C@H](O)CCCCCCCCCCC",
"smiles_sanitized":"CCCCCCCCCCCC(O)CC(=O)NC1C(OP(=O)(O)O)OC(COC2OC(COC3(C(=O)O)CC(OC4(C(=O)O)CC(O)C(O)C(C(O)CO)O4)C(O)C(C(O)CO)O3)C(OP(=O)(O)O)C(OC(=O)CC(O)CCCCCCCCCCC)C2NC(=O)CC(O)CCCCCCCCCCC)C(O)C1OC(=O)CC(O)CCCCCCCCCCC"
}
},
{
"id":"C18_1_C9__3HYDROXYACP_C",
"name":"cis-3-hydroxyoctadec-9-enoyl-acp",
"compartment":"C"
},
{
"id":"cpd00096_C",
"name":"CDP",
"compartment":"C",
"annotation":{
"smiles":"Nc1ccn([C@@H]2O[C@H](COP(=O)([O-])OP(=O)([O-])O)[C@@H](O)[C@H]2O)c(=O)n1",
"smiles_sanitized":"Nc1ccn(C2OC(COP(=O)(O)OP(=O)(O)O)C(O)C2O)c(=O)n1"
}
},
{
"id":"cpd00379_C",
"name":"Glutarate",
"compartment":"C",
"annotation":{
"smiles":"O=C([O-])CCCC(=O)[O-]",
"smiles_sanitized":"O=C(O)CCCC(=O)O"
}
},
{
"id":"cpd03123_C",
"name":"3-Oxohexanoyl-CoA",
"compartment":"C",
"annotation":{
"smiles":"CCCC(=O)CC(=O)SCCNC(=O)CCNC(=O)[C@H](O)C(C)(C)COP(=O)([O-])OP(=O)([O-])OC[C@H]1O[C@@H](n2cnc3c(N)ncnc32)[C@H](O)[C@@H]1OP(=O)([O-])[O-]",
"smiles_sanitized":"CCCC(=O)CC(=O)SCCNC(=O)CCNC(=O)C(O)C(C)(C)COP(=O)(O)OP(=O)(O)OCC1OC(n2cnc3c(N)ncnc32)C(O)C1OP(=O)(O)O"
}
},
{
"id":"cpd00297_C",
"name":"dTDP",
"compartment":"C",
"annotation":{
"smiles":"Cc1cn([C@H]2C[C@H](O)[C@@H](COP(=O)([O-])OP(=O)([O-])O)O2)c(=O)[nH]c1=O",
"smiles_sanitized":"Cc1cn(C2CC(O)C(COP(=O)(O)OP(=O)(O)O)O2)c(=O)[nH]c1=O"
}
},
{
"id":"CHARGED_VAL_TRNAS_C",
"name":"charged valine trna",
"compartment":"C"
},
{
"id":"cpd02964_C",
"name":"UDP-N-acetylmuramoyl-L-alanyl-D-gamma-glutamyl-meso-2-6-diaminopimelate",
"compartment":"C",
"annotation":{
"smiles":"CC(=O)N[C@H]1[C@@H](OP(=O)([O-])OP(=O)([O-])OC[C@H]2O[C@@H](n3ccc(=O)[nH]c3=O)[C@H](O)[C@@H]2O)O[C@H](CO)[C@@H](O)[C@@H]1O[C@H](C)C(=O)N[C@@H](C)C(=O)N[C@H](CCC(=O)N[C@@H](CCC[C@@H]([NH3+])C(=O)[O-])C(=O)[O-])C(=O)[O-]",
"smiles_sanitized":"CC(=O)NC1C(OP(=O)(O)OP(=O)(O)OCC2OC(n3ccc(=O)[nH]c3=O)C(O)C2O)OC(CO)C(O)C1OC(C)C(=O)NC(C)C(=O)NC(CCC(=O)NC(CCCC(N)C(=O)O)C(=O)O)C(=O)O"
}
},
{
"id":"cpd00957_C",
"name":"2,5-Diamino-6-(5'-phosphoribosylamino)-4-pyrimidineone",
"compartment":"C",
"annotation":{
"smiles":"Nc1nc(N[C@@H]2O[C@H](COP(=O)([O-])[O-])[C@@H](O)[C@H]2O)c(N)c(=O)[nH]1",
"smiles_sanitized":"Nc1nc(NC2OC(COP(=O)(O)O)C(O)C2O)c(N)c(=O)[nH]1"
}
},
{
"id":"cpd00092_E",
"name":"Uracil",
"compartment":"E",
"annotation":{
"smiles":"O=c1cc[nH]c(=O)[nH]1",
"smiles_sanitized":"O=c1cc[nH]c(=O)[nH]1"
}
},
{
"id":"cpd00536_E",
"name":"Oleate",
"compartment":"E",
"annotation":{
"smiles":"CCCCCCCC/C=C\\CCCCCCCC(=O)[O-]",
"smiles_sanitized":"CCCCCCCCC=CCCCCCCCC(=O)O"
}
},
{
"id":"cpd02113_C",
"name":"dTDP-rhamnose",
"compartment":"C",
"annotation":{
"smiles":"Cc1cn([C@H]2C[C@H](O)[C@@H](COP(=O)([O-])OP(=O)([O-])O[C@H]3O[C@@H](C)[C@H](O)[C@@H](O)[C@H]3O)O2)c(=O)[nH]c1=O",
"smiles_sanitized":"Cc1cn(C2CC(O)C(COP(=O)(O)OP(=O)(O)OC3OC(C)C(O)C(O)C3O)O2)c(=O)[nH]c1=O"
}
},
{
"id":"cpd00032_C",
"name":"Oxaloacetate",
"compartment":"C",
"annotation":{
"smiles":"O=C([O-])CC(=O)C(=O)[O-]",
"smiles_sanitized":"O=C(O)CC(=O)C(=O)O"
}
},
{
"id":"cpd00320_C",
"name":"D-Aspartate",
"compartment":"C",
"annotation":{
"smiles":"[NH3+][C@H](CC(=O)[O-])C(=O)[O-]",
"smiles_sanitized":"NC(CC(=O)O)C(=O)O"
}
},
{
"id":"C4_0_2ENOYLACP_C",
"name":"trans-but-2-enoyl-acp",
"compartment":"C"
},
{
"id":"cpd00277_C",
"name":"Deoxyguanosine",
"compartment":"C",
"annotation":{
"smiles":"Nc1nc2c(ncn2[C@H]2C[C@H](O)[C@@H](CO)O2)c(=O)[nH]1",
"smiles_sanitized":"Nc1nc2c(ncn2C2CC(O)C(CO)O2)c(=O)[nH]1"
}
},
{
"id":"cpd00064_C",
"name":"Ornithine",
"compartment":"C",
"annotation":{
"smiles":"[NH3+]CCC[C@H]([NH3+])C(=O)[O-]",
"smiles_sanitized":"NCCCC(N)C(=O)O"
}
},
{
"id":"cpd00699_C",
"name":"5-Dehydroquinate",
"compartment":"C",
"annotation":{
"smiles":"O=C1C[C@@](O)(C(=O)[O-])C[C@@H](O)[C@@H]1O",
"smiles_sanitized":"O=C1CC(O)(C(=O)O)CC(O)C1O"
}
},
{
"id":"CYS_TRNAS_C",
"name":"cysteine trna",
"compartment":"C"
},
{
"id":"cpd01630_C",
"name":"cis,cis-Muconate",
"compartment":"C",
"annotation":{
"smiles":"O=C([O-])/C=C\\C=C/C(=O)[O-]",
"smiles_sanitized":"O=C(O)C=CC=CC(=O)O"
}
},
{
"id":"cpd03846_E",
"name":"octanoate",
"compartment":"E",
"annotation":{
"smiles":"CCCCCCCC(=O)[O-]",
"smiles_sanitized":"CCCCCCCC(=O)O"
}
},
{
"id":"cpd16644_C",
"name":"4-carboxy-3-oxobutanoyl-coa",
"compartment":"C",
"annotation":{
"smiles":"",
"smiles_sanitized":""
}
},
{
"id":"cpd01324_C",
"name":"L-Histidinal",
"compartment":"C",
"annotation":{
"smiles":"[NH3+][C@H](C=O)Cc1c[nH]cn1",
"smiles_sanitized":"NC(C=O)Cc1c[nH]cn1"
}
},
{
"id":"cpd00086_C",
"name":"Propionyl-CoA",
"compartment":"C",
"annotation":{
"smiles":"CCC(=O)SCCNC(=O)CCNC(=O)[C@H](O)C(C)(C)COP(=O)([O-])OP(=O)([O-])OC[C@H]1O[C@@H](n2cnc3c(N)ncnc32)[C@H](O)[C@@H]1OP(=O)([O-])[O-]",
"smiles_sanitized":"CCC(=O)SCCNC(=O)CCNC(=O)C(O)C(C)(C)COP(=O)(O)OP(=O)(O)OCC1OC(n2cnc3c(N)ncnc32)C(O)C1OP(=O)(O)O"
}
},
{
"id":"cpd00012_C",
"name":"PPi",
"compartment":"C",
"annotation":{
"smiles":"O=P([O-])([O-])OP(=O)([O-])O",
"smiles_sanitized":"O=P(O)(O)OP(=O)(O)O"
}
},
{
"id":"ACETOACETYL_S_ACP_C",
"name":"acetoacetyl-acp",
"compartment":"C"
},
{
"id":"cpd00102_C",
"name":"Glyceraldehyde3-phosphate",
"compartment":"C",
"annotation":{
"smiles":"O=C[C@H](O)COP(=O)([O-])[O-]",
"smiles_sanitized":"O=CC(O)COP(=O)(O)O"
}
},
{
"id":"cpd00218_C",
"name":"Niacin",
"compartment":"C",
"annotation":{
"smiles":"O=C([O-])c1cccnc1",
"smiles_sanitized":"O=C(O)c1cccnc1"
}
},
{
"id":"cpd04073_E",
"name":"Vanillate",
"compartment":"E",
"annotation":{
"smiles":"COc1cc(C(=O)[O-])ccc1O",
"smiles_sanitized":"COc1cc(C(=O)O)ccc1O"
}
},
{
"id":"cpd00129_E",
"name":"L-Proline",
"compartment":"E",
"annotation":{
"smiles":"O=C([O-])[C@@H]1CCC[NH2+]1",
"smiles_sanitized":"O=C(O)C1CCCN1"
}
},
{
"id":"cpd00797_E",
"name":"(R)-3-Hydroxybutanoate",
"compartment":"E",
"annotation":{
"smiles":"C[C@@H](O)CC(=O)[O-]",
"smiles_sanitized":"CC(O)CC(=O)O"
}
},
{
"id":"cpd02698_C",
"name":"N-Succinyl-L-2,6-diaminopimelate",
"compartment":"C",
"annotation":{
"smiles":"[NH3+][C@@H](CCC[C@H](NC(=O)CCC(=O)[O-])C(=O)[O-])C(=O)[O-]",
"smiles_sanitized":"NC(CCCC(NC(=O)CCC(=O)O)C(=O)O)C(=O)O"
}
},
{
"id":"cpd00015_C",
"name":"FAD",
"compartment":"C",
"annotation":{
"smiles":"Cc1cc2nc3c(=O)[n-]c(=O)nc-3n(C[C@H](O)[C@H](O)[C@H](O)COP(=O)([O-])OP(=O)([O-])OC[C@H]3O[C@@H](n4cnc5c(N)ncnc54)[C@H](O)[C@@H]3O)c2cc1C",
"smiles_sanitized":"Cc1cc2nc3c(=O)[nH]c(=O)nc-3n(CC(O)C(O)C(O)COP(=O)(O)OP(=O)(O)OCC3OC(n4cnc5c(N)ncnc54)C(O)C3O)c2cc1C"
}
},
{
"id":"AFATTYACID_WAX_C",
"name":"AFATTYACID_WAX",
"compartment":"C"
},
{
"id":"cpd00041_C",
"name":"L-Aspartate",
"compartment":"C",
"annotation":{
"smiles":"[NH3+][C@@H](CC(=O)[O-])C(=O)[O-]",
"smiles_sanitized":"NC(CC(=O)O)C(=O)O"
}
},
{
"id":"C14_0_ACP_C",
"name":"tetradecanoyl-acp",
"compartment":"C"
},
{
"id":"cpd00790_C",
"name":"O-Acetyl-L-homoserine",
"compartment":"C",
"annotation":{
"smiles":"CC(=O)OCC[C@H]([NH3+])C(=O)[O-]",
"smiles_sanitized":"CC(=O)OCCC(N)C(=O)O"
}
},
{
"id":"cpd00137_E",
"name":"Citrate",
"compartment":"E",
"annotation":{
"smiles":"O=C([O-])CC(O)(CC(=O)[O-])C(=O)[O-]",
"smiles_sanitized":"O=C(O)CC(O)(CC(=O)O)C(=O)O"
}
},
{
"id":"cpd00021_E",
"name":"Fe",
"compartment":"E",
"annotation":{
"smiles":"[Fe]",
"smiles_sanitized":"[Fe]"
}
},
{
"id":"cpd03846_C",
"name":"octanoate",
"compartment":"C",
"annotation":{
"smiles":"CCCCCCCC(=O)[O-]",
"smiles_sanitized":"CCCCCCCC(=O)O"
}
},
{
"id":"cpd02152_C",
"name":"L-Argininosuccinate",
"compartment":"C",
"annotation":{
"smiles":"[NH2+]=C(NCCC[C@H]([NH3+])C(=O)[O-])NC(CC(=O)[O-])C(=O)[O-]",
"smiles_sanitized":"N=C(NCCCC(N)C(=O)O)NC(CC(=O)O)C(=O)O"
}
},
{
"id":"cpd00081_C",
"name":"Sulfite",
"compartment":"C",
"annotation":{
"smiles":"O=S([O-])O",
"smiles_sanitized":"O=S(O)O"
}
},
{
"id":"cpd00219_C",
"name":"Prephenate",
"compartment":"C",
"annotation":{
"smiles":"O=C([O-])C(=O)C[C@]1(C(=O)[O-])C=C[C@@H](O)C=C1",
"smiles_sanitized":"O=C(O)C(=O)CC1(C(=O)O)C=CC(O)C=C1"
}
},
{
"id":"cpd00093_E",
"name":"Anthranilate",
"compartment":"E",
"annotation":{
"smiles":"Nc1ccccc1C(=O)[O-]",
"smiles_sanitized":"Nc1ccccc1C(=O)O"
}
},
{
"id":"C10_0_3HYDROXYACP_C",
"name":"3-hydroxydecanoyl-acp",
"compartment":"C"
},
{
"id":"CHARGED_LEU_TRNAS_C",
"name":"charged leucine trna",
"compartment":"C"
},
{
"id":"cpd11422_C",
"name":"1,2-diacyl-sn-glycerol 3-phosphate",
"compartment":"C",
"annotation":{
"smiles":"*C(=O)OCC(COP(=O)([O-])[O-])OC(*)=O",
"smiles_sanitized":"*C(=O)OCC(COP(=O)(O)O)OC(*)=O"
}
},
{
"id":"cpd03115_C",
"name":"(S)-3-Hydroxytetradecanoyl-CoA",
"compartment":"C",
"annotation":{
"smiles":"CCCCCCCCCCC[C@H](O)CC(=O)SCCNC(=O)CCNC(=O)[C@H](O)C(C)(C)COP(=O)([O-])OP(=O)([O-])OC[C@H]1O[C@@H](n2cnc3c(N)ncnc32)[C@H](O)[C@@H]1OP(=O)([O-])[O-]",
"smiles_sanitized":"CCCCCCCCCCCC(O)CC(=O)SCCNC(=O)CCNC(=O)C(O)C(C)(C)COP(=O)(O)OP(=O)(O)OCC1OC(n2cnc3c(N)ncnc32)C(O)C1OP(=O)(O)O"
}
},
{
"id":"POLYSACCH_FREE_W_C",
"name":"POLYSACCH_FREE_W",
"compartment":"C"
},
{
"id":"cpd02820_C",
"name":"UDP-N-acetylglucosamine enolpyruvate",
"compartment":"C",
"annotation":{
"smiles":"C=C(O[C@H]1[C@H](O)[C@@H](CO)O[C@H](OP(=O)([O-])OP(=O)([O-])OC[C@H]2O[C@@H](n3ccc(=O)[nH]c3=O)[C@H](O)[C@@H]2O)[C@@H]1NC(C)=O)C(=O)[O-]",
"smiles_sanitized":"C=C(OC1C(O)C(CO)OC(OP(=O)(O)OP(=O)(O)OCC2OC(n3ccc(=O)[nH]c3=O)C(O)C2O)C1NC(C)=O)C(=O)O"
}
},
{
"id":"cpd11193_C",
"name":"benzylacetate",
"compartment":"C",
"annotation":{
"smiles":"CC(=O)OCc1ccccc1",
"smiles_sanitized":"CC(=O)OCc1ccccc1"
}
},
{
"id":"cpd01335_C",
"name":"Octanoyl-CoA",
"compartment":"C",
"annotation":{
"smiles":"CCCCCCCC(=O)SCCNC(=O)CCNC(=O)[C@H](O)C(C)(C)COP(=O)([O-])OP(=O)([O-])OC[C@H]1O[C@@H](n2cnc3c(N)ncnc32)[C@H](O)[C@@H]1OP(=O)([O-])[O-]",
"smiles_sanitized":"CCCCCCCC(=O)SCCNC(=O)CCNC(=O)C(O)C(C)(C)COP(=O)(O)OP(=O)(O)OCC1OC(n2cnc3c(N)ncnc32)C(O)C1OP(=O)(O)O"
}
},
{
"id":"cpd00066_E",
"name":"L-Phenylalanine",
"compartment":"E",
"annotation":{
"smiles":"[NH3+][C@@H](Cc1ccccc1)C(=O)[O-]",
"smiles_sanitized":"NC(Cc1ccccc1)C(=O)O"
}
},
{
"id":"C18_0_2ENOYLACP_C",
"name":"trans-octadec-2-enoyl-acp",
"compartment":"C"
},
{
"id":"cpd00210_E",
"name":"Taurine",
"compartment":"E",
"annotation":{
"smiles":"[NH3+]CCS(=O)(=O)[O-]",
"smiles_sanitized":"NCCS(=O)(=O)O"
}
},
{
"id":"cpd00858_C",
"name":"L-Glutamate5-semialdehyde",
"compartment":"C",
"annotation":{
"smiles":"[NH3+][C@@H](CCC=O)C(=O)[O-]",
"smiles_sanitized":"NC(CCC=O)C(=O)O"
}
},
{
"id":"cpd02566_C",
"name":"2,3-Dihydroxy-2,3-dihydrobenzoate",
"compartment":"C",
"annotation":{
"smiles":"O=C([O-])C1=CC=C[C@H](O)[C@H]1O",
"smiles_sanitized":"O=C(O)C1=CC=CC(O)C1O"
}
},
{
"id":"cpd00051_C",
"name":"L-Arginine",
"compartment":"C",
"annotation":{
"smiles":"NC(=[NH2+])NCCC[C@H]([NH3+])C(=O)[O-]",
"smiles_sanitized":"N=C(N)NCCCC(N)C(=O)O"
}
},
{
"id":"cpd15378_E",
"name":"4-Hydroxy-benzylalcohol",
"compartment":"E",
"annotation":{
"smiles":"OCc1ccc(O)cc1",
"smiles_sanitized":"OCc1ccc(O)cc1"
}
},
{
"id":"cpd00177_C",
"name":"dADP",
"compartment":"C",
"annotation":{
"smiles":"Nc1ncnc2c1ncn2[C@H]1C[C@H](O)[C@@H](COP(=O)([O-])OP(=O)([O-])O)O1",
"smiles_sanitized":"Nc1ncnc2c1ncn2C1CC(O)C(COP(=O)(O)OP(=O)(O)O)O1"
}
},
{
"id":"cpd00066_C",
"name":"L-Phenylalanine",
"compartment":"C",
"annotation":{
"smiles":"[NH3+][C@@H](Cc1ccccc1)C(=O)[O-]",
"smiles_sanitized":"NC(Cc1ccccc1)C(=O)O"
}
},
{
"id":"cpd00047_C",
"name":"Formate",
"compartment":"C",
"annotation":{
"smiles":"O=C[O-]",
"smiles_sanitized":"O=CO"
}
},
{
"id":"GLY_TRNAS_C",
"name":"glycine trna",
"compartment":"C"
},
{
"id":"cpd00003_C",
"name":"NAD",
"compartment":"C",
"annotation":{
"smiles":"NC(=O)c1ccc[n+]([C@@H]2O[C@H](COP(=O)([O-])OP(=O)([O-])OC[C@H]3O[C@@H](n4cnc5c(N)ncnc54)[C@H](O)[C@@H]3O)[C@@H](O)[C@H]2O)c1",
"smiles_sanitized":"NC(=O)c1ccc[n+](C2OC(COP(=O)(O)OP(=O)(O)OCC3OC(n4cnc5c(N)ncnc54)C(O)C3O)C(O)C2O)c1"
}
},
{
"id":"cpd00522_C",
"name":"dTDP-4-oxo-L-rhamnose",
"compartment":"C",
"annotation":{
"smiles":"Cc1cn([C@H]2C[C@H](O)[C@@H](COP(=O)([O-])OP(=O)([O-])O[C@H]3O[C@@H](C)C(=O)[C@@H](O)[C@H]3O)O2)c(=O)[nH]c1=O",
"smiles_sanitized":"Cc1cn(C2CC(O)C(COP(=O)(O)OP(=O)(O)OC3OC(C)C(=O)C(O)C3O)O2)c(=O)[nH]c1=O"
}
},
{
"id":"cpd16648_E",
"name":"sorbate",
"compartment":"E",
"annotation":{
"smiles":"C/C=C/C=C/C(=O)[O-]",
"smiles_sanitized":"CC=CC=CC(=O)O"
}
},
{
"id":"cpd00322_C",
"name":"L-Isoleucine",
"compartment":"C",
"annotation":{
"smiles":"CC[C@H](C)[C@H]([NH3+])C(=O)[O-]",
"smiles_sanitized":"CCC(C)C(N)C(=O)O"
}
},
{
"id":"cpd00100_C",
"name":"Glycerol",
"compartment":"C",
"annotation":{
"smiles":"OCC(O)CO",
"smiles_sanitized":"OCC(O)CO"
}
},
{
"id":"cpd09844_C",
"name":"(3S)-3-Hydroxyadipyl-CoA",
"compartment":"C",
"annotation":{
"smiles":"CC(C)(COP(=O)([O-])OP(=O)([O-])OC[C@H]1O[C@@H](n2cnc3c(N)ncnc32)[C@H](O)[C@@H]1OP(=O)([O-])[O-])[C@@H](O)C(=O)NCCC(=O)NCCSC(=O)C[C@@H](O)CCC(=O)[O-]",
"smiles_sanitized":"CC(C)(COP(=O)(O)OP(=O)(O)OCC1OC(n2cnc3c(N)ncnc32)C(O)C1OP(=O)(O)O)C(O)C(=O)NCCC(=O)NCCSC(=O)CC(O)CCC(=O)O"
}
},
{
"id":"cpd01716_P",
"name":"3-Dehydroshikimate",
"compartment":"P",
"annotation":{
"smiles":"O=C([O-])C1=CC(=O)[C@@H](O)[C@H](O)C1",
"smiles_sanitized":"O=C(O)C1=CC(=O)C(O)C(O)C1"
}
},
{
"id":"cpd02666_C",
"name":"(R)-4'-Phosphopantothenoyl-L-cysteine",
"compartment":"C",
"annotation":{
"smiles":"CC(C)(COP(=O)([O-])[O-])[C@@H](O)C(=O)NCCC(=O)N[C@@H](CS)C(=O)[O-]",
"smiles_sanitized":"CC(C)(COP(=O)(O)O)C(O)C(=O)NCCC(=O)NC(CS)C(=O)O"
}
},
{
"id":"cpd16658_E",
"name":"4-hydroxybenzylacetate",
"compartment":"E",
"annotation":{
"smiles":"",
"smiles_sanitized":""
}
},
{
"id":"cpd02060_C",
"name":"(2E)-Dodecenoyl-CoA",
"compartment":"C",
"annotation":{
"smiles":"CCCCCCCCC/C=C/C(=O)SCCNC(=O)CCNC(=O)C(O)C(C)(C)COP(=O)([O-])OP(=O)([O-])OC[C@H]1OC(n2cnc3c(N)ncnc32)[C@H](O)[C@@H]1OP(=O)([O-])[O-]",
"smiles_sanitized":"CCCCCCCCCC=CC(=O)SCCNC(=O)CCNC(=O)C(O)C(C)(C)COP(=O)(O)OP(=O)(O)OCC1OC(n2cnc3c(N)ncnc32)C(O)C1OP(=O)(O)O"
}
},
{
"id":"cpd02147_C",
"name":"4-Phosphoerythronate",
"compartment":"C",
"annotation":{
"smiles":"O=C([O-])[C@H](O)[C@H](O)COP(=O)([O-])[O-]",
"smiles_sanitized":"O=C(O)C(O)C(O)COP(=O)(O)O"
}
},
{
"id":"CHARGED_PHE_TRNAS_C",
"name":"charged phenylalanine trna",
"compartment":"C"
},
{
"id":"cpd00039_C",
"name":"L-Lysine",
"compartment":"C",
"annotation":{
"smiles":"[NH3+]CCCC[C@H]([NH3+])C(=O)[O-]",
"smiles_sanitized":"NCCCCC(N)C(=O)O"
}
},
{
"id":"cpd00242_C",
"name":"H2CO3",
"compartment":"C",
"annotation":{
"smiles":"O=C([O-])O",
"smiles_sanitized":"O=C(O)O"
}
},
{
"id":"cpd00782_C",
"name":"Pimeloyl-CoA",
"compartment":"C",
"annotation":{
"smiles":"CC(C)(COP(=O)([O-])OP(=O)([O-])OC[C@H]1O[C@@H](n2cnc3c(N)ncnc32)[C@H](O)[C@@H]1OP(=O)([O-])[O-])[C@@H](O)C(=O)NCCC(=O)NCCSC(=O)CCCCCC(=O)[O-]",
"smiles_sanitized":"CC(C)(COP(=O)(O)OP(=O)(O)OCC1OC(n2cnc3c(N)ncnc32)C(O)C1OP(=O)(O)O)C(O)C(=O)NCCC(=O)NCCSC(=O)CCCCCC(=O)O"
}
},
{
"id":"cpd03122_C",
"name":"(S)-Hydroxyhexanoyl-CoA",
"compartment":"C",
"annotation":{
"smiles":"CCC[C@H](O)CC(=O)SCCNC(=O)CCNC(=O)C(O)C(C)(C)COP(=O)([O-])OP(=O)([O-])OC[C@H]1O[C@@H](n2cnc3c(N)ncnc32)[C@H](O)[C@@H]1OP(=O)([O-])[O-]",
"smiles_sanitized":"CCCC(O)CC(=O)SCCNC(=O)CCNC(=O)C(O)C(C)(C)COP(=O)(O)OP(=O)(O)OCC1OC(n2cnc3c(N)ncnc32)C(O)C1OP(=O)(O)O"
}
},
{
"id":"cpd11193_E",
"name":"benzylacetate",
"compartment":"E",
"annotation":{
"smiles":"CC(=O)OCc1ccccc1",
"smiles_sanitized":"CC(=O)OCc1ccccc1"
}
},
{
"id":"cpd00532_C",
"name":"O2-",
"compartment":"C",
"annotation":{
"smiles":"[O][O-]",
"smiles_sanitized":"[O]O"
}
},
{
"id":"cpd02394_C",
"name":"GAR",
"compartment":"C",
"annotation":{
"smiles":"[NH3+]CC(=O)N[C@@H]1O[C@H](COP(=O)([O-])[O-])[C@@H](O)[C@H]1O",
"smiles_sanitized":"NCC(=O)NC1OC(COP(=O)(O)O)C(O)C1O"
}
},
{
"id":"cpd00060_E",
"name":"L-Methionine",
"compartment":"E",
"annotation":{
"smiles":"CSCC[C@H]([NH3+])C(=O)[O-]",
"smiles_sanitized":"CSCCC(N)C(=O)O"
}
},
{
"id":"cpd01034_C",
"name":"Toluol",
"compartment":"C",
"annotation":{
"smiles":"Cc1ccccc1",
"smiles_sanitized":"Cc1ccccc1"
}
},
{
"id":"cpd00016_C",
"name":"Pyridoxal phosphate",
"compartment":"C",
"annotation":{
"smiles":"Cc1ncc(COP(=O)([O-])[O-])c(C=O)c1O",
"smiles_sanitized":"Cc1ncc(COP(=O)(O)O)c(C=O)c1O"
}
},
{
"id":"cpd00330_C",
"name":"Dihydrofolate",
"compartment":"C",
"annotation":{
"smiles":"Nc1nc2c(c(=O)[nH]1)N=C(CNc1ccc(C(=O)N[C@@H](CCC(=O)[O-])C(=O)[O-])cc1)CN2",
"smiles_sanitized":"Nc1nc2c(c(=O)[nH]1)N=C(CNc1ccc(C(=O)NC(CCC(=O)O)C(=O)O)cc1)CN2"
}
},
{
"id":"cpd00540_C",
"name":"BET",
"compartment":"C",
"annotation":{
"smiles":"C[N+](C)(C)CC(=O)[O-]",
"smiles_sanitized":"C[N+](C)(C)CC(=O)O"
}
},
{
"id":"cpd03447_C",
"name":"2-Octaprenyl-6-methoxy-1,4-benzoquinone",
"compartment":"C",
"annotation":{
"smiles":"COC1=CC(=O)C=C(C/C=C(\\C)CC/C=C(\\C)CC/C=C(\\C)CC/C=C(\\C)CC/C=C(\\C)CC/C=C(\\C)CC/C=C(\\C)CCC=C(C)C)C1=O",
"smiles_sanitized":"COC1=CC(=O)C=C(CC=C(C)CCC=C(C)CCC=C(C)CCC=C(C)CCC=C(C)CCC=C(C)CCC=C(C)CCC=C(C)C)C1=O"
}
},
{
"id":"cpd00604_E",
"name":"4-Coumarate",
"compartment":"E",
"annotation":{
"smiles":"O=C([O-])/C=C/c1ccc(O)cc1",
"smiles_sanitized":"O=C(O)C=Cc1ccc(O)cc1"
}
},
{
"id":"cpd00011_C",
"name":"CO2",
"compartment":"C",
"annotation":{
"smiles":"O=C=O",
"smiles_sanitized":"O=C=O"
}
},
{
"id":"cpd00338_C",
"name":"5-Aminolevulinate",
"compartment":"C",
"annotation":{
"smiles":"[NH3+]CC(=O)CCC(=O)[O-]",
"smiles_sanitized":"NCC(=O)CCC(=O)O"
}
},
{
"id":"cpd00073_E",
"name":"Urea",
"compartment":"E",
"annotation":{
"smiles":"NC(N)=O",
"smiles_sanitized":"NC(N)=O"
}
},
{
"id":"cpd09842_C",
"name":"Adipyl-CoA",
"compartment":"C",
"annotation":{
"smiles":"CC(C)(COP(=O)([O-])OP(=O)([O-])OC[C@H]1O[C@@H](n2cnc3c(N)ncnc32)[C@H](O)[C@@H]1OP(=O)([O-])[O-])[C@@H](O)C(=O)NCCC(=O)NCCSC(=O)CCCCC(=O)[O-]",
"smiles_sanitized":"CC(C)(COP(=O)(O)OP(=O)(O)OCC1OC(n2cnc3c(N)ncnc32)C(O)C1OP(=O)(O)O)C(O)C(=O)NCCC(=O)NCCSC(=O)CCCCC(=O)O"
}
},
{
"id":"C18_1_C9__2ENOYLACP_C",
"name":"trans,cis-octadec-2,9-enoyl-acp",
"compartment":"C"
},
{
"id":"cpd00438_E",
"name":"Deoxyadenosine",
"compartment":"E",
"annotation":{
"smiles":"Nc1ncnc2c1ncn2[C@H]1C[C@H](O)[C@@H](CO)O1",
"smiles_sanitized":"Nc1ncnc2c1ncn2C1CC(O)C(CO)O1"
}
},
{
"id":"cpd00971_C",
"name":"Na+",
"compartment":"C",
"annotation":{
"smiles":"[Na+]",
"smiles_sanitized":"[Na+]"
}
},
{
"id":"cpd16626_C",
"name":"cis-hexadec-7-enal",
"compartment":"C",
"annotation":{
"smiles":"",
"smiles_sanitized":""
}
},
{
"id":"cpd00361_C",
"name":"ACTN",
"compartment":"C",
"annotation":{
"smiles":"CC(=O)[C@H](C)O",
"smiles_sanitized":"CC(=O)C(C)O"
}
},
{
"id":"cpd16651_C",
"name":"7-carboxy-3-oxoheptanoyl-coa",
"compartment":"C",
"annotation":{
"smiles":"",
"smiles_sanitized":""
}
},
{
"id":"cpd00435_E",
"name":"Phenylcarbinol",
"compartment":"E",
"annotation":{
"smiles":"OCc1ccccc1",
"smiles_sanitized":"OCc1ccccc1"
}
},
{
"id":"cpd00010_C",
"name":"CoA",
"compartment":"C",
"annotation":{
"smiles":"CC(C)(COP(=O)([O-])OP(=O)([O-])OC[C@H]1O[C@@H](n2cnc3c(N)ncnc32)[C@H](O)[C@@H]1OP(=O)([O-])[O-])[C@@H](O)C(=O)NCCC(=O)NCCS",
"smiles_sanitized":"CC(C)(COP(=O)(O)OP(=O)(O)OCC1OC(n2cnc3c(N)ncnc32)C(O)C1OP(=O)(O)O)C(O)C(=O)NCCC(=O)NCCS"
}
},
{
"id":"C16_0_ALDEHYDE_C",
"name":"C16:0-ALDEHYDE",
"compartment":"C"
},
{
"id":"cpd00307_C",
"name":"Cytosine",
"compartment":"C",
"annotation":{
"smiles":"Nc1cc[nH]c(=O)n1",
"smiles_sanitized":"Nc1cc[nH]c(=O)n1"
}
},
{
"id":"cpd34075_C",
"name":"(S)-3-hydroxy-(9Z)-octadecenoyl-CoA",
"compartment":"C",
"annotation":{
"smiles":"CCCCCCCC/C=C\\CCCCC[C@H](O)CC(=O)SCCNC(=O)CCNC(=O)[C@H](O)C(C)(C)COP(=O)([O-])OP(=O)([O-])OC[C@H]1O[C@@H](n2cnc3c(N)ncnc32)[C@H](O)[C@@H]1OP(=O)([O-])[O-]",
"smiles_sanitized":"CCCCCCCCC=CCCCCCC(O)CC(=O)SCCNC(=O)CCNC(=O)C(O)C(C)(C)COP(=O)(O)OP(=O)(O)OCC1OC(n2cnc3c(N)ncnc32)C(O)C1OP(=O)(O)O"
}
},
{
"id":"cpd03584_C",
"name":"UDP-3-O-(beta-hydroxymyristoyl)-D-glucosamine",
"compartment":"C",
"annotation":{
"smiles":"CCCCCCCCCCC[C@@H](O)CC(=O)O[C@H]1[C@H](O)[C@@H](CO)O[C@H](OP(=O)([O-])OP(=O)([O-])OC[C@H]2O[C@@H](n3ccc(=O)[nH]c3=O)[C@H](O)[C@@H]2O)[C@@H]1[NH3+]",
"smiles_sanitized":"CCCCCCCCCCCC(O)CC(=O)OC1C(N)C(OP(=O)(O)OP(=O)(O)OCC2OC(n3ccc(=O)[nH]c3=O)C(O)C2O)OC(CO)C1O"
}
},
{
"id":"cpd00774_C",
"name":"UroporphyrinogenIII",
"compartment":"C",
"annotation":{
"smiles":"O=C([O-])CCc1c2[nH]c(c1CC(=O)[O-])Cc1[nH]c(c(CC(=O)[O-])c1CCC(=O)[O-])Cc1[nH]c(c(CC(=O)[O-])c1CCC(=O)[O-])Cc1[nH]c(c(CCC(=O)[O-])c1CC(=O)[O-])C2",
"smiles_sanitized":"O=C(O)CCc1c2[nH]c(c1CC(=O)O)Cc1[nH]c(c(CC(=O)O)c1CCC(=O)O)Cc1[nH]c(c(CC(=O)O)c1CCC(=O)O)Cc1[nH]c(c(CCC(=O)O)c1CC(=O)O)C2"
}
},
{
"id":"cpd00918_C",
"name":"2-Acetamido-5-oxopentanoate",
"compartment":"C",
"annotation":{
"smiles":"CC(=O)N[C@@H](CCC=O)C(=O)[O-]",
"smiles_sanitized":"CC(=O)NC(CCC=O)C(=O)O"
}
},
{
"id":"cpd03119_C",
"name":"3-Oxodecanoyl-CoA",
"compartment":"C",
"annotation":{
"smiles":"CCCCCCCC(=O)CC(=O)SCCNC(=O)CCNC(=O)[C@H](O)C(C)(C)COP(=O)([O-])OP(=O)([O-])OC[C@H]1O[C@@H](n2cnc3c(N)ncnc32)[C@H](O)[C@@H]1OP(=O)([O-])[O-]",
"smiles_sanitized":"CCCCCCCC(=O)CC(=O)SCCNC(=O)CCNC(=O)C(O)C(C)(C)COP(=O)(O)OP(=O)(O)OCC1OC(n2cnc3c(N)ncnc32)C(O)C1OP(=O)(O)O"
}
},
{
"id":"cpd00655_C",
"name":"Dephospho-CoA",
"compartment":"C",
"annotation":{
"smiles":"CC(C)(COP(=O)([O-])OP(=O)([O-])OC[C@H]1O[C@@H](n2cnc3c(N)ncnc32)[C@H](O)[C@@H]1O)[C@@H](O)C(=O)NCCC(=O)NCCS",
"smiles_sanitized":"CC(C)(COP(=O)(O)OP(=O)(O)OCC1OC(n2cnc3c(N)ncnc32)C(O)C1O)C(O)C(=O)NCCC(=O)NCCS"
}
},
{
"id":"cpd00239_C",
"name":"H2S",
"compartment":"C",
"annotation":{
"smiles":"[SH-]",
"smiles_sanitized":"S"
}
},
{
"id":"cpd00028_C",
"name":"Heme",
"compartment":"C",
"annotation":{
"smiles":"C=CC1=C(C)C2=N/C1=C\\c1c(C)c(CCC(=O)[O-])c3n1[Fe]n1/c(c(C)c(C=C)/c1=C/2)=C\\C1=N/C(=C\\3)C(CCC(=O)[O-])=C1C",
"smiles_sanitized":"C=CC1=C(C)C2=NC1=Cc1c(C)c(CCC(=O)O)c3n1[Fe]n1c(c(C)c(C=C)c1=C2)=CC1=NC(=C3)C(CCC(=O)O)=C1C"
}
},
{
"id":"CHARGED_GLY_TRNAS_C",
"name":"charged glycine trna",
"compartment":"C"
},
{
"id":"cpd00485_C",
"name":"D-Mannose1-phosphate",
"compartment":"C",
"annotation":{
"smiles":"O=P([O-])([O-])O[C@H]1O[C@H](CO)[C@@H](O)[C@H](O)[C@@H]1O",
"smiles_sanitized":"O=P(O)(O)OC1OC(CO)C(O)C(O)C1O"
}
},
{
"id":"CHARGED_ILE_TRNAS_C",
"name":"charged isoleucine trna",
"compartment":"C"
},
{
"id":"cpd00035_C",
"name":"L-Alanine",
"compartment":"C",
"annotation":{
"smiles":"C[C@H]([NH3+])C(=O)[O-]",
"smiles_sanitized":"CC(N)C(=O)O"
}
},
{
"id":"cpd03606_C",
"name":"2-Oxo-3-hydroxy-4-phosphobutanoate",
"compartment":"C",
"annotation":{
"smiles":"O=C([O-])C(=O)[C@H](O)COP(=O)([O-])[O-]",
"smiles_sanitized":"O=C(O)C(=O)C(O)COP(=O)(O)O"
}
},
{
"id":"cpd00435_C",
"name":"Phenylcarbinol",
"compartment":"C",
"annotation":{
"smiles":"OCc1ccccc1",
"smiles_sanitized":"OCc1ccccc1"
}
},
{
"id":"cpd00085_C",
"name":"beta-Alanine",
"compartment":"C",
"annotation":{
"smiles":"[NH3+]CCC(=O)[O-]",
"smiles_sanitized":"NCCC(=O)O"
}
},
{
"id":"cpd02882_C",
"name":"4--1-D-Ribitylamino-5-aminouracil",
"compartment":"C",
"annotation":{
"smiles":"Nc1c(NC[C@H](O)[C@H](O)[C@H](O)CO)[nH]c(=O)[nH]c1=O",
"smiles_sanitized":"Nc1c(NCC(O)C(O)C(O)CO)[nH]c(=O)[nH]c1=O"
}
},
{
"id":"ASPARTYL_TRNAASN_C",
"name":"l-aspartyl-trna(asn)",
"compartment":"C"
},
{
"id":"cpd03585_C",
"name":"KDO-lipid IV(A)",
"compartment":"C",
"annotation":{
"smiles":"CCCCCCCCCCC[C@@H](O)CC(=O)N[C@H]1[C@@H](OP(=O)([O-])[O-])O[C@H](CO[C@@H]2O[C@H](CO[C@]3(C(=O)[O-])C[C@@H](O)[C@@H](O)[C@@H]([C@H](O)CO)O3)[C@@H](OP(=O)([O-])[O-])[C@H](OC(=O)C[C@H](O)CCCCCCCCCCC)[C@H]2NC(=O)C[C@H](O)CCCCCCCCCCC)[C@@H](O)[C@@H]1OC(=O)C[C@H](O)CCCCCCCCCCC",
"smiles_sanitized":"CCCCCCCCCCCC(O)CC(=O)NC1C(OP(=O)(O)O)OC(COC2OC(COC3(C(=O)O)CC(O)C(O)C(C(O)CO)O3)C(OP(=O)(O)O)C(OC(=O)CC(O)CCCCCCCCCCC)C2NC(=O)CC(O)CCCCCCCCCCC)C(O)C1OC(=O)CC(O)CCCCCCCCCCC"
}
},
{
"id":"cpd04104_C",
"name":"3-Ketopimelyl-CoA",
"compartment":"C",
"annotation":{
"smiles":"CC(C)(COP(=O)([O-])OP(=O)([O-])OC[C@H]1O[C@@H](n2cnc3c(N)ncnc32)[C@H](O)[C@@H]1OP(=O)([O-])[O-])[C@@H](O)C(=O)NCCC(=O)NCCSC(=O)CC(=O)CCCC(=O)[O-]",
"smiles_sanitized":"CC(C)(COP(=O)(O)OP(=O)(O)OCC1OC(n2cnc3c(N)ncnc32)C(O)C1OP(=O)(O)O)C(O)C(=O)NCCC(=O)NCCSC(=O)CC(=O)CCCC(=O)O"
}
},
{
"id":"cpd02345_C",
"name":"L-Glutamate1-semialdehyde",
"compartment":"C",
"annotation":{
"smiles":"[NH3+][C@H](C=O)CCC(=O)[O-]",
"smiles_sanitized":"NC(C=O)CCC(=O)O"
}
},
{
"id":"cpd02920_C",
"name":"2-Amino-4-hydroxy-6-hydroxymethyl-7-8-dihydropteridinediphosphate",
"compartment":"C",
"annotation":{
"smiles":"Nc1nc2c(c(=O)[nH]1)N=C(COP(=O)([O-])OP(=O)([O-])[O-])CN2",
"smiles_sanitized":"Nc1nc2c(c(=O)[nH]1)N=C(COP(=O)(O)OP(=O)(O)O)CN2"
}
},
{
"id":"cpd01006_E",
"name":"Aspirin",
"compartment":"E",
"annotation":{
"smiles":"CC(=O)Oc1ccccc1C(=O)[O-]",
"smiles_sanitized":"CC(=O)Oc1ccccc1C(=O)O"
}
},
{
"id":"cpd00699_P",
"name":"5-Dehydroquinate",
"compartment":"P",
"annotation":{
"smiles":"O=C1C[C@@](O)(C(=O)[O-])C[C@@H](O)[C@@H]1O",
"smiles_sanitized":"O=C1CC(O)(C(=O)O)CC(O)C1O"
}
},
{
"id":"THR_TRNAS_C",
"name":"threonine trna",
"compartment":"C"
},
{
"id":"cpd00018_C",
"name":"AMP",
"compartment":"C",
"annotation":{
"smiles":"Nc1ncnc2c1ncn2[C@@H]1O[C@H](COP(=O)([O-])[O-])[C@@H](O)[C@H]1O",
"smiles_sanitized":"Nc1ncnc2c1ncn2C1OC(COP(=O)(O)O)C(O)C1O"
}
},
{
"id":"cpd00238_C",
"name":"Sedoheptulose7-phosphate",
"compartment":"C",
"annotation":{
"smiles":"O=C(CO)[C@@H](O)[C@H](O)[C@H](O)[C@H](O)COP(=O)([O-])[O-]",
"smiles_sanitized":"O=C(CO)C(O)C(O)C(O)C(O)COP(=O)(O)O"
}
},
{
"id":"cpd00247_C",
"name":"Orotate",
"compartment":"C",
"annotation":{
"smiles":"O=C([O-])c1cc(=O)[nH]c(=O)[nH]1",
"smiles_sanitized":"O=C(O)c1cc(=O)[nH]c(=O)[nH]1"
}
},
{
"id":"cpd02557_C",
"name":"Farnesylfarnesylgeraniol",
"compartment":"C",
"annotation":{
"smiles":"CC(C)=CCC/C(C)=C/CC/C(C)=C/CC/C(C)=C/CC/C(C)=C/CC/C(C)=C/CC/C(C)=C/CC/C(C)=C/COP(=O)([O-])OP(=O)([O-])O",
"smiles_sanitized":"CC(C)=CCCC(C)=CCCC(C)=CCCC(C)=CCCC(C)=CCCC(C)=CCCC(C)=CCCC(C)=CCOP(=O)(O)OP(=O)(O)O"
}
},
{
"id":"cpd05193_E",
"name":"suberate",
"compartment":"E",
"annotation":{
"smiles":"O=C([O-])CCCCCCC(=O)[O-]",
"smiles_sanitized":"O=C(O)CCCCCCC(=O)O"
}
},
{
"id":"cpd03587_C",
"name":"kdo2-lipid a",
"compartment":"C",
"annotation":{
"smiles":"CCCCCCCCCCCCCC(=O)O[C@H](CCCCCCCCCCC)CC(=O)O[C@H]1[C@H](OP(=O)([O-])[O-])[C@@H](CO[C@]2(C(=O)[O-])C[C@@H](O[C@]3(C(=O)[O-])C[C@@H](O)[C@@H](O)[C@@H]([C@H](O)CO)O3)[C@@H](O)[C@@H]([C@H](O)CO)O2)O[C@@H](OC[C@H]2O[C@H](OP(=O)([O-])[O-])[C@H](NC(=O)C[C@H](O)CCCCCCCCCCC)[C@@H](OC(=O)C[C@H](O)CCCCCCCCCCC)[C@@H]2O)[C@@H]1NC(=O)C[C@@H](CCCCCCCCCCC)OC(=O)CCCCCCCCCCC",
"smiles_sanitized":"CCCCCCCCCCCCCC(=O)OC(CCCCCCCCCCC)CC(=O)OC1C(NC(=O)CC(CCCCCCCCCCC)OC(=O)CCCCCCCCCCC)C(OCC2OC(OP(=O)(O)O)C(NC(=O)CC(O)CCCCCCCCCCC)C(OC(=O)CC(O)CCCCCCCCCCC)C2O)OC(COC2(C(=O)O)CC(OC3(C(=O)O)CC(O)C(O)C(C(O)CO)O3)C(O)C(C(O)CO)O2)C1OP(=O)(O)O"
}
},
{
"id":"cpd00613_C",
"name":"Hexadecanol",
"compartment":"C",
"annotation":{
"smiles":"CCCCCCCCCCCCCCCCO",
"smiles_sanitized":"CCCCCCCCCCCCCCCCO"
}
},
{
"id":"ACYL_COA_TAG_C",
"name":"ACYL-COA_TAG",
"compartment":"C"
},
{
"id":"cpd02255_C",
"name":"3-oxoadipate-enol-lactone",
"compartment":"C",
"annotation":{
"smiles":"O=C([O-])CC1=CCC(=O)O1",
"smiles_sanitized":"O=C(O)CC1=CCC(=O)O1"
}
},
{
"id":"CHARGED_GLT_TRNAS_C",
"name":"charged glutamate trna",
"compartment":"C"
},
{
"id":"cpd00587_C",
"name":"D-Cysteine",
"compartment":"C",
"annotation":{
"smiles":"[NH3+][C@H](CS)C(=O)[O-]",
"smiles_sanitized":"NC(CS)C(=O)O"
}
},
{
"id":"cpd03279_C",
"name":"Deoxyinosine",
"compartment":"C",
"annotation":{
"smiles":"O=c1[nH]cnc2c1ncn2[C@H]1C[C@H](O)[C@@H](CO)O1",
"smiles_sanitized":"O=c1[nH]cnc2c1ncn2C1CC(O)C(CO)O1"
}
},
{
"id":"cpd11310_C",
"name":"5-phosphoribosyl-5-carboxyaminoimidazole",
"compartment":"C",
"annotation":{
"smiles":"O=C([O-])Nc1cncn1[C@@H]1O[C@H](COP(=O)([O-])[O-])[C@@H](O)[C@H]1O",
"smiles_sanitized":"O=C(O)Nc1cncn1C1OC(COP(=O)(O)O)C(O)C1O"
}
},
{
"id":"cpd01844_C",
"name":"Propionyl phosphate",
"compartment":"C",
"annotation":{
"smiles":"CCC(=O)OP(=O)([O-])[O-]",
"smiles_sanitized":"CCC(=O)OP(=O)(O)O"
}
},
{
"id":"cpd00065_C",
"name":"L-Tryptophan",
"compartment":"C",
"annotation":{
"smiles":"[NH3+][C@@H](Cc1c[nH]c2ccccc12)C(=O)[O-]",
"smiles_sanitized":"NC(Cc1c[nH]c2ccccc12)C(=O)O"
}
},
{
"id":"cpd12689_C",
"name":"3-Oxotetradecanoyl-CoA",
"compartment":"C",
"annotation":{
"smiles":"CCCCCCCCCCCC(=O)CC(=O)SCCNC(=O)CCNC(=O)[C@H](O)C(C)(C)COP(=O)([O-])OP(=O)([O-])OC[C@H]1O[C@@H](n2cnc3c(N)ncnc32)[C@H](O)[C@@H]1OP(=O)([O-])[O-]",
"smiles_sanitized":"CCCCCCCCCCCC(=O)CC(=O)SCCNC(=O)CCNC(=O)C(O)C(C)(C)COP(=O)(O)OP(=O)(O)OCC1OC(n2cnc3c(N)ncnc32)C(O)C1OP(=O)(O)O"
}
},
{
"id":"MALONYL_ACP_C",
"name":"malonyl-acp",
"compartment":"C"
},
{
"id":"cpd00309_C",
"name":"XAN",
"compartment":"C",
"annotation":{
"smiles":"O=c1[nH]c(=O)c2[nH]cnc2[nH]1",
"smiles_sanitized":"O=c1[nH]c(=O)c2[nH]cnc2[nH]1"
}
},
{
"id":"cpd00287_C",
"name":"3-Oxo-2-methylpropanoate",
"compartment":"C",
"annotation":{
"smiles":"CC(C=O)C(=O)[O-]",
"smiles_sanitized":"CC(C=O)C(=O)O"
}
},
{
"id":"cpd00498_C",
"name":"2-Aceto-2-hydroxybutanoate",
"compartment":"C",
"annotation":{
"smiles":"CC[C@](O)(C(C)=O)C(=O)[O-]",
"smiles_sanitized":"CCC(O)(C(C)=O)C(=O)O"
}
},
{
"id":"cpd16643_C",
"name":"4-carboxy-3-hydroxybutanoyl-coa",
"compartment":"C",
"annotation":{
"smiles":"",
"smiles_sanitized":""
}
},
{
"id":"cpd00478_C",
"name":"Pyridoxine phosphate",
"compartment":"C",
"annotation":{
"smiles":"Cc1ncc(COP(=O)([O-])[O-])c(CO)c1O",
"smiles_sanitized":"Cc1ncc(COP(=O)(O)O)c(CO)c1O"
}
},
{
"id":"cpd00053_E",
"name":"L-Glutamine",
"compartment":"E",
"annotation":{
"smiles":"NC(=O)CC[C@H]([NH3+])C(=O)[O-]",
"smiles_sanitized":"NC(=O)CCC(N)C(=O)O"
}
},
{
"id":"cpd00056_C",
"name":"TPP",
"compartment":"C",
"annotation":{
"smiles":"Cc1ncc(C[n+]2csc(CCOP(=O)([O-])OP(=O)([O-])O)c2C)c(N)n1",
"smiles_sanitized":"Cc1ncc(C[n+]2csc(CCOP(=O)(O)OP(=O)(O)O)c2C)c(N)n1"
}
},
{
"id":"NICOTINATE_NUCLEOTIDE_C",
"name":"NICOTINATE_NUCLEOTIDE",
"compartment":"C"
},
{
"id":"cpd00035_E",
"name":"L-Alanine",
"compartment":"E",
"annotation":{
"smiles":"C[C@H]([NH3+])C(=O)[O-]",
"smiles_sanitized":"CC(N)C(=O)O"
}
},
{
"id":"cpd00098_C",
"name":"Choline",
"compartment":"C",
"annotation":{
"smiles":"C[N+](C)(C)CCO",
"smiles_sanitized":"C[N+](C)(C)CCO"
}
},
{
"id":"CHARGED_TRP_TRNAS_C",
"name":"charged tryptophan trna",
"compartment":"C"
},
{
"id":"cpd00036_C",
"name":"Succinate",
"compartment":"C",
"annotation":{
"smiles":"O=C([O-])CCC(=O)[O-]",
"smiles_sanitized":"O=C(O)CCC(=O)O"
}
},
{
"id":"cpd00299_C",
"name":"dUMP",
"compartment":"C",
"annotation":{
"smiles":"O=c1ccn([C@H]2C[C@H](O)[C@@H](COP(=O)([O-])[O-])O2)c(=O)[nH]1",
"smiles_sanitized":"O=c1ccn(C2CC(O)C(COP(=O)(O)O)O2)c(=O)[nH]1"
}
},
{
"id":"cpd23921_C",
"name":"beta-ketovaleryl-coa",
"compartment":"C",
"annotation":{
"smiles":"CCC(=O)CC(=O)SCCNC(=O)CCNC(=O)[C@H](O)C(C)(C)COP(=O)([O-])OP(=O)([O-])OC[C@H]1O[C@@H](n2cnc3c(N)ncnc32)[C@H](O)[C@@H]1OP(=O)([O-])[O-]",
"smiles_sanitized":"CCC(=O)CC(=O)SCCNC(=O)CCNC(=O)C(O)C(C)(C)COP(=O)(O)OP(=O)(O)OCC1OC(n2cnc3c(N)ncnc32)C(O)C1OP(=O)(O)O"
}
},
{
"id":"cpd00093_C",
"name":"Anthranilate",
"compartment":"C",
"annotation":{
"smiles":"Nc1ccccc1C(=O)[O-]",
"smiles_sanitized":"Nc1ccccc1C(=O)O"
}
},
{
"id":"cpd00448_C",
"name":"D-Glyceraldehyde",
"compartment":"C",
"annotation":{
"smiles":"O=C[C@H](O)CO",
"smiles_sanitized":"O=CC(O)CO"
}
},
{
"id":"cpd00197_P",
"name":"Protocatechuate",
"compartment":"P",
"annotation":{
"smiles":"O=C([O-])c1ccc(O)c(O)c1",
"smiles_sanitized":"O=C(O)c1ccc(O)c(O)c1"
}
},
{
"id":"cpd00200_C",
"name":"4MOP",
"compartment":"C",
"annotation":{
"smiles":"CC(C)CC(=O)C(=O)[O-]",
"smiles_sanitized":"CC(C)CC(=O)C(=O)O"
}
},
{
"id":"C16_1_C7__3HYDROXYACP_C",
"name":"cis-3-hydroxyhexadec-7-enoyl-acp",
"compartment":"C"
},
{
"id":"ACP_C",
"name":"acyl carrier protein",
"compartment":"C"
},
{
"id":"cpd00856_C",
"name":"3-Carboxy-cis,cis-muconate",
"compartment":"C",
"annotation":{
"smiles":"O=C([O-])/C=C\\C(=C/C(=O)[O-])C(=O)[O-]",
"smiles_sanitized":"O=C(O)C=CC(=CC(=O)O)C(=O)O"
}
},
{
"id":"cpd01433_C",
"name":"2-Nitropropane",
"compartment":"C",
"annotation":{
"smiles":"CC(C)[N+](=O)[O-]",
"smiles_sanitized":"CC(C)[N+](=O)[O-]"
}
},
{
"id":"cpd00260_C",
"name":"Isocitrate",
"compartment":"C",
"annotation":{
"smiles":"O=C([O-])CC(C(=O)[O-])C(O)C(=O)[O-]",
"smiles_sanitized":"O=C(O)CC(C(=O)O)C(O)C(=O)O"
}
},
{
"id":"cpd00346_C",
"name":"L-Aspartate4-semialdehyde",
"compartment":"C",
"annotation":{
"smiles":"[NH3+][C@@H](CC=O)C(=O)[O-]",
"smiles_sanitized":"NC(CC=O)C(=O)O"
}
},
{
"id":"C10_0_ACP_C",
"name":"decanoyl-acp",
"compartment":"C"
},
{
"id":"cpd00523_C",
"name":"Trehalose 6-phosphate",
"compartment":"C",
"annotation":{
"smiles":"O=P([O-])([O-])OC[C@H]1O[C@H](O[C@H]2O[C@H](CO)[C@@H](O)[C@H](O)[C@H]2O)[C@H](O)[C@@H](O)[C@@H]1O",
"smiles_sanitized":"O=P(O)(O)OCC1OC(OC2OC(CO)C(O)C(O)C2O)C(O)C(O)C1O"
}
},
{
"id":"cpd00125_C",
"name":"5-10-Methylenetetrahydrofolate",
"compartment":"C",
"annotation":{
"smiles":"Nc1nc2c(c(=O)[nH]1)N1CN(c3ccc(C(=O)N[C@@H](CCC(=O)[O-])C(=O)[O-])cc3)C[C@H]1CN2",
"smiles_sanitized":"Nc1nc2c(c(=O)[nH]1)N1CN(c3ccc(C(=O)NC(CCC(=O)O)C(=O)O)cc3)CC1CN2"
}
},
{
"id":"cpd02333_C",
"name":"Quinolinate",
"compartment":"C",
"annotation":{
"smiles":"O=C([O-])c1cccnc1C(=O)[O-]",
"smiles_sanitized":"O=C(O)c1cccnc1C(=O)O"
}
},
{
"id":"3_OHMYRISTOYL_ACP_C",
"name":"3-hydroxytetradecanoyl-acp",
"compartment":"C"
},
{
"id":"cpd00199_C",
"name":"4-Oxobutanoate",
"compartment":"C",
"annotation":{
"smiles":"O=CCCC(=O)[O-]",
"smiles_sanitized":"O=CCCC(=O)O"
}
},
{
"id":"cpd28230_C",
"name":"trans-2,3-dehydroadipyl-coA",
"compartment":"C",
"annotation":{
"smiles":"CC(C)(COP(=O)([O-])OP(=O)([O-])OC[C@H]1O[C@@H](n2cnc3c(N)ncnc32)[C@H](O)[C@@H]1OP(=O)([O-])[O-])[C@@H](O)C(=O)NCCC(=O)NCCSC(=O)/C=C/CCC(=O)[O-]",
"smiles_sanitized":"CC(C)(COP(=O)(O)OP(=O)(O)OCC1OC(n2cnc3c(N)ncnc32)C(O)C1OP(=O)(O)O)C(O)C(=O)NCCC(=O)NCCSC(=O)C=CCCC(=O)O"
}
},
{
"id":"cpd16652_C",
"name":"7-carboxyheptanoyl-coa",
"compartment":"C",
"annotation":{
"smiles":"",
"smiles_sanitized":""
}
},
{
"id":"cpd03443_C",
"name":"3-Octaprenyl-4-hydroxybenzoate",
"compartment":"C",
"annotation":{
"smiles":"CC(C)=CCC/C(C)=C/CC/C(C)=C/CC/C(C)=C/CC/C(C)=C/CC/C(C)=C/CC/C(C)=C/CC/C(C)=C/Cc1cc(C(=O)[O-])ccc1O",
"smiles_sanitized":"CC(C)=CCCC(C)=CCCC(C)=CCCC(C)=CCCC(C)=CCCC(C)=CCCC(C)=CCCC(C)=CCc1cc(C(=O)O)ccc1O"
}
},
{
"id":"cpd00650_C",
"name":"Crotonyl-CoA",
"compartment":"C",
"annotation":{
"smiles":"C/C=C/C(=O)SCCNC(=O)CCNC(=O)C(O)C(C)(C)COP(=O)([O-])OP(=O)([O-])OC[C@H]1O[C@@H](n2cnc3c(N)ncnc32)[C@H](O)[C@@H]1OP(=O)([O-])[O-]",
"smiles_sanitized":"CC=CC(=O)SCCNC(=O)CCNC(=O)C(O)C(C)(C)COP(=O)(O)OP(=O)(O)OCC1OC(n2cnc3c(N)ncnc32)C(O)C1OP(=O)(O)O"
}
},
{
"id":"cpd00107_C",
"name":"L-Leucine",
"compartment":"C",
"annotation":{
"smiles":"CC(C)C[C@H]([NH3+])C(=O)[O-]",
"smiles_sanitized":"CC(C)CC(N)C(=O)O"
}
},
{
"id":"cpd00039_E",
"name":"L-Lysine",
"compartment":"E",
"annotation":{
"smiles":"[NH3+]CCCC[C@H]([NH3+])C(=O)[O-]",
"smiles_sanitized":"NCCCCC(N)C(=O)O"
}
},
{
"id":"cpd00551_C",
"name":"DTYL",
"compartment":"C",
"annotation":{
"smiles":"CC(=O)C(C)=O",
"smiles_sanitized":"CC(=O)C(C)=O"
}
},
{
"id":"ASP_TRNAS_C",
"name":"aspartate trna",
"compartment":"C"
},
{
"id":"cpd00027_E",
"name":"D-Glucose",
"compartment":"E",
"annotation":{
"smiles":"OC[C@H]1OC(O)[C@H](O)[C@@H](O)[C@@H]1O",
"smiles_sanitized":"OCC1OC(O)C(O)C(O)C1O"
}
},
{
"id":"CHARGED_SER_TRNAS_C",
"name":"charged serine trna",
"compartment":"C"
},
{
"id":"cpd00159_E",
"name":"L-Lactate",
"compartment":"E",
"annotation":{
"smiles":"C[C@H](O)C(=O)[O-]",
"smiles_sanitized":"CC(O)C(=O)O"
}
},
{
"id":"cpd02370_C",
"name":"D-Galactosamine 1-phosphate",
"compartment":"C",
"annotation":{
"smiles":"[NH3+][C@H]1C(OP(=O)([O-])[O-])O[C@H](CO)[C@H](O)[C@@H]1O",
"smiles_sanitized":"NC1C(OP(=O)(O)O)OC(CO)C(O)C1O"
}
},
{
"id":"cpd00008_C",
"name":"ADP",
"compartment":"C",
"annotation":{
"smiles":"Nc1ncnc2c1ncn2[C@@H]1O[C@H](COP(=O)([O-])OP(=O)([O-])O)[C@@H](O)[C@H]1O",
"smiles_sanitized":"Nc1ncnc2c1ncn2C1OC(COP(=O)(O)OP(=O)(O)O)C(O)C1O"
}
},
{
"id":"cpd02555_C",
"name":"Tetrahydropteroyltri-L-glutamate",
"compartment":"C",
"annotation":{
"smiles":"Nc1nc2c(c(=O)[nH]1)N[C@@H](CNc1ccc(C(=O)N[C@@H](CCC(=O)N[C@@H](CCC(=O)N[C@@H](CCC(=O)[O-])C(=O)[O-])C(=O)[O-])C(=O)[O-])cc1)CN2",
"smiles_sanitized":"Nc1nc2c(c(=O)[nH]1)NC(CNc1ccc(C(=O)NC(CCC(=O)NC(CCC(=O)NC(CCC(=O)O)C(=O)O)C(=O)O)C(=O)O)cc1)CN2"
}
},
{
"id":"cpd02738_C",
"name":"5-Methyltetrahydropteroyltri-L-glutamate",
"compartment":"C",
"annotation":{
"smiles":"CN1c2c(nc(N)[nH]c2=O)NC[C@@H]1CNc1ccc(C(=O)N[C@@H](CCC(=O)N[C@@H](CCC(=O)N[C@@H](CCC(=O)[O-])C(=O)[O-])C(=O)[O-])C(=O)[O-])cc1",
"smiles_sanitized":"CN1c2c(nc(N)[nH]c2=O)NCC1CNc1ccc(C(=O)NC(CCC(=O)NC(CCC(=O)NC(CCC(=O)O)C(=O)O)C(=O)O)C(=O)O)cc1"
}
},
{
"id":"cpd00360_E",
"name":"Purine",
"compartment":"E",
"annotation":{
"smiles":"c1ncc2[nH]cnc2n1",
"smiles_sanitized":"c1ncc2[nH]cnc2n1"
}
},
{
"id":"MET_TRNAS_C",
"name":"methionine trna",
"compartment":"C"
},
{
"id":"cpd00047_E",
"name":"Formate",
"compartment":"E",
"annotation":{
"smiles":"O=C[O-]",
"smiles_sanitized":"O=CO"
}
},
{
"id":"cpd00024_C",
"name":"2-Oxoglutarate",
"compartment":"C",
"annotation":{
"smiles":"O=C([O-])CCC(=O)C(=O)[O-]",
"smiles_sanitized":"O=C(O)CCC(=O)C(=O)O"
}
},
{
"id":"L_1_PHOSPHATIDYL_GLYCEROL_C",
"name":"generic l-1-phosphatidyl-glycerol",
"compartment":"C"
},
{
"id":"cpd03114_C",
"name":"3-Oxopalmitoyl-CoA",
"compartment":"C",
"annotation":{
"smiles":"CCCCCCCCCCCCCC(=O)CC(=O)SCCNC(=O)CCNC(=O)[C@H](O)C(C)(C)COP(=O)([O-])OP(=O)([O-])OC[C@H]1O[C@@H](n2cnc3c(N)ncnc32)[C@H](O)[C@@H]1OP(=O)([O-])[O-]",
"smiles_sanitized":"CCCCCCCCCCCCCC(=O)CC(=O)SCCNC(=O)CCNC(=O)C(O)C(C)(C)COP(=O)(O)OP(=O)(O)OCC1OC(n2cnc3c(N)ncnc32)C(O)C1OP(=O)(O)O"
}
},
{
"id":"cpd00235_C",
"name":"D-mannose-6-phosphate",
"compartment":"C",
"annotation":{
"smiles":"O=P([O-])([O-])OC[C@H]1OC(O)[C@@H](O)[C@@H](O)[C@@H]1O",
"smiles_sanitized":"O=P(O)(O)OCC1OC(O)C(O)C(O)C1O"
}
},
{
"id":"cpd00126_C",
"name":"GMP",
"compartment":"C",
"annotation":{
"smiles":"Nc1nc2c(ncn2[C@@H]2O[C@H](COP(=O)([O-])[O-])[C@@H](O)[C@H]2O)c(=O)[nH]1",
"smiles_sanitized":"Nc1nc2c(ncn2C2OC(COP(=O)(O)O)C(O)C2O)c(=O)[nH]1"
}
},
{
"id":"cpd03847_E",
"name":"Myristic acid",
"compartment":"E",
"annotation":{
"smiles":"CCCCCCCCCCCCCC(=O)[O-]",
"smiles_sanitized":"CCCCCCCCCCCCCC(=O)O"
}
},
{
"id":"cpd00480_C",
"name":"2-Dehydro-D-gluconate",
"compartment":"C",
"annotation":{
"smiles":"O=C([O-])C(=O)[C@@H](O)[C@H](O)[C@H](O)CO",
"smiles_sanitized":"O=C(O)C(=O)C(O)C(O)C(O)CO"
}
},
{
"id":"ILE_TRNAS_C",
"name":"isoleucine trna",
"compartment":"C"
},
{
"id":"cpd00031_C",
"name":"GDP",
"compartment":"C",
"annotation":{
"smiles":"Nc1nc2c(ncn2[C@@H]2O[C@H](COP(=O)([O-])OP(=O)([O-])O)[C@@H](O)[C@H]2O)c(=O)[nH]1",
"smiles_sanitized":"Nc1nc2c(ncn2C2OC(COP(=O)(O)OP(=O)(O)O)C(O)C2O)c(=O)[nH]1"
}
},
{
"id":"cpd00412_C",
"name":"Deoxyuridine",
"compartment":"C",
"annotation":{
"smiles":"O=c1ccn([C@H]2C[C@H](O)[C@@H](CO)O2)c(=O)[nH]1",
"smiles_sanitized":"O=c1ccn(C2CC(O)C(CO)O2)c(=O)[nH]1"
}
},
{
"id":"AFATTYACID_FREE_W_C",
"name":"AFATTYACID_FREE_W",
"compartment":"C"
},
{
"id":"C16_0_3HYDROXYACP_C",
"name":"3-hydroxyhexadecanoyl-acp",
"compartment":"C"
},
{
"id":"cpd03448_C",
"name":"2-Octaprenyl-3-methyl-6-methoxy-1,4-benzoquinone",
"compartment":"C",
"annotation":{
"smiles":"COC1=CC(=O)C(C)=C(C/C=C(\\C)CC/C=C(\\C)CC/C=C(\\C)CC/C=C(\\C)CC/C=C(\\C)CC/C=C(\\C)CC/C=C(\\C)CCC=C(C)C)C1=O",
"smiles_sanitized":"COC1=CC(=O)C(C)=C(CC=C(C)CCC=C(C)CCC=C(C)CCC=C(C)CCC=C(C)CCC=C(C)CCC=C(C)CCC=C(C)C)C1=O"
}
},
{
"id":"cpd00081_E",
"name":"Sulfite",
"compartment":"E",
"annotation":{
"smiles":"O=S([O-])O",
"smiles_sanitized":"O=S(O)O"
}
},
{
"id":"cpd02083_C",
"name":"CoproporphyrinogenIII",
"compartment":"C",
"annotation":{
"smiles":"Cc1c2[nH]c(c1CCC(=O)[O-])Cc1[nH]c(c(CCC(=O)[O-])c1C)Cc1[nH]c(c(CCC(=O)[O-])c1C)Cc1[nH]c(c(C)c1CCC(=O)[O-])C2",
"smiles_sanitized":"Cc1c2[nH]c(c1CCC(=O)O)Cc1[nH]c(c(CCC(=O)O)c1C)Cc1[nH]c(c(CCC(=O)O)c1C)Cc1[nH]c(c(C)c1CCC(=O)O)C2"
}
},
{
"id":"C6_C",
"name":"undecaprenyl-pyrophosphoryl-murnac-(pentapeptide)-n-acetylglucosamine",
"compartment":"C"
},
{
"id":"cpd00064_E",
"name":"Ornithine",
"compartment":"E",
"annotation":{
"smiles":"[NH3+]CCC[C@H]([NH3+])C(=O)[O-]",
"smiles_sanitized":"NCCCC(N)C(=O)O"
}
},
{
"id":"cpd00060_C",
"name":"L-Methionine",
"compartment":"C",
"annotation":{
"smiles":"CSCC[C@H]([NH3+])C(=O)[O-]",
"smiles_sanitized":"CSCCC(N)C(=O)O"
}
},
{
"id":"cpd00843_C",
"name":"Tartronate semialdehyde",
"compartment":"C",
"annotation":{
"smiles":"O=CC(O)C(=O)[O-]",
"smiles_sanitized":"O=CC(O)C(=O)O"
}
},
{
"id":"WAX_ESTER_W_C",
"name":"WAX-ESTER_W",
"compartment":"C"
},
{
"id":"C16_1_C7__2ENOYLACP_C",
"name":"trans,cis-hexadec-2,7-enoyl-acp",
"compartment":"C"
},
{
"id":"LYS_TRNAS_C",
"name":"lysine trna",
"compartment":"C"
},
{
"id":"cpd03847_C",
"name":"Myristic acid",
"compartment":"C",
"annotation":{
"smiles":"CCCCCCCCCCCCCC(=O)[O-]",
"smiles_sanitized":"CCCCCCCCCCCCCC(=O)O"
}
},
{
"id":"cpd00042_C",
"name":"GSH",
"compartment":"C",
"annotation":{
"smiles":"[NH3+][C@@H](CCC(=O)N[C@@H](CS)C(=O)NCC(=O)[O-])C(=O)[O-]",
"smiles_sanitized":"NC(CCC(=O)NC(CS)C(=O)NCC(=O)O)C(=O)O"
}
},
{
"id":"cpd00022_C",
"name":"Acetyl-CoA",
"compartment":"C",
"annotation":{
"smiles":"CC(=O)SCCNC(=O)CCNC(=O)[C@H](O)C(C)(C)COP(=O)([O-])OP(=O)([O-])OC[C@H]1O[C@@H](n2cnc3c(N)ncnc32)[C@H](O)[C@@H]1OP(=O)([O-])[O-]",
"smiles_sanitized":"CC(=O)SCCNC(=O)CCNC(=O)C(O)C(C)(C)COP(=O)(O)OP(=O)(O)OCC1OC(n2cnc3c(N)ncnc32)C(O)C1OP(=O)(O)O"
}
},
{
"id":"C6_0_3OXOACP_C",
"name":"3-oxoxyhexanoyl-acp",
"compartment":"C"
},
{
"id":"cpd02857_C",
"name":"DAHP",
"compartment":"C",
"annotation":{
"smiles":"O=C([O-])C(=O)C[C@@H](O)[C@H](O)[C@H](O)COP(=O)([O-])[O-]",
"smiles_sanitized":"O=C(O)C(=O)CC(O)C(O)C(O)COP(=O)(O)O"
}
},
{
"id":"C10_0_2ENOYLACP_C",
"name":"trans-dec-2-enoyl-acp",
"compartment":"C"
},
{
"id":"cpd00438_C",
"name":"Deoxyadenosine",
"compartment":"C",
"annotation":{
"smiles":"Nc1ncnc2c1ncn2[C@H]1C[C@H](O)[C@@H](CO)O1",
"smiles_sanitized":"Nc1ncnc2c1ncn2C1CC(O)C(CO)O1"
}
},
{
"id":"DNA_W_C",
"name":"DNA_W",
"compartment":"C"
},
{
"id":"cpd01733_E",
"name":"Ferulaldehyde",
"compartment":"E",
"annotation":{
"smiles":"COc1cc(/C=C/C=O)ccc1O",
"smiles_sanitized":"COc1cc(C=CC=O)ccc1O"
}
},
{
"id":"GLN_TRNAS_C",
"name":"glutamine trna",
"compartment":"C"
},
{
"id":"cpd00209_E",
"name":"Nitrate",
"compartment":"E",
"annotation":{
"smiles":"O=[N+]([O-])[O-]",
"smiles_sanitized":"O=[N+]([O-])[O-]"
}
},
{
"id":"cpd00658_C",
"name":"Isochorismate",
"compartment":"C",
"annotation":{
"smiles":"C=C(O[C@H]1C=CC=C(C(=O)[O-])[C@@H]1O)C(=O)[O-]",
"smiles_sanitized":"C=C(OC1C=CC=C(C(=O)O)C1O)C(=O)O"
}
},
{
"id":"cpd03127_C",
"name":"(2E)-Tetradecenoyl-CoA",
"compartment":"C",
"annotation":{
"smiles":"CCCCCCCCCCC/C=C/C(=O)SCCNC(=O)CCNC(=O)[C@H](O)C(C)(C)COP(=O)([O-])OP(=O)([O-])OC[C@H]1O[C@@H](n2cnc3c(N)ncnc32)[C@H](O)[C@@H]1OP(=O)([O-])[O-]",
"smiles_sanitized":"CCCCCCCCCCCC=CC(=O)SCCNC(=O)CCNC(=O)C(O)C(C)(C)COP(=O)(O)OP(=O)(O)OCC1OC(n2cnc3c(N)ncnc32)C(O)C1OP(=O)(O)O"
}
},
{
"id":"cpd16646_C",
"name":"trans-3-hydroxyhex-4-enoyl-coa",
"compartment":"C",
"annotation":{
"smiles":"",
"smiles_sanitized":""
}
},
{
"id":"cpd00137_C",
"name":"Citrate",
"compartment":"C",
"annotation":{
"smiles":"O=C([O-])CC(O)(CC(=O)[O-])C(=O)[O-]",
"smiles_sanitized":"O=C(O)CC(O)(CC(=O)O)C(=O)O"
}
},
{
"id":"cpd34461_E",
"name":"(7Z)-hexadec-7-enoate",
"compartment":"E",
"annotation":{
"smiles":"CCCCCCCC/C=C\\CCCCCC(=O)[O-]",
"smiles_sanitized":"CCCCCCCCC=CCCCCCC(=O)O"
}
},
{
"id":"cpd01113_C",
"name":"Hexanoate",
"compartment":"C",
"annotation":{
"smiles":"CCCCCC(=O)[O-]",
"smiles_sanitized":"CCCCCC(=O)O"
}
},
{
"id":"cpd03642_C",
"name":"Adipate",
"compartment":"C",
"annotation":{
"smiles":"O=C([O-])CCCCC(=O)[O-]",
"smiles_sanitized":"O=C(O)CCCCC(=O)O"
}
},
{
"id":"cpd00868_C",
"name":"p-hydroxyphenylpyruvate",
"compartment":"C",
"annotation":{
"smiles":"O=C([O-])C(=O)Cc1ccc(O)cc1",
"smiles_sanitized":"O=C(O)C(=O)Cc1ccc(O)cc1"
}
},
{
"id":"cpd02991_C",
"name":"phosphoribulosylformimino-AICAR-phosphate",
"compartment":"C",
"annotation":{
"smiles":"NC(=O)c1ncn([C@@H]2O[C@H](COP(=O)([O-])[O-])[C@@H](O)[C@H]2O)c1N/C=N/CC(=O)[C@H](O)[C@H](O)COP(=O)([O-])[O-]",
"smiles_sanitized":"NC(=O)c1ncn(C2OC(COP(=O)(O)O)C(O)C2O)c1NC=NCC(=O)C(O)C(O)COP(=O)(O)O"
}
},
{
"id":"cpd00144_C",
"name":"UDPglucuronate",
"compartment":"C",
"annotation":{
"smiles":"O=C([O-])[C@H]1O[C@H](OP(=O)([O-])OP(=O)([O-])OC[C@H]2O[C@@H](n3ccc(=O)[nH]c3=O)[C@H](O)[C@@H]2O)[C@H](O)[C@@H](O)[C@@H]1O",
"smiles_sanitized":"O=C(O)C1OC(OP(=O)(O)OP(=O)(O)OCC2OC(n3ccc(=O)[nH]c3=O)C(O)C2O)C(O)C(O)C1O"
}
},
{
"id":"cpd03126_C",
"name":"(2E)-Hexadecenoyl-CoA",
"compartment":"C",
"annotation":{
"smiles":"CCCCCCCCCCCCC/C=C/C(=O)SCCNC(=O)CCNC(=O)[C@H](O)C(C)(C)COP(=O)([O-])OP(=O)([O-])OC[C@H]1O[C@@H](n2cnc3c(N)ncnc32)[C@H](O)[C@@H]1OP(=O)([O-])[O-]",
"smiles_sanitized":"CCCCCCCCCCCCCC=CC(=O)SCCNC(=O)CCNC(=O)C(O)C(C)(C)COP(=O)(O)OP(=O)(O)OCC1OC(n2cnc3c(N)ncnc32)C(O)C1OP(=O)(O)O"
}
},
{
"id":"cpd02431_C",
"name":"1-Pyrroline-5-carboxylate",
"compartment":"C",
"annotation":{
"smiles":"O=C([O-])[C@@H]1CCC=N1",
"smiles_sanitized":"O=C(O)C1CCC=N1"
}
},
{
"id":"cpd02978_C",
"name":"7,8-Dihydroneopterin 3'-triphosphate",
"compartment":"C",
"annotation":{
"smiles":"Nc1nc2c(c(=O)[nH]1)N=C([C@H](O)[C@H](O)COP(=O)([O-])OP(=O)([O-])OP(=O)([O-])[O-])CN2",
"smiles_sanitized":"Nc1nc2c(c(=O)[nH]1)N=C(C(O)C(O)COP(=O)(O)OP(=O)(O)OP(=O)(O)O)CN2"
}
},
{
"id":"ASN_TRNAS_C",
"name":"asparagine trna",
"compartment":"C"
},
{
"id":"ALA_TRNAS_C",
"name":"alanine trna",
"compartment":"C"
},
{
"id":"HEME_O_C",
"name":"HEME_O",
"compartment":"C"
},
{
"id":"cpd23520_C",
"name":"9-octadecenal",
"compartment":"C",
"annotation":{
"smiles":"CCCCCCCC/C=C\\CCCCCCCC=O",
"smiles_sanitized":"CCCCCCCCC=CCCCCCCCC=O"
}
},
{
"id":"TYR_TRNAS_C",
"name":"tyrosine trna",
"compartment":"C"
},
{
"id":"cpd11423_C",
"name":"1,2-Diacyl-sn-glycerol",
"compartment":"C",
"annotation":{
"smiles":"*C(=O)OCC(CO)OC(*)=O",
"smiles_sanitized":"*C(=O)OCC(CO)OC(*)=O"
}
},
{
"id":"cpd00497_C",
"name":"XMP",
"compartment":"C",
"annotation":{
"smiles":"O=c1[nH]c(=O)c2ncn([C@@H]3O[C@H](COP(=O)([O-])[O-])[C@@H](O)[C@H]3O)c2[nH]1",
"smiles_sanitized":"O=c1[nH]c(=O)c2ncn(C3OC(COP(=O)(O)O)C(O)C3O)c2[nH]1"
}
},
{
"id":"cpd00279_C",
"name":"Acetoacetyl-CoA",
"compartment":"C",
"annotation":{
"smiles":"CC(=O)CC(=O)SCCNC(=O)CCNC(=O)[C@H](O)C(C)(C)COP(=O)([O-])OP(=O)([O-])OC[C@H]1O[C@@H](n2cnc3c(N)ncnc32)[C@H](O)[C@@H]1OP(=O)([O-])[O-]",
"smiles_sanitized":"CC(=O)CC(=O)SCCNC(=O)CCNC(=O)C(O)C(C)(C)COP(=O)(O)OP(=O)(O)OCC1OC(n2cnc3c(N)ncnc32)C(O)C1OP(=O)(O)O"
}
},
{
"id":"cpd00139_C",
"name":"Glycolate",
"compartment":"C",
"annotation":{
"smiles":"O=C([O-])CO",
"smiles_sanitized":"O=C(O)CO"
}
},
{
"id":"cpd02720_C",
"name":"5-Amino-6--5-phosphoribitylaminouracil",
"compartment":"C",
"annotation":{
"smiles":"Nc1c(NC[C@H](O)[C@H](O)[C@H](O)COP(=O)([O-])[O-])[nH]c(=O)[nH]c1=O",
"smiles_sanitized":"Nc1c(NCC(O)C(O)C(O)COP(=O)(O)O)[nH]c(=O)[nH]c1=O"
}
},
{
"id":"cpd00504_C",
"name":"LL-2,6-Diaminopimelate",
"compartment":"C",
"annotation":{
"smiles":"[NH3+][C@@H](CCC[C@H]([NH3+])C(=O)[O-])C(=O)[O-]",
"smiles_sanitized":"NC(CCCC(N)C(=O)O)C(=O)O"
}
},
{
"id":"cpd00443_C",
"name":"ABEE",
"compartment":"C",
"annotation":{
"smiles":"Nc1ccc(C(=O)[O-])cc1",
"smiles_sanitized":"Nc1ccc(C(=O)O)cc1"
}
},
{
"id":"cpd00652_E",
"name":"D-Mucic acid",
"compartment":"E",
"annotation":{
"smiles":"O=C([O-])[C@@H](O)[C@H](O)[C@H](O)[C@@H](O)C(=O)[O-]",
"smiles_sanitized":"O=C(O)C(O)C(O)C(O)C(O)C(=O)O"
}
},
{
"id":"cpd00797_C",
"name":"(R)-3-Hydroxybutanoate",
"compartment":"C",
"annotation":{
"smiles":"C[C@@H](O)CC(=O)[O-]",
"smiles_sanitized":"CC(O)CC(=O)O"
}
},
{
"id":"cpd00493_C",
"name":"Pyridoxamine phosphate",
"compartment":"C",
"annotation":{
"smiles":"Cc1ncc(COP(=O)([O-])[O-])c(C[NH3+])c1O",
"smiles_sanitized":"Cc1ncc(COP(=O)(O)O)c(CN)c1O"
}
},
{
"id":"COFACTOR_W_C",
"name":"COFACTOR_W",
"compartment":"C"
},
{
"id":"cpd08286_C",
"name":"2-C-methyl-D-erythritol4-phosphate",
"compartment":"C",
"annotation":{
"smiles":"C[C@](O)(CO)[C@H](O)COP(=O)([O-])[O-]",
"smiles_sanitized":"CC(O)(CO)C(O)COP(=O)(O)O"
}
},
{
"id":"cpd00249_E",
"name":"Uridine",
"compartment":"E",
"annotation":{
"smiles":"O=c1ccn([C@@H]2O[C@H](CO)[C@@H](O)[C@H]2O)c(=O)[nH]1",
"smiles_sanitized":"O=c1ccn(C2OC(CO)C(O)C2O)c(=O)[nH]1"
}
},
{
"id":"C12_0_ACP_C",
"name":"dodecanoyl-acp",
"compartment":"C"
},
{
"id":"PROTEIN_W_C",
"name":"PROTEIN_W",
"compartment":"C"
},
{
"id":"cpd00075_E",
"name":"Nitrite",
"compartment":"E",
"annotation":{
"smiles":"O=NO",
"smiles_sanitized":"O=NO"
}
},
{
"id":"cpd00727_C",
"name":"2-Phosphoglycolate",
"compartment":"C",
"annotation":{
"smiles":"O=C([O-])COP(=O)([O-])[O-]",
"smiles_sanitized":"O=C(O)COP(=O)(O)O"
}
},
{
"id":"cpd03120_C",
"name":"(S)-Hydroxyoctanoyl-CoA",
"compartment":"C",
"annotation":{
"smiles":"CCCCC[C@H](O)CC(=O)SCCNC(=O)CCNC(=O)C(O)C(C)(C)COP(=O)([O-])OP(=O)([O-])OC[C@H]1O[C@@H](n2cnc3c(N)ncnc32)[C@H](O)[C@@H]1OP(=O)([O-])[O-]",
"smiles_sanitized":"CCCCCC(O)CC(=O)SCCNC(=O)CCNC(=O)C(O)C(C)(C)COP(=O)(O)OP(=O)(O)OCC1OC(n2cnc3c(N)ncnc32)C(O)C1OP(=O)(O)O"
}
},
{
"id":"cpd00139_E",
"name":"Glycolate",
"compartment":"E",
"annotation":{
"smiles":"O=C([O-])CO",
"smiles_sanitized":"O=C(O)CO"
}
},
{
"id":"cpd00284_C",
"name":"6-Phospho-D-gluconate",
"compartment":"C",
"annotation":{
"smiles":"O=C([O-])[C@H](O)[C@@H](O)[C@H](O)[C@H](O)COP(=O)([O-])[O-]",
"smiles_sanitized":"O=C(O)C(O)C(O)C(O)C(O)COP(=O)(O)O"
}
},
{
"id":"cpd02103_C",
"name":"N2-Succinyl-L-arginine",
"compartment":"C",
"annotation":{
"smiles":"NC(=[NH2+])NCCC[C@H](NC(=O)CCC(=O)[O-])C(=O)[O-]",
"smiles_sanitized":"N=C(N)NCCCC(NC(=O)CCC(=O)O)C(=O)O"
}
},
{
"id":"FRUCTOSE_E",
"name":"external FRUCTOSE",
"compartment":"E",
"charge":0
},
{
"id":"FRUCTOSE_C",
"name":"cyotplasmic FRUCTOSE",
"compartment":"C",
"charge":0
}
],
"reactions":[
{
"id":"CARBOXYLESTERASE_RXN",
"name":"carboxylesterase",
"metabolites":{
"ACYL_OH_WAX_C":1.0,
"AFATTYACID_WAX_C":1.0,
"WAX_ESTER_C":-1.0,
"cpd00001_C":-1.0
},
"lower_bound":0.0,
"upper_bound":1000.0,
"gene_reaction_rule":"ACIAD1134 or ACIAD3648"
},
{
"id":"DIAMINOPIMEPIM_RXN",
"name":"diaminopimelate epimerase",
"metabolites":{
"cpd00504_C":-1.0,
"cpd00516_C":1.0
},
"lower_bound":-1000.0,
"upper_bound":1000.0,
"gene_reaction_rule":"ACIAD2659"
},
{
"id":"EXF_2_KETOGLUTARATE_E",
"name":"exchange flux of 2-oxoglutarate",
"metabolites":{
"cpd00024_E":-1.0
},
"lower_bound":-0.0,
"upper_bound":1000.0,
"gene_reaction_rule":""
},
{
"id":"DEPHOSPHOCOAKIN_RXN",
"name":"dephospho-coa kinase",
"metabolites":{
"cpd00002_C":-1.0,
"cpd00008_C":1.0,
"cpd00010_C":1.0,
"cpd00655_C":-1.0
},
"lower_bound":-1000.0,
"upper_bound":1000.0,
"gene_reaction_rule":"ACIAD0359"
},
{
"id":"4_1_1_32_RXN",
"name":"phosphoenolpyruvate carboxykinase (gtp)",
"metabolites":{
"cpd00011_C":1.0,
"cpd00031_C":1.0,
"cpd00032_C":-1.0,
"cpd00038_C":-1.0,
"cpd00061_C":1.0
},
"lower_bound":0.0,
"upper_bound":1000.0,
"gene_reaction_rule":"ACIAD2842"
},
{
"id":"GARTRANSFORMYL2_RXN",
"name":"phosphoribosylglycinamide formyltransferase 2",
"metabolites":{
"cpd00002_C":-1.0,
"cpd00008_C":1.0,
"cpd00009_C":1.0,
"cpd00047_C":-1.0,
"cpd02394_C":-1.0,
"cpd02678_C":1.0
},
"lower_bound":0.0,
"upper_bound":1000.0,
"gene_reaction_rule":"ACIAD2078"
},
{
"id":"DIHYDROXYISOVALDEHYDRAT_RXN",
"name":"dihydroxy-acid dehydratase",
"metabolites":{
"cpd00001_C":1.0,
"cpd00123_C":1.0,
"cpd02498_C":-1.0
},
"lower_bound":-1000.0,
"upper_bound":1000.0,
"gene_reaction_rule":"ACIAD1266 or ACIAD3636"
},
{
"id":"ISOCITDEH_RXN",
"name":"isocitrate dehydrogenase (nadp+)",
"metabolites":{
"cpd00005_C":1.0,
"cpd00006_C":-1.0,
"cpd00011_C":1.0,
"cpd00024_C":1.0,
"cpd00260_C":-1.0
},
"lower_bound":0.0,
"upper_bound":1000.0,
"gene_reaction_rule":"ACIAD1187 or ACIAD1190"
},
{
"id":"DGDPKIN_RXN",
"name":"dgdp kinase",
"metabolites":{
"cpd00002_C":-1.0,
"cpd00008_C":1.0,
"cpd00241_C":1.0,
"cpd00295_C":-1.0
},
"lower_bound":-1000.0,
"upper_bound":1000.0,
"gene_reaction_rule":"ACIAD0556"
},
{
"id":"GALACTARDEHYDRA_RXN",
"name":"galactarate dehydratase",
"metabolites":{
"cpd00001_C":1.0,
"cpd00515_C":1.0,
"cpd00652_C":-1.0
},
"lower_bound":0.0,
"upper_bound":1000.0,
"gene_reaction_rule":"ACIAD0126"
},
{
"id":"C16_1_C7_ALDEHYDE_REDUCTASE_RXN",
"name":"long chain aldehyde reductase (c16:1(c7))",
"metabolites":{
"cpd00003_C":1.0,
"cpd00004_C":-1.0,
"cpd16626_C":-1.0,
"cpd16628_C":1.0
},
"lower_bound":-1000.0,
"upper_bound":1000.0,
"gene_reaction_rule":""
},
{
"id":"C12_0_ACP_HYDROLASE_RXN",
"name":"acyl-acp hydrolase (c12:0)",
"metabolites":{
"ACP_C":1.0,
"C12_0_ACP_C":-1.0,
"cpd00001_C":-1.0,
"cpd01741_C":1.0
},
"lower_bound":0.0,
"upper_bound":1000.0,
"gene_reaction_rule":""
},
{
"id":"L_ASPARTATE_OXID_RXN",
"name":"l-aspartate oxidase (producing iminoaspartate)",
"metabolites":{
"cpd00007_C":-1.0,
"cpd00025_C":1.0,
"cpd00041_C":-1.0,
"cpd03470_C":1.0
},
"lower_bound":0.0,
"upper_bound":1000.0,
"gene_reaction_rule":"ACIAD2587"
},
{
"id":"RXN0_902",
"name":"hypoxanthine dehydrogenase",
"metabolites":{
"cpd00001_C":-1.0,
"cpd00003_C":-1.0,
"cpd00004_C":1.0,
"cpd00226_C":-1.0,
"cpd00309_C":1.0
},
"lower_bound":-1000.0,
"upper_bound":1000.0,
"gene_reaction_rule":"ACIAD2466 and ACIAD2467 and ACIAD2468"
},
{
"id":"C18_0_3OXOACP_REDUCTASE_RXN",
"name":"3oxoacyl-acp reductase (c18:0)",
"metabolites":{
"C18_0_3HYDROXYACP_C":1.0,
"C18_0_3OXOACP_C":-1.0,
"cpd00005_C":-1.0,
"cpd00006_C":1.0
},
"lower_bound":0.0,
"upper_bound":1000.0,
"gene_reaction_rule":"ACIAD0871"
},
{
"id":"C16_0_COATRANSFERASE_RXN",
"name":"acyl coa transferase (c16:0)",
"metabolites":{
"cpd00002_C":-1.0,
"cpd00010_C":-1.0,
"cpd00012_C":1.0,
"cpd00018_C":1.0,
"cpd00134_C":1.0,
"cpd00214_C":-1.0
},
"lower_bound":0.0,
"upper_bound":1000.0,
"gene_reaction_rule":"ACIAD1818 or ACIAD0235 or ACIAD1724 or ACIAD2106"
},
{
"id":"ACETOACETYL_COA_TRANSFER_RXN",
"name":"acetoacetyl-coa transferase",
"metabolites":{
"cpd00022_C":-1.0,
"cpd00029_C":1.0,
"cpd00142_C":-1.0,
"cpd00279_C":1.0
},
"lower_bound":-1000.0,
"upper_bound":1000.0,
"gene_reaction_rule":"ACIAD2513 and ACIAD2514"
},
{
"id":"ATP_DRAIN_RXN",
"name":"atp maintenance requirement",
"metabolites":{
"cpd00001_C":-1.0,
"cpd00002_C":-1.0,
"cpd00008_C":1.0,
"cpd00009_C":1.0,
"cpd00067_C":1.0
},
"lower_bound":0.0,
"upper_bound":1000.0,
"gene_reaction_rule":""
},
{
"id":"DHHB_METHYLTRANSFER_RXN",
"name":"3-demethylubiquinone-9 3-o-methyltransferase",
"metabolites":{
"cpd00017_C":-1.0,
"cpd00019_C":1.0,
"cpd03449_C":-1.0,
"cpd15561_C":1.0
},
"lower_bound":0.0,
"upper_bound":1000.0,
"gene_reaction_rule":"ACIAD0044"
},
{
"id":"EXF_COUMARATE_E",
"name":"exchange flux of 4-coumarate",
"metabolites":{
"cpd00604_E":-1.0
},
"lower_bound":-0.0,
"upper_bound":1000.0,
"gene_reaction_rule":""
},
{
"id":"ALKANAL_MONOOXYGENASE_FMN_LINKED_RXN",
"name":"octanal monooxygenase (fmn-linked)",
"metabolites":{
"cpd00001_C":1.0,
"cpd00007_C":-1.0,
"cpd00050_C":1.0,
"cpd01088_C":-1.0,
"cpd01270_C":-1.0,
"cpd03846_C":1.0
},
"lower_bound":0.0,
"upper_bound":1000.0,
"gene_reaction_rule":"ACIAD1830"
},
{
"id":"INOSINE_NUCLEOSIDASE_RXN",
"name":"inosine nucleosidase",
"metabolites":{
"cpd00001_C":-1.0,
"cpd00105_C":1.0,
"cpd00226_C":1.0,
"cpd00246_C":-1.0
},
"lower_bound":0.0,
"upper_bound":1000.0,
"gene_reaction_rule":"ACIAD2386"
},
{
"id":"CYT_UBIQUINOL_OXID_O_RXN",
"name":"cytochrome oxidase bo",
"metabolites":{
"cpd00001_C":2.0,
"cpd00007_C":-1.0,
"cpd00067_C":-8.0,
"cpd00067_P":8.0,
"cpd15560_C":2.0,
"cpd15561_C":-2.0
},
"lower_bound":0.0,
"upper_bound":1000.0,
"gene_reaction_rule":"ACIAD2428 and ACIAD2427 and ACIAD2426 and ACIAD2425"
},
{
"id":"UDP_NACMURALGLDAPLIG_RXN",
"name":"udp-n-acetylmuramoylalanyl-d-glutamate--2,6-diaminopimelate ligase",
"metabolites":{
"cpd00002_C":-1.0,
"cpd00008_C":1.0,
"cpd00009_C":1.0,
"cpd00516_C":-1.0,
"cpd00525_C":-1.0,
"cpd02964_C":1.0
},
"lower_bound":0.0,
"upper_bound":1000.0,
"gene_reaction_rule":"ACIAD3365"
},
{
"id":"PYRROLINECARBREDUCT_RXN",
"name":"pyrroline-5-carboxylate reductase",
"metabolites":{
"cpd00005_C":-1.0,
"cpd00006_C":1.0,
"cpd00129_C":1.0,
"cpd02431_C":-1.0
},
"lower_bound":0.0,
"upper_bound":1000.0,
"gene_reaction_rule":"ACIAD3032"
},
{
"id":"UREIDOGLYCOLATE_HYDROLASE_RXN",
"name":"ureidoglycolate hydrolase",
"metabolites":{
"cpd00001_C":-1.0,
"cpd00011_C":1.0,
"cpd00013_C":2.0,
"cpd00040_C":1.0,
"cpd00465_C":-1.0
},
"lower_bound":0.0,
"upper_bound":1000.0,
"gene_reaction_rule":"ACIAD3542"
},
{
"id":"HISTPRATPHYD_RXN",
"name":"phosphoribosyl-atp pyrophosphatase",
"metabolites":{
"cpd00001_C":-1.0,
"cpd00012_C":1.0,
"cpd01775_C":-1.0,
"cpd01777_C":1.0
},
"lower_bound":0.0,
"upper_bound":1000.0,
"gene_reaction_rule":"ACIAD0380"
},
{
"id":"C18_1_C9_3OXOCOA_THIOLASE_RXN",
"name":"3oxoacyl-coa thiolase (c18:1(c9))",
"metabolites":{
"cpd00010_C":-1.0,
"cpd00022_C":1.0,
"cpd16627_C":1.0,
"cpd32253_C":-1.0
},
"lower_bound":-1000.0,
"upper_bound":1000.0,
"gene_reaction_rule":"ACIAD0334 or ACIAD1450 or ACIAD1706 or ACIAD1977 or ACIAD1689 or ACIAD2988 or ACIAD1569"
},
{
"id":"TRANS_RXN_URIDINE",
"name":"transport of uridine [extraorganism;cytosol](secondary symport)",
"metabolites":{
"cpd00067_C":1.0,
"cpd00067_P":-1.0,
"cpd00249_C":1.0,
"cpd00249_E":-1.0
},
"lower_bound":0.0,
"upper_bound":1000.0,
"gene_reaction_rule":"ACIAD2389"
},
{
"id":"TRANS_RXN_C6_0",
"name":"transport of hexanoate [extraorganism;cytosol](secondary symport)",
"metabolites":{
"cpd00067_C":1.0,
"cpd00067_P":-1.0,
"cpd01113_C":1.0,
"cpd01113_E":-1.0
},
"lower_bound":-1000.0,
"upper_bound":1000.0,
"gene_reaction_rule":"ACIAD0835 or ACIAD1427 or ACIAD2788 or ACIAD3392"
},
{
"id":"C8_0_3OXOACP_SYNTHASE_RXN",
"name":"3oxoacyl-acp synthase (c8:0)",
"metabolites":{
"ACP_C":1.0,
"C6_0_ACP_C":-1.0,
"C8_0_3OXOACP_C":1.0,
"MALONYL_ACP_C":-1.0,
"cpd00011_C":1.0
},
"lower_bound":0.0,
"upper_bound":1000.0,
"gene_reaction_rule":"ACIAD0580 or ACIAD0879 or ACIAD0583 or ACIAD2101"
},
{
"id":"GLUTATHIONE_PEROXIDASE_RXN",
"name":"glutathione peroxidase",
"metabolites":{
"cpd00001_C":2.0,
"cpd00025_C":-1.0,
"cpd00042_C":-2.0,
"cpd00111_C":1.0
},
"lower_bound":0.0,
"upper_bound":1000.0,
"gene_reaction_rule":"ACIAD0193 or ACIAD2085"
},
{
"id":"R142_RXN",
"name":"4-diphosphocytidyl-2-c-methyl-d-erythritol kinase",
"metabolites":{
"cpd00002_C":-1.0,
"cpd00008_C":1.0,
"cpd08287_C":-1.0,
"cpd08288_C":1.0
},
"lower_bound":0.0,
"upper_bound":1000.0,
"gene_reaction_rule":"ACIAD2903"
},
{
"id":"DIHYDROFOLATESYNTH_RXN",
"name":"dihydrofolate synthase",
"metabolites":{
"cpd00002_C":-1.0,
"cpd00008_C":1.0,
"cpd00009_C":1.0,
"cpd00023_C":-1.0,
"cpd00330_C":1.0,
"cpd00683_C":-1.0
},
"lower_bound":0.0,
"upper_bound":1000.0,
"gene_reaction_rule":"ACIAD0644"
},
{
"id":"MALONYL_ACPDECARBOX_RXN",
"name":"malonyl-acp decarboxylase",
"metabolites":{
"ACETYL_ACP_C":1.0,
"MALONYL_ACP_C":-1.0,
"cpd00011_C":1.0
},
"lower_bound":0.0,
"upper_bound":1000.0,
"gene_reaction_rule":"ACIAD0879"
},
{
"id":"DHSHIKIMATE_DEHYDRO_RXN",
"name":"3-dehydroshikimate dehydratase",
"metabolites":{
"cpd00001_P":1.0,
"cpd00197_P":1.0,
"cpd01716_P":-1.0
},
"lower_bound":-1000.0,
"upper_bound":1000.0,
"gene_reaction_rule":"ACIAD1714"
},
{
"id":"GLUCARDEHYDRA_RXN",
"name":"glucarate dehydratase",
"metabolites":{
"cpd00001_C":1.0,
"cpd00515_C":1.0,
"cpd00571_C":-1.0
},
"lower_bound":0.0,
"upper_bound":1000.0,
"gene_reaction_rule":"ACIAD0128"
},
{
"id":"2_OCTAPRENYLPHENOL_HYDROX_RXN",
"name":"2-octaprenylphenol hydroxylase",
"metabolites":{
"cpd00001_C":1.0,
"cpd00005_C":-1.0,
"cpd00006_C":1.0,
"cpd00007_C":-1.0,
"cpd00067_C":-1.0,
"cpd03444_C":-1.0,
"cpd03445_C":1.0
},
"lower_bound":0.0,
"upper_bound":1000.0,
"gene_reaction_rule":"ACIAD0382"
},
{
"id":"DALADEHYDROG_RXN",
"name":"d-alanine dehydrogenase",
"metabolites":{
"cpd00001_C":-1.0,
"cpd00013_C":1.0,
"cpd00015_C":-1.0,
"cpd00020_C":1.0,
"cpd00117_C":-1.0,
"cpd00982_C":1.0
},
"lower_bound":-1000.0,
"upper_bound":1000.0,
"gene_reaction_rule":"ACIAD1998 or ACIAD0922 or ACIAD0115"
},
{
"id":"C14_0_3HYDROXYCOA_DEHYDROGENASE_RXN",
"name":"hydroxyacyl-coa dehydrogenase (c14:0)",
"metabolites":{
"cpd00003_C":-1.0,
"cpd00004_C":1.0,
"cpd03115_C":-1.0,
"cpd12689_C":1.0
},
"lower_bound":0.0,
"upper_bound":1000.0,
"gene_reaction_rule":"ACIAD0335 or ACIAD1565 or ACIAD2416"
},
{
"id":"TETRAACYLDISACC4KIN_RXN",
"name":"tetraacyldisaccharide 4'-kinase",
"metabolites":{
"cpd00002_C":-1.0,
"cpd00008_C":1.0,
"cpd02993_C":1.0,
"cpd03002_C":-1.0
},
"lower_bound":0.0,
"upper_bound":1000.0,
"gene_reaction_rule":"ACIAD2363"
},
{
"id":"PHOSPHAGLYPSYN_RXN",
"name":"cdp-diacylglycerol--glycerol-3-phosphate 3-phosphatidyltransferase",
"metabolites":{
"CDPDIACYLGLYCEROL_C":-1.0,
"L_1_PHOSPHATIDYL_GLYCEROL_P_C":1.0,
"cpd00046_C":1.0,
"cpd00080_C":-1.0
},
"lower_bound":0.0,
"upper_bound":1000.0,
"gene_reaction_rule":"ACIAD0342"
},
{
"id":"THREDEHYD_RXN",
"name":"threonine dehydratase",
"metabolites":{
"cpd00013_C":1.0,
"cpd00094_C":1.0,
"cpd00161_C":-1.0
},
"lower_bound":0.0,
"upper_bound":1000.0,
"gene_reaction_rule":"ACIAD1357"
},
{
"id":"OCTAPRENYL_METHYL_METHOXY_BENZOQ_OH_RXN",
"name":"2-octaprenyl-3-methyl-6-methoxy-1,4-benzoquinol hydroxylase",
"metabolites":{
"cpd00007_C":-1.0,
"cpd03448_C":-1.0,
"cpd03449_C":1.0
},
"lower_bound":0.0,
"upper_bound":1000.0,
"gene_reaction_rule":"ACIAD1128"
},
{
"id":"CARDIOLIPSYN_RXN",
"name":"cardiolipin synthase",
"metabolites":{
"L_1_PHOSPHATIDYL_GLYCEROL_C":-2.0,
"cpd00100_C":1.0,
"cpd12801_C":1.0
},
"lower_bound":-1000.0,
"upper_bound":1000.0,
"gene_reaction_rule":""
},
{
"id":"RXNIM_6099",
"name":"caffeoyl-coa lyase",
"metabolites":{
"cpd00001_C":-1.0,
"cpd00270_C":-1.0,
"cpd33217_C":1.0
},
"lower_bound":-1000.0,
"upper_bound":1000.0,
"gene_reaction_rule":"ACIAD1726"
},
{
"id":"GLUTAMINESYN_RXN",
"name":"glutamate--ammonia ligase",
"metabolites":{
"cpd00002_C":-1.0,
"cpd00008_C":1.0,
"cpd00009_C":1.0,
"cpd00013_C":-1.0,
"cpd00023_C":-1.0,
"cpd00053_C":1.0
},
"lower_bound":0.0,
"upper_bound":1000.0,
"gene_reaction_rule":"ACIAD2458 or ACIAD2528"
},
{
"id":"GLUTARYL_COA_DEHYDROGENASE_RXN",
"name":"glutaryl-coa dehydrogenase (producing crotonyl-coa)",
"metabolites":{
"cpd00011_C":1.0,
"cpd00015_C":-1.0,
"cpd00413_C":-1.0,
"cpd00650_C":1.0,
"cpd00982_C":1.0
},
"lower_bound":0.0,
"upper_bound":1000.0,
"gene_reaction_rule":"ACIAD1660"
},
{
"id":"ACETYLORNDEACET_RXN",
"name":"acetylornithine deacetylase",
"metabolites":{
"cpd00001_C":-1.0,
"cpd00029_C":1.0,
"cpd00064_C":1.0,
"cpd00342_C":-1.0
},
"lower_bound":0.0,
"upper_bound":1000.0,
"gene_reaction_rule":"ACIAD3489"
},
{
"id":"RXNIM_6092",
"name":"o-acylhomoserine sulfhydrylase",
"metabolites":{
"cpd00029_C":1.0,
"cpd00084_C":-1.0,
"cpd00424_C":1.0,
"cpd00790_C":-1.0
},
"lower_bound":0.0,
"upper_bound":1000.0,
"gene_reaction_rule":"ACIAD2314"
},
{
"id":"RXN0_277",
"name":"2-c-methyl-d-erythritol 4-phosphate cytidylyltransferase",
"metabolites":{
"cpd00012_C":1.0,
"cpd00052_C":-1.0,
"cpd08286_C":-1.0,
"cpd08287_C":1.0
},
"lower_bound":0.0,
"upper_bound":1000.0,
"gene_reaction_rule":"ACIAD1999"
},
{
"id":"RXNIM_6094",
"name":"3,4-dihydroxyphenyl-beta-hydroxypropionyl-coa lyase",
"metabolites":{
"cpd00022_C":1.0,
"cpd16507_C":1.0,
"cpd33217_C":-1.0
},
"lower_bound":-1000.0,
"upper_bound":1000.0,
"gene_reaction_rule":"ACIAD1726"
},
{
"id":"RXNIM_6095",
"name":"coumaryl-coa lyase",
"metabolites":{
"cpd00001_C":-1.0,
"cpd00192_C":-1.0,
"cpd23287_C":1.0
},
"lower_bound":-1000.0,
"upper_bound":1000.0,
"gene_reaction_rule":"ACIAD1726"
},
{
"id":"TRANS_RXN_ASN",
"name":"transport of l-asparagine [extraorganism;cytosol](passive)",
"metabolites":{
"cpd00132_C":1.0,
"cpd00132_E":-1.0
},
"lower_bound":0.0,
"upper_bound":1000.0,
"gene_reaction_rule":""
},
{
"id":"EXF_ASN_E",
"name":"exchange flux of l-asparagine",
"metabolites":{
"cpd00132_E":-1.0
},
"lower_bound":0.0,
"upper_bound":1000.0,
"gene_reaction_rule":""
},
{
"id":"DLACTDEHYDROGFAD_RXN",
"name":"d-lactate dehydrogenase, fad",
"metabolites":{
"cpd00015_C":-1.0,
"cpd00020_C":1.0,
"cpd00221_C":-1.0,
"cpd00982_C":1.0
},
"lower_bound":0.0,
"upper_bound":1000.0,
"gene_reaction_rule":"ACIAD0109"
},
{
"id":"HISTIDINE_AMMONIA_LYASE_RXN",
"name":"histidine ammonia-lyase",
"metabolites":{
"cpd00013_C":1.0,
"cpd00119_C":-1.0,
"cpd00581_C":1.0
},
"lower_bound":0.0,
"upper_bound":1000.0,
"gene_reaction_rule":"ACIAD0574"
},
{
"id":"TRANS_RXN_100",
"name":"transport of l-carnitine [extraorganism;cytosol](secondary symport)",
"metabolites":{
"cpd00067_C":1.0,
"cpd00067_P":-1.0,
"cpd00266_C":1.0,
"cpd00266_E":-1.0
},
"lower_bound":0.0,
"upper_bound":1000.0,
"gene_reaction_rule":"ACIAD3460 or ACIAD1011"
},
{
"id":"C10_0_3OXOACP_SYNTHASE_RXN",
"name":"3oxoacyl-acp synthase (c10:0)",
"metabolites":{
"ACP_C":1.0,
"C10_0_3OXOACP_C":1.0,
"C8_0_ACP_C":-1.0,
"MALONYL_ACP_C":-1.0,
"cpd00011_C":1.0
},
"lower_bound":0.0,
"upper_bound":1000.0,
"gene_reaction_rule":"ACIAD0580 or ACIAD0879 or ACIAD0583 or ACIAD2101"
},
{
"id":"METHYLENETHFREDUCT_RXN",
"name":"5,10-methylenetetrahydrofolate reductase (fadh)",
"metabolites":{
"cpd00003_C":-1.0,
"cpd00004_C":1.0,
"cpd00125_C":1.0,
"cpd00345_C":-1.0
},
"lower_bound":-1000.0,
"upper_bound":1000.0,
"gene_reaction_rule":"ACIAD2283"
},
{
"id":"NAD_SYNTH_GLN_RXN",
"name":"nad(+) synthase (glutamine-hydrolysing)",
"metabolites":{
"cpd00001_C":-1.0,
"cpd00002_C":-1.0,
"cpd00003_C":1.0,
"cpd00012_C":1.0,
"cpd00018_C":1.0,
"cpd00023_C":1.0,
"cpd00053_C":-1.0,
"cpd00638_C":-1.0
},
"lower_bound":-1000.0,
"upper_bound":1000.0,
"gene_reaction_rule":"ACIAD2888"
},
{
"id":"ORNDECARBOX_RXN",
"name":"ornithine decarboxylase",
"metabolites":{
"cpd00011_C":1.0,
"cpd00064_C":-1.0,
"cpd00118_C":1.0
},
"lower_bound":0.0,
"upper_bound":1000.0,
"gene_reaction_rule":"ACIAD2768"
},
{
"id":"C18_0_3OXOACP_SYNTHASE_RXN",
"name":"3oxoacyl-acp synthase (c18:0)",
"metabolites":{
"ACP_C":1.0,
"C16_0_ACP_C":-1.0,
"C18_0_3OXOACP_C":1.0,
"MALONYL_ACP_C":-1.0,
"cpd00011_C":1.0
},
"lower_bound":0.0,
"upper_bound":1000.0,
"gene_reaction_rule":"ACIAD0580 or ACIAD0879 or ACIAD0583 or ACIAD2101"
},
{
"id":"TRANSALDOL_RXN",
"name":"transaldolase",
"metabolites":{
"cpd00102_C":-1.0,
"cpd00236_C":1.0,
"cpd00238_C":-1.0,
"cpd15394_C":1.0
},
"lower_bound":-1000.0,
"upper_bound":1000.0,
"gene_reaction_rule":"ACIAD1980"
},
{
"id":"TRANS_RXN_ADENOSINE",
"name":"transport of adenosine [extraorganism;cytosol](secondary symport)",
"metabolites":{
"cpd00067_C":1.0,
"cpd00067_P":-1.0,
"cpd00182_C":1.0,
"cpd00182_E":-1.0
},
"lower_bound":-1000.0,
"upper_bound":1000.0,
"gene_reaction_rule":"ACIAD2389"
},
{
"id":"TRANS_RXN_DEOXYURIDINE",
"name":"transport of deoxyuridine [extraorganism;cytosol](secondary symport)",
"metabolites":{
"cpd00067_C":1.0,
"cpd00067_P":-1.0,
"cpd00412_C":1.0,
"cpd00412_E":-1.0
},
"lower_bound":0.0,
"upper_bound":1000.0,
"gene_reaction_rule":"ACIAD2389"
},
{
"id":"ARGSUCCINSYN_RXN",
"name":"argininosuccinate synthase",
"metabolites":{
"cpd00002_C":-1.0,
"cpd00012_C":1.0,
"cpd00018_C":1.0,
"cpd00041_C":-1.0,
"cpd00274_C":-1.0,
"cpd02152_C":1.0
},
"lower_bound":-1000.0,
"upper_bound":1000.0,
"gene_reaction_rule":"ACIAD1151"
},
{
"id":"DTDPDEHYRHAMREDUCT_RXN",
"name":"dtdp-4-dehydrorhamnose reductase",
"metabolites":{
"cpd00005_C":-1.0,
"cpd00006_C":1.0,
"cpd00522_C":-1.0,
"cpd02113_C":1.0
},
"lower_bound":-1000.0,
"upper_bound":1000.0,
"gene_reaction_rule":"ACIAD0078"
},
{
"id":"DGTPTRIPHYDRO_RXN",
"name":"deoxyguanosinetriphosphate triphosphohydrolase",
"metabolites":{
"cpd00001_C":-1.0,
"cpd00241_C":-1.0,
"cpd00277_C":1.0,
"cpd00421_C":1.0
},
"lower_bound":0.0,
"upper_bound":1000.0,
"gene_reaction_rule":"ACIAD2613"
},
{
"id":"TRIACYLGLYCEROL_LIPASE_RXN",
"name":"triacylglycerol lipase",
"metabolites":{
"AFATTYACID_TAG_C":1.0,
"cpd00001_C":-1.0,
"cpd11423_C":1.0,
"cpd11677_C":-1.0
},
"lower_bound":0.0,
"upper_bound":1000.0,
"gene_reaction_rule":"ACIAD3309 or ACIAD1121"
},
{
"id":"EXF_CIS_CIS_MUCONATE_E",
"name":"exchange flux of cis,cis-muconate",
"metabolites":{
"cpd01630_E":-1.0
},
"lower_bound":-0.0,
"upper_bound":1000.0,
"gene_reaction_rule":""
},
{
"id":"EXF_CIT_E",
"name":"exchange flux of citrate",
"metabolites":{
"cpd00137_E":-1.0
},
"lower_bound":-0.0,
"upper_bound":1000.0,
"gene_reaction_rule":""
},
{
"id":"TRANS_RXN_GLN",
"name":"transport of l-glutamine [extraorganism;cytosol](passive)",
"metabolites":{
"cpd00053_C":1.0,
"cpd00053_E":-1.0
},
"lower_bound":0.0,
"upper_bound":1000.0,
"gene_reaction_rule":""
},
{
"id":"2_8_1_6_RXN",
"name":"biotin synthase",
"metabolites":{
"cpd00035_C":1.0,
"cpd00084_C":-1.0,
"cpd00104_C":1.0,
"cpd01311_C":-1.0
},
"lower_bound":-1000.0,
"upper_bound":1000.0,
"gene_reaction_rule":"ACIAD2045"
},
{
"id":"DIOHBUTANONEPSYN_RXN",
"name":"3,4-dihydroxy-2-butanone-4-phosphate synthase",
"metabolites":{
"cpd00047_C":1.0,
"cpd00067_C":1.0,
"cpd00171_C":-1.0,
"cpd11225_C":1.0
},
"lower_bound":-1000.0,
"upper_bound":1000.0,
"gene_reaction_rule":"ACIAD2913 or ACIAD3570"
},
{
"id":"EXF_CPD_205_E",
"name":"exchange flux of pimelate",
"metabolites":{
"cpd01727_E":-1.0
},
"lower_bound":-0.0,
"upper_bound":1000.0,
"gene_reaction_rule":""
},
{
"id":"METHIONINE_TRNA_LIGASE_RXN",
"name":"methionine--trna ligase",
"metabolites":{
"CHARGED_MET_TRNAS_C":1.0,
"MET_TRNAS_C":-1.0,
"cpd00002_C":-1.0,
"cpd00012_C":1.0,
"cpd00018_C":1.0,
"cpd00060_C":-1.0
},
"lower_bound":0.0,
"upper_bound":1000.0,
"gene_reaction_rule":"ACIAD0768"
},
{
"id":"C14_0_3OXOCOA_THIOLASE_RXN",
"name":"3oxoacyl-coa thiolase (c14:0)",
"metabolites":{
"cpd00010_C":-1.0,
"cpd00022_C":1.0,
"cpd01260_C":1.0,
"cpd12689_C":-1.0
},
"lower_bound":-1000.0,
"upper_bound":1000.0,
"gene_reaction_rule":"ACIAD0334 or ACIAD1450 or ACIAD1706 or ACIAD1977 or ACIAD1689 or ACIAD2988 or ACIAD1569"
},
{
"id":"GSAAMINOTRANS_RXN",
"name":"glutamate-1-semialdehyde 2,1-aminomutase",
"metabolites":{
"cpd00338_C":1.0,
"cpd02345_C":-1.0
},
"lower_bound":-1000.0,
"upper_bound":1000.0,
"gene_reaction_rule":"ACIAD1201"
},
{
"id":"PHOSMANMUT_RXN",
"name":"phosphomannomutase",
"metabolites":{
"cpd00235_C":1.0,
"cpd00485_C":-1.0
},
"lower_bound":-1000.0,
"upper_bound":1000.0,
"gene_reaction_rule":"ACIAD0902 or ACIAD0104"
},
{
"id":"EXF_SULFATE_E",
"name":"exchange flux of so42-",
"metabolites":{
"cpd00048_E":-1.0
},
"lower_bound":-1000.0,
"upper_bound":1000.0,
"gene_reaction_rule":""
},
{
"id":"RXN0_302",
"name":"2c-methyl-d-erythritol 2,4-cyclodiphosphate synthase",
"metabolites":{
"cpd00046_C":1.0,
"cpd08288_C":-1.0,
"cpd08301_C":1.0
},
"lower_bound":0.0,
"upper_bound":1000.0,
"gene_reaction_rule":"ACIAD1996"
},
{
"id":"DAMP_NUCLEOTIDASE_RXN",
"name":"damp nucleotidase",
"metabolites":{
"cpd00001_C":-1.0,
"cpd00009_C":1.0,
"cpd00294_C":-1.0,
"cpd00438_C":1.0
},
"lower_bound":0.0,
"upper_bound":1000.0,
"gene_reaction_rule":"ACIAD0015 or ACIAD1227"
},
{
"id":"RXN0_305",
"name":"hydroxypyruvate isomerase",
"metabolites":{
"cpd00145_C":-1.0,
"cpd00843_C":1.0
},
"lower_bound":-1000.0,
"upper_bound":1000.0,
"gene_reaction_rule":"ACIAD3075"
},
{
"id":"UNBAL_PROTON",
"name":"unbalance flux of h+",
"metabolites":{
"cpd00067_C":-1.0
},
"lower_bound":-1000.0,
"upper_bound":1000.0,
"gene_reaction_rule":""
},
{
"id":"HYDROXYBENZALDEHYDE_OXIDATION_NAD_RXN",
"name":"hydroxybenzaldehyde dehydrogenase",
"metabolites":{
"cpd00001_C":-1.0,
"cpd00003_C":-1.0,
"cpd00004_C":1.0,
"cpd00136_C":1.0,
"cpd00484_C":-1.0
},
"lower_bound":-1000.0,
"upper_bound":1000.0,
"gene_reaction_rule":"ACIAD1430 or ACIAD1725"
},
{
"id":"4_CARBOXYMUCONOLACTONE_DECARBOXYLASE_RXN",
"name":"4-carboxymuconolactone decarboxylase",
"metabolites":{
"cpd00011_C":1.0,
"cpd00938_C":-1.0,
"cpd02255_C":1.0
},
"lower_bound":0.0,
"upper_bound":1000.0,
"gene_reaction_rule":"ACIAD1710"
},
{
"id":"TRANS_RXN_29A",
"name":"transport of betaine [extraorganism;cytosol](secondary symport)",
"metabolites":{
"cpd00067_C":1.0,
"cpd00067_P":-1.0,
"cpd00540_C":1.0,
"cpd00540_E":-1.0
},
"lower_bound":0.0,
"upper_bound":1000.0,
"gene_reaction_rule":"ACIAD3460 or ACIAD1011"
},
{
"id":"MANNPGUANYLTRANGDP_RXN",
"name":"mannose-1-phosphate guanylyltransferase (gdp)",
"metabolites":{
"cpd00009_C":1.0,
"cpd00031_C":-1.0,
"cpd00083_C":1.0,
"cpd00485_C":-1.0
},
"lower_bound":0.0,
"upper_bound":1000.0,
"gene_reaction_rule":"ACIAD0086"
},
{
"id":"SUCCORNTRANSAM_RXN",
"name":"succinylornithine transaminase",
"metabolites":{
"cpd00023_C":1.0,
"cpd00024_C":-1.0,
"cpd02160_C":-1.0,
"cpd03526_C":1.0
},
"lower_bound":-1000.0,
"upper_bound":1000.0,
"gene_reaction_rule":"ACIAD1284"
},
{
"id":"NAG1P_URIDYLTRANS_RXN",
"name":"udp-n-acetylglucosamine pyrophosphorylase",
"metabolites":{
"cpd00012_C":1.0,
"cpd00037_C":1.0,
"cpd00062_C":-1.0,
"cpd02611_C":-1.0
},
"lower_bound":0.0,
"upper_bound":1000.0,
"gene_reaction_rule":"ACIAD3575"
},
{
"id":"GLYOXI_RXN",
"name":"lactoylglutathione lyase",
"metabolites":{
"cpd00042_C":-1.0,
"cpd00428_C":-1.0,
"cpd02182_C":1.0
},
"lower_bound":-1000.0,
"upper_bound":1000.0,
"gene_reaction_rule":"ACIAD2213"
},
{
"id":"R9_RXN",
"name":"crotonyl-coa hydratase",
"metabolites":{
"cpd00001_C":-1.0,
"cpd00650_C":-1.0,
"cpd00842_C":1.0
},
"lower_bound":-1000.0,
"upper_bound":1000.0,
"gene_reaction_rule":"ACIAD0335 or ACIAD1692 or ACIAD1726 or ACIAD1820 or ACIAD1829 or ACIAD2351 or ACIAD3073 or ACIAD1609 or ACIAD1608 or ACIAD1701"
},
{
"id":"EXF_CPD_335_E",
"name":"exchange flux of (r)-3-hydroxybutanoate",
"metabolites":{
"cpd00797_E":-1.0
},
"lower_bound":-0.0,
"upper_bound":1000.0,
"gene_reaction_rule":""
},
{
"id":"TRANS_RXN_CPD_302",
"name":"transport of d-aspartate [extraorganism;cytosol](passive)",
"metabolites":{
"cpd00320_C":1.0,
"cpd00320_E":-1.0
},
"lower_bound":0.0,
"upper_bound":1000.0,
"gene_reaction_rule":""
},
{
"id":"PSERTRANSAM_RXN",
"name":"phosphoserine aminotransferase",
"metabolites":{
"cpd00023_C":-1.0,
"cpd00024_C":1.0,
"cpd00738_C":1.0,
"cpd02069_C":-1.0
},
"lower_bound":-1000.0,
"upper_bound":1000.0,
"gene_reaction_rule":"ACIAD2647"
},
{
"id":"RXN0_5107",
"name":"dttp pyrophosphorylase",
"metabolites":{
"cpd00001_C":-1.0,
"cpd00012_C":1.0,
"cpd00298_C":1.0,
"cpd00357_C":-1.0
},
"lower_bound":0.0,
"upper_bound":1000.0,
"gene_reaction_rule":"ACIAD3065"
},
{
"id":"ACETYL_COA_ACETYLTRANSFER_RXN",
"name":"acetyl-coa c-acetyltransferase",
"metabolites":{
"cpd00010_C":-1.0,
"cpd00022_C":2.0,
"cpd00279_C":-1.0
},
"lower_bound":-1000.0,
"upper_bound":1000.0,
"gene_reaction_rule":"ACIAD2516 or ACIAD0694"
},
{
"id":"C10_0_3OXOACP_REDUCTASE_RXN",
"name":"3oxoacyl-acp reductase (c10:0)",
"metabolites":{
"C10_0_3HYDROXYACP_C":1.0,
"C10_0_3OXOACP_C":-1.0,
"cpd00005_C":-1.0,
"cpd00006_C":1.0
},
"lower_bound":0.0,
"upper_bound":1000.0,
"gene_reaction_rule":"ACIAD0871"
},
{
"id":"GLUTRNAREDUCT_RXN",
"name":"glutamyl trna reductase",
"metabolites":{
"CHARGED_GLT_TRNAS_C":-1.0,
"GLT_TRNAS_C":1.0,
"cpd00005_C":-1.0,
"cpd00006_C":1.0,
"cpd02345_C":1.0
},
"lower_bound":-1000.0,
"upper_bound":1000.0,
"gene_reaction_rule":"ACIAD2900"
},
{
"id":"_R_R_BUTANEDIOL_DEHYDROGENASE_RXN",
"name":"(r,r)-butanediol dehydrogenase",
"metabolites":{
"cpd00003_C":-1.0,
"cpd00004_C":1.0,
"cpd00361_C":1.0,
"cpd01947_C":-1.0
},
"lower_bound":-1000.0,
"upper_bound":1000.0,
"gene_reaction_rule":"ACIAD1021"
},
{
"id":"ENOYL_COA_DELTA_ISOM_RXN",
"name":"dodecenoyl-coa delta-isomerase",
"metabolites":{
"cpd01887_C":-1.0,
"cpd02060_C":1.0
},
"lower_bound":-1000.0,
"upper_bound":1000.0,
"gene_reaction_rule":"ACIAD0335"
},
{
"id":"UDPGLUCEPIM_RXN",
"name":"udp-glucose 4-epimerase",
"metabolites":{
"cpd00026_C":-1.0,
"cpd00043_C":1.0
},
"lower_bound":-1000.0,
"upper_bound":1000.0,
"gene_reaction_rule":"ACIAD1891 or ACIAD0102 or ACIAD0096"
},
{
"id":"EXF_UREA_E",
"name":"exchange flux of urea",
"metabolites":{
"cpd00073_E":-1.0
},
"lower_bound":-0.0,
"upper_bound":1000.0,
"gene_reaction_rule":""
},
{
"id":"EXF_CPDIM_6116_E",
"name":"exchange flux of 3,4-dihydroxyphenylpropionate",
"metabolites":{
"cpd07333_E":-1.0
},
"lower_bound":-0.0,
"upper_bound":1000.0,
"gene_reaction_rule":""
},
{
"id":"EXF_DEOXYINOSINE_E",
"name":"exchange flux of deoxyinosine",
"metabolites":{
"cpd03279_E":-1.0
},
"lower_bound":-0.0,
"upper_bound":1000.0,
"gene_reaction_rule":""
},
{
"id":"C6_0_COATRANSFERASE_RXN",
"name":"acyl coa transferase (c6:0)",
"metabolites":{
"cpd00002_C":-1.0,
"cpd00010_C":-1.0,
"cpd00012_C":1.0,
"cpd00018_C":1.0,
"cpd01113_C":-1.0,
"cpd03124_C":1.0
},
"lower_bound":0.0,
"upper_bound":1000.0,
"gene_reaction_rule":"ACIAD1818 or ACIAD0235 or ACIAD1724 or ACIAD2106"
},
{
"id":"RXNIM_6205",
"name":"benzylacetate esterase",
"metabolites":{
"cpd00001_C":-1.0,
"cpd00029_C":1.0,
"cpd00435_C":1.0,
"cpd11193_C":-1.0
},
"lower_bound":0.0,
"upper_bound":1000.0,
"gene_reaction_rule":"ACIAD1428"
},
{
"id":"CARBPSYN_RXN",
"name":"carbamoyl-phosphate synthase (glutamine-hydrolysing)",
"metabolites":{
"cpd00001_C":-1.0,
"cpd00002_C":-2.0,
"cpd00008_C":2.0,
"cpd00009_C":1.0,
"cpd00023_C":1.0,
"cpd00053_C":-1.0,
"cpd00067_C":-1.0,
"cpd00146_C":1.0,
"cpd00242_C":-1.0
},
"lower_bound":0.0,
"upper_bound":1000.0,
"gene_reaction_rule":"ACIAD2861 and ACIAD2860"
},
{
"id":"RXNIM_6207",
"name":"salicyl alcohol dehydrogenase",
"metabolites":{
"cpd00003_C":-1.0,
"cpd00004_C":1.0,
"cpd01553_C":-1.0,
"cpd03709_C":1.0
},
"lower_bound":-1000.0,
"upper_bound":1000.0,
"gene_reaction_rule":"ACIAD1429 or ACIAD1578"
},
{
"id":"RXNIM_6206",
"name":"ethylsalicylate esterase",
"metabolites":{
"cpd00001_C":-1.0,
"cpd00363_C":1.0,
"cpd00599_C":1.0,
"cpd16657_C":-1.0
},
"lower_bound":0.0,
"upper_bound":1000.0,
"gene_reaction_rule":"ACIAD1426"
},
{
"id":"DTDPGLUCOSEPP_RXN",
"name":"glucose-1-phosphate thymidylyltransferase",
"metabolites":{
"cpd00012_C":1.0,
"cpd00089_C":-1.0,
"cpd00357_C":-1.0,
"cpd00626_C":1.0
},
"lower_bound":-1000.0,
"upper_bound":1000.0,
"gene_reaction_rule":"ACIAD0079"
},
{
"id":"ACYL_ACP_LPS_ASSEMBLY_RXN",
"name":"generic acyl acp of lps assembly",
"metabolites":{
"ACYL_ACP_LPS_C":1.0,
"C12_0_ACP_C":-0.62,
"C16_0_ACP_C":-0.18,
"C18_0_ACP_C":-0.1,
"C18_1_C9__ACP_C":-0.1
},
"lower_bound":-1000.0,
"upper_bound":1000.0,
"gene_reaction_rule":""
},
{
"id":"DGMP_NUCLEOTIDASE_RXN",
"name":"dgmp nucleotidase",
"metabolites":{
"cpd00001_C":-1.0,
"cpd00009_C":1.0,
"cpd00277_C":1.0,
"cpd00296_C":-1.0
},
"lower_bound":0.0,
"upper_bound":1000.0,
"gene_reaction_rule":"ACIAD0015 or ACIAD1227"
},
{
"id":"RXNIM_3886",
"name":"glucose dehydrogenase [pyrroloquinoline-quinone]",
"metabolites":{
"cpd00027_P":-1.0,
"cpd00097_P":-1.0,
"cpd00170_P":1.0,
"cpd00986_P":1.0
},
"lower_bound":0.0,
"upper_bound":1000.0,
"gene_reaction_rule":"ACIAD2983"
},
{
"id":"RXNIM_3991",
"name":"malonate decarboxylase",
"metabolites":{
"cpd00011_C":1.0,
"cpd00029_C":1.0,
"cpd00308_C":-1.0
},
"lower_bound":0.0,
"upper_bound":1000.0,
"gene_reaction_rule":"ACIAD1757 and ACIAD1756 and ACIAD1755 and ACIAD1753 and ACIAD1754 and ACIAD1758 and ACIAD1759"
},
{
"id":"RXNIM_6209",
"name":"4-hydroxybenzylacetate esterase",
"metabolites":{
"cpd00001_C":-1.0,
"cpd00029_C":1.0,
"cpd15378_C":1.0,
"cpd16658_C":-1.0
},
"lower_bound":0.0,
"upper_bound":1000.0,
"gene_reaction_rule":"ACIAD1428"
},
{
"id":"RXNIM_6208",
"name":"salicylacetate esterase",
"metabolites":{
"cpd00001_C":-1.0,
"cpd00029_C":1.0,
"cpd01006_C":-1.0,
"cpd01553_C":1.0
},
"lower_bound":0.0,
"upper_bound":1000.0,
"gene_reaction_rule":"ACIAD1428"
},
{
"id":"1_ACYLGLYCEROL_3_P_ACYLTRANSFER_TAG_RXN",
"name":"glycerol-3-phosphate acyltransferase",
"metabolites":{
"ACP_C":1.0,
"ACYL_ACP_TAG_C":-1.0,
"ACYL_SN_GLYCEROL_3P_TAG_C":-1.0,
"L_PHOSPHATIDATE_TAG_C":1.0
},
"lower_bound":0.0,
"upper_bound":1000.0,
"gene_reaction_rule":"ACIAD3232"
},
{
"id":"C8_0_CARBOXY3OXOCOA_THIOLASE_RXN",
"name":"dicarboxylate 3oxoacyl-coa thiolase (c8:0)",
"metabolites":{
"cpd00010_C":-1.0,
"cpd00022_C":1.0,
"cpd09842_C":1.0,
"cpd16651_C":-1.0
},
"lower_bound":-1000.0,
"upper_bound":1000.0,
"gene_reaction_rule":"ACIAD1689"
},
{
"id":"GLUTCYSLIG_RXN",
"name":"glutamate--cysteine ligase",
"metabolites":{
"cpd00002_C":-1.0,
"cpd00008_C":1.0,
"cpd00009_C":1.0,
"cpd00023_C":-1.0,
"cpd00084_C":-1.0,
"cpd00506_C":1.0
},
"lower_bound":0.0,
"upper_bound":1000.0,
"gene_reaction_rule":"ACIAD3549"
},
{
"id":"KDO_8PSYNTH_RXN",
"name":"2-dehydro-3-deoxyphosphooctonate aldolase",
"metabolites":{
"cpd00001_C":-1.0,
"cpd00009_C":1.0,
"cpd00061_C":-1.0,
"cpd00817_C":-1.0,
"cpd02730_C":1.0
},
"lower_bound":0.0,
"upper_bound":1000.0,
"gene_reaction_rule":"ACIAD2002"
},
{
"id":"UREASE_RXN",
"name":"urease",
"metabolites":{
"cpd00001_C":-1.0,
"cpd00011_C":1.0,
"cpd00013_C":2.0,
"cpd00073_C":-1.0
},
"lower_bound":0.0,
"upper_bound":1000.0,
"gene_reaction_rule":"ACIAD1089 and ACIAD1090 and ACIAD1091"
},
{
"id":"CYCLOHEXADIENYL_DEHYDROGENASE_RXN",
"name":"cyclohexadienyl dehydrogenase",
"metabolites":{
"cpd00003_C":-1.0,
"cpd00004_C":1.0,
"cpd00011_C":1.0,
"cpd00069_C":1.0,
"cpd00616_C":-1.0
},
"lower_bound":0.0,
"upper_bound":1000.0,
"gene_reaction_rule":"ACIAD2222"
},
{
"id":"C16_1_C7_ACP_HYDROLASE_RXN",
"name":"acyl-acp hydrolase (c16:1(c7))",
"metabolites":{
"ACP_C":1.0,
"C16_1_C7__ACP_C":-1.0,
"cpd00001_C":-1.0,
"cpd34461_C":1.0
},
"lower_bound":0.0,
"upper_bound":1000.0,
"gene_reaction_rule":""
},
{
"id":"TRANS_RXN_BENZYL_ALCOHOL",
"name":"transport of benzyl alcohol [extraorganism;cytosol](passive)",
"metabolites":{
"cpd00435_C":1.0,
"cpd00435_E":-1.0
},
"lower_bound":0.0,
"upper_bound":1000.0,
"gene_reaction_rule":""
},
{
"id":"C12_0_3OXOCOA_THIOLASE_RXN",
"name":"3oxoacyl-coa thiolase (c12:0)",
"metabolites":{
"cpd00010_C":-1.0,
"cpd00022_C":1.0,
"cpd03117_C":-1.0,
"cpd03128_C":1.0
},
"lower_bound":-1000.0,
"upper_bound":1000.0,
"gene_reaction_rule":"ACIAD0334 or ACIAD1450 or ACIAD1706 or ACIAD1977 or ACIAD1689 or ACIAD2988 or ACIAD1569"
},
{
"id":"1TRANSKETO_RXN",
"name":"transketolase (ribose-5p)",
"metabolites":{
"cpd00101_C":-1.0,
"cpd00102_C":1.0,
"cpd00198_C":-1.0,
"cpd00238_C":1.0
},
"lower_bound":-1000.0,
"upper_bound":1000.0,
"gene_reaction_rule":"ACIAD2035"
},
{
"id":"EXF_CYTOSINE_E",
"name":"exchange flux of cytosine",
"metabolites":{
"cpd00307_E":-1.0
},
"lower_bound":-0.0,
"upper_bound":1000.0,
"gene_reaction_rule":""
},
{
"id":"PEPSYNTH_RXN",
"name":"pyruvate,water dikinase",
"metabolites":{
"cpd00001_C":-1.0,
"cpd00002_C":-1.0,
"cpd00009_C":1.0,
"cpd00018_C":1.0,
"cpd00020_C":-1.0,
"cpd00061_C":1.0
},
"lower_bound":0.0,
"upper_bound":1000.0,
"gene_reaction_rule":"ACIAD2423"
},
{
"id":"GLUTAMIDOTRANS_RXN",
"name":"imidazole glycerol phosphate synthase",
"metabolites":{
"cpd00023_C":1.0,
"cpd00053_C":-1.0,
"cpd02843_C":1.0,
"cpd02851_C":1.0,
"cpd02991_C":-1.0
},
"lower_bound":0.0,
"upper_bound":1000.0,
"gene_reaction_rule":"ACIAD3404 and ACIAD3396"
},
{
"id":"C18_0_2ENOYLACP_REDUCTASE_RXN",
"name":"2enoyl-acp reductase (c18:0)",
"metabolites":{
"C18_0_2ENOYLACP_C":-1.0,
"C18_0_ACP_C":1.0,
"cpd00003_C":1.0,
"cpd00004_C":-1.0
},
"lower_bound":0.0,
"upper_bound":1000.0,
"gene_reaction_rule":"ACIAD3116"
},
{
"id":"3_DEHYDROQUINATE_DEHYDRATASE_CATABOLIC_RXN",
"name":"3-dehydroquinate dehydratase",
"metabolites":{
"cpd00001_P":1.0,
"cpd00699_P":-1.0,
"cpd01716_P":1.0
},
"lower_bound":-1000.0,
"upper_bound":1000.0,
"gene_reaction_rule":"ACIAD1713"
},
{
"id":"EXF_CPD_592_E",
"name":"exchange flux of 4-guanidinobutanoate",
"metabolites":{
"cpd00762_E":-1.0
},
"lower_bound":-0.0,
"upper_bound":1000.0,
"gene_reaction_rule":""
},
{
"id":"EXF_CPDIM_6112_E",
"name":"exchange flux of ethylsalicylate",
"metabolites":{
"cpd16657_E":-1.0
},
"lower_bound":-0.0,
"upper_bound":1000.0,
"gene_reaction_rule":""
},
{
"id":"SPONTPRO_RXN",
"name":"l-glutamate 5-semialdehyde dehydratase",
"metabolites":{
"cpd00001_C":1.0,
"cpd00858_C":-1.0,
"cpd02431_C":1.0
},
"lower_bound":0.0,
"upper_bound":1000.0,
"gene_reaction_rule":"SPONT"
},
{
"id":"EXF_URACIL_E",
"name":"exchange flux of uracil",
"metabolites":{
"cpd00092_E":-1.0
},
"lower_bound":-0.0,
"upper_bound":1000.0,
"gene_reaction_rule":""
},
{
"id":"CHORISMATEMUT_RXN",
"name":"chorismate mutase",
"metabolites":{
"cpd00216_C":-1.0,
"cpd00219_C":1.0
},
"lower_bound":-1000.0,
"upper_bound":1000.0,
"gene_reaction_rule":"ACIAD2223"
},
{
"id":"2_OCTAPRENYL_6_OHPHENOL_METHY_RXN",
"name":"2-octaprenyl-6-hydroxyphenol methylase",
"metabolites":{
"cpd00017_C":-1.0,
"cpd00019_C":1.0,
"cpd03445_C":-1.0,
"cpd03446_C":1.0
},
"lower_bound":0.0,
"upper_bound":1000.0,
"gene_reaction_rule":"ACIAD0044"
},
{
"id":"SHIKIMATE_PQQ_RXN",
"name":"shikimate dehydrogenase, pqq",
"metabolites":{
"cpd00097_P":-1.0,
"cpd00383_P":-1.0,
"cpd00986_P":1.0,
"cpd01716_P":1.0
},
"lower_bound":-1000.0,
"upper_bound":1000.0,
"gene_reaction_rule":"ACIAD1716"
},
{
"id":"EXF_CYTIDINE_E",
"name":"exchange flux of cytidine",
"metabolites":{
"cpd00367_E":-1.0
},
"lower_bound":-0.0,
"upper_bound":1000.0,
"gene_reaction_rule":""
},
{
"id":"TRANS_RXN_3_KETOBUTYRATE",
"name":"transport of acetoacetate [extraorganism;cytosol](passive)",
"metabolites":{
"cpd00142_C":1.0,
"cpd00142_E":-1.0
},
"lower_bound":0.0,
"upper_bound":1000.0,
"gene_reaction_rule":""
},
{
"id":"CDPKIN_RXN",
"name":"cdp kinase",
"metabolites":{
"cpd00002_C":-1.0,
"cpd00008_C":1.0,
"cpd00052_C":1.0,
"cpd00096_C":-1.0
},
"lower_bound":-1000.0,
"upper_bound":1000.0,
"gene_reaction_rule":"ACIAD0556"
},
{
"id":"4_1_2_41_RXN",
"name":"vanillin synthase",
"metabolites":{
"cpd00022_C":1.0,
"cpd00561_C":1.0,
"cpd04511_C":-1.0
},
"lower_bound":-1000.0,
"upper_bound":1000.0,
"gene_reaction_rule":"ACIAD1726"
},
{
"id":"GLUTAMATESYN_RXN",
"name":"glutamate synthase (nadph)",
"metabolites":{
"cpd00005_C":-1.0,
"cpd00006_C":1.0,
"cpd00023_C":2.0,
"cpd00024_C":-1.0,
"cpd00053_C":-1.0
},
"lower_bound":0.0,
"upper_bound":1000.0,
"gene_reaction_rule":"(ACIAD3350 and ACIAD3349) or (ACIAD2525 and ACIAD2526 and ACIAD2527)"
},
{
"id":"TRANS_RXN_C8_0",
"name":"transport of octanoate [extraorganism;cytosol](secondary symport)",
"metabolites":{
"cpd00067_C":1.0,
"cpd00067_P":-1.0,
"cpd03846_C":1.0,
"cpd03846_E":-1.0
},
"lower_bound":-1000.0,
"upper_bound":1000.0,
"gene_reaction_rule":"ACIAD0835 or ACIAD1427 or ACIAD2788 or ACIAD3392"
},
{
"id":"CMPKI_RXN",
"name":"cmp kinase",
"metabolites":{
"cpd00002_C":-1.0,
"cpd00008_C":1.0,
"cpd00046_C":-1.0,
"cpd00096_C":1.0
},
"lower_bound":-1000.0,
"upper_bound":1000.0,
"gene_reaction_rule":"ACIAD2348"
},
{
"id":"PYRNUTRANSHYDROGEN_RXN",
"name":"nad(p)(+) transhydrogenase (b-specific)",
"metabolites":{
"cpd00003_C":-1.0,
"cpd00004_C":1.0,
"cpd00005_C":-1.0,
"cpd00006_C":1.0
},
"lower_bound":0.0,
"upper_bound":1000.0,
"gene_reaction_rule":"ACIAD2274"
},
{
"id":"2TRANSKETO_RXN",
"name":"transketolase (erythrose-4p)",
"metabolites":{
"cpd00102_C":1.0,
"cpd00198_C":-1.0,
"cpd00236_C":-1.0,
"cpd15394_C":1.0
},
"lower_bound":-1000.0,
"upper_bound":1000.0,
"gene_reaction_rule":"ACIAD2035"
},
{
"id":"GLYCOLALD_DEHYDROG_RXN",
"name":"glycolaldehyde dehydrogenase",
"metabolites":{
"cpd00001_C":-1.0,
"cpd00003_C":-1.0,
"cpd00004_C":1.0,
"cpd00139_C":1.0,
"cpd00229_C":-1.0
},
"lower_bound":-1000.0,
"upper_bound":1000.0,
"gene_reaction_rule":"ACIAD2018 or ACIAD2542"
},
{
"id":"DIACYLGLYCEROL_O_ACYLTRANSFERASE_RXN",
"name":"diacylglycerol o-acyltransferase",
"metabolites":{
"ACYL_COA_TAG_C":-1.0,
"cpd00010_C":1.0,
"cpd11423_C":-1.0,
"cpd11677_C":1.0
},
"lower_bound":0.0,
"upper_bound":1000.0,
"gene_reaction_rule":"ACIAD0832"
},
{
"id":"RXN_3641",
"name":"beta-ketoadipyl coa thiolase",
"metabolites":{
"cpd00010_C":-1.0,
"cpd00022_C":1.0,
"cpd00078_C":1.0,
"cpd01507_C":-1.0
},
"lower_bound":0.0,
"upper_bound":1000.0,
"gene_reaction_rule":"ACIAD1450 or ACIAD1689 or ACIAD1706"
},
{
"id":"NAD_KIN_RXN",
"name":"nad(+) kinase",
"metabolites":{
"cpd00002_C":-1.0,
"cpd00003_C":-1.0,
"cpd00006_C":1.0,
"cpd00008_C":1.0
},
"lower_bound":-1000.0,
"upper_bound":1000.0,
"gene_reaction_rule":"ACIAD2231"
},
{
"id":"C4_0_3OXOACP_REDUCTASE_RXN",
"name":"3oxoacyl-acp reductase (c4:0)",
"metabolites":{
"ACETOACETYL_S_ACP_C":-1.0,
"C4_0_3HYDROXYACP_C":1.0,
"cpd00005_C":-1.0,
"cpd00006_C":1.0
},
"lower_bound":0.0,
"upper_bound":1000.0,
"gene_reaction_rule":"ACIAD0871"
},
{
"id":"CYTIDEAM2_RXN",
"name":"cytidine deaminase",
"metabolites":{
"cpd00001_C":-1.0,
"cpd00013_C":1.0,
"cpd00249_C":1.0,
"cpd00367_C":-1.0
},
"lower_bound":0.0,
"upper_bound":1000.0,
"gene_reaction_rule":"ACIAD3297 or ACIAD0204"
},
{
"id":"C12_0_3HYDROXYACP_DEHYDRATASE_RXN",
"name":"3hydroxyacyl-acp dehydratase (c12:0)",
"metabolites":{
"3_OHLAUROYL_ACP_C":-1.0,
"C12_0_2ENOYLACP_C":1.0,
"cpd00001_C":1.0
},
"lower_bound":0.0,
"upper_bound":1000.0,
"gene_reaction_rule":"ACIAD1381"
},
{
"id":"C8_0_3OXOACP_REDUCTASE_RXN",
"name":"3oxoacyl-acp reductase (c8:0)",
"metabolites":{
"C8_0_3HYDROXYACP_C":1.0,
"C8_0_3OXOACP_C":-1.0,
"cpd00005_C":-1.0,
"cpd00006_C":1.0
},
"lower_bound":0.0,
"upper_bound":1000.0,
"gene_reaction_rule":"ACIAD0871"
},
{
"id":"2_DEHYDROPANTOATE_REDUCT_RXN",
"name":"2-dehydropantoate 2-reductase",
"metabolites":{
"cpd00005_C":1.0,
"cpd00006_C":-1.0,
"cpd00408_C":-1.0,
"cpd00712_C":1.0
},
"lower_bound":-1000.0,
"upper_bound":1000.0,
"gene_reaction_rule":"ACIAD3102"
},
{
"id":"RXN_1241",
"name":"coniferyl-aldehyde dehydrogenase",
"metabolites":{
"cpd00001_C":-1.0,
"cpd00003_C":-1.0,
"cpd00004_C":1.0,
"cpd01059_C":1.0,
"cpd01733_C":-1.0
},
"lower_bound":-1000.0,
"upper_bound":1000.0,
"gene_reaction_rule":"ACIAD0503"
},
{
"id":"ACETOLACTREDUCTOISOM_RXN",
"name":"ketol-acid reductoisomerase",
"metabolites":{
"cpd00005_C":-1.0,
"cpd00006_C":1.0,
"cpd00668_C":-1.0,
"cpd02498_C":1.0
},
"lower_bound":-1000.0,
"upper_bound":1000.0,
"gene_reaction_rule":"ACIAD3102"
},
{
"id":"EXF_CAFFEATE_E",
"name":"exchange flux of caffeate",
"metabolites":{
"cpd00881_E":-1.0
},
"lower_bound":-0.0,
"upper_bound":1000.0,
"gene_reaction_rule":""
},
{
"id":"ARGSUCCINLYA_RXN",
"name":"argininosuccinate lyase",
"metabolites":{
"cpd00051_C":1.0,
"cpd00106_C":1.0,
"cpd02152_C":-1.0
},
"lower_bound":-1000.0,
"upper_bound":1000.0,
"gene_reaction_rule":"ACIAD0283"
},
{
"id":"LIPIDADISACCHARIDESYNTH_RXN",
"name":"lipid-a-disaccharide synthase",
"metabolites":{
"cpd00014_C":1.0,
"cpd02835_C":-1.0,
"cpd02930_C":-1.0,
"cpd03002_C":1.0
},
"lower_bound":0.0,
"upper_bound":1000.0,
"gene_reaction_rule":"ACIAD2324"
},
{
"id":"GLYOHMETRANS_RXN",
"name":"glycine hydroxymethyltransferase",
"metabolites":{
"cpd00001_C":-1.0,
"cpd00033_C":-1.0,
"cpd00054_C":1.0,
"cpd00087_C":1.0,
"cpd00125_C":-1.0
},
"lower_bound":-1000.0,
"upper_bound":1000.0,
"gene_reaction_rule":"ACIAD2255"
},
{
"id":"UDPNACETYLGLUCOSAMENOLPYRTRANS_RXN",
"name":"udp-n-acetylglucosamine 1-carboxyvinyltransferase",
"metabolites":{
"cpd00009_C":1.0,
"cpd00037_C":-1.0,
"cpd00061_C":-1.0,
"cpd02820_C":1.0
},
"lower_bound":0.0,
"upper_bound":1000.0,
"gene_reaction_rule":"ACIAD0660"
},
{
"id":"URACIL_PRIBOSYLTRANS_RXN",
"name":"uracil phosphoribosyltransferase",
"metabolites":{
"cpd00012_C":1.0,
"cpd00091_C":1.0,
"cpd00092_C":-1.0,
"cpd00103_C":-1.0
},
"lower_bound":0.0,
"upper_bound":1000.0,
"gene_reaction_rule":"ACIAD0744"
},
{
"id":"3_OCTAPRENYL_4_OHBENZOATE_DECARBOX_RXN",
"name":"octaprenyl-hydroxybenzoate decarboxylase",
"metabolites":{
"cpd00011_C":1.0,
"cpd03443_C":-1.0,
"cpd03444_C":1.0
},
"lower_bound":0.0,
"upper_bound":1000.0,
"gene_reaction_rule":""
},
{
"id":"GLUTKIN_RXN",
"name":"glutamate 5-kinase",
"metabolites":{
"cpd00002_C":-1.0,
"cpd00008_C":1.0,
"cpd00023_C":-1.0,
"cpd02097_C":1.0
},
"lower_bound":0.0,
"upper_bound":1000.0,
"gene_reaction_rule":"ACIAD2560"
},
{
"id":"3PGAREARR_RXN",
"name":"phosphoglycerate mutase",
"metabolites":{
"cpd00169_C":-1.0,
"cpd00482_C":1.0
},
"lower_bound":-1000.0,
"upper_bound":1000.0,
"gene_reaction_rule":"ACIAD0256"
},
{
"id":"EXF_CONIFERYL_ALDEHYDE_E",
"name":"exchange flux of coniferyl aldehyde",
"metabolites":{
"cpd01733_E":-1.0
},
"lower_bound":-0.0,
"upper_bound":1000.0,
"gene_reaction_rule":""
},
{
"id":"RXN_681",
"name":"l-2,4-diaminobutyrate decarboxylase",
"metabolites":{
"cpd00011_C":1.0,
"cpd00726_C":1.0,
"cpd02095_C":-1.0
},
"lower_bound":0.0,
"upper_bound":1000.0,
"gene_reaction_rule":"ACIAD1211"
},
{
"id":"TRANS_RXN_242A",
"name":"transport of homoserine [cytosol;extraorganism](secondary antiport)",
"metabolites":{
"cpd00067_C":1.0,
"cpd00067_P":-1.0,
"cpd00227_C":-1.0,
"cpd00227_E":1.0
},
"lower_bound":0.0,
"upper_bound":1000.0,
"gene_reaction_rule":"ACIAD0173 or ACIAD3166"
},
{
"id":"R221_RXN",
"name":"octanol dehydrogenase",
"metabolites":{
"cpd00003_C":-1.0,
"cpd00004_C":1.0,
"cpd00562_C":-1.0,
"cpd01088_C":1.0
},
"lower_bound":-1000.0,
"upper_bound":1000.0,
"gene_reaction_rule":""
},
{
"id":"PLSYN_RXN",
"name":"generic phospholipid assembly",
"metabolites":{
"L_1_PHOSPHATIDYL_ETHANOLAMINE_C":-0.735,
"L_1_PHOSPHATIDYL_GLYCEROL_C":-0.116,
"PL_W_C":0.8359,
"cpd12801_C":-0.149
},
"lower_bound":-1000.0,
"upper_bound":1000.0,
"gene_reaction_rule":""
},
{
"id":"SUCC_FUM_OXRED_RXN",
"name":"succinate dehydrogenase",
"metabolites":{
"cpd00036_C":-1.0,
"cpd00106_C":1.0,
"cpd15560_C":-1.0,
"cpd15561_C":1.0
},
"lower_bound":-1000.0,
"upper_bound":1000.0,
"gene_reaction_rule":"ACIAD2882 and ACIAD2881 and ACIAD2880 and ACIAD2879"
},
{
"id":"DIHYDRODIPICSYN_RXN",
"name":"dihydrodipicolinate synthase",
"metabolites":{
"cpd00001_C":2.0,
"cpd00020_C":-1.0,
"cpd00346_C":-1.0,
"cpd02120_C":1.0
},
"lower_bound":-1000.0,
"upper_bound":1000.0,
"gene_reaction_rule":"ACIAD3585"
},
{
"id":"UDPACYLGLCNACDEACETYL_RXN",
"name":"udp-3-o-acyl-n-acetylglucosamine deacetylase",
"metabolites":{
"cpd00001_C":-1.0,
"cpd00029_C":1.0,
"cpd02886_C":-1.0,
"cpd03584_C":1.0
},
"lower_bound":0.0,
"upper_bound":1000.0,
"gene_reaction_rule":"ACIAD3510"
},
{
"id":"1_3_1_62_RXN",
"name":"pimeloyl-coa dehydrogenase",
"metabolites":{
"cpd00015_C":-1.0,
"cpd00782_C":-1.0,
"cpd00982_C":1.0,
"cpd04111_C":1.0
},
"lower_bound":0.0,
"upper_bound":1000.0,
"gene_reaction_rule":"ACIAD1693"
},
{
"id":"6_CARBOXYHEXANOATE_COA_LIGASE_RXN",
"name":"6-carboxyhexanoate--coa ligase",
"metabolites":{
"cpd00002_C":-1.0,
"cpd00010_C":-1.0,
"cpd00012_C":1.0,
"cpd00018_C":1.0,
"cpd00782_C":1.0,
"cpd01727_C":-1.0
},
"lower_bound":-1000.0,
"upper_bound":1000.0,
"gene_reaction_rule":"ACIAD1826"
},
{
"id":"EXF_XANTHINE_E",
"name":"exchange flux of xanthine",
"metabolites":{
"cpd00309_E":-1.0
},
"lower_bound":-0.0,
"upper_bound":1000.0,
"gene_reaction_rule":""
},
{
"id":"TRANS_RXNIM_5747_BIS",
"name":"transport of l-proline [extraorganism;cytosol](passive)",
"metabolites":{
"cpd00129_C":1.0,
"cpd00129_E":-1.0
},
"lower_bound":-1000.0,
"upper_bound":1000.0,
"gene_reaction_rule":"ACIAD2280 or ACIAD1036"
},
{
"id":"C12_0_COA_THIOLASE_RXN",
"name":"acyl-coa thiolase (c12:0)",
"metabolites":{
"cpd00001_C":-1.0,
"cpd00010_C":1.0,
"cpd01260_C":-1.0,
"cpd01741_C":1.0
},
"lower_bound":0.0,
"upper_bound":1000.0,
"gene_reaction_rule":"ACIAD0341 or ACIAD3231 or ACIAD1023"
},
{
"id":"EXF_GLN_E",
"name":"exchange flux of l-glutamine",
"metabolites":{
"cpd00053_E":-1.0
},
"lower_bound":-0.0,
"upper_bound":1000.0,
"gene_reaction_rule":""
},
{
"id":"C10_0_COA_THIOLASE_RXN",
"name":"acyl-coa thiolase (c10:0)",
"metabolites":{
"cpd00001_C":-1.0,
"cpd00010_C":1.0,
"cpd01107_C":1.0,
"cpd03128_C":-1.0
},
"lower_bound":0.0,
"upper_bound":1000.0,
"gene_reaction_rule":"ACIAD0341 or ACIAD3231 or ACIAD1023"
},
{
"id":"TRANS_RXN_DEOXYADENOSINE",
"name":"transport of deoxyadenosine [extraorganism;cytosol](secondary symport)",
"metabolites":{
"cpd00067_C":1.0,
"cpd00067_P":-1.0,
"cpd00438_C":1.0,
"cpd00438_E":-1.0
},
"lower_bound":0.0,
"upper_bound":1000.0,
"gene_reaction_rule":"ACIAD2389"
},
{
"id":"MUCONOLACTONE_DELTA_ISOMERASE_RXN",
"name":"muconolactone delta-isomerase",
"metabolites":{
"cpd02255_C":1.0,
"cpd02536_C":-1.0
},
"lower_bound":-1000.0,
"upper_bound":1000.0,
"gene_reaction_rule":"ACIAD1447"
},
{
"id":"ADOMET_DMK_METHYLTRANSFER_RXN",
"name":"s-adenosylmethione:2-demethylmenaquinol methyltransferase",
"metabolites":{
"cpd00017_C":-1.0,
"cpd00019_C":1.0,
"cpd15352_C":-1.0,
"cpd25799_C":1.0
},
"lower_bound":0.0,
"upper_bound":1000.0,
"gene_reaction_rule":"ACIAD2118 or ACIAD0384 or ACIAD1391"
},
{
"id":"PHEAMINOTRANS_RXN",
"name":"phenylalanine aminotransferase",
"metabolites":{
"cpd00023_C":-1.0,
"cpd00024_C":1.0,
"cpd00066_C":1.0,
"cpd00143_C":-1.0
},
"lower_bound":-1000.0,
"upper_bound":1000.0,
"gene_reaction_rule":"ACIAD2574 or ACIAD0597 or ACIAD0112 or ACIAD0664"
},
{
"id":"RXNIM_3878",
"name":"acetoin dehydrogenase",
"metabolites":{
"cpd00003_C":-1.0,
"cpd00004_C":1.0,
"cpd00010_C":-1.0,
"cpd00022_C":1.0,
"cpd00067_C":1.0,
"cpd00071_C":1.0,
"cpd00361_C":-1.0
},
"lower_bound":0.0,
"upper_bound":1000.0,
"gene_reaction_rule":"(ACIAD1017 and ACIAD1018) and ACIAD1019 and (ACIAD1020 or ACIAD2874)"
},
{
"id":"RXN0_299",
"name":"taurine dioxygenase",
"metabolites":{
"cpd00007_C":-1.0,
"cpd00011_C":1.0,
"cpd00024_C":-1.0,
"cpd00036_C":1.0,
"cpd00081_C":1.0,
"cpd00210_C":-1.0,
"cpd04122_C":1.0
},
"lower_bound":0.0,
"upper_bound":1000.0,
"gene_reaction_rule":"ACIAD1592 or ACIAD1600"
},
{
"id":"ADENODEAMIN_RXN",
"name":"adenosine deaminase",
"metabolites":{
"cpd00001_C":-1.0,
"cpd00013_C":1.0,
"cpd00182_C":-1.0,
"cpd00246_C":1.0
},
"lower_bound":0.0,
"upper_bound":1000.0,
"gene_reaction_rule":"ACIAD1245"
},
{
"id":"RXNIM_3877",
"name":"transport of beta-d-glucose [extraorganism;periplasm](passive)",
"metabolites":{
"cpd00027_E":-1.0,
"cpd00027_P":1.0
},
"lower_bound":-1000.0,
"upper_bound":1000.0,
"gene_reaction_rule":"ACIAD2984 or ACIAD1715"
},
{
"id":"TRANS_RXN_SHIKIMATE",
"name":"transport of shikimate [extraorganism;periplasm](passive)",
"metabolites":{
"cpd00383_E":-1.0,
"cpd00383_P":1.0
},
"lower_bound":-1000.0,
"upper_bound":1000.0,
"gene_reaction_rule":"ACIAD1715 or ACIAD2984"
},
{
"id":"CTPSYN_RXN",
"name":"ctp synthase",
"metabolites":{
"cpd00001_C":-1.0,
"cpd00002_C":-1.0,
"cpd00008_C":1.0,
"cpd00009_C":1.0,
"cpd00023_C":1.0,
"cpd00052_C":1.0,
"cpd00053_C":-1.0,
"cpd00062_C":-1.0
},
"lower_bound":0.0,
"upper_bound":1000.0,
"gene_reaction_rule":"ACIAD2003"
},
{
"id":"TRANS_RXN_DEOXYINOSINE",
"name":"transport of deoxyinosine [extraorganism;cytosol](secondary symport)",
"metabolites":{
"cpd00067_C":1.0,
"cpd00067_P":-1.0,
"cpd03279_C":1.0,
"cpd03279_E":-1.0
},
"lower_bound":0.0,
"upper_bound":1000.0,
"gene_reaction_rule":"ACIAD2389"
},
{
"id":"ATP_PYROPHOSPHATASE_RXN",
"name":"atp pyrophosphatase",
"metabolites":{
"cpd00001_C":-1.0,
"cpd00002_C":-1.0,
"cpd00012_C":1.0,
"cpd00018_C":1.0
},
"lower_bound":0.0,
"upper_bound":1000.0,
"gene_reaction_rule":"ACIAD3065"
},
{
"id":"PSERTRANSAMPYR_RXN",
"name":"phosphohydroxythreonine transaminase",
"metabolites":{
"cpd00023_C":-1.0,
"cpd00024_C":1.0,
"cpd03606_C":-1.0,
"cpd03607_C":1.0
},
"lower_bound":-1000.0,
"upper_bound":1000.0,
"gene_reaction_rule":"ACIAD2647"
},
{
"id":"EXF_C8_0_E",
"name":"exchange flux of octanoate",
"metabolites":{
"cpd03846_E":-1.0
},
"lower_bound":-0.0,
"upper_bound":1000.0,
"gene_reaction_rule":""
},
{
"id":"2_OCTAPRENYL_METHOXY_BENZOQ_METH_RXN",
"name":"2-octaprenyl-6-methoxy-benzoquinol methylase",
"metabolites":{
"cpd00017_C":-1.0,
"cpd00019_C":1.0,
"cpd03447_C":-1.0,
"cpd03448_C":1.0
},
"lower_bound":0.0,
"upper_bound":1000.0,
"gene_reaction_rule":"ACIAD0384"
},
{
"id":"ALLANTOICASE_RXN",
"name":"allantoicase",
"metabolites":{
"cpd00001_C":-1.0,
"cpd00073_C":1.0,
"cpd00388_C":-1.0,
"cpd00465_C":1.0
},
"lower_bound":-1000.0,
"upper_bound":1000.0,
"gene_reaction_rule":"ACIAD3541"
},
{
"id":"ACYL_OH_WAX_ASSEMBLY_RXN",
"name":"generic acyl-alcohol of wax assembly",
"metabolites":{
"ACYL_OH_WAX_C":1.0,
"cpd00613_C":-0.052,
"cpd16628_C":-0.475,
"cpd16639_C":-0.473
},
"lower_bound":-1000.0,
"upper_bound":1000.0,
"gene_reaction_rule":""
},
{
"id":"DLACTDEHYDROGNAD_RXN",
"name":"d-lactate dehydrogenase, nad",
"metabolites":{
"cpd00003_C":-1.0,
"cpd00004_C":1.0,
"cpd00020_C":1.0,
"cpd00221_C":-1.0
},
"lower_bound":0.0,
"upper_bound":1000.0,
"gene_reaction_rule":"ACIAD0109"
},
{
"id":"THYMIDYLATE_5_APOST_PHOSPHATASE_RXN",
"name":"dtmp nucleotidase",
"metabolites":{
"cpd00001_C":-1.0,
"cpd00009_C":1.0,
"cpd00184_C":1.0,
"cpd00298_C":-1.0
},
"lower_bound":0.0,
"upper_bound":1000.0,
"gene_reaction_rule":"ACIAD0015 or ACIAD1227"
},
{
"id":"EXF_SO3_E",
"name":"exchange flux of so3-2",
"metabolites":{
"cpd00081_E":-1.0
},
"lower_bound":-0.0,
"upper_bound":1000.0,
"gene_reaction_rule":""
},
{
"id":"BENZALDEHYDE_DEHYDROGENASE_NAD_RXN",
"name":"benzaldehyde dehydrogenase (nad+)",
"metabolites":{
"cpd00001_C":-1.0,
"cpd00003_C":-1.0,
"cpd00004_C":1.0,
"cpd00153_C":1.0,
"cpd00225_C":-1.0
},
"lower_bound":-1000.0,
"upper_bound":1000.0,
"gene_reaction_rule":"ACIAD1430 or ACIAD1725"
},
{
"id":"C12_0_3OXOACP_REDUCTASE_RXN",
"name":"3oxoacyl-acp reductase (c12:0)",
"metabolites":{
"3_OHLAUROYL_ACP_C":1.0,
"C12_0_3OXOACP_C":-1.0,
"cpd00005_C":-1.0,
"cpd00006_C":1.0
},
"lower_bound":0.0,
"upper_bound":1000.0,
"gene_reaction_rule":"ACIAD0871"
},
{
"id":"KDPGALDOL_RXN",
"name":"2-dehydro-3-deoxyphosphogluconate aldolase",
"metabolites":{
"cpd00020_C":1.0,
"cpd00102_C":1.0,
"cpd00945_C":-1.0
},
"lower_bound":-1000.0,
"upper_bound":1000.0,
"gene_reaction_rule":"ACIAD0543"
},
{
"id":"EXF_C6_0_E",
"name":"exchange flux of hexanoate",
"metabolites":{
"cpd01113_E":-1.0
},
"lower_bound":-0.0,
"upper_bound":1000.0,
"gene_reaction_rule":""
},
{
"id":"C8_0_CARBOXY3HYDROXYCOA_DEHYDROGENASE_RXN",
"name":"dicarboxylate hydroxyacyl-coa dehydrogenase (c8:0)",
"metabolites":{
"cpd00003_C":-1.0,
"cpd00004_C":1.0,
"cpd16650_C":-1.0,
"cpd16651_C":1.0
},
"lower_bound":0.0,
"upper_bound":1000.0,
"gene_reaction_rule":"ACIAD1690"
},
{
"id":"C12_0_3OXOACP_SYNTHASE_RXN",
"name":"3oxoacyl-acp synthase (c12:0)",
"metabolites":{
"ACP_C":1.0,
"C10_0_ACP_C":-1.0,
"C12_0_3OXOACP_C":1.0,
"MALONYL_ACP_C":-1.0,
"cpd00011_C":1.0
},
"lower_bound":0.0,
"upper_bound":1000.0,
"gene_reaction_rule":"ACIAD0580 or ACIAD0879 or ACIAD0583 or ACIAD2101"
},
{
"id":"ACETYLORNTRANSAM_RXN",
"name":"acetylornithine aminotransferase",
"metabolites":{
"cpd00023_C":-1.0,
"cpd00024_C":1.0,
"cpd00342_C":1.0,
"cpd00918_C":-1.0
},
"lower_bound":-1000.0,
"upper_bound":1000.0,
"gene_reaction_rule":"ACIAD1231 or ACIAD1284"
},
{
"id":"CHORPYRLY_RXN",
"name":"chorismate pyruvate lyase",
"metabolites":{
"cpd00020_C":1.0,
"cpd00136_C":1.0,
"cpd00216_C":-1.0
},
"lower_bound":0.0,
"upper_bound":1000.0,
"gene_reaction_rule":"ACIAD2456"
},
{
"id":"LIPIDXSYNTHESIS_RXN",
"name":"udp-2,3-diacylglucosamine hydrolase",
"metabolites":{
"cpd00001_C":-1.0,
"cpd00091_C":1.0,
"cpd02835_C":-1.0,
"cpd02930_C":1.0
},
"lower_bound":0.0,
"upper_bound":1000.0,
"gene_reaction_rule":"ACIAD1922"
},
{
"id":"C18_0_COA_DEHYDROGENASE_RXN",
"name":"acyl-coa dehydrogenase (c18:0)",
"metabolites":{
"cpd00015_C":-1.0,
"cpd00327_C":-1.0,
"cpd00982_C":1.0,
"cpd14937_C":1.0
},
"lower_bound":0.0,
"upper_bound":1000.0,
"gene_reaction_rule":"ACIAD0452 or ACIAD3262 or ACIAD2537 or ACIAD2277 or ACIAD1886 or ACIAD1821 or ACIAD1819 or ACIAD1700 or ACIAD1693 or ACIAD1607 or ACIAD1413 or ACIAD0623 or ACIAD0624 or ACIAD0428 or ACIAD0427 or ACIAD1723 or ACIAD2226 or ACIAD3191 or ACIAD1566"
},
{
"id":"C16_1_C7_COATRANSFERASE_RXN",
"name":"acyl coa transferase (c16:1(c7))",
"metabolites":{
"cpd00002_C":-1.0,
"cpd00010_C":-1.0,
"cpd00012_C":1.0,
"cpd00018_C":1.0,
"cpd16627_C":1.0,
"cpd34461_C":-1.0
},
"lower_bound":0.0,
"upper_bound":1000.0,
"gene_reaction_rule":"ACIAD1818 or ACIAD0235 or ACIAD1724 or ACIAD2106"
},
{
"id":"DIHYDROXYMETVALDEHYDRAT_RXN",
"name":"dihydroxy-acid dehydratase",
"metabolites":{
"cpd00001_C":1.0,
"cpd00508_C":1.0,
"cpd02535_C":-1.0
},
"lower_bound":-1000.0,
"upper_bound":1000.0,
"gene_reaction_rule":"ACIAD3636 or ACIAD1266"
},
{
"id":"EXF_C18_0_E",
"name":"exchange flux of octadenoate",
"metabolites":{
"cpd01080_E":-1.0
},
"lower_bound":-0.0,
"upper_bound":1000.0,
"gene_reaction_rule":""
},
{
"id":"PDXA_RXN",
"name":"4- (phosphohydroxy)-l-threonine dehydrogenase",
"metabolites":{
"cpd00003_C":-1.0,
"cpd00004_C":1.0,
"cpd00011_C":1.0,
"cpd03607_C":-1.0,
"cpd08449_C":1.0
},
"lower_bound":0.0,
"upper_bound":1000.0,
"gene_reaction_rule":"ACIAD3010"
},
{
"id":"TRIACYLGLYCEROL_W_RXN",
"name":"conversion to \"mass\" metabolite of triacylglycerol",
"metabolites":{
"TRIACYLGLYCEROL_W_C":0.8205,
"cpd11677_C":-1.0
},
"lower_bound":-1000.0,
"upper_bound":1000.0,
"gene_reaction_rule":""
},
{
"id":"AMIDASE_RXN",
"name":"propanamide amidase",
"metabolites":{
"cpd00001_C":-1.0,
"cpd00013_C":1.0,
"cpd00141_C":1.0,
"cpd27812_C":-1.0
},
"lower_bound":0.0,
"upper_bound":1000.0,
"gene_reaction_rule":"ACIAD1618"
},
{
"id":"C8_0_CARBOXY_COATRANSFERASE_RXN",
"name":"dicarboxylate acyl coa transferase (c8:0)",
"metabolites":{
"cpd00002_C":-1.0,
"cpd00010_C":-1.0,
"cpd00012_C":1.0,
"cpd00018_C":1.0,
"cpd05193_C":-1.0,
"cpd16652_C":1.0
},
"lower_bound":0.0,
"upper_bound":1000.0,
"gene_reaction_rule":"ACIAD1695 and ACIAD1696"
},
{
"id":"GLYCERATE_DEHYDROGENASE_RXN",
"name":"glycerate dehydrogenase",
"metabolites":{
"cpd00003_C":-1.0,
"cpd00004_C":1.0,
"cpd00145_C":1.0,
"cpd00223_C":-1.0
},
"lower_bound":-1000.0,
"upper_bound":1000.0,
"gene_reaction_rule":"ACIAD3278 or ACIAD1301"
},
{
"id":"ANTHRANSYN_RXN",
"name":"anthranilate synthase",
"metabolites":{
"cpd00020_C":1.0,
"cpd00023_C":1.0,
"cpd00053_C":-1.0,
"cpd00093_C":1.0,
"cpd00216_C":-1.0
},
"lower_bound":-1000.0,
"upper_bound":1000.0,
"gene_reaction_rule":"ACIAD0297 and ACIAD2461"
},
{
"id":"PROTOCATECHUATE_3_4_DIOXYGENASE_RXN",
"name":"protocatechuate 3,4-dioxygenase",
"metabolites":{
"cpd00007_C":-1.0,
"cpd00197_C":-1.0,
"cpd00856_C":1.0
},
"lower_bound":0.0,
"upper_bound":1000.0,
"gene_reaction_rule":"ACIAD1711 and ACIAD1712"
},
{
"id":"SIROHEME_FERROCHELAT_RXN",
"name":"sirohydrochlorin ferrochelatase",
"metabolites":{
"cpd00021_C":-1.0,
"cpd00067_C":2.0,
"cpd00557_C":1.0,
"cpd03426_C":-1.0
},
"lower_bound":0.0,
"upper_bound":1000.0,
"gene_reaction_rule":"ACIAD2934"
},
{
"id":"C10_0_2ENOYLACP_REDUCTASE_RXN",
"name":"2enoyl-acp reductase (c10:0)",
"metabolites":{
"C10_0_2ENOYLACP_C":-1.0,
"C10_0_ACP_C":1.0,
"cpd00003_C":1.0,
"cpd00004_C":-1.0
},
"lower_bound":0.0,
"upper_bound":1000.0,
"gene_reaction_rule":"ACIAD3116"
},
{
"id":"EXF_PYRUVATE_E",
"name":"exchange flux of pyruvate",
"metabolites":{
"cpd00020_E":-1.0
},
"lower_bound":-0.0,
"upper_bound":1000.0,
"gene_reaction_rule":""
},
{
"id":"2_3_1_155_RXN",
"name":"3oxoacyl-coa thiolase (c16:0)",
"metabolites":{
"cpd00010_C":-1.0,
"cpd00022_C":1.0,
"cpd01695_C":1.0,
"cpd03114_C":-1.0
},
"lower_bound":-1000.0,
"upper_bound":1000.0,
"gene_reaction_rule":"ACIAD0334 or ACIAD1450 or ACIAD1706 or ACIAD1977 or ACIAD1689 or ACIAD2988 or ACIAD1569"
},
{
"id":"S_ADENMETSYN_RXN",
"name":"methionine adenosyltransferase",
"metabolites":{
"cpd00001_C":-1.0,
"cpd00002_C":-1.0,
"cpd00009_C":1.0,
"cpd00012_C":1.0,
"cpd00017_C":1.0,
"cpd00060_C":-1.0
},
"lower_bound":0.0,
"upper_bound":1000.0,
"gene_reaction_rule":"ACIAD2037"
},
{
"id":"EXF_GLYCEROL_E",
"name":"exchange flux of glycerol",
"metabolites":{
"cpd00100_E":-1.0
},
"lower_bound":-0.0,
"upper_bound":1000.0,
"gene_reaction_rule":""
},
{
"id":"TRANS_RXN_PI_ABC",
"name":"transport of phosphate [extraorganism;cytosol](secondary antiport)",
"metabolites":{
"cpd00001_C":-1.0,
"cpd00002_C":-1.0,
"cpd00008_C":1.0,
"cpd00009_C":2.0,
"cpd00009_E":-1.0
},
"lower_bound":0.0,
"upper_bound":1000.0,
"gene_reaction_rule":"ACIAD1212 and ACIAD1213 and ACIAD1214 and ACIAD1215"
},
{
"id":"TETHYDPICSUCC_RXN",
"name":"2,3,4,5-tetrahydropyridine-2-carboxylate n-succinyltransferase",
"metabolites":{
"cpd00001_C":-1.0,
"cpd00010_C":1.0,
"cpd00078_C":-1.0,
"cpd02465_C":-1.0,
"cpd02724_C":1.0
},
"lower_bound":-1000.0,
"upper_bound":1000.0,
"gene_reaction_rule":"ACIAD2599"
},
{
"id":"C4_0_2ENOYLACP_REDUCTASE_RXN",
"name":"2enoyl-acp reductase (c4:0)",
"metabolites":{
"C4_0_2ENOYLACP_C":-1.0,
"C4_0_ACP_C":1.0,
"cpd00003_C":1.0,
"cpd00004_C":-1.0
},
"lower_bound":0.0,
"upper_bound":1000.0,
"gene_reaction_rule":"ACIAD3116"
},
{
"id":"UDPNACETYLMURAMATEDEHYDROG_RXN",
"name":"udp-n-acetylmuramate dehydrogenase",
"metabolites":{
"cpd00005_C":-1.0,
"cpd00006_C":1.0,
"cpd00773_C":1.0,
"cpd02820_C":-1.0
},
"lower_bound":-1000.0,
"upper_bound":1000.0,
"gene_reaction_rule":"ACIAD1945"
},
{
"id":"TRANS_RXNIM_5855",
"name":"transport of l-aspartate [extraorganism;cytosol](secondary symport)",
"metabolites":{
"cpd00041_C":1.0,
"cpd00041_E":-1.0,
"cpd00067_C":1.0,
"cpd00067_P":-1.0
},
"lower_bound":0.0,
"upper_bound":1000.0,
"gene_reaction_rule":"ACIAD0481"
},
{
"id":"EXF_CPD_302_E",
"name":"exchange flux of d-aspartate",
"metabolites":{
"cpd00320_E":-1.0
},
"lower_bound":-0.0,
"upper_bound":1000.0,
"gene_reaction_rule":""
},
{
"id":"EXF_D_SERINE_E",
"name":"exchange flux of d-serine",
"metabolites":{
"cpd00550_E":-1.0
},
"lower_bound":-0.0,
"upper_bound":1000.0,
"gene_reaction_rule":""
},
{
"id":"RXNIM_3902",
"name":"gluconate permease",
"metabolites":{
"cpd00097_P":1.0,
"cpd00222_C":1.0,
"cpd00222_P":-1.0,
"cpd00986_P":-1.0
},
"lower_bound":0.0,
"upper_bound":1000.0,
"gene_reaction_rule":"ACIAD0544"
},
{
"id":"C14_0_ACP_HYDROLASE_RXN",
"name":"acyl-acp hydrolase (c14:0)",
"metabolites":{
"ACP_C":1.0,
"C14_0_ACP_C":-1.0,
"cpd00001_C":-1.0,
"cpd03847_C":1.0
},
"lower_bound":0.0,
"upper_bound":1000.0,
"gene_reaction_rule":""
},
{
"id":"C10_0_CARBOXYCOA_DEHYDROGENASE_RXN",
"name":"dicarboxylate acyl-coa dehydrogenase (c10:0)",
"metabolites":{
"cpd00015_C":-1.0,
"cpd00982_C":1.0,
"cpd16604_C":1.0,
"cpd16607_C":-1.0
},
"lower_bound":0.0,
"upper_bound":1000.0,
"gene_reaction_rule":"ACIAD1693"
},
{
"id":"NITRITREDUCT_RXN",
"name":"nitrite reductase (nad(p)h)",
"metabolites":{
"cpd00001_C":2.0,
"cpd00003_C":3.0,
"cpd00004_C":-3.0,
"cpd00013_C":1.0,
"cpd00067_C":-5.0,
"cpd00075_C":-1.0
},
"lower_bound":0.0,
"upper_bound":1000.0,
"gene_reaction_rule":"ACIAD1909 and ACIAD1910"
},
{
"id":"PROTOHEMEFERROCHELAT_RXN",
"name":"ferrochelatase",
"metabolites":{
"cpd00021_C":-1.0,
"cpd00028_C":1.0,
"cpd00067_C":2.0,
"cpd01476_C":-1.0
},
"lower_bound":-1000.0,
"upper_bound":1000.0,
"gene_reaction_rule":"ACIAD3255"
},
{
"id":"ISOCHORMAT_RXN",
"name":"isochorismatase",
"metabolites":{
"cpd00001_C":-1.0,
"cpd00020_C":1.0,
"cpd00658_C":-1.0,
"cpd02566_C":1.0
},
"lower_bound":-1000.0,
"upper_bound":1000.0,
"gene_reaction_rule":"ACIAD0028 or ACIAD3473 or ACIAD2775"
},
{
"id":"TRANS_RXN_ETHANOL_AMINE",
"name":"transport of ethanol-amine [extraorganism;cytosol](secondary symport)",
"metabolites":{
"cpd00067_C":1.0,
"cpd00067_P":-1.0,
"cpd00162_C":1.0,
"cpd00162_E":-1.0
},
"lower_bound":0.0,
"upper_bound":1000.0,
"gene_reaction_rule":"ACIAD1458"
},
{
"id":"TRANS_RXNIM_5812",
"name":"transport of l-arginine [extraorganism;cytosol](secondary symport)",
"metabolites":{
"cpd00051_C":1.0,
"cpd00051_E":-1.0,
"cpd00067_C":1.0,
"cpd00067_P":-1.0
},
"lower_bound":0.0,
"upper_bound":1000.0,
"gene_reaction_rule":"ACIAD1417"
},
{
"id":"LYSINE_TRNA_LIGASE_RXN",
"name":"lysine--trna ligase",
"metabolites":{
"CHARGED_LYS_TRNAS_C":1.0,
"LYS_TRNAS_C":-1.0,
"cpd00002_C":-1.0,
"cpd00012_C":1.0,
"cpd00018_C":1.0,
"cpd00039_C":-1.0
},
"lower_bound":0.0,
"upper_bound":1000.0,
"gene_reaction_rule":"ACIAD1069"
},
{
"id":"BRANCHED_CHAINAMINOTRANSFERVAL_RXN",
"name":"valine aminotransferase",
"metabolites":{
"cpd00023_C":-1.0,
"cpd00024_C":1.0,
"cpd00123_C":-1.0,
"cpd00156_C":1.0
},
"lower_bound":-1000.0,
"upper_bound":1000.0,
"gene_reaction_rule":"ACIAD0597"
},
{
"id":"TRANS_RXNIM_5837",
"name":"transport of vanillate [extraorganism;cytosol](secondary symport)",
"metabolites":{
"cpd00067_C":1.0,
"cpd00067_P":-1.0,
"cpd04073_C":1.0,
"cpd04073_E":-1.0
},
"lower_bound":-1000.0,
"upper_bound":1000.0,
"gene_reaction_rule":"ACIAD0982"
},
{
"id":"TRANS_RXN_PYRUVATE",
"name":"transport of pyruvate [extraorganism;cytosol](passive)",
"metabolites":{
"cpd00020_C":1.0,
"cpd00020_E":-1.0
},
"lower_bound":-1000.0,
"upper_bound":1000.0,
"gene_reaction_rule":""
},
{
"id":"EXF_PRO_E",
"name":"exchange flux of l-proline",
"metabolites":{
"cpd00129_E":-1.0
},
"lower_bound":-0.0,
"upper_bound":1000.0,
"gene_reaction_rule":""
},
{
"id":"EXF_CPD_218_E",
"name":"exchange flux of d-methionine",
"metabolites":{
"cpd00637_E":-1.0
},
"lower_bound":-0.0,
"upper_bound":1000.0,
"gene_reaction_rule":""
},
{
"id":"TRANS_RXN_C18_0",
"name":"transport of octadenoate [extraorganism;cytosol](secondary symport)",
"metabolites":{
"cpd00067_C":1.0,
"cpd00067_P":-1.0,
"cpd01080_C":1.0,
"cpd01080_E":-1.0
},
"lower_bound":-1000.0,
"upper_bound":1000.0,
"gene_reaction_rule":"ACIAD0835 or ACIAD1427 or ACIAD2788 or ACIAD3392"
},
{
"id":"ARGININE_TRNA_LIGASE_RXN",
"name":"arginine--trna ligase",
"metabolites":{
"ARG_TRNAS_C":-1.0,
"CHARGED_ARG_TRNAS_C":1.0,
"cpd00002_C":-1.0,
"cpd00012_C":1.0,
"cpd00018_C":1.0,
"cpd00051_C":-1.0
},
"lower_bound":0.0,
"upper_bound":1000.0,
"gene_reaction_rule":"ACIAD0164"
},
{
"id":"TRANS_RXN_C16_0",
"name":"transport of hexadecanoate [extraorganism;cytosol](secondary symport)",
"metabolites":{
"cpd00067_C":1.0,
"cpd00067_P":-1.0,
"cpd00214_C":1.0,
"cpd00214_E":-1.0
},
"lower_bound":-1000.0,
"upper_bound":1000.0,
"gene_reaction_rule":"ACIAD0835 or ACIAD1427 or ACIAD2788 or ACIAD3392"
},
{
"id":"EXF_ARG_E",
"name":"exchange flux of l-arginine",
"metabolites":{
"cpd00051_E":-1.0
},
"lower_bound":-0.0,
"upper_bound":1000.0,
"gene_reaction_rule":""
},
{
"id":"DCDPKIN_RXN",
"name":"dcdp kinase",
"metabolites":{
"cpd00002_C":-1.0,
"cpd00008_C":1.0,
"cpd00356_C":1.0,
"cpd00533_C":-1.0
},
"lower_bound":-1000.0,
"upper_bound":1000.0,
"gene_reaction_rule":"ACIAD0556"
},
{
"id":"HISTIDPHOS_RXN",
"name":"histidinol-phosphatase",
"metabolites":{
"cpd00001_C":-1.0,
"cpd00009_C":1.0,
"cpd00641_C":1.0,
"cpd00807_C":-1.0
},
"lower_bound":0.0,
"upper_bound":1000.0,
"gene_reaction_rule":""
},
{
"id":"WAX_ESTERS_W_RXN",
"name":"conversion to \"mass\" metabolite of wax",
"metabolites":{
"WAX_ESTER_C":-1.0,
"WAX_ESTER_W_C":0.5013
},
"lower_bound":-1000.0,
"upper_bound":1000.0,
"gene_reaction_rule":""
},
{
"id":"EXF_OXYGEN_MOLECULE_E",
"name":"exchange flux of o2",
"metabolites":{
"cpd00007_E":-1.0
},
"lower_bound":-1000.0,
"upper_bound":1000.0,
"gene_reaction_rule":""
},
{
"id":"MALIC_NAD_RXN",
"name":"malate dehydrogenase (oxaloacetate decarboxylating)",
"metabolites":{
"cpd00003_C":-1.0,
"cpd00004_C":1.0,
"cpd00011_C":1.0,
"cpd00020_C":1.0,
"cpd00130_C":-1.0
},
"lower_bound":0.0,
"upper_bound":1000.0,
"gene_reaction_rule":"ACIAD0166"
},
{
"id":"DIHYDROOROT_RXN",
"name":"dihydroorotase",
"metabolites":{
"cpd00001_C":-1.0,
"cpd00282_C":-1.0,
"cpd00343_C":1.0
},
"lower_bound":-1000.0,
"upper_bound":1000.0,
"gene_reaction_rule":"ACIAD1150 or ACIAD1419"
},
{
"id":"MANNPISOM_RXN",
"name":"mannose-6-phosphate isomerase",
"metabolites":{
"cpd00235_C":-1.0,
"cpd15394_C":1.0
},
"lower_bound":-1000.0,
"upper_bound":1000.0,
"gene_reaction_rule":"ACIAD0086"
},
{
"id":"TRANS_RXN_62B",
"name":"transport of glycine [extraorganism;cytosol](secondary symport)",
"metabolites":{
"cpd00033_C":1.0,
"cpd00033_E":-1.0,
"cpd00067_C":1.0,
"cpd00067_P":-1.0
},
"lower_bound":0.0,
"upper_bound":1000.0,
"gene_reaction_rule":"ACIAD0118 or ACIAD2662"
},
{
"id":"3_OXOADIPATE_COA_TRANSFERASE_RXN",
"name":"3-oxoadipate coa-transferase",
"metabolites":{
"cpd00036_C":1.0,
"cpd00078_C":-1.0,
"cpd00630_C":-1.0,
"cpd01507_C":1.0
},
"lower_bound":-1000.0,
"upper_bound":1000.0,
"gene_reaction_rule":"(ACIAD1695 and ACIAD1696) or (ACIAD1448 and ACIAD1449) or (ACIAD1704 and ACIAD1705)"
},
{
"id":"TRANS_RXN_SUC",
"name":"transport of succinate [extraorganism;cytosol](secondary symport)",
"metabolites":{
"cpd00036_C":1.0,
"cpd00036_E":-1.0,
"cpd00067_C":1.0,
"cpd00067_P":-1.0
},
"lower_bound":0.0,
"upper_bound":1000.0,
"gene_reaction_rule":"ACIAD2227"
},
{
"id":"HOMOSERDEHYDROG_RXN",
"name":"homoserine dehydrogenase",
"metabolites":{
"cpd00005_C":-1.0,
"cpd00006_C":1.0,
"cpd00227_C":1.0,
"cpd00346_C":-1.0
},
"lower_bound":-1000.0,
"upper_bound":1000.0,
"gene_reaction_rule":"ACIAD0264"
},
{
"id":"EXF_GLT_E",
"name":"exchange flux of l-glutamate",
"metabolites":{
"cpd00023_E":-1.0
},
"lower_bound":-0.0,
"upper_bound":1000.0,
"gene_reaction_rule":""
},
{
"id":"C14_1_C5_3OXOACP_SYNTHASE_RXN",
"name":"3oxoacyl-acp synthase (c14:1(c5))",
"metabolites":{
"ACP_C":1.0,
"C12_1_C3__ACP_C":-1.0,
"C14_1_C5__3OXOACP_C":1.0,
"MALONYL_ACP_C":-1.0,
"cpd00011_C":1.0
},
"lower_bound":0.0,
"upper_bound":1000.0,
"gene_reaction_rule":"ACIAD0580 or ACIAD0879 or ACIAD0583 or ACIAD2101"
},
{
"id":"RNA_SYNTH_RXN",
"name":"rna synthesis reaction",
"metabolites":{
"RNA_W_C":0.319,
"cpd00002_C":-0.62,
"cpd00008_C":0.4,
"cpd00009_C":0.4,
"cpd00012_C":1.0,
"cpd00038_C":-0.222,
"cpd00052_C":-0.299,
"cpd00062_C":-0.259
},
"lower_bound":0.0,
"upper_bound":1000.0,
"gene_reaction_rule":""
},
{
"id":"PREPHENATEDEHYDROG_RXN",
"name":"prephenate dehydrogenase",
"metabolites":{
"cpd00003_C":-1.0,
"cpd00004_C":1.0,
"cpd00011_C":1.0,
"cpd00219_C":-1.0,
"cpd00868_C":1.0
},
"lower_bound":0.0,
"upper_bound":1000.0,
"gene_reaction_rule":"ACIAD2222"
},
{
"id":"RIBOPHOSPHAT_RXN",
"name":"pyrimidine phosphatase",
"metabolites":{
"cpd00001_C":-1.0,
"cpd00009_C":1.0,
"cpd02720_C":-1.0,
"cpd02882_C":1.0
},
"lower_bound":0.0,
"upper_bound":1000.0,
"gene_reaction_rule":""
},
{
"id":"TRANS_RXN_CONIFERYL_ALDEHYDE",
"name":"transport of coniferyl aldehyde [extraorganism;cytosol](secondary symport)",
"metabolites":{
"cpd00067_C":1.0,
"cpd00067_P":-1.0,
"cpd01733_C":1.0,
"cpd01733_E":-1.0
},
"lower_bound":-1000.0,
"upper_bound":1000.0,
"gene_reaction_rule":"ACIAD1709"
},
{
"id":"EXF_CARNITINE_E",
"name":"exchange flux of l-carnitine",
"metabolites":{
"cpd00266_E":-1.0
},
"lower_bound":-0.0,
"upper_bound":1000.0,
"gene_reaction_rule":""
},
{
"id":"PHOSPHATIDATE_PHOSPHATASE_RXN",
"name":"phosphatidate phosphatase",
"metabolites":{
"L_PHOSPHATIDATE_TAG_C":-1.0,
"cpd00001_C":-1.0,
"cpd00009_C":1.0,
"cpd11423_C":1.0
},
"lower_bound":0.0,
"upper_bound":1000.0,
"gene_reaction_rule":""
},
{
"id":"C14_0_3OXOACP_REDUCTASE_RXN",
"name":"3oxoacyl-acp reductase (c14:0)",
"metabolites":{
"3_OHMYRISTOYL_ACP_C":1.0,
"C14_0_3OXOACP_C":-1.0,
"cpd00005_C":-1.0,
"cpd00006_C":1.0
},
"lower_bound":0.0,
"upper_bound":1000.0,
"gene_reaction_rule":"ACIAD0871"
},
{
"id":"METHENYLTHFCYCLOHYDRO_RXN",
"name":"methenyltetrahydrofolate cyclohydrolase",
"metabolites":{
"cpd00001_C":-1.0,
"cpd00201_C":1.0,
"cpd00347_C":-1.0
},
"lower_bound":-1000.0,
"upper_bound":1000.0,
"gene_reaction_rule":"ACIAD2553 or ACIAD2847"
},
{
"id":"CYTDEAM_RXN",
"name":"cytosine deaminase",
"metabolites":{
"cpd00001_C":-1.0,
"cpd00013_C":1.0,
"cpd00092_C":1.0,
"cpd00307_C":-1.0
},
"lower_bound":0.0,
"upper_bound":1000.0,
"gene_reaction_rule":"ACIAD0137"
},
{
"id":"L_ASPARTATE_OXIDASE_RXN",
"name":"l-aspartate oxidase (producing oxaloacetate)",
"metabolites":{
"cpd00001_C":-1.0,
"cpd00003_C":-1.0,
"cpd00004_C":1.0,
"cpd00013_C":1.0,
"cpd00032_C":1.0,
"cpd00041_C":-1.0,
"cpd00067_C":1.0
},
"lower_bound":0.0,
"upper_bound":1000.0,
"gene_reaction_rule":"ACIAD2587"
},
{
"id":"TRANS_RXN_CAFFEATE_PERIP",
"name":"transport of caffeate [periplasm;cytosol](secondary symport)",
"metabolites":{
"cpd00067_C":1.0,
"cpd00067_P":-1.0,
"cpd00881_C":1.0,
"cpd00881_P":-1.0
},
"lower_bound":-1000.0,
"upper_bound":1000.0,
"gene_reaction_rule":"ACIAD1709"
},
{
"id":"GUANIDINOBUTYRASE_RXN",
"name":"guanidinobutyrase",
"metabolites":{
"cpd00001_C":-1.0,
"cpd00073_C":1.0,
"cpd00281_C":1.0,
"cpd00762_C":-1.0
},
"lower_bound":0.0,
"upper_bound":1000.0,
"gene_reaction_rule":"ACIAD1299"
},
{
"id":"TRANS_RXN_C10_0",
"name":"transport of decanoate [extraorganism;cytosol](secondary symport)",
"metabolites":{
"cpd00067_C":1.0,
"cpd00067_P":-1.0,
"cpd01107_C":1.0,
"cpd01107_E":-1.0
},
"lower_bound":-1000.0,
"upper_bound":1000.0,
"gene_reaction_rule":"ACIAD0835 or ACIAD1427 or ACIAD2788 or ACIAD3392"
},
{
"id":"PDXJ_RXN",
"name":"pyridoxine 5'-phosphate synthase",
"metabolites":{
"cpd00001_C":2.0,
"cpd00009_C":1.0,
"cpd00478_C":1.0,
"cpd08289_C":-1.0,
"cpd08449_C":-1.0
},
"lower_bound":-1000.0,
"upper_bound":1000.0,
"gene_reaction_rule":"ACIAD2577"
},
{
"id":"TYRAMINOTRANS_RXN",
"name":"tyrosine aminotransferase",
"metabolites":{
"cpd00023_C":-1.0,
"cpd00024_C":1.0,
"cpd00069_C":1.0,
"cpd00868_C":-1.0
},
"lower_bound":-1000.0,
"upper_bound":1000.0,
"gene_reaction_rule":"ACIAD0597 or ACIAD2574 or ACIAD0112"
},
{
"id":"SUCCINYLDIAMINOPIMTRANS_RXN",
"name":"succinyldiaminopimelate aminotransferase",
"metabolites":{
"cpd00023_C":1.0,
"cpd00024_C":-1.0,
"cpd02698_C":-1.0,
"cpd02724_C":1.0
},
"lower_bound":-1000.0,
"upper_bound":1000.0,
"gene_reaction_rule":"ACIAD1231 or ACIAD1284"
},
{
"id":"POLYSACCH_FREESYN_RXN",
"name":"generic free polysaccharides synthesis",
"metabolites":{
"POLYSACCH_FREE_W_C":0.8,
"UDP_POLYSACCH_LPS_C":-1.0,
"cpd00014_C":1.0
},
"lower_bound":0.0,
"upper_bound":1000.0,
"gene_reaction_rule":""
},
{
"id":"EXF_L_LACTATE_E",
"name":"exchange flux of lactate",
"metabolites":{
"cpd00159_E":-1.0
},
"lower_bound":-0.0,
"upper_bound":1000.0,
"gene_reaction_rule":""
},
{
"id":"F16BDEPHOS_RXN",
"name":"fructose-bisphosphatase",
"metabolites":{
"cpd00001_C":-1.0,
"cpd00009_C":1.0,
"cpd00290_C":-1.0,
"cpd15394_C":1.0
},
"lower_bound":0.0,
"upper_bound":1000.0,
"gene_reaction_rule":"ACIAD2625"
},
{
"id":"GLUTAMATE_N_ACETYLTRANSFERASE_RXN",
"name":"glutamate n-acetyltransferase",
"metabolites":{
"cpd00023_C":-1.0,
"cpd00064_C":1.0,
"cpd00342_C":-1.0,
"cpd00477_C":1.0
},
"lower_bound":0.0,
"upper_bound":1000.0,
"gene_reaction_rule":"ACIAD0650"
},
{
"id":"C16_0_ALDEHYDE_REDUCTASE_RXN",
"name":"long chain aldehyde reductase (c16:0)",
"metabolites":{
"C16_0_ALDEHYDE_C":-1.0,
"cpd00003_C":1.0,
"cpd00004_C":-1.0,
"cpd00613_C":1.0
},
"lower_bound":-1000.0,
"upper_bound":1000.0,
"gene_reaction_rule":""
},
{
"id":"TREHALOSE6PSYN_RXN",
"name":"alpha,alpha-trehalose-phosphate synthase (udp-forming)",
"metabolites":{
"cpd00014_C":1.0,
"cpd00026_C":-1.0,
"cpd00079_C":-1.0,
"cpd00523_C":1.0
},
"lower_bound":0.0,
"upper_bound":1000.0,
"gene_reaction_rule":"ACIAD1774"
},
{
"id":"DNA_SYNTH_RXN",
"name":"dna synthesis reaction",
"metabolites":{
"DNA_W_C":0.3087,
"cpd00002_C":-1.37,
"cpd00008_C":1.37,
"cpd00009_C":1.37,
"cpd00012_C":1.0,
"cpd00115_C":-0.298,
"cpd00241_C":-0.202,
"cpd00356_C":-0.202,
"cpd00357_C":-0.298
},
"lower_bound":0.0,
"upper_bound":1000.0,
"gene_reaction_rule":""
},
{
"id":"UDP_NACMUR_ALA_LIG_RXN",
"name":"udp-n-acetylmuramate--alanine ligase",
"metabolites":{
"cpd00002_C":-1.0,
"cpd00008_C":1.0,
"cpd00009_C":1.0,
"cpd00035_C":-1.0,
"cpd00773_C":-1.0,
"cpd00890_C":1.0
},
"lower_bound":0.0,
"upper_bound":1000.0,
"gene_reaction_rule":"ACIAD3516"
},
{
"id":"PMPOXI_RXN",
"name":"pyridoxamine-phosphate oxidase",
"metabolites":{
"cpd00001_C":-1.0,
"cpd00007_C":-1.0,
"cpd00013_C":1.0,
"cpd00016_C":1.0,
"cpd00025_C":1.0,
"cpd00493_C":-1.0
},
"lower_bound":0.0,
"upper_bound":1000.0,
"gene_reaction_rule":"ACIAD3501"
},
{
"id":"EXF_NITRITE_E",
"name":"exchange flux of no2-",
"metabolites":{
"cpd00075_E":-1.0
},
"lower_bound":-0.0,
"upper_bound":1000.0,
"gene_reaction_rule":""
},
{
"id":"BRANCHED_CHAINAMINOTRANSFERLEU_RXN",
"name":"leucine aminotransferase",
"metabolites":{
"cpd00023_C":-1.0,
"cpd00024_C":1.0,
"cpd00107_C":1.0,
"cpd00200_C":-1.0
},
"lower_bound":-1000.0,
"upper_bound":1000.0,
"gene_reaction_rule":"ACIAD0112 or ACIAD0597"
},
{
"id":"TRANS_RXNIM_5788",
"name":"transport of p-hydroxybenzoate [extraorganism;cytosol](secondary symport)",
"metabolites":{
"cpd00067_C":1.0,
"cpd00067_P":-1.0,
"cpd00136_C":1.0,
"cpd00136_E":-1.0
},
"lower_bound":-1000.0,
"upper_bound":1000.0,
"gene_reaction_rule":"ACIAD1709"
},
{
"id":"C14_1_C5_3HYDROXYCOA_DEHYDROGENASE_RXN",
"name":"hydroxyacyl-coa dehydrogenase (c14:1(c5))",
"metabolites":{
"cpd00003_C":-1.0,
"cpd00004_C":1.0,
"cpd16613_C":-1.0,
"cpd16615_C":1.0
},
"lower_bound":0.0,
"upper_bound":1000.0,
"gene_reaction_rule":"ACIAD0335 or ACIAD1565 or ACIAD2416"
},
{
"id":"BADH_RXN",
"name":"betaine-aldehyde dehydrogenase",
"metabolites":{
"cpd00001_C":-1.0,
"cpd00003_C":-1.0,
"cpd00004_C":1.0,
"cpd00447_C":-1.0,
"cpd00540_C":1.0
},
"lower_bound":-1000.0,
"upper_bound":1000.0,
"gene_reaction_rule":"ACIAD1009"
},
{
"id":"MALSYN_RXN",
"name":"malate synthase",
"metabolites":{
"cpd00001_C":-1.0,
"cpd00010_C":1.0,
"cpd00022_C":-1.0,
"cpd00040_C":-1.0,
"cpd00130_C":1.0
},
"lower_bound":0.0,
"upper_bound":1000.0,
"gene_reaction_rule":"ACIAD2335"
},
{
"id":"TRANS_RXNIM_5786",
"name":"transport of malonate [extraorganism;cytosol](secondary symport)",
"metabolites":{
"cpd00308_C":1.0,
"cpd00308_E":-1.0,
"cpd00971_C":1.0,
"cpd00971_E":-1.0
},
"lower_bound":-1000.0,
"upper_bound":1000.0,
"gene_reaction_rule":"ACIAD1760 and ACIAD1761"
},
{
"id":"TRANS_RXN_1",
"name":"transport of formate [cytosol;extraorganism](passive)",
"metabolites":{
"cpd00047_C":-1.0,
"cpd00047_E":1.0
},
"lower_bound":0.0,
"upper_bound":1000.0,
"gene_reaction_rule":"ACIAD1800 or ACIAD3593"
},
{
"id":"ALARACECAT_RXN",
"name":"alanine racemase",
"metabolites":{
"cpd00035_C":-1.0,
"cpd00117_C":1.0
},
"lower_bound":-1000.0,
"upper_bound":1000.0,
"gene_reaction_rule":"ACIAD2434 or ACIAD0116"
},
{
"id":"PTAALT_RXN",
"name":"phosphate acetyltransferase",
"metabolites":{
"cpd00009_C":1.0,
"cpd00010_C":-1.0,
"cpd00086_C":1.0,
"cpd01844_C":-1.0
},
"lower_bound":-1000.0,
"upper_bound":1000.0,
"gene_reaction_rule":"ACIAD0540"
},
{
"id":"TRANS_RXN_NMN",
"name":"transport of nicotinamide nucleotide [extraorganism;cytosol](passive)",
"metabolites":{
"cpd00355_C":1.0,
"cpd00355_E":-1.0
},
"lower_bound":0.0,
"upper_bound":1000.0,
"gene_reaction_rule":"ACIAD1033"
},
{
"id":"SUPEROX_DISMUT_RXN",
"name":"superoxide dismutase",
"metabolites":{
"cpd00007_C":1.0,
"cpd00025_C":1.0,
"cpd00067_C":-2.0,
"cpd00532_C":-2.0
},
"lower_bound":0.0,
"upper_bound":1000.0,
"gene_reaction_rule":"ACIAD1070 or ACIAD1346 or ACIAD2072"
},
{
"id":"ATPSYN_RXN",
"name":"h(+)-transporting atp synthase",
"metabolites":{
"cpd00001_C":1.0,
"cpd00002_C":1.0,
"cpd00008_C":-1.0,
"cpd00009_C":-1.0,
"cpd00067_C":3.0,
"cpd00067_P":-4.0
},
"lower_bound":-1000.0,
"upper_bound":1000.0,
"gene_reaction_rule":"ACIAD0178 and ACIAD0180 and ACIAD0182 and ACIAD0183 and ACIAD0184 and ACIAD0185 and ACIAD0186 and ACIAD0187 and ACIAD0188"
},
{
"id":"ACYL_COA_TAG_ASSEMBLY_RXN",
"name":"generic acyl-coa of triacylglycerol assembly",
"metabolites":{
"ACYL_COA_TAG_C":1.0,
"cpd00134_C":-0.717,
"cpd00327_C":-0.06,
"cpd01695_C":-0.022,
"cpd16627_C":-0.052,
"cpd16638_C":-0.149
},
"lower_bound":-1000.0,
"upper_bound":1000.0,
"gene_reaction_rule":""
},
{
"id":"C6_0_2ENOYLACP_REDUCTASE_RXN",
"name":"2enoyl-acp reductase (c6:0)",
"metabolites":{
"C6_0_2ENOYLACP_C":-1.0,
"C6_0_ACP_C":1.0,
"cpd00003_C":1.0,
"cpd00004_C":-1.0
},
"lower_bound":0.0,
"upper_bound":1000.0,
"gene_reaction_rule":"ACIAD3116"
},
{
"id":"TRANS_RXNIM_5771",
"name":"transport of no3- [extraorganism;cytosol](primary active atp)",
"metabolites":{
"cpd00001_C":-1.0,
"cpd00002_C":-1.0,
"cpd00008_C":1.0,
"cpd00009_C":1.0,
"cpd00209_C":1.0,
"cpd00209_E":-1.0
},
"lower_bound":0.0,
"upper_bound":1000.0,
"gene_reaction_rule":"ACIAD1914 and ACIAD1463 and ACIAD1465 and ACIAD1464"
},
{
"id":"PANTEPADENYLYLTRAN_RXN",
"name":"pantetheine-phosphate adenylyltransferase",
"metabolites":{
"cpd00002_C":-1.0,
"cpd00012_C":1.0,
"cpd00655_C":1.0,
"cpd00834_C":-1.0
},
"lower_bound":-1000.0,
"upper_bound":1000.0,
"gene_reaction_rule":"ACIAD2893"
},
{
"id":"TRANS_RXNIM_5773",
"name":"transport of no3- [extraorganism;cytosol](secondary symport)",
"metabolites":{
"cpd00067_C":1.0,
"cpd00067_P":-1.0,
"cpd00209_C":1.0,
"cpd00209_E":-1.0
},
"lower_bound":-1000.0,
"upper_bound":1000.0,
"gene_reaction_rule":"ACIAD1911"
},
{
"id":"C10_0_2ENOYLCOA_HYDRATASE_RXN",
"name":"enoyl-coa hydratase (c10:0)",
"metabolites":{
"cpd00001_C":-1.0,
"cpd03118_C":1.0,
"cpd03129_C":-1.0
},
"lower_bound":0.0,
"upper_bound":1000.0,
"gene_reaction_rule":"ACIAD1692 or ACIAD1726 or ACIAD1820 or ACIAD1829 or ACIAD2351 or ACIAD3073 or ACIAD1609 or ACIAD0335 or ACIAD1608 or ACIAD1701 or ACIAD1565 or ACIAD2416"
},
{
"id":"TRANS_RXN_QUINATE",
"name":"transport of quinate [extraorganism;periplasm](passive)",
"metabolites":{
"cpd00248_E":-1.0,
"cpd00248_P":1.0
},
"lower_bound":-1000.0,
"upper_bound":1000.0,
"gene_reaction_rule":"ACIAD1715 or ACIAD2984"
},
{
"id":"FOLYLPOLYGLUTAMATESYNTH_RXN",
"name":"folylpolyglutamate synthase",
"metabolites":{
"cpd00002_C":-2.0,
"cpd00008_C":2.0,
"cpd00009_C":2.0,
"cpd00023_C":-2.0,
"cpd00087_C":-1.0,
"cpd02555_C":1.0
},
"lower_bound":0.0,
"upper_bound":1000.0,
"gene_reaction_rule":"ACIAD0644"
},
{
"id":"AMP_DEPHOSPHORYLATION_RXN",
"name":"amp nucleotidase",
"metabolites":{
"cpd00001_C":-1.0,
"cpd00009_C":1.0,
"cpd00018_C":-1.0,
"cpd00182_C":1.0
},
"lower_bound":0.0,
"upper_bound":1000.0,
"gene_reaction_rule":"ACIAD0015 or ACIAD1227"
},
{
"id":"SUCCARGDIHYDRO_RXN",
"name":"succinylarginine dihydrolase",
"metabolites":{
"cpd00001_C":-2.0,
"cpd00011_C":1.0,
"cpd00013_C":2.0,
"cpd02103_C":-1.0,
"cpd02160_C":1.0
},
"lower_bound":0.0,
"upper_bound":1000.0,
"gene_reaction_rule":"ACIAD1288"
},
{
"id":"C5_0_CARBOXY3HYDROXYCOA_DEHYDROGENASE_RXN",
"name":"dicarboxylate hydroxyacyl-coa dehydrogenase (c5:0)",
"metabolites":{
"cpd00003_C":-1.0,
"cpd00004_C":1.0,
"cpd16643_C":-1.0,
"cpd16644_C":1.0
},
"lower_bound":0.0,
"upper_bound":1000.0,
"gene_reaction_rule":"ACIAD1690"
},
{
"id":"INORGPYROPHOSPHAT_RXN",
"name":"inorganic pyrophosphatase",
"metabolites":{
"cpd00001_C":-1.0,
"cpd00009_C":2.0,
"cpd00012_C":-1.0
},
"lower_bound":0.0,
"upper_bound":1000.0,
"gene_reaction_rule":"ACIAD0239"
},
{
"id":"EXF_FUM_E",
"name":"exchange flux of fumarate",
"metabolites":{
"cpd00106_E":-1.0
},
"lower_bound":-0.0,
"upper_bound":1000.0,
"gene_reaction_rule":""
},
{
"id":"BUTYRYL_COA_DEHYDROGENASE_RXN",
"name":"butyryl-coa dehydrogenase",
"metabolites":{
"cpd00015_C":-1.0,
"cpd00120_C":-1.0,
"cpd00650_C":1.0,
"cpd00982_C":1.0
},
"lower_bound":0.0,
"upper_bound":1000.0,
"gene_reaction_rule":"ACIAD1412"
},
{
"id":"GLUTAMINE_TRNA_LIGASE_RXN",
"name":"glutamine--trna ligase",
"metabolites":{
"CHARGED_GLN_TRNAS_C":1.0,
"GLN_TRNAS_C":-1.0,
"cpd00002_C":-1.0,
"cpd00012_C":1.0,
"cpd00018_C":1.0,
"cpd00053_C":-1.0
},
"lower_bound":0.0,
"upper_bound":1000.0,
"gene_reaction_rule":"ACIAD1920"
},
{
"id":"C14_0_2ENOYLACP_REDUCTASE_RXN",
"name":"2enoyl-acp reductase (c14:0)",
"metabolites":{
"C14_0_ACP_C":1.0,
"cpd00003_C":1.0,
"cpd00004_C":-1.0,
"cpd03127_C":-1.0
},
"lower_bound":0.0,
"upper_bound":1000.0,
"gene_reaction_rule":"ACIAD3116"
},
{
"id":"N_ACETYLGLUTPREDUCT_RXN",
"name":"n-acetyl-gamma-glutamyl-phosphate reductase",
"metabolites":{
"cpd00005_C":-1.0,
"cpd00006_C":1.0,
"cpd00009_C":1.0,
"cpd00918_C":1.0,
"cpd02552_C":-1.0
},
"lower_bound":-1000.0,
"upper_bound":1000.0,
"gene_reaction_rule":"ACIAD1360"
},
{
"id":"TRANS_RXN_INOSINE",
"name":"transport of inosine [extraorganism;cytosol](secondary symport)",
"metabolites":{
"cpd00067_C":1.0,
"cpd00067_P":-1.0,
"cpd00246_C":1.0,
"cpd00246_E":-1.0
},
"lower_bound":-1000.0,
"upper_bound":1000.0,
"gene_reaction_rule":"ACIAD2389"
},
{
"id":"C16_1_C7_2ENOYLACP_REDUCTASE_RXN",
"name":"2enoyl-acp reductase (c16:1(c7))",
"metabolites":{
"C16_1_C7__2ENOYLACP_C":-1.0,
"C16_1_C7__ACP_C":1.0,
"cpd00003_C":1.0,
"cpd00004_C":-1.0
},
"lower_bound":0.0,
"upper_bound":1000.0,
"gene_reaction_rule":"ACIAD3116"
},
{
"id":"EXF_C18_1_C9_E",
"name":"exchange flux of cis-octadec-9-enoate",
"metabolites":{
"cpd00536_E":-1.0
},
"lower_bound":-0.0,
"upper_bound":1000.0,
"gene_reaction_rule":""
},
{
"id":"PYRUVOXID_RXN",
"name":"pyruvate dehydrogenase (cytochrome)",
"metabolites":{
"cpd00001_C":-1.0,
"cpd00011_C":1.0,
"cpd00020_C":-1.0,
"cpd00029_C":1.0,
"cpd15560_C":-1.0,
"cpd15561_C":1.0
},
"lower_bound":0.0,
"upper_bound":1000.0,
"gene_reaction_rule":"ACIAD3381"
},
{
"id":"CHORISMATE_SYNTHASE_RXN",
"name":"chorismate synthase",
"metabolites":{
"cpd00009_C":1.0,
"cpd00216_C":1.0,
"cpd00932_C":-1.0
},
"lower_bound":0.0,
"upper_bound":1000.0,
"gene_reaction_rule":"ACIAD2028"
},
{
"id":"GLUTARATE_COA_LIGASE_RXN",
"name":"glutarate--coa ligase",
"metabolites":{
"cpd00002_C":-1.0,
"cpd00010_C":-1.0,
"cpd00012_C":1.0,
"cpd00018_C":1.0,
"cpd00379_C":-1.0,
"cpd00413_C":1.0
},
"lower_bound":0.0,
"upper_bound":1000.0,
"gene_reaction_rule":"ACIAD1695 and ACIAD1696"
},
{
"id":"TRANS_RXN_CPD_335",
"name":"transport of (r)-3-hydroxybutanoate [extraorganism;cytosol](passive)",
"metabolites":{
"cpd00797_C":1.0,
"cpd00797_E":-1.0
},
"lower_bound":-1000.0,
"upper_bound":1000.0,
"gene_reaction_rule":"ACIAD2510"
},
{
"id":"GLUCOSAMINE1PACETYL_RXN",
"name":"glucosamine-1-phosphate N-acetyltransferase",
"metabolites":{
"cpd00010_C":1.0,
"cpd00022_C":-1.0,
"cpd02370_C":-1.0,
"cpd02611_C":1.0
},
"lower_bound":0.0,
"upper_bound":1000.0,
"gene_reaction_rule":"ACIAD3575"
},
{
"id":"PROPIONATE_COA_LIGASE_RXN",
"name":"propionate--coa ligase",
"metabolites":{
"cpd00002_C":-1.0,
"cpd00010_C":-1.0,
"cpd00012_C":1.0,
"cpd00018_C":1.0,
"cpd00086_C":1.0,
"cpd00141_C":-1.0
},
"lower_bound":0.0,
"upper_bound":1000.0,
"gene_reaction_rule":"ACIAD3475"
},
{
"id":"ACYL_ACP_TAG_ASSEMBLY_RXN",
"name":"generic acyl-acp of triacylglycerol assembly",
"metabolites":{
"ACYL_ACP_TAG_C":1.0,
"C14_0_ACP_C":-0.022,
"C16_0_ACP_C":-0.717,
"C16_1_C7__ACP_C":-0.052,
"C18_0_ACP_C":-0.06,
"C18_1_C9__ACP_C":-0.149
},
"lower_bound":-1000.0,
"upper_bound":1000.0,
"gene_reaction_rule":""
},
{
"id":"ISOCHORSYN_RXN",
"name":"isochorismate mutase",
"metabolites":{
"cpd00216_C":-1.0,
"cpd00658_C":1.0
},
"lower_bound":0.0,
"upper_bound":1000.0,
"gene_reaction_rule":"ACIAD2776"
},
{
"id":"C6_1_C4_2ENOYLCOA_HYDRATASE_RXN",
"name":"enoyl-coa hydratase (c6:1(c4))",
"metabolites":{
"C6_1_C4__2ENOYLCOA_C":-1.0,
"cpd00001_C":-1.0,
"cpd16646_C":1.0
},
"lower_bound":0.0,
"upper_bound":1000.0,
"gene_reaction_rule":"ACIAD1692 or ACIAD1726 or ACIAD1820 or ACIAD1829 or ACIAD2351 or ACIAD3073 or ACIAD1609 or ACIAD0335 or ACIAD1608 or ACIAD1701 or ACIAD1565 or ACIAD2416"
},
{
"id":"DAPASYN_RXN",
"name":"adenosylmethionine--8-amino-7-oxononanoate aminotransferase",
"metabolites":{
"cpd00017_C":-1.0,
"cpd00764_C":1.0,
"cpd00800_C":-1.0,
"cpd02701_C":1.0
},
"lower_bound":-1000.0,
"upper_bound":1000.0,
"gene_reaction_rule":"ACIAD0856"
},
{
"id":"EXF_THR_E",
"name":"exchange flux of l-threonine",
"metabolites":{
"cpd00161_E":-1.0
},
"lower_bound":-0.0,
"upper_bound":1000.0,
"gene_reaction_rule":""
},
{
"id":"PROLDEHYDROG_RXN",
"name":"proline dehydrogenase",
"metabolites":{
"cpd00015_C":-1.0,
"cpd00129_C":-1.0,
"cpd00982_C":1.0,
"cpd02431_C":1.0
},
"lower_bound":0.0,
"upper_bound":1000.0,
"gene_reaction_rule":"ACIAD1646"
},
{
"id":"GLYRIBONUCSYN_RXN",
"name":"phosphoribosylamine--glycine ligase",
"metabolites":{
"cpd00002_C":-1.0,
"cpd00008_C":1.0,
"cpd00009_C":1.0,
"cpd00033_C":-1.0,
"cpd01982_C":-1.0,
"cpd02394_C":1.0
},
"lower_bound":0.0,
"upper_bound":1000.0,
"gene_reaction_rule":"ACIAD2448"
},
{
"id":"GLUCONOKIN_RXN",
"name":"gluconokinase",
"metabolites":{
"cpd00002_C":-1.0,
"cpd00008_C":1.0,
"cpd00222_C":-1.0,
"cpd00284_C":1.0
},
"lower_bound":-1000.0,
"upper_bound":1000.0,
"gene_reaction_rule":"ACIAD0545"
},
{
"id":"TRANS_RXN_104",
"name":"transport of lactate [extraorganism;cytosol](secondary symport)",
"metabolites":{
"cpd00067_C":1.0,
"cpd00067_P":-1.0,
"cpd00159_C":1.0,
"cpd00159_E":-1.0
},
"lower_bound":0.0,
"upper_bound":1000.0,
"gene_reaction_rule":"ACIAD0106"
},
{
"id":"GLYCEROL_3_P_ACYLTRANSFER_TAG_RXN",
"name":"glycerol-3-phosphate acyltransferase",
"metabolites":{
"ACP_C":1.0,
"ACYL_ACP_TAG_C":-1.0,
"ACYL_SN_GLYCEROL_3P_TAG_C":1.0,
"cpd00080_C":-1.0
},
"lower_bound":0.0,
"upper_bound":1000.0,
"gene_reaction_rule":"ACIAD3232"
},
{
"id":"TRANS_RXN_132",
"name":"transport of uracil [extraorganism;cytosol](secondary symport)",
"metabolites":{
"cpd00067_C":1.0,
"cpd00067_P":-1.0,
"cpd00092_C":1.0,
"cpd00092_E":-1.0
},
"lower_bound":0.0,
"upper_bound":1000.0,
"gene_reaction_rule":"ACIAD3628 or ACIAD0031 or ACIAD0136 or ACIAD0702 or ACIAD0762 or ACIAD1243"
},
{
"id":"TRANS_RXN_ETOH",
"name":"transport of ethanol [extraorganism;cytosol](passive)",
"metabolites":{
"cpd00363_C":1.0,
"cpd00363_E":-1.0
},
"lower_bound":-1000.0,
"upper_bound":1000.0,
"gene_reaction_rule":""
},
{
"id":"C16_0_2ENOYLCOA_HYDRATASE_RXN",
"name":"enoyl-coa hydratase (c16:0)",
"metabolites":{
"cpd00001_C":-1.0,
"cpd03113_C":1.0,
"cpd03126_C":-1.0
},
"lower_bound":0.0,
"upper_bound":1000.0,
"gene_reaction_rule":"ACIAD1692 or ACIAD1726 or ACIAD1820 or ACIAD1829 or ACIAD2351 or ACIAD3073 or ACIAD1609 or ACIAD0335 or ACIAD1608 or ACIAD1701 or ACIAD1565 or ACIAD2416"
},
{
"id":"C5_0_3HYDROXYCOA_DEHYDROGENASE_RXN",
"name":"hydroxyacyl-coa dehydrogenase (c5:0)",
"metabolites":{
"cpd00003_C":-1.0,
"cpd00004_C":1.0,
"cpd23920_C":-1.0,
"cpd23921_C":1.0
},
"lower_bound":0.0,
"upper_bound":1000.0,
"gene_reaction_rule":"ACIAD0335 or ACIAD1565 or ACIAD2416"
},
{
"id":"TRANS_RXN_1_BIS",
"name":"transport of no2- [cytosol;extraorganism](passive)",
"metabolites":{
"cpd00075_C":-1.0,
"cpd00075_E":1.0
},
"lower_bound":-1000.0,
"upper_bound":1000.0,
"gene_reaction_rule":"ACIAD1800 or ACIAD3593"
},
{
"id":"URIDINE_NUCLEOSIDASE_RXN",
"name":"uridine nucleosidase",
"metabolites":{
"cpd00001_C":-1.0,
"cpd00092_C":1.0,
"cpd00105_C":1.0,
"cpd00249_C":-1.0
},
"lower_bound":0.0,
"upper_bound":1000.0,
"gene_reaction_rule":"ACIAD2386"
},
{
"id":"UNDECAPRENOL_KINASE_RXN",
"name":"undecaprenol kinase",
"metabolites":{
"cpd00002_C":-1.0,
"cpd00008_C":1.0,
"cpd00286_C":1.0,
"cpd01352_C":-1.0
},
"lower_bound":0.0,
"upper_bound":1000.0,
"gene_reaction_rule":"ACIAD0678 or ACIAD0914 or ACIAD1642"
},
{
"id":"DADPKIN_RXN",
"name":"dadp kinase",
"metabolites":{
"cpd00002_C":-1.0,
"cpd00008_C":1.0,
"cpd00115_C":1.0,
"cpd00177_C":-1.0
},
"lower_bound":-1000.0,
"upper_bound":1000.0,
"gene_reaction_rule":"ACIAD0556"
},
{
"id":"RXN0_884",
"name":"1-hydroxy-2-methyl-2-(e)-butenyl 4-diphosphate reductase (dmpp)",
"metabolites":{
"cpd00001_C":1.0,
"cpd00003_C":1.0,
"cpd00004_C":-1.0,
"cpd00067_C":-1.0,
"cpd00202_C":1.0,
"cpd08615_C":-1.0
},
"lower_bound":-1000.0,
"upper_bound":1000.0,
"gene_reaction_rule":"ACIAD3322"
},
{
"id":"KDOTRANS2_RXN",
"name":"3-deoxy-d-manno-octulosonic acid transferase",
"metabolites":{
"cpd00046_C":1.0,
"cpd02546_C":-1.0,
"cpd03585_C":-1.0,
"cpd03586_C":1.0
},
"lower_bound":0.0,
"upper_bound":1000.0,
"gene_reaction_rule":"ACIAD3479"
},
{
"id":"EXF_4_HYDROXYBENZOATE_E",
"name":"exchange flux of p-hydroxybenzoate",
"metabolites":{
"cpd00136_E":-1.0
},
"lower_bound":-0.0,
"upper_bound":1000.0,
"gene_reaction_rule":""
},
{
"id":"TRANS_RXN_CPD_6501",
"name":"transport of benzylacetate [extraorganism;cytosol](passive)",
"metabolites":{
"cpd11193_C":1.0,
"cpd11193_E":-1.0
},
"lower_bound":-1000.0,
"upper_bound":1000.0,
"gene_reaction_rule":""
},
{
"id":"C16_1_C7_3HYDROXYACP_DEHYDRATASE_RXN",
"name":"3hydroxyacyl-acp dehydratase (c16:1(c7))",
"metabolites":{
"C16_1_C7__2ENOYLACP_C":1.0,
"C16_1_C7__3HYDROXYACP_C":-1.0,
"cpd00001_C":1.0
},
"lower_bound":0.0,
"upper_bound":1000.0,
"gene_reaction_rule":"ACIAD1381"
},
{
"id":"EXF_TAURINE_E",
"name":"exchange flux of taurine",
"metabolites":{
"cpd00210_E":-1.0
},
"lower_bound":-0.0,
"upper_bound":1000.0,
"gene_reaction_rule":""
},
{
"id":"LPSSYN_RXN",
"name":"generic lps assembly",
"metabolites":{
"LPS_W_C":3.005,
"UDP_POLYSACCH_LPS_C":-1.0,
"cpd00014_C":1.0,
"cpd03587_C":-1.0
},
"lower_bound":0.0,
"upper_bound":1000.0,
"gene_reaction_rule":""
},
{
"id":"DARAB5PISOM_RXN",
"name":"arabinose-5-phosphate isomerase",
"metabolites":{
"cpd00171_C":1.0,
"cpd00817_C":-1.0
},
"lower_bound":-1000.0,
"upper_bound":1000.0,
"gene_reaction_rule":"ACIAD1482"
},
{
"id":"RXN_2425",
"name":"trans-2,3-dehydroadipyl-coa hydratase",
"metabolites":{
"cpd00001_C":-1.0,
"cpd09844_C":1.0,
"cpd28230_C":-1.0
},
"lower_bound":0.0,
"upper_bound":1000.0,
"gene_reaction_rule":"ACIAD1692"
},
{
"id":"2_5_DIOXOVALERATE_DEHYDROGENASE_RXN",
"name":"2,5-dioxovalerate dehydrogenase",
"metabolites":{
"cpd00001_C":-1.0,
"cpd00005_C":1.0,
"cpd00006_C":-1.0,
"cpd00024_C":1.0,
"cpd00340_C":-1.0
},
"lower_bound":-1000.0,
"upper_bound":1000.0,
"gene_reaction_rule":"ACIAD0131"
},
{
"id":"C14_1_C5_3OXOACP_REDUCTASE_RXN",
"name":"3oxoacyl-acp reductase (c14:1(c5))",
"metabolites":{
"C14_1_C5__3HYDROXYACP_C":1.0,
"C14_1_C5__3OXOACP_C":-1.0,
"cpd00005_C":-1.0,
"cpd00006_C":1.0
},
"lower_bound":0.0,
"upper_bound":1000.0,
"gene_reaction_rule":"ACIAD0871"
},
{
"id":"EXF_BUTANEDIOL_E",
"name":"exchange flux of 2,3-butanediol",
"metabolites":{
"cpd01947_E":-1.0
},
"lower_bound":-0.0,
"upper_bound":1000.0,
"gene_reaction_rule":""
},
{
"id":"TRANS_RXN_MAL",
"name":"transport of malate [extraorganism;cytosol](secondary symport) TDT family",
"metabolites":{
"cpd00067_C":1.0,
"cpd00067_P":-1.0,
"cpd00130_C":1.0,
"cpd00130_E":-1.0
},
"lower_bound":0.0,
"upper_bound":1000.0,
"gene_reaction_rule":"ACIAD0024"
},
{
"id":"CITSYN_RXN",
"name":"citrate (si)-synthase",
"metabolites":{
"cpd00001_C":-1.0,
"cpd00010_C":1.0,
"cpd00022_C":-1.0,
"cpd00032_C":-1.0,
"cpd00137_C":1.0
},
"lower_bound":0.0,
"upper_bound":1000.0,
"gene_reaction_rule":"ACIAD2886"
},
{
"id":"PNPOXI_RXN",
"name":"pyridoxine 5'-phosphate oxidase",
"metabolites":{
"cpd00007_C":-1.0,
"cpd00016_C":1.0,
"cpd00025_C":1.0,
"cpd00478_C":-1.0
},
"lower_bound":-1000.0,
"upper_bound":1000.0,
"gene_reaction_rule":"ACIAD3501"
},
{
"id":"CATECHOL_1_2_DIOXYGENASE_RXN",
"name":"catechol 1,2-dioxygenase",
"metabolites":{
"cpd00007_C":-1.0,
"cpd00077_C":-1.0,
"cpd01630_C":1.0
},
"lower_bound":0.0,
"upper_bound":1000.0,
"gene_reaction_rule":"ACIAD1442"
},
{
"id":"TRANS_RXN_CIT_MFS",
"name":"transport of citrate [extraorganism;cytosol](secondary symport)",
"metabolites":{
"cpd00067_C":1.0,
"cpd00067_P":-1.0,
"cpd00137_C":1.0,
"cpd00137_E":-1.0
},
"lower_bound":-1000.0,
"upper_bound":1000.0,
"gene_reaction_rule":"ACIAD0395 or ACIAD1541"
},
{
"id":"TRANS_RXN_CHLOROGENATE",
"name":"transport of chlorogenate [extraorganism;periplasm](passive)",
"metabolites":{
"cpd00634_E":-1.0,
"cpd00634_P":1.0
},
"lower_bound":0.0,
"upper_bound":1000.0,
"gene_reaction_rule":""
},
{
"id":"UDP_NACMURALGLDAPAALIG_RXN",
"name":"udp-n-acetylmuramoylalanyl-d-glutamyl-2,6-diaminopimelate--d-alanyl- d-alanine ligase",
"metabolites":{
"cpd00002_C":-1.0,
"cpd00008_C":1.0,
"cpd00009_C":1.0,
"cpd00731_C":-1.0,
"cpd02932_C":1.0,
"cpd02964_C":-1.0
},
"lower_bound":0.0,
"upper_bound":1000.0,
"gene_reaction_rule":"ACIAD3364"
},
{
"id":"UROGENIIISYN_RXN",
"name":"uroporphyrinogen-iii synthase",
"metabolites":{
"cpd00001_C":1.0,
"cpd00755_C":-1.0,
"cpd00774_C":1.0
},
"lower_bound":-1000.0,
"upper_bound":1000.0,
"gene_reaction_rule":"ACIAD0287"
},
{
"id":"4OHBENZOATE_OCTAPRENYLTRANSFER_RXN",
"name":"hydroxybenzoate octaprenyltransferase",
"metabolites":{
"cpd00012_C":1.0,
"cpd00136_C":-1.0,
"cpd02557_C":-1.0,
"cpd03443_C":1.0
},
"lower_bound":-1000.0,
"upper_bound":1000.0,
"gene_reaction_rule":"ACIAD2455"
},
{
"id":"PSERPHOSPHA_RXN",
"name":"phosphoserine phosphatase",
"metabolites":{
"cpd00001_C":-1.0,
"cpd00009_C":1.0,
"cpd00054_C":1.0,
"cpd00738_C":-1.0
},
"lower_bound":0.0,
"upper_bound":1000.0,
"gene_reaction_rule":"ACIAD3567"
},
{
"id":"TRANS_RXN_PHE",
"name":"transport of l-phenylalanine [extraorganism;cytosol](secondary symport)",
"metabolites":{
"cpd00066_C":1.0,
"cpd00066_E":-1.0,
"cpd00067_C":1.0,
"cpd00067_P":-1.0
},
"lower_bound":-1000.0,
"upper_bound":1000.0,
"gene_reaction_rule":"ACIAD1343 or ACIAD2969"
},
{
"id":"AICARTRANSFORM_RXN",
"name":"phosphoribosylaminoimidazolecarboxamide formyltransferase",
"metabolites":{
"cpd00087_C":1.0,
"cpd00201_C":-1.0,
"cpd02851_C":-1.0,
"cpd02884_C":1.0
},
"lower_bound":0.0,
"upper_bound":1000.0,
"gene_reaction_rule":"ACIAD2447"
},
{
"id":"GLUC1PURIDYLTRANS_RXN",
"name":"utp--glucose-1-phosphate uridylyltransferase",
"metabolites":{
"cpd00012_C":1.0,
"cpd00026_C":1.0,
"cpd00062_C":-1.0,
"cpd00089_C":-1.0
},
"lower_bound":0.0,
"upper_bound":1000.0,
"gene_reaction_rule":"ACIAD0099"
},
{
"id":"SAICARSYN_RXN",
"name":"phosphoribosylaminoimidazole-succinocarboxamide synthase",
"metabolites":{
"cpd00002_C":-1.0,
"cpd00008_C":1.0,
"cpd00009_C":1.0,
"cpd00041_C":-1.0,
"cpd02893_C":-1.0,
"cpd02921_C":1.0
},
"lower_bound":0.0,
"upper_bound":1000.0,
"gene_reaction_rule":"ACIAD3587"
},
{
"id":"RXN0_311",
"name":"methyl-cis-aconitic acid hydratase",
"metabolites":{
"cpd00001_C":-1.0,
"cpd02597_C":-1.0,
"cpd02799_C":1.0
},
"lower_bound":-1000.0,
"upper_bound":1000.0,
"gene_reaction_rule":"ACIAD2755"
},
{
"id":"EXF_4_HYDROXY_BENZYL_ALCOHOL_E",
"name":"exchange flux of 4-hydroxybenzyl alcohol",
"metabolites":{
"cpd15378_E":-1.0
},
"lower_bound":-0.0,
"upper_bound":1000.0,
"gene_reaction_rule":""
},
{
"id":"GPPSYN_RXN",
"name":"dimethylallyltransferase",
"metabolites":{
"cpd00012_C":1.0,
"cpd00113_C":-1.0,
"cpd00202_C":-1.0,
"cpd00283_C":1.0
},
"lower_bound":-1000.0,
"upper_bound":1000.0,
"gene_reaction_rule":"ACIAD2968 or ACIAD2940 or ACIAD1374"
},
{
"id":"EXF_GLC_E",
"name":"exchange flux of beta-d-glucose",
"metabolites":{
"cpd00027_E":-1.0
},
"lower_bound":-0.0,
"upper_bound":1000.0,
"gene_reaction_rule":""
},
{
"id":"CHLOROGENATE_HYDROLASE_RXN",
"name":"chlorogenate hydrolase",
"metabolites":{
"cpd00001_P":-1.0,
"cpd00248_P":1.0,
"cpd00634_P":-1.0,
"cpd00881_P":1.0
},
"lower_bound":-1000.0,
"upper_bound":1000.0,
"gene_reaction_rule":"ACIAD1720"
},
{
"id":"EXF_QUINATE_E",
"name":"exchange flux of quinate",
"metabolites":{
"cpd00248_E":-1.0
},
"lower_bound":-0.0,
"upper_bound":1000.0,
"gene_reaction_rule":""
},
{
"id":"PEPTIDOSYN_RXN",
"name":"peptidoglycan synthesis",
"metabolites":{
"C6_C":-1.0,
"PEPTIDO_W_E":0.9208,
"cpd00117_E":0.8,
"cpd02229_C":1.0
},
"lower_bound":0.0,
"upper_bound":1000.0,
"gene_reaction_rule":""
},
{
"id":"C4_0_3OXOACP_SYNTHASE_RXN",
"name":"3oxoacyl-acp synthase (c4:0)",
"metabolites":{
"ACETOACETYL_S_ACP_C":1.0,
"ACETYL_ACP_C":-1.0,
"ACP_C":1.0,
"MALONYL_ACP_C":-1.0,
"cpd00011_C":1.0
},
"lower_bound":0.0,
"upper_bound":1000.0,
"gene_reaction_rule":"ACIAD0580 or ACIAD0879 or ACIAD0583 or ACIAD2101"
},
{
"id":"TRANS_RXN_TYR",
"name":"transport of l-tyrosine [extraorganism;cytosol](secondary symport)",
"metabolites":{
"cpd00067_C":1.0,
"cpd00067_P":-1.0,
"cpd00069_C":1.0,
"cpd00069_E":-1.0
},
"lower_bound":-1000.0,
"upper_bound":1000.0,
"gene_reaction_rule":"ACIAD1343 or ACIAD2969"
},
{
"id":"CATAL_RXN",
"name":"catalase",
"metabolites":{
"cpd00001_C":2.0,
"cpd00007_C":1.0,
"cpd00025_C":-2.0
},
"lower_bound":0.0,
"upper_bound":1000.0,
"gene_reaction_rule":"ACIAD0451"
},
{
"id":"QUINOPRIBOTRANS_RXN",
"name":"nicotinate-nucleotide pyrophosphorylase (carboxylating)",
"metabolites":{
"NICOTINATE_NUCLEOTIDE_C":1.0,
"cpd00011_C":1.0,
"cpd00012_C":1.0,
"cpd00103_C":-1.0,
"cpd02333_C":-1.0
},
"lower_bound":0.0,
"upper_bound":1000.0,
"gene_reaction_rule":"ACIAD0062"
},
{
"id":"5_APOST_NUCLEOTID_RXN",
"name":"ump nucleotidase",
"metabolites":{
"cpd00001_C":-1.0,
"cpd00009_C":1.0,
"cpd00091_C":-1.0,
"cpd00249_C":1.0
},
"lower_bound":0.0,
"upper_bound":1000.0,
"gene_reaction_rule":"ACIAD0015 or ACIAD1227"
},
{
"id":"3_DEHYDROQUINATE_DEHYDRATASE_RXN",
"name":"3-dehydroquinate dehydratase",
"metabolites":{
"cpd00001_C":1.0,
"cpd00699_C":-1.0,
"cpd01716_C":1.0
},
"lower_bound":-1000.0,
"upper_bound":1000.0,
"gene_reaction_rule":"ACIAD1738"
},
{
"id":"TRANS_RXN_L_LACTATE",
"name":"transport of lactate [extraorganism;cytosol](passive)",
"metabolites":{
"cpd00159_C":1.0,
"cpd00159_E":-1.0
},
"lower_bound":-1000.0,
"upper_bound":1000.0,
"gene_reaction_rule":""
},
{
"id":"MALIC_NADP_RXN",
"name":"malate dehydrogenase (oxaloacetate decarboxylating) (nadp+)",
"metabolites":{
"cpd00005_C":1.0,
"cpd00006_C":-1.0,
"cpd00011_C":1.0,
"cpd00020_C":1.0,
"cpd00130_C":-1.0
},
"lower_bound":0.0,
"upper_bound":1000.0,
"gene_reaction_rule":"ACIAD2287"
},
{
"id":"EXF_D_LACTATE_E",
"name":"exchange flux of d-lactate",
"metabolites":{
"cpd00221_E":-1.0
},
"lower_bound":-0.0,
"upper_bound":1000.0,
"gene_reaction_rule":""
},
{
"id":"TRANS_RXN_C16_1_C7_",
"name":"transport of cis-hexadec-7-enoate [extraorganism;cytosol](secondary symport)",
"metabolites":{
"cpd00067_C":1.0,
"cpd00067_P":-1.0,
"cpd34461_C":1.0,
"cpd34461_E":-1.0
},
"lower_bound":-1000.0,
"upper_bound":1000.0,
"gene_reaction_rule":"ACIAD0835 or ACIAD1427 or ACIAD2788 or ACIAD3392"
},
{
"id":"PGLUCISOM_RXN",
"name":"glucose-6-phosphate isomerase",
"metabolites":{
"cpd00079_C":-1.0,
"cpd15394_C":1.0
},
"lower_bound":-1000.0,
"upper_bound":1000.0,
"gene_reaction_rule":"ACIAD0101"
},
{
"id":"RXN0_882",
"name":"1-hydroxy-2-methyl-2-(e)-butenyl 4-diphosphate synthase",
"metabolites":{
"cpd00001_C":1.0,
"cpd00067_C":-1.0,
"cpd08301_C":-1.0,
"cpd08615_C":1.0
},
"lower_bound":-1000.0,
"upper_bound":1000.0,
"gene_reaction_rule":"ACIAD0561"
},
{
"id":"DIHYDROPICRED_RXN",
"name":"dihydrodipicolinate reductase",
"metabolites":{
"cpd00005_C":-1.0,
"cpd00006_C":1.0,
"cpd02120_C":-1.0,
"cpd02465_C":1.0
},
"lower_bound":0.0,
"upper_bound":1000.0,
"gene_reaction_rule":"ACIAD3619"
},
{
"id":"SULFATE_ADENYLYLTRANS_RXN",
"name":"sulfate adenylyltransferase",
"metabolites":{
"APS_C":1.0,
"cpd00002_C":-1.0,
"cpd00012_C":1.0,
"cpd00048_C":-1.0
},
"lower_bound":-1000.0,
"upper_bound":1000.0,
"gene_reaction_rule":"ACIAD1073 and ACIAD1072"
},
{
"id":"PRIBFAICARPISOM_RXN",
"name":"n-(5'-phospho-d-ribosylformimino)-5-amino-1-(5''-phosphoribosyl)-4- imidazole carboxamide isomerase",
"metabolites":{
"cpd02979_C":-1.0,
"cpd02991_C":1.0
},
"lower_bound":0.0,
"upper_bound":1000.0,
"gene_reaction_rule":"ACIAD3398"
},
{
"id":"FPPSYN_RXN",
"name":"geranyltranstransferase",
"metabolites":{
"cpd00012_C":1.0,
"cpd00113_C":-1.0,
"cpd00283_C":-1.0,
"cpd00350_C":1.0
},
"lower_bound":-1000.0,
"upper_bound":1000.0,
"gene_reaction_rule":"ACIAD2968 or ACIAD2940 or ACIAD1374"
},
{
"id":"TRANS_RXN_NA_CPA1",
"name":"transport of na+ [extraorganism;cytosol](secondary antiport)",
"metabolites":{
"cpd00067_C":-1.0,
"cpd00067_P":1.0,
"cpd00971_C":1.0,
"cpd00971_E":-1.0
},
"lower_bound":-1000.0,
"upper_bound":1000.0,
"gene_reaction_rule":"ACIAD2867"
},
{
"id":"TRANS_RXN_C5_0",
"name":"transport of pentanoate [extraorganism;cytosol](secondary symport)",
"metabolites":{
"cpd00067_C":1.0,
"cpd00067_P":-1.0,
"cpd00597_C":1.0,
"cpd00597_E":-1.0
},
"lower_bound":-1000.0,
"upper_bound":1000.0,
"gene_reaction_rule":"ACIAD0835 or ACIAD1427 or ACIAD2788 or ACIAD3392"
},
{
"id":"C18_0_COATRANSFERASE_RXN",
"name":"acyl coa transferase (c18:0)",
"metabolites":{
"cpd00002_C":-1.0,
"cpd00010_C":-1.0,
"cpd00012_C":1.0,
"cpd00018_C":1.0,
"cpd00327_C":1.0,
"cpd01080_C":-1.0
},
"lower_bound":0.0,
"upper_bound":1000.0,
"gene_reaction_rule":"ACIAD1818 or ACIAD0235 or ACIAD1724 or ACIAD2106"
},
{
"id":"EXF_TOLUENE_E",
"name":"exchange flux of toluene",
"metabolites":{
"cpd01034_E":-1.0
},
"lower_bound":-0.0,
"upper_bound":1000.0,
"gene_reaction_rule":""
},
{
"id":"HISTOLDEHYD_RXN",
"name":"histidinol dehydrogenase",
"metabolites":{
"cpd00003_C":-1.0,
"cpd00004_C":1.0,
"cpd00641_C":-1.0,
"cpd01324_C":1.0
},
"lower_bound":0.0,
"upper_bound":1000.0,
"gene_reaction_rule":"ACIAD0663"
},
{
"id":"C8_0_2ENOYLCOA_HYDRATASE_RXN",
"name":"enoyl-coa hydratase (c8:0)",
"metabolites":{
"cpd00001_C":-1.0,
"cpd03120_C":1.0,
"cpd03130_C":-1.0
},
"lower_bound":0.0,
"upper_bound":1000.0,
"gene_reaction_rule":"ACIAD1692 or ACIAD1726 or ACIAD1820 or ACIAD1829 or ACIAD2351 or ACIAD3073 or ACIAD1609 or ACIAD0335 or ACIAD1608 or ACIAD1701 or ACIAD1565 or ACIAD2416"
},
{
"id":"RIBOFLAVINSYNREDUC_RXN",
"name":"5-amino-6-(5- phosphoribosylamino)uracil reductase (htp reductase)",
"metabolites":{
"cpd00005_C":-1.0,
"cpd00006_C":1.0,
"cpd00931_C":-1.0,
"cpd02720_C":1.0
},
"lower_bound":-1000.0,
"upper_bound":1000.0,
"gene_reaction_rule":"ACIAD0247"
},
{
"id":"GLY3KIN_RXN",
"name":"glycerate kinase",
"metabolites":{
"cpd00002_C":-1.0,
"cpd00008_C":1.0,
"cpd00169_C":1.0,
"cpd00223_C":-1.0
},
"lower_bound":0.0,
"upper_bound":1000.0,
"gene_reaction_rule":"ACIAD2850"
},
{
"id":"3_ISOPROPYLMALISOM_RXN",
"name":"3-isopropylmalate dehydratase",
"metabolites":{
"cpd01646_C":-1.0,
"cpd02693_C":1.0
},
"lower_bound":-1000.0,
"upper_bound":1000.0,
"gene_reaction_rule":"ACIAD0466 and ACIAD0463"
},
{
"id":"TRANS_RXN_FE2",
"name":"transport of fe+2 [extraorganism;cytosol](passive)",
"metabolites":{
"cpd00021_C":1.0,
"cpd00021_E":-1.0
},
"lower_bound":-1000.0,
"upper_bound":1000.0,
"gene_reaction_rule":"ACIAD0268 or ACIAD1471 or ACIAD0531"
},
{
"id":"C6_0_3OXOACP_REDUCTASE_RXN",
"name":"3oxoacyl-acp reductase (c6:0)",
"metabolites":{
"C6_0_3HYDROXYACP_C":1.0,
"C6_0_3OXOACP_C":-1.0,
"cpd00005_C":-1.0,
"cpd00006_C":1.0
},
"lower_bound":0.0,
"upper_bound":1000.0,
"gene_reaction_rule":"ACIAD0871"
},
{
"id":"C18_1_C9_2ENOYLACP_REDUCTASE_RXN",
"name":"2enoyl-acp reductase (c18:1(c9))",
"metabolites":{
"C18_1_C9__2ENOYLACP_C":-1.0,
"C18_1_C9__ACP_C":1.0,
"cpd00003_C":1.0,
"cpd00004_C":-1.0
},
"lower_bound":0.0,
"upper_bound":1000.0,
"gene_reaction_rule":"ACIAD3116"
},
{
"id":"C14_1_C5_3OXOCOA_THIOLASE_RXN",
"name":"3oxoacyl-coa thiolase (c14:1(c5))",
"metabolites":{
"cpd00010_C":-1.0,
"cpd00022_C":1.0,
"cpd01887_C":1.0,
"cpd16615_C":-1.0
},
"lower_bound":-1000.0,
"upper_bound":1000.0,
"gene_reaction_rule":"ACIAD0334 or ACIAD1450 or ACIAD1706 or ACIAD1977 or ACIAD1689 or ACIAD2988 or ACIAD1569"
},
{
"id":"ASPCARBTRANS_RXN",
"name":"aspartate carbamoyltransferase",
"metabolites":{
"cpd00009_C":1.0,
"cpd00041_C":-1.0,
"cpd00146_C":-1.0,
"cpd00343_C":1.0
},
"lower_bound":0.0,
"upper_bound":1000.0,
"gene_reaction_rule":"ACIAD1270 and ACIAD1271"
},
{
"id":"C18_1_C9_COA_THIOLASE_RXN",
"name":"acyl-coa thiolase (c18:1(c9))",
"metabolites":{
"cpd00001_C":-1.0,
"cpd00010_C":1.0,
"cpd00536_C":1.0,
"cpd16638_C":-1.0
},
"lower_bound":0.0,
"upper_bound":1000.0,
"gene_reaction_rule":"ACIAD0341 or ACIAD3231 or ACIAD1023"
},
{
"id":"ACETYLHOMOSER_CYS_RXN",
"name":"o-acetylhomoserine sulfhydrolase",
"metabolites":{
"cpd00029_C":1.0,
"cpd00135_C":1.0,
"cpd00239_C":-1.0,
"cpd00790_C":-1.0
},
"lower_bound":0.0,
"upper_bound":1000.0,
"gene_reaction_rule":"ACIAD3382 or ACIAD2314"
},
{
"id":"4_2_1_101_RXN",
"name":"trans-feruloyl-coa hydratase",
"metabolites":{
"cpd00001_C":-1.0,
"cpd00321_C":-1.0,
"cpd04511_C":1.0
},
"lower_bound":-1000.0,
"upper_bound":1000.0,
"gene_reaction_rule":"ACIAD1726"
},
{
"id":"SUCCCOASYN_RXN",
"name":"succinate--coa ligase (adp-forming)",
"metabolites":{
"cpd00002_C":1.0,
"cpd00008_C":-1.0,
"cpd00009_C":-1.0,
"cpd00010_C":1.0,
"cpd00036_C":1.0,
"cpd00078_C":-1.0
},
"lower_bound":-1000.0,
"upper_bound":1000.0,
"gene_reaction_rule":"ACIAD2873 and ACIAD2872"
},
{
"id":"EXF_VANILLATE_E",
"name":"exchange flux of vanillate",
"metabolites":{
"cpd04073_E":-1.0
},
"lower_bound":-0.0,
"upper_bound":1000.0,
"gene_reaction_rule":""
},
{
"id":"TRANS_RXN_ACET",
"name":"transport of acetate [extraorganism;cytosol](passive)",
"metabolites":{
"cpd00029_C":1.0,
"cpd00029_E":-1.0
},
"lower_bound":-1000.0,
"upper_bound":1000.0,
"gene_reaction_rule":""
},
{
"id":"CPM_KDOSYNTH_RXN",
"name":"3-deoxy-manno-octulosonate cytidylyltransferase",
"metabolites":{
"cpd00012_C":1.0,
"cpd00052_C":-1.0,
"cpd00875_C":-1.0,
"cpd02546_C":1.0
},
"lower_bound":0.0,
"upper_bound":1000.0,
"gene_reaction_rule":"ACIAD2362"
},
{
"id":"EXF_CARBON_DIOXIDE_E",
"name":"exchange flux of co2",
"metabolites":{
"cpd00011_E":-1.0
},
"lower_bound":-0.0,
"upper_bound":1000.0,
"gene_reaction_rule":""
},
{
"id":"6_3_5_6_RXN",
"name":"asparaginyl-trna synthetase (glutamine-hydrolyzing)",
"metabolites":{
"ASPARTYL_TRNAASN_C":-1.0,
"CHARGED_ASN_TRNAS_C":1.0,
"cpd00002_C":-1.0,
"cpd00008_C":1.0,
"cpd00009_C":1.0,
"cpd00023_C":1.0,
"cpd00053_C":-1.0
},
"lower_bound":0.0,
"upper_bound":1000.0,
"gene_reaction_rule":"ACIAD0822 and ACIAD0823 and ACIAD0824"
},
{
"id":"RIBOFLAVIN_SYN_RXN",
"name":"riboflavin synthase",
"metabolites":{
"cpd00220_C":1.0,
"cpd02656_C":-2.0,
"cpd02882_C":1.0
},
"lower_bound":-1000.0,
"upper_bound":1000.0,
"gene_reaction_rule":"ACIAD0249"
},
{
"id":"5_5_1_2_RXN",
"name":"3-carboxy-cis,cis-muconate cycloisomerase",
"metabolites":{
"cpd00856_C":1.0,
"cpd00938_C":-1.0
},
"lower_bound":-1000.0,
"upper_bound":1000.0,
"gene_reaction_rule":"ACIAD1707"
},
{
"id":"UNDECAPRENYL_DIPHOSPHATASE_RXN",
"name":"undecaprenyl-diphosphatase",
"metabolites":{
"cpd00001_C":-1.0,
"cpd00009_C":1.0,
"cpd00286_C":1.0,
"cpd02229_C":-1.0
},
"lower_bound":0.0,
"upper_bound":1000.0,
"gene_reaction_rule":"ACIAD1642 or ACIAD0678"
},
{
"id":"TRANS_RXN_FE_NRAMP",
"name":"transport of fe+2 [extraorganism;cytosol](secondary symport)",
"metabolites":{
"cpd00021_C":1.0,
"cpd00021_E":-1.0,
"cpd00067_C":1.0,
"cpd00067_P":-1.0
},
"lower_bound":0.0,
"upper_bound":1000.0,
"gene_reaction_rule":"ACIAD2521"
},
{
"id":"AFATTYACID_FREE_ASSEMBLY_RXN",
"name":"generic free fatty acid assembly",
"metabolites":{
"AFATTYACID_FREE_W_C":0.2556,
"cpd00214_C":-0.28,
"cpd00536_C":-0.09,
"cpd03847_C":-0.056,
"cpd34461_C":-0.574
},
"lower_bound":-1000.0,
"upper_bound":1000.0,
"gene_reaction_rule":""
},
{
"id":"ASPAMINOTRANS_RXN",
"name":"aspartate aminotransferase",
"metabolites":{
"cpd00023_C":-1.0,
"cpd00024_C":1.0,
"cpd00032_C":-1.0,
"cpd00041_C":1.0
},
"lower_bound":-1000.0,
"upper_bound":1000.0,
"gene_reaction_rule":"ACIAD2574 or ACIAD2087"
},
{
"id":"H2PTEROATESYNTH_RXN",
"name":"dihydropteroate synthase",
"metabolites":{
"cpd00012_C":1.0,
"cpd00443_C":-1.0,
"cpd00683_C":1.0,
"cpd02920_C":-1.0
},
"lower_bound":0.0,
"upper_bound":1000.0,
"gene_reaction_rule":"ACIAD2852"
},
{
"id":"RXN_7609",
"name":"dgmp nucleotidase",
"metabolites":{
"cpd00001_C":-1.0,
"cpd00009_C":1.0,
"cpd00126_C":-1.0,
"cpd00311_C":1.0
},
"lower_bound":0.0,
"upper_bound":1000.0,
"gene_reaction_rule":"ACIAD0015 or ACIAD1227"
},
{
"id":"RIBOKIN_RXN",
"name":"ribokinase",
"metabolites":{
"cpd00002_C":-1.0,
"cpd00008_C":1.0,
"cpd00101_C":1.0,
"cpd00105_C":-1.0
},
"lower_bound":0.0,
"upper_bound":1000.0,
"gene_reaction_rule":"ACIAD2392"
},
{
"id":"TRANS_RXN_TRP",
"name":"transport of l-tryptophan [extraorganism;cytosol](secondary symport)",
"metabolites":{
"cpd00065_C":1.0,
"cpd00065_E":-1.0,
"cpd00067_C":1.0,
"cpd00067_P":-1.0
},
"lower_bound":-1000.0,
"upper_bound":1000.0,
"gene_reaction_rule":"ACIAD1343 or ACIAD2969"
},
{
"id":"ACYL_COA_WAX_ASSEMBLY_RXN",
"name":"generic acyl-coa of wax assembly",
"metabolites":{
"ACYL_COA_WAX_C":1.0,
"cpd00134_C":-0.24,
"cpd16627_C":-0.315,
"cpd16638_C":-0.445
},
"lower_bound":-1000.0,
"upper_bound":1000.0,
"gene_reaction_rule":""
},
{
"id":"EXF_PURINE_E",
"name":"exchange flux of purine",
"metabolites":{
"cpd00360_E":-1.0
},
"lower_bound":-0.0,
"upper_bound":1000.0,
"gene_reaction_rule":""
},
{
"id":"TRANS_RXNIM_5731",
"name":"transport of d-serine [extraorganism;cytosol](passive)",
"metabolites":{
"cpd00550_C":1.0,
"cpd00550_E":-1.0
},
"lower_bound":0.0,
"upper_bound":1000.0,
"gene_reaction_rule":"ACIAD0118 or ACIAD2662"
},
{
"id":"TRANS_RXNIM_5733",
"name":"transport of d-alanine [extraorganism;cytosol](passive)",
"metabolites":{
"cpd00117_C":1.0,
"cpd00117_E":-1.0
},
"lower_bound":0.0,
"upper_bound":1000.0,
"gene_reaction_rule":"ACIAD0118 or ACIAD2662"
},
{
"id":"TRANS_RXN_ANTHRANILATE",
"name":"transport of 2-aminobenzoate [extraorganism;cytosol](passive)",
"metabolites":{
"cpd00093_C":1.0,
"cpd00093_E":-1.0
},
"lower_bound":-1000.0,
"upper_bound":1000.0,
"gene_reaction_rule":""
},
{
"id":"C12_0_3HYDROXYCOA_DEHYDROGENASE_RXN",
"name":"3oxoacyl-coa thiolase (c10:0)",
"metabolites":{
"cpd00003_C":-1.0,
"cpd00004_C":1.0,
"cpd03116_C":-1.0,
"cpd03117_C":1.0
},
"lower_bound":0.0,
"upper_bound":1000.0,
"gene_reaction_rule":"ACIAD0335 or ACIAD1565 or ACIAD2416"
},
{
"id":"GLUCONOLACT_RXN",
"name":"gluconolactonase",
"metabolites":{
"cpd00001_P":-1.0,
"cpd00170_P":-1.0,
"cpd00222_P":1.0
},
"lower_bound":-1000.0,
"upper_bound":1000.0,
"gene_reaction_rule":"ACIAD2819 or SPONT"
},
{
"id":"MALONYL_COA_ACP_TRANSACYL_RXN",
"name":"malonyl-coa-acp transacylase",
"metabolites":{
"ACP_C":-1.0,
"MALONYL_ACP_C":1.0,
"cpd00010_C":1.0,
"cpd00070_C":-1.0
},
"lower_bound":-1000.0,
"upper_bound":1000.0,
"gene_reaction_rule":"ACIAD0870 and ACIAD0872"
},
{
"id":"TRANS_RXN_MALONATE",
"name":"transport of malonate [extraorganism;cytosol](secondary symport)",
"metabolites":{
"cpd00067_C":1.0,
"cpd00067_P":-1.0,
"cpd00308_C":1.0,
"cpd00308_E":-1.0
},
"lower_bound":-1000.0,
"upper_bound":1000.0,
"gene_reaction_rule":"ACIAD0710 or ACIAD2956"
},
{
"id":"PANTOATE_BETA_ALANINE_LIG_RXN",
"name":"pantoate--beta-alanine ligase",
"metabolites":{
"cpd00002_C":-1.0,
"cpd00012_C":1.0,
"cpd00018_C":1.0,
"cpd00085_C":-1.0,
"cpd00408_C":-1.0,
"cpd00644_C":1.0
},
"lower_bound":-1000.0,
"upper_bound":1000.0,
"gene_reaction_rule":"ACIAD3060"
},
{
"id":"EXF_BENZOATE_E",
"name":"exchange flux of benzoate",
"metabolites":{
"cpd00153_E":-1.0
},
"lower_bound":-0.0,
"upper_bound":1000.0,
"gene_reaction_rule":""
},
{
"id":"ADENYLOSUCCINATE_SYNTHASE_RXN",
"name":"adenylosuccinate synthase",
"metabolites":{
"cpd00009_C":1.0,
"cpd00031_C":1.0,
"cpd00038_C":-1.0,
"cpd00041_C":-1.0,
"cpd00114_C":-1.0,
"cpd02375_C":1.0
},
"lower_bound":-1000.0,
"upper_bound":1000.0,
"gene_reaction_rule":"ACIAD1258"
},
{
"id":"PYRROLINECARBDEHYDROG_RXN",
"name":"1-pyrroline-5-carboxylate dehydrogenase",
"metabolites":{
"cpd00001_C":-2.0,
"cpd00003_C":-1.0,
"cpd00004_C":1.0,
"cpd00023_C":1.0,
"cpd02431_C":-1.0
},
"lower_bound":0.0,
"upper_bound":1000.0,
"gene_reaction_rule":"ACIAD1646"
},
{
"id":"F16ALDOLASE_RXN",
"name":"fructose-bisphosphate aldolase",
"metabolites":{
"cpd00095_C":-1.0,
"cpd00102_C":-1.0,
"cpd00290_C":1.0
},
"lower_bound":-1000.0,
"upper_bound":1000.0,
"gene_reaction_rule":"ACIAD1925"
},
{
"id":"TRANS_RXN_GLYCOLLATE_OUT",
"name":"transport of glycolate [cytosol;extraorganism](passive)",
"metabolites":{
"cpd00139_C":-1.0,
"cpd00139_E":1.0
},
"lower_bound":0.0,
"upper_bound":1000.0,
"gene_reaction_rule":""
},
{
"id":"GDPKIN_RXN",
"name":"gdp kinase",
"metabolites":{
"cpd00002_C":-1.0,
"cpd00008_C":1.0,
"cpd00031_C":-1.0,
"cpd00038_C":1.0
},
"lower_bound":-1000.0,
"upper_bound":1000.0,
"gene_reaction_rule":"ACIAD0556"
},
{
"id":"DXS_RXN",
"name":"1-deoxy-d-xylulose 5-phosphate synthase",
"metabolites":{
"cpd00011_C":1.0,
"cpd00020_C":-1.0,
"cpd00102_C":-1.0,
"cpd08289_C":1.0
},
"lower_bound":0.0,
"upper_bound":1000.0,
"gene_reaction_rule":"ACIAD3247"
},
{
"id":"CDPREDUCT_RXN",
"name":"cdp reductase",
"metabolites":{
"cpd00001_C":1.0,
"cpd00096_C":-1.0,
"cpd00533_C":1.0,
"cpd11420_C":1.0,
"cpd11421_C":-1.0
},
"lower_bound":-1000.0,
"upper_bound":1000.0,
"gene_reaction_rule":"ACIAD0724 and ACIAD0722"
},
{
"id":"TRANS_RXN_C12_0",
"name":"transport of dodecanoate [extraorganism;cytosol](secondary symport)",
"metabolites":{
"cpd00067_C":1.0,
"cpd00067_P":-1.0,
"cpd01741_C":1.0,
"cpd01741_E":-1.0
},
"lower_bound":-1000.0,
"upper_bound":1000.0,
"gene_reaction_rule":"ACIAD0835 or ACIAD1427 or ACIAD2788 or ACIAD3392"
},
{
"id":"CYSTEINE_TRNA_LIGASE_RXN",
"name":"cysteine--trna ligase",
"metabolites":{
"CHARGED_CYS_TRNAS_C":1.0,
"CYS_TRNAS_C":-1.0,
"cpd00002_C":-1.0,
"cpd00012_C":1.0,
"cpd00018_C":1.0,
"cpd00084_C":-1.0
},
"lower_bound":0.0,
"upper_bound":1000.0,
"gene_reaction_rule":"ACIAD1481"
},
{
"id":"GART_RXN",
"name":"phosphoribosylglycinamide formyltransferase",
"metabolites":{
"cpd00087_C":1.0,
"cpd00201_C":-1.0,
"cpd02394_C":-1.0,
"cpd02678_C":1.0
},
"lower_bound":0.0,
"upper_bound":1000.0,
"gene_reaction_rule":"ACIAD2635"
},
{
"id":"GLYCINE_TRNA_LIGASE_RXN",
"name":"glycine--trna ligase",
"metabolites":{
"CHARGED_GLY_TRNAS_C":1.0,
"GLY_TRNAS_C":-1.0,
"cpd00002_C":-1.0,
"cpd00012_C":1.0,
"cpd00018_C":1.0,
"cpd00033_C":-1.0
},
"lower_bound":0.0,
"upper_bound":1000.0,
"gene_reaction_rule":"ACIAD3268 and ACIAD3269"
},
{
"id":"ALKANE_1_MONOOXYGENASE_RXN",
"name":"octane-1 monooxygenase",
"metabolites":{
"cpd00001_C":1.0,
"cpd00003_C":1.0,
"cpd00004_C":-1.0,
"cpd00007_C":-1.0,
"cpd00562_C":1.0,
"cpd00998_C":-1.0
},
"lower_bound":0.0,
"upper_bound":1000.0,
"gene_reaction_rule":"ACIAD1411 and ACIAD1065"
},
{
"id":"GUANINE_DEAMINASE_RXN",
"name":"guanine deaminase",
"metabolites":{
"cpd00001_C":-1.0,
"cpd00013_C":1.0,
"cpd00207_C":-1.0,
"cpd00309_C":1.0
},
"lower_bound":0.0,
"upper_bound":1000.0,
"gene_reaction_rule":"ACIAD3668"
},
{
"id":"ALLOPHANATE_HYDROLASE_RXN",
"name":"allophanate hydrolase",
"metabolites":{
"cpd00001_C":-1.0,
"cpd00011_C":2.0,
"cpd00013_C":2.0,
"cpd00742_C":-1.0
},
"lower_bound":0.0,
"upper_bound":1000.0,
"gene_reaction_rule":"ACIAD2518"
},
{
"id":"UDP_POLYSACCH_LPS_ASSEMBLY_RXN",
"name":"generic udp-polysaccharide of lps assembly",
"metabolites":{
"UDP_POLYSACCH_LPS_C":1.0,
"cpd00014_C":0.67,
"cpd00026_C":-1.67,
"cpd00031_C":1.66,
"cpd00083_C":-1.66,
"cpd00297_C":1.67,
"cpd02113_C":-1.67
},
"lower_bound":0.0,
"upper_bound":1000.0,
"gene_reaction_rule":""
},
{
"id":"TRANS_RXN_OXYGEN_MOLECULE",
"name":"transport of o2 [extraorganism;cytosol](passive)",
"metabolites":{
"cpd00007_C":1.0,
"cpd00007_E":-1.0
},
"lower_bound":-1000.0,
"upper_bound":1000.0,
"gene_reaction_rule":""
},
{
"id":"PORPHOBILSYNTH_RXN",
"name":"porphobilinogen synthase",
"metabolites":{
"cpd00001_C":2.0,
"cpd00338_C":-2.0,
"cpd00689_C":1.0
},
"lower_bound":-1000.0,
"upper_bound":1000.0,
"gene_reaction_rule":"ACIAD0923"
},
{
"id":"EXF_PHE_E",
"name":"exchange flux of l-phenylalanine",
"metabolites":{
"cpd00066_E":-1.0
},
"lower_bound":-0.0,
"upper_bound":1000.0,
"gene_reaction_rule":""
},
{
"id":"ADENOSYLHOMOCYSTEINASE_RXN",
"name":"adenosylhomocysteinase",
"metabolites":{
"cpd00001_C":-1.0,
"cpd00019_C":-1.0,
"cpd00135_C":1.0,
"cpd00182_C":1.0
},
"lower_bound":0.0,
"upper_bound":1000.0,
"gene_reaction_rule":"ACIAD2282"
},
{
"id":"HYPOXANPRIBOSYLTRAN_RXN",
"name":"hypoxanthine phosphoribosyltransferase",
"metabolites":{
"cpd00012_C":1.0,
"cpd00103_C":-1.0,
"cpd00114_C":1.0,
"cpd00226_C":-1.0
},
"lower_bound":0.0,
"upper_bound":1000.0,
"gene_reaction_rule":"ACIAD3669"
},
{
"id":"TRANS_RXN_PROPIONATE",
"name":"transport of propionate [extraorganism;cytosol](passive)",
"metabolites":{
"cpd00141_C":1.0,
"cpd00141_E":-1.0
},
"lower_bound":-1000.0,
"upper_bound":1000.0,
"gene_reaction_rule":""
},
{
"id":"TRANS_RXN_GUANOSINE",
"name":"transport of guanosine [extraorganism;cytosol](secondary symport)",
"metabolites":{
"cpd00067_C":1.0,
"cpd00067_P":-1.0,
"cpd00311_C":1.0,
"cpd00311_E":-1.0
},
"lower_bound":0.0,
"upper_bound":1000.0,
"gene_reaction_rule":"ACIAD2389"
},
{
"id":"R11_RXN",
"name":"acetate coa-transferase",
"metabolites":{
"cpd00022_C":1.0,
"cpd00029_C":-1.0,
"cpd00120_C":-1.0,
"cpd00211_C":1.0
},
"lower_bound":-1000.0,
"upper_bound":1000.0,
"gene_reaction_rule":"ACIAD2514 and ACIAD2513"
},
{
"id":"TRANS_RXNIM_5895",
"name":"transport of toluene [extraorganism;cytosol](primary active atp)",
"metabolites":{
"cpd00001_C":-1.0,
"cpd00002_C":-1.0,
"cpd00008_C":1.0,
"cpd00009_C":1.0,
"cpd01034_C":1.0,
"cpd01034_E":-1.0
},
"lower_bound":0.0,
"upper_bound":1000.0,
"gene_reaction_rule":"ACIAD3241 and ACIAD3242 and ACIAD3243 and ACIAD3244"
},
{
"id":"GLUTARYL_COA_DEHYDROG_RXN",
"name":"glutaryl-coa dehydrogenase",
"metabolites":{
"cpd00015_C":-1.0,
"cpd00413_C":-1.0,
"cpd00982_C":1.0,
"cpd01599_C":1.0
},
"lower_bound":0.0,
"upper_bound":1000.0,
"gene_reaction_rule":"ACIAD1693"
},
{
"id":"ASPDECARBOX_RXN",
"name":"aspartate 1-decarboxylase",
"metabolites":{
"cpd00011_C":1.0,
"cpd00041_C":-1.0,
"cpd00085_C":1.0
},
"lower_bound":0.0,
"upper_bound":1000.0,
"gene_reaction_rule":"ACIAD2911"
},
{
"id":"THIAZOLSYN2_RXN",
"name":"thiazole phosphate synthesis",
"metabolites":{
"cpd00001_C":1.0,
"cpd00002_C":-1.0,
"cpd00011_C":1.0,
"cpd00012_C":1.0,
"cpd00018_C":1.0,
"cpd00035_C":1.0,
"cpd00069_C":-1.0,
"cpd00084_C":-1.0,
"cpd02654_C":1.0,
"cpd08289_C":-1.0,
"cpd15378_C":1.0
},
"lower_bound":-1000.0,
"upper_bound":1000.0,
"gene_reaction_rule":"ACIAD1308 and ACIAD1309 and ACIAD2411 and ACIAD1404"
},
{
"id":"TRANS_RXN_C6_2_c2_c4_",
"name":"transport of sorbate [extraorganism;cytosol](secondary symport)",
"metabolites":{
"cpd00067_C":1.0,
"cpd00067_P":-1.0,
"cpd16648_C":1.0,
"cpd16648_E":-1.0
},
"lower_bound":-1000.0,
"upper_bound":1000.0,
"gene_reaction_rule":"ACIAD0835 or ACIAD1427 or ACIAD2788 or ACIAD3392"
},
{
"id":"TRANS_RXN_CPD_205",
"name":"transport of pimelate [extraorganism;cytosol](secondary symport)",
"metabolites":{
"cpd00067_C":1.0,
"cpd00067_P":-1.0,
"cpd01727_C":1.0,
"cpd01727_E":-1.0
},
"lower_bound":-1000.0,
"upper_bound":1000.0,
"gene_reaction_rule":"ACIAD1694"
},
{
"id":"4OH2OXOGLUTARALDOL_RXN",
"name":"4-hydroxy-2-oxoglutarate aldolase",
"metabolites":{
"cpd00020_C":1.0,
"cpd00040_C":1.0,
"cpd00830_C":-1.0
},
"lower_bound":-1000.0,
"upper_bound":1000.0,
"gene_reaction_rule":"ACIAD0543"
},
{
"id":"DIHYDROFOLATEREDUCT_RXN",
"name":"dihydrofolate reductase",
"metabolites":{
"cpd00005_C":1.0,
"cpd00006_C":-1.0,
"cpd00087_C":-1.0,
"cpd00330_C":1.0
},
"lower_bound":-1000.0,
"upper_bound":1000.0,
"gene_reaction_rule":"ACIAD0514"
},
{
"id":"C16_1_C7_3OXOCOA_THIOLASE_RXN",
"name":"3oxoacyl-coa thiolase (c16:1(c7))",
"metabolites":{
"cpd00010_C":-1.0,
"cpd00022_C":1.0,
"cpd16617_C":1.0,
"cpd32368_C":-1.0
},
"lower_bound":-1000.0,
"upper_bound":1000.0,
"gene_reaction_rule":"ACIAD0334 or ACIAD1450 or ACIAD1706 or ACIAD1977 or ACIAD1689 or ACIAD2988 or ACIAD1569"
},
{
"id":"NITRILE_HYDRATASE_RXN",
"name":"propane nitrile hydratase",
"metabolites":{
"cpd00001_C":-1.0,
"cpd25464_C":-1.0,
"cpd27812_C":1.0
},
"lower_bound":0.0,
"upper_bound":1000.0,
"gene_reaction_rule":"ACIAD1615 and ACIAD1616"
},
{
"id":"C12_0_2ENOYLCOA_HYDRATASE_RXN",
"name":"enoyl-coa hydratase (c12:0)",
"metabolites":{
"cpd00001_C":-1.0,
"cpd02060_C":-1.0,
"cpd03116_C":1.0
},
"lower_bound":0.0,
"upper_bound":1000.0,
"gene_reaction_rule":"ACIAD1692 or ACIAD1726 or ACIAD1820 or ACIAD1829 or ACIAD2351 or ACIAD3073 or ACIAD1609 or ACIAD0335 or ACIAD1608 or ACIAD1701 or ACIAD1565 or ACIAD2416"
},
{
"id":"GLURS_RXN",
"name":"glutamate--trna ligase",
"metabolites":{
"CHARGED_GLT_TRNAS_C":1.0,
"GLT_TRNAS_C":-1.0,
"cpd00002_C":-1.0,
"cpd00012_C":1.0,
"cpd00018_C":1.0,
"cpd00023_C":-1.0
},
"lower_bound":-1000.0,
"upper_bound":1000.0,
"gene_reaction_rule":"ACIAD3371"
},
{
"id":"ERYTHRON4PDEHYDROG_RXN",
"name":"erythronate-4-phosphate dehydrogenase",
"metabolites":{
"cpd00003_C":-1.0,
"cpd00004_C":1.0,
"cpd02147_C":-1.0,
"cpd03606_C":1.0
},
"lower_bound":-1000.0,
"upper_bound":1000.0,
"gene_reaction_rule":"ACIAD2667"
},
{
"id":"GUANPRIBOSYLTRAN_RXN",
"name":"guanine phosphoribosyltransferase",
"metabolites":{
"cpd00012_C":1.0,
"cpd00103_C":-1.0,
"cpd00126_C":1.0,
"cpd00207_C":-1.0
},
"lower_bound":0.0,
"upper_bound":1000.0,
"gene_reaction_rule":"ACIAD3669"
},
{
"id":"EXF_C5_0_E",
"name":"exchange flux of pentanoate",
"metabolites":{
"cpd00597_E":-1.0
},
"lower_bound":-0.0,
"upper_bound":1000.0,
"gene_reaction_rule":""
},
{
"id":"HEMEOSYN_RXN",
"name":"protoheme ix farnesyltransferase",
"metabolites":{
"HEME_O_C":1.0,
"cpd00001_C":-1.0,
"cpd00012_C":1.0,
"cpd00028_C":-1.0,
"cpd00350_C":-1.0
},
"lower_bound":-1000.0,
"upper_bound":1000.0,
"gene_reaction_rule":"ACIAD2429"
},
{
"id":"DCYSDESULF_RXN",
"name":"d-cysteine desulfhydrase",
"metabolites":{
"cpd00001_C":-1.0,
"cpd00013_C":1.0,
"cpd00020_C":1.0,
"cpd00239_C":1.0,
"cpd00587_C":-1.0
},
"lower_bound":0.0,
"upper_bound":1000.0,
"gene_reaction_rule":"ACIAD0753"
},
{
"id":"DTDPGLUCDEHYDRAT_RXN",
"name":"dtdp-glucose 4,6-dehydratase",
"metabolites":{
"cpd00001_C":1.0,
"cpd00521_C":1.0,
"cpd00626_C":-1.0
},
"lower_bound":0.0,
"upper_bound":1000.0,
"gene_reaction_rule":"ACIAD0076"
},
{
"id":"EXF_BETAINE_E",
"name":"exchange flux of betaine",
"metabolites":{
"cpd00540_E":-1.0
},
"lower_bound":-0.0,
"upper_bound":1000.0,
"gene_reaction_rule":""
},
{
"id":"RXN_7607",
"name":"imp nucleotidase",
"metabolites":{
"cpd00001_C":-1.0,
"cpd00009_C":1.0,
"cpd00114_C":-1.0,
"cpd00246_C":1.0
},
"lower_bound":0.0,
"upper_bound":1000.0,
"gene_reaction_rule":"ACIAD0015 or ACIAD1227"
},
{
"id":"C10_0_3OXOCOA_THIOLASE_RXN",
"name":"3oxoacyl-coa thiolase (c10:0)",
"metabolites":{
"cpd00010_C":-1.0,
"cpd00022_C":1.0,
"cpd01335_C":1.0,
"cpd03119_C":-1.0
},
"lower_bound":-1000.0,
"upper_bound":1000.0,
"gene_reaction_rule":"ACIAD0334 or ACIAD1450 or ACIAD1706 or ACIAD1977 or ACIAD1689 or ACIAD2988 or ACIAD1569"
},
{
"id":"FGAMSYN_RXN",
"name":"phosphoribosylformylglycinamidine synthase",
"metabolites":{
"cpd00001_C":-1.0,
"cpd00002_C":-1.0,
"cpd00008_C":1.0,
"cpd00009_C":1.0,
"cpd00023_C":1.0,
"cpd00053_C":-1.0,
"cpd02678_C":-1.0,
"cpd02826_C":1.0
},
"lower_bound":0.0,
"upper_bound":1000.0,
"gene_reaction_rule":"ACIAD2612"
},
{
"id":"1_1_99_25_RXN",
"name":"quinate dehydrogenase (pyrroloquinoline-quinone)",
"metabolites":{
"cpd00097_P":-1.0,
"cpd00248_P":-1.0,
"cpd00699_P":1.0,
"cpd00986_P":1.0
},
"lower_bound":-1000.0,
"upper_bound":1000.0,
"gene_reaction_rule":"ACIAD1716"
},
{
"id":"C8_0_3OXOCOA_THIOLASE_RXN",
"name":"3oxoacyl-coa thiolase (c8:0)",
"metabolites":{
"cpd00010_C":-1.0,
"cpd00022_C":1.0,
"cpd03121_C":-1.0,
"cpd03124_C":1.0
},
"lower_bound":-1000.0,
"upper_bound":1000.0,
"gene_reaction_rule":"ACIAD0334 or ACIAD1450 or ACIAD1706 or ACIAD1977 or ACIAD1689 or ACIAD2988 or ACIAD1569"
},
{
"id":"ADDALT_RXN",
"name":"deoxyadenosine deaminase",
"metabolites":{
"cpd00001_C":-1.0,
"cpd00013_C":1.0,
"cpd00438_C":-1.0,
"cpd03279_C":1.0
},
"lower_bound":0.0,
"upper_bound":1000.0,
"gene_reaction_rule":"ACIAD1245"
},
{
"id":"DUTP_PYROP_RXN",
"name":"dutp pyrophosphatase",
"metabolites":{
"cpd00001_C":-1.0,
"cpd00012_C":1.0,
"cpd00299_C":1.0,
"cpd00358_C":-1.0
},
"lower_bound":0.0,
"upper_bound":1000.0,
"gene_reaction_rule":"ACIAD0901"
},
{
"id":"SUCCDIAMINOPIMDESUCC_RXN",
"name":"succinyl-diaminopimelate desuccinylase",
"metabolites":{
"cpd00001_C":-1.0,
"cpd00036_C":1.0,
"cpd00504_C":1.0,
"cpd02698_C":-1.0
},
"lower_bound":-1000.0,
"upper_bound":1000.0,
"gene_reaction_rule":"ACIAD0791"
},
{
"id":"DETHIOBIOTIN_SYN_RXN",
"name":"dethiobiotin synthase",
"metabolites":{
"cpd00002_C":-1.0,
"cpd00008_C":1.0,
"cpd00009_C":1.0,
"cpd00011_C":-1.0,
"cpd00764_C":-1.0,
"cpd01311_C":1.0
},
"lower_bound":0.0,
"upper_bound":1000.0,
"gene_reaction_rule":"ACIAD0859"
},
{
"id":"EXF_AMMONIUM_E",
"name":"exchange flux of nh4+",
"metabolites":{
"cpd00013_E":-1.0
},
"lower_bound":-1000.0,
"upper_bound":1000.0,
"gene_reaction_rule":""
},
{
"id":"RXN_7913",
"name":"dcmp kinase",
"metabolites":{
"cpd00002_C":-1.0,
"cpd00008_C":1.0,
"cpd00206_C":-1.0,
"cpd00533_C":1.0
},
"lower_bound":0.0,
"upper_bound":1000.0,
"gene_reaction_rule":"ACIAD2348"
},
{
"id":"THIOREDOXIN_REDUCT_NADPH_RXN",
"name":"thioredoxin reductase (nadph)",
"metabolites":{
"cpd00005_C":-1.0,
"cpd00006_C":1.0,
"cpd11420_C":-1.0,
"cpd11421_C":1.0
},
"lower_bound":-1000.0,
"upper_bound":1000.0,
"gene_reaction_rule":"ACIAD0890"
},
{
"id":"C7_0_CARBOXY2ENOYLCOA_HYDRATASE_RXN",
"name":"dicarboxylate enoyl-coa hydratase (c7:0)",
"metabolites":{
"cpd00001_C":-1.0,
"cpd04103_C":1.0,
"cpd04111_C":-1.0
},
"lower_bound":0.0,
"upper_bound":1000.0,
"gene_reaction_rule":"ACIAD1692"
},
{
"id":"C8_0_COA_THIOLASE_RXN",
"name":"acyl-coa thiolase (c8:0)",
"metabolites":{
"cpd00001_C":-1.0,
"cpd00010_C":1.0,
"cpd01335_C":-1.0,
"cpd03846_C":1.0
},
"lower_bound":0.0,
"upper_bound":1000.0,
"gene_reaction_rule":"ACIAD0341 or ACIAD3231 or ACIAD1023"
},
{
"id":"1_1_1_259_RXN",
"name":"3-hydroxypimeloyl-coa dehydrogenase",
"metabolites":{
"cpd00003_C":-1.0,
"cpd00004_C":1.0,
"cpd04103_C":-1.0,
"cpd04104_C":1.0
},
"lower_bound":0.0,
"upper_bound":1000.0,
"gene_reaction_rule":"ACIAD1690"
},
{
"id":"METHYLISOCITRATE_LYASE_RXN",
"name":"methylisocitrate lyase",
"metabolites":{
"cpd00020_C":1.0,
"cpd00036_C":1.0,
"cpd02799_C":-1.0
},
"lower_bound":0.0,
"upper_bound":1000.0,
"gene_reaction_rule":"ACIAD2758"
},
{
"id":"C12_0_2ENOYLACP_REDUCTASE_RXN",
"name":"2enoyl-acp reductase (c12:0)",
"metabolites":{
"C12_0_2ENOYLACP_C":-1.0,
"C12_0_ACP_C":1.0,
"cpd00003_C":1.0,
"cpd00004_C":-1.0
},
"lower_bound":0.0,
"upper_bound":1000.0,
"gene_reaction_rule":"ACIAD3116"
},
{
"id":"ACONITATEHYDR_RXN",
"name":"aconitase (isocitrate hydro-lyase)",
"metabolites":{
"cpd00001_C":-1.0,
"cpd00260_C":1.0,
"cpd00331_C":-1.0
},
"lower_bound":-1000.0,
"upper_bound":1000.0,
"gene_reaction_rule":"ACIAD3090 or ACIAD2395 or ACIAD2755"
},
{
"id":"ATPPHOSPHORIBOSYLTRANS_RXN",
"name":"atp phosphoribosyltransferase",
"metabolites":{
"cpd00002_C":-1.0,
"cpd00012_C":1.0,
"cpd00103_C":-1.0,
"cpd01775_C":1.0
},
"lower_bound":-1000.0,
"upper_bound":1000.0,
"gene_reaction_rule":"ACIAD0661 and ACIAD1257"
},
{
"id":"1_1_1_215_RXN",
"name":"gluconate 2-dehydrogenase",
"metabolites":{
"cpd00005_C":1.0,
"cpd00006_C":-1.0,
"cpd00222_C":-1.0,
"cpd00480_C":1.0
},
"lower_bound":-1000.0,
"upper_bound":1000.0,
"gene_reaction_rule":"ACIAD1327"
},
{
"id":"PRPPAMIDOTRANS_RXN",
"name":"amidophosphoribosyltransferase",
"metabolites":{
"cpd00001_C":-1.0,
"cpd00012_C":1.0,
"cpd00023_C":1.0,
"cpd00053_C":-1.0,
"cpd00103_C":-1.0,
"cpd01982_C":1.0
},
"lower_bound":0.0,
"upper_bound":1000.0,
"gene_reaction_rule":"ACIAD1323"
},
{
"id":"C5_0_COATRANSFERASE_RXN",
"name":"acyl coa transferase (c5:0)",
"metabolites":{
"cpd00002_C":-1.0,
"cpd00010_C":-1.0,
"cpd00012_C":1.0,
"cpd00018_C":1.0,
"cpd00597_C":-1.0,
"cpd00660_C":1.0
},
"lower_bound":0.0,
"upper_bound":1000.0,
"gene_reaction_rule":"ACIAD1818 or ACIAD0235 or ACIAD1724 or ACIAD2106"
},
{
"id":"UROPORIIIMETHYLTRANSA_RXN",
"name":"uroporphyrinogen-iii c-methyltransferase",
"metabolites":{
"cpd00017_C":-2.0,
"cpd00019_C":2.0,
"cpd00774_C":-1.0,
"cpd01620_C":1.0
},
"lower_bound":0.0,
"upper_bound":1000.0,
"gene_reaction_rule":"ACIAD2934"
},
{
"id":"METHYLENETHFDEHYDROG_NADP_RXN",
"name":"methylenetetrahydrofolate dehydrogenase (nadp+)",
"metabolites":{
"cpd00005_C":1.0,
"cpd00006_C":-1.0,
"cpd00125_C":-1.0,
"cpd00347_C":1.0
},
"lower_bound":-1000.0,
"upper_bound":1000.0,
"gene_reaction_rule":"ACIAD2553 or ACIAD2847 or ACIAD1783"
},
{
"id":"ISPH2_RXN",
"name":"1-hydroxy-2-methyl-2-(e)-butenyl 4-diphosphate reductase (ipdp)",
"metabolites":{
"cpd00001_C":1.0,
"cpd00015_C":2.0,
"cpd00067_C":-2.0,
"cpd00113_C":1.0,
"cpd00982_C":-2.0,
"cpd08615_C":-1.0
},
"lower_bound":-1000.0,
"upper_bound":1000.0,
"gene_reaction_rule":"ACIAD3322"
},
{
"id":"N_ACETYLTRANSFER_RXN",
"name":"amino-acid n-acetyltransferase",
"metabolites":{
"cpd00010_C":1.0,
"cpd00022_C":-1.0,
"cpd00023_C":-1.0,
"cpd00477_C":1.0
},
"lower_bound":-1000.0,
"upper_bound":1000.0,
"gene_reaction_rule":"ACIAD0650 or ACIAD0039"
},
{
"id":"THRESYN_RXN",
"name":"threonine synthase",
"metabolites":{
"cpd00001_C":-1.0,
"cpd00009_C":1.0,
"cpd00161_C":1.0,
"cpd00809_C":-1.0
},
"lower_bound":0.0,
"upper_bound":1000.0,
"gene_reaction_rule":"ACIAD0263"
},
{
"id":"DCTP_PYROPHOSPHATASE_RXN",
"name":"dctp pyrophosphatase",
"metabolites":{
"cpd00001_C":-1.0,
"cpd00012_C":1.0,
"cpd00206_C":1.0,
"cpd00356_C":-1.0
},
"lower_bound":0.0,
"upper_bound":1000.0,
"gene_reaction_rule":"ACIAD3065"
},
{
"id":"ASPARTATE_TRNA_LIGASE_RXN",
"name":"aspartate--trna ligase",
"metabolites":{
"ASP_TRNAS_C":-1.0,
"CHARGED_ASP_TRNAS_C":1.0,
"cpd00002_C":-1.0,
"cpd00012_C":1.0,
"cpd00018_C":1.0,
"cpd00041_C":-1.0
},
"lower_bound":0.0,
"upper_bound":1000.0,
"gene_reaction_rule":"ACIAD0609"
},
{
"id":"IMP_DEHYDROG_RXN",
"name":"imp dehydrogenase",
"metabolites":{
"cpd00001_C":-1.0,
"cpd00003_C":-1.0,
"cpd00004_C":1.0,
"cpd00114_C":-1.0,
"cpd00497_C":1.0
},
"lower_bound":-1000.0,
"upper_bound":1000.0,
"gene_reaction_rule":"ACIAD3503"
},
{
"id":"DIACYLGLYKIN_RXN",
"name":"diacylglycerol kinase",
"metabolites":{
"L_PHOSPHATIDATE_TAG_C":1.0,
"cpd00002_C":-1.0,
"cpd00008_C":1.0,
"cpd11423_C":-1.0
},
"lower_bound":0.0,
"upper_bound":1000.0,
"gene_reaction_rule":"ACIAD2837"
},
{
"id":"EXF_TYR_E",
"name":"exchange flux of l-tyrosine",
"metabolites":{
"cpd00069_E":-1.0
},
"lower_bound":-0.0,
"upper_bound":1000.0,
"gene_reaction_rule":""
},
{
"id":"THI_P_SYN_RXN",
"name":"thiamine-phosphate pyrophosphorylase",
"metabolites":{
"cpd00012_C":1.0,
"cpd00793_C":1.0,
"cpd02654_C":-1.0,
"cpd02894_C":-1.0
},
"lower_bound":-1000.0,
"upper_bound":1000.0,
"gene_reaction_rule":"ACIAD1200"
},
{
"id":"EXF_C6_2_C2_C4_E",
"name":"exchange flux of sorbate",
"metabolites":{
"cpd16648_E":-1.0
},
"lower_bound":-0.0,
"upper_bound":1000.0,
"gene_reaction_rule":""
},
{
"id":"EXF_CPD_110_E",
"name":"exchange flux of salicylate",
"metabolites":{
"cpd00599_E":-1.0
},
"lower_bound":-0.0,
"upper_bound":1000.0,
"gene_reaction_rule":""
},
{
"id":"RIB5PISOM_RXN",
"name":"ribose 5-phosphate epimerase",
"metabolites":{
"cpd00101_C":-1.0,
"cpd00171_C":1.0
},
"lower_bound":-1000.0,
"upper_bound":1000.0,
"gene_reaction_rule":"ACIAD1358"
},
{
"id":"UTP_PYROPHOSPHATASE_RXN",
"name":"utp pyrophosphorylase",
"metabolites":{
"cpd00001_C":-1.0,
"cpd00012_C":1.0,
"cpd00062_C":-1.0,
"cpd00091_C":1.0
},
"lower_bound":0.0,
"upper_bound":1000.0,
"gene_reaction_rule":"ACIAD3065"
},
{
"id":"GUANYL_KIN_RXN",
"name":"gmp kinase",
"metabolites":{
"cpd00002_C":-1.0,
"cpd00008_C":1.0,
"cpd00031_C":1.0,
"cpd00126_C":-1.0
},
"lower_bound":-1000.0,
"upper_bound":1000.0,
"gene_reaction_rule":"ACIAD3324"
},
{
"id":"TRANS_RXNIM_5808",
"name":"transport of benzoate [extraorganism;periplasm](passive)",
"metabolites":{
"cpd00153_E":-1.0,
"cpd00153_P":1.0
},
"lower_bound":-1000.0,
"upper_bound":1000.0,
"gene_reaction_rule":"ACIAD1433"
},
{
"id":"3_OXOADIPATE_ENOL_LACTONASE_RXN",
"name":"3-oxoadipate enol-lactonase",
"metabolites":{
"cpd00001_C":-1.0,
"cpd00630_C":1.0,
"cpd02255_C":-1.0
},
"lower_bound":-1000.0,
"upper_bound":1000.0,
"gene_reaction_rule":"ACIAD1708 or ACIAD1451"
},
{
"id":"TRYPTOPHAN_TRNA_LIGASE_RXN",
"name":"tryptophan--trna ligase",
"metabolites":{
"CHARGED_TRP_TRNAS_C":1.0,
"TRP_TRNAS_C":-1.0,
"cpd00002_C":-1.0,
"cpd00012_C":1.0,
"cpd00018_C":1.0,
"cpd00065_C":-1.0
},
"lower_bound":0.0,
"upper_bound":1000.0,
"gene_reaction_rule":"ACIAD2871"
},
{
"id":"2_7_7_1_RXN",
"name":"nicotinamide-nucleotide adenylyltransferase",
"metabolites":{
"cpd00002_C":-1.0,
"cpd00003_C":1.0,
"cpd00012_C":1.0,
"cpd00355_C":-1.0
},
"lower_bound":-1000.0,
"upper_bound":1000.0,
"gene_reaction_rule":"ACIAD2606"
},
{
"id":"IMPCYCLOHYDROLASE_RXN",
"name":"imp cyclohydrolase",
"metabolites":{
"cpd00001_C":-1.0,
"cpd00114_C":-1.0,
"cpd02884_C":1.0
},
"lower_bound":-1000.0,
"upper_bound":1000.0,
"gene_reaction_rule":"ACIAD2447"
},
{
"id":"C14_0_2ENOYLCOA_HYDRATASE_RXN",
"name":"enoyl-coa hydratase (c14:0)",
"metabolites":{
"C14_0_2ENOYLCOA_C":-1.0,
"cpd00001_C":-1.0,
"cpd03115_C":1.0
},
"lower_bound":0.0,
"upper_bound":1000.0,
"gene_reaction_rule":"ACIAD1692 or ACIAD1726 or ACIAD1820 or ACIAD1829 or ACIAD2351 or ACIAD3073 or ACIAD1609 or ACIAD0335 or ACIAD1608 or ACIAD1701 or ACIAD1565 or ACIAD2416"
},
{
"id":"DIHYDROOROTOX_RXN",
"name":"dihydroorotate oxidase",
"metabolites":{
"cpd00007_C":-1.0,
"cpd00025_C":1.0,
"cpd00247_C":1.0,
"cpd00282_C":-1.0
},
"lower_bound":0.0,
"upper_bound":1000.0,
"gene_reaction_rule":"ACIAD1321"
},
{
"id":"EXF_GLUTARATE_E",
"name":"exchange flux of glutarate",
"metabolites":{
"cpd00379_E":-1.0
},
"lower_bound":-0.0,
"upper_bound":1000.0,
"gene_reaction_rule":""
},
{
"id":"C18_1_C9_3HYDROXYACP_DEHYDRATASE_RXN",
"name":"3hydroxyacyl-acp dehydratase (c18:1(c9))",
"metabolites":{
"C18_1_C9__2ENOYLACP_C":1.0,
"C18_1_C9__3HYDROXYACP_C":-1.0,
"cpd00001_C":1.0
},
"lower_bound":0.0,
"upper_bound":1000.0,
"gene_reaction_rule":"ACIAD1381"
},
{
"id":"EXF_CPDIM_6114_E",
"name":"exchange flux of salicylacetate",
"metabolites":{
"cpd01006_E":-1.0
},
"lower_bound":-0.0,
"upper_bound":1000.0,
"gene_reaction_rule":""
},
{
"id":"NACGLCTRANS_RXN",
"name":"udp-n-acetylglucosamine:n-acetylmuramyl-(pentapeptide) pyrophosphoryl-undecaprenol n-acetylglucosamine transferase",
"metabolites":{
"C6_C":1.0,
"cpd00014_C":1.0,
"cpd00037_C":-1.0,
"cpd03494_C":-1.0
},
"lower_bound":0.0,
"upper_bound":1000.0,
"gene_reaction_rule":"ACIAD3517"
},
{
"id":"UMPKI_RXN",
"name":"ump kinase",
"metabolites":{
"cpd00002_C":-1.0,
"cpd00008_C":1.0,
"cpd00014_C":1.0,
"cpd00091_C":-1.0
},
"lower_bound":-1000.0,
"upper_bound":1000.0,
"gene_reaction_rule":"ACIAD1372"
},
{
"id":"CYT_UBIQUINOL_OXID_D_RXN",
"name":"cytochrome oxidase bd",
"metabolites":{
"cpd00001_C":2.0,
"cpd00007_C":-1.0,
"cpd00067_C":-4.0,
"cpd00067_P":4.0,
"cpd15560_C":2.0,
"cpd15561_C":-2.0
},
"lower_bound":0.0,
"upper_bound":1000.0,
"gene_reaction_rule":"ACIAD2290 and ACIAD2291"
},
{
"id":"TRANS_RXN_SUBERATE",
"name":"transport of suberate [extraorganism;cytosol](secondary symport)",
"metabolites":{
"cpd00067_C":1.0,
"cpd00067_P":-1.0,
"cpd05193_C":1.0,
"cpd05193_E":-1.0
},
"lower_bound":-1000.0,
"upper_bound":1000.0,
"gene_reaction_rule":"ACIAD1694"
},
{
"id":"EXF_TRP_E",
"name":"exchange flux of l-tryptophan",
"metabolites":{
"cpd00065_E":-1.0
},
"lower_bound":-0.0,
"upper_bound":1000.0,
"gene_reaction_rule":""
},
{
"id":"ADENYL_KIN_RXN",
"name":"adenylate kinase",
"metabolites":{
"cpd00002_C":-1.0,
"cpd00008_C":2.0,
"cpd00018_C":-1.0
},
"lower_bound":-1000.0,
"upper_bound":1000.0,
"gene_reaction_rule":"ACIAD1105"
},
{
"id":"TRANS_RXN_FUM",
"name":"transport of fumarate [extraorganism;cytosol](secondary symport)",
"metabolites":{
"cpd00067_C":1.0,
"cpd00067_P":-1.0,
"cpd00106_C":1.0,
"cpd00106_E":-1.0
},
"lower_bound":0.0,
"upper_bound":1000.0,
"gene_reaction_rule":"ACIAD1026 or ACIAD2227 or ACIAD1191 or ACIAD1866"
},
{
"id":"RXN0_363",
"name":"xanthosine hydrolase",
"metabolites":{
"cpd00001_C":-1.0,
"cpd00105_C":1.0,
"cpd00309_C":1.0,
"cpd01217_C":-1.0
},
"lower_bound":0.0,
"upper_bound":1000.0,
"gene_reaction_rule":"ACIAD2386"
},
{
"id":"AMPSYN_RXN",
"name":"adenylosuccinate lyase",
"metabolites":{
"cpd00018_C":1.0,
"cpd00106_C":1.0,
"cpd02375_C":-1.0
},
"lower_bound":-1000.0,
"upper_bound":1000.0,
"gene_reaction_rule":"ACIAD1219"
},
{
"id":"PYRIMSYN3_RXN",
"name":"phosphomethylpyrimidine kinase",
"metabolites":{
"cpd00002_C":-1.0,
"cpd00008_C":1.0,
"cpd02775_C":-1.0,
"cpd02894_C":1.0
},
"lower_bound":-1000.0,
"upper_bound":1000.0,
"gene_reaction_rule":"ACIAD0875"
},
{
"id":"CO2_DISS_RXN",
"name":"dissolution of carbon dioxide",
"metabolites":{
"cpd00001_C":-1.0,
"cpd00011_C":-1.0,
"cpd00067_C":1.0,
"cpd00242_C":1.0
},
"lower_bound":-1000.0,
"upper_bound":1000.0,
"gene_reaction_rule":"SPONT"
},
{
"id":"GLUTSEMIALDEHYDROG_RXN",
"name":"glutamate-5-semialdehyde dehydrogenase",
"metabolites":{
"cpd00005_C":-1.0,
"cpd00006_C":1.0,
"cpd00009_C":1.0,
"cpd00858_C":1.0,
"cpd02097_C":-1.0
},
"lower_bound":0.0,
"upper_bound":1000.0,
"gene_reaction_rule":"ACIAD0547"
},
{
"id":"IGPSYN_RXN",
"name":"indole-3-glycerol-phosphate synthase",
"metabolites":{
"cpd00001_C":1.0,
"cpd00011_C":1.0,
"cpd00956_C":-1.0,
"cpd02210_C":1.0
},
"lower_bound":0.0,
"upper_bound":1000.0,
"gene_reaction_rule":"ACIAD2463"
},
{
"id":"QUINOLINATE_SYNTHA_RXN",
"name":"quinolinate synthetase a",
"metabolites":{
"cpd00001_C":2.0,
"cpd00009_C":1.0,
"cpd00095_C":-1.0,
"cpd02333_C":1.0,
"cpd03470_C":-1.0
},
"lower_bound":0.0,
"upper_bound":1000.0,
"gene_reaction_rule":"ACIAD0651"
},
{
"id":"EXF_L_ASPARTATE_E",
"name":"exchange flux of l-aspartate",
"metabolites":{
"cpd00041_E":-1.0
},
"lower_bound":-0.0,
"upper_bound":1000.0,
"gene_reaction_rule":""
},
{
"id":"ACONITATEDEHYDR_RXN",
"name":"aconitase (citrate hydro-lyase)",
"metabolites":{
"cpd00001_C":1.0,
"cpd00137_C":-1.0,
"cpd00331_C":1.0
},
"lower_bound":-1000.0,
"upper_bound":1000.0,
"gene_reaction_rule":"ACIAD2395 or ACIAD3090 or ACIAD2755"
},
{
"id":"TRANS_RXN_3_4_DIHYDROXYBENZOATE_PERIP",
"name":"3,4-dihydroxybenzoate periplasm-cytoplasm transport",
"metabolites":{
"cpd00197_C":1.0,
"cpd00197_P":-1.0
},
"lower_bound":-1000.0,
"upper_bound":1000.0,
"gene_reaction_rule":"ACIAD1709 or ACIAD0982"
},
{
"id":"C14_1_C5_2ENOYLCOA_HYDRATASE_RXN",
"name":"enoyl-coa hydratase (c14:1(c5))",
"metabolites":{
"cpd00001_C":-1.0,
"cpd16613_C":1.0,
"cpd35448_C":-1.0
},
"lower_bound":0.0,
"upper_bound":1000.0,
"gene_reaction_rule":"ACIAD1692 or ACIAD1726 or ACIAD1820 or ACIAD1829 or ACIAD2351 or ACIAD3073 or ACIAD1609 or ACIAD0335 or ACIAD1608 or ACIAD1701 or ACIAD1565 or ACIAD2416"
},
{
"id":"C16_0_COA_REDUCTASE_RXN",
"name":"fatty acyl-coa reductase (c16:0)",
"metabolites":{
"C16_0_ALDEHYDE_C":1.0,
"cpd00005_C":-1.0,
"cpd00006_C":1.0,
"cpd00010_C":1.0,
"cpd00134_C":-1.0
},
"lower_bound":-1000.0,
"upper_bound":1000.0,
"gene_reaction_rule":"ACIAD3383"
},
{
"id":"EXF_CPD_148_E",
"name":"exchange flux of octane",
"metabolites":{
"cpd00998_E":-1.0
},
"lower_bound":-0.0,
"upper_bound":1000.0,
"gene_reaction_rule":""
},
{
"id":"GDP_MANNOSE_6_DEHYDROGENASE_RXN",
"name":"gdp-mannose 6-dehydrogenase",
"metabolites":{
"cpd00001_C":-1.0,
"cpd00003_C":-2.0,
"cpd00004_C":2.0,
"cpd00083_C":-1.0,
"cpd00719_C":1.0
},
"lower_bound":-1000.0,
"upper_bound":1000.0,
"gene_reaction_rule":"ACIAD0087"
},
{
"id":"R222_RXN",
"name":"octanal dehydrogenase",
"metabolites":{
"cpd00001_C":-1.0,
"cpd00003_C":-1.0,
"cpd00004_C":1.0,
"cpd01088_C":-1.0,
"cpd03846_C":1.0
},
"lower_bound":-1000.0,
"upper_bound":1000.0,
"gene_reaction_rule":"ACIAD2018 or ACIAD2542"
},
{
"id":"DTMPKI_RXN",
"name":"dtmp kinase",
"metabolites":{
"cpd00002_C":-1.0,
"cpd00008_C":1.0,
"cpd00297_C":1.0,
"cpd00298_C":-1.0
},
"lower_bound":-1000.0,
"upper_bound":1000.0,
"gene_reaction_rule":"ACIAD2588"
},
{
"id":"PHOSGLYPHOS_RXN",
"name":"phosphoglycerate kinase",
"metabolites":{
"cpd00002_C":1.0,
"cpd00008_C":-1.0,
"cpd00169_C":1.0,
"cpd00203_C":-1.0
},
"lower_bound":-1000.0,
"upper_bound":1000.0,
"gene_reaction_rule":"ACIAD1927"
},
{
"id":"GLUTATHIONE_SYN_RXN",
"name":"glutathione synthase",
"metabolites":{
"cpd00002_C":-1.0,
"cpd00008_C":1.0,
"cpd00009_C":1.0,
"cpd00033_C":-1.0,
"cpd00042_C":1.0,
"cpd00506_C":-1.0
},
"lower_bound":0.0,
"upper_bound":1000.0,
"gene_reaction_rule":"ACIAD3518"
},
{
"id":"RXN0_280",
"name":"ethane sulfonate monooxygenase",
"metabolites":{
"cpd00001_C":1.0,
"cpd00007_C":-1.0,
"cpd00050_C":1.0,
"cpd00071_C":1.0,
"cpd00081_C":1.0,
"cpd01270_C":-1.0,
"cpd11579_C":-1.0
},
"lower_bound":0.0,
"upper_bound":1000.0,
"gene_reaction_rule":"ACIAD0036 or ACIAD1505 or ACIAD1518 or ACIAD1527 or ACIAD1535 or ACIAD3471"
},
{
"id":"EXF_ADENOSINE_E",
"name":"exchange flux of adenosine",
"metabolites":{
"cpd00182_E":-1.0
},
"lower_bound":-0.0,
"upper_bound":1000.0,
"gene_reaction_rule":""
},
{
"id":"ACETYL_COA_CARBOXYLTRANSFER_RXN",
"name":"acetyl-coa carboxylase",
"metabolites":{
"cpd00002_C":-1.0,
"cpd00008_C":1.0,
"cpd00009_C":1.0,
"cpd00022_C":-1.0,
"cpd00067_C":-1.0,
"cpd00070_C":1.0,
"cpd00242_C":-1.0
},
"lower_bound":-1000.0,
"upper_bound":1000.0,
"gene_reaction_rule":"(ACIAD3035 and ACIAD0643) and ACIAD0841 and ACIAD1736 and ACIAD1737"
},
{
"id":"L_LACTDEHYDROGFMN_RXN",
"name":"lactate oxidation",
"metabolites":{
"cpd00020_C":1.0,
"cpd00159_C":-1.0,
"cpd15560_C":-1.0,
"cpd15561_C":1.0
},
"lower_bound":0.0,
"upper_bound":1000.0,
"gene_reaction_rule":"ACIAD0108"
},
{
"id":"HISTCYCLOHYD_RXN",
"name":"phosphoribosyl-amp cyclohydrolase",
"metabolites":{
"cpd00001_C":-1.0,
"cpd01777_C":-1.0,
"cpd02979_C":1.0
},
"lower_bound":0.0,
"upper_bound":1000.0,
"gene_reaction_rule":"ACIAD0380"
},
{
"id":"C7_0_CARBOXY3OXOCOA_THIOLASE_RXN",
"name":"dicarboxylate 3oxoacyl-coa thiolase (c7:0)",
"metabolites":{
"cpd00010_C":-1.0,
"cpd00022_C":1.0,
"cpd00413_C":1.0,
"cpd04104_C":-1.0
},
"lower_bound":-1000.0,
"upper_bound":1000.0,
"gene_reaction_rule":"ACIAD1689"
},
{
"id":"EXF_C10_0_E",
"name":"exchange flux of decanoate",
"metabolites":{
"cpd01107_E":-1.0
},
"lower_bound":-0.0,
"upper_bound":1000.0,
"gene_reaction_rule":""
},
{
"id":"TRANS_RXN_CAFFEATE",
"name":"transport of caffeate [extraorganism;periplasm](passive)",
"metabolites":{
"cpd00881_E":-1.0,
"cpd00881_P":1.0
},
"lower_bound":-1000.0,
"upper_bound":1000.0,
"gene_reaction_rule":""
},
{
"id":"P_PANTOCYSDECARB_RXN",
"name":"phosphopantothenoylcysteine decarboxylase",
"metabolites":{
"cpd00011_C":1.0,
"cpd00834_C":1.0,
"cpd02666_C":-1.0
},
"lower_bound":0.0,
"upper_bound":1000.0,
"gene_reaction_rule":"ACIAD3125"
},
{
"id":"C16_1_C7_3OXOACP_REDUCTASE_RXN",
"name":"3oxoacyl-acp reductase (c16:1(c7))",
"metabolites":{
"C16_1_C7__3HYDROXYACP_C":1.0,
"C16_1_C7__3OXOACP_C":-1.0,
"cpd00005_C":-1.0,
"cpd00006_C":1.0
},
"lower_bound":0.0,
"upper_bound":1000.0,
"gene_reaction_rule":"ACIAD0871"
},
{
"id":"PREPHENATEDEHYDRAT_RXN",
"name":"prephenate dehydratase",
"metabolites":{
"cpd00001_C":1.0,
"cpd00011_C":1.0,
"cpd00143_C":1.0,
"cpd00219_C":-1.0
},
"lower_bound":0.0,
"upper_bound":1000.0,
"gene_reaction_rule":"ACIAD2223"
},
{
"id":"TRANS_RXNIM_5794",
"name":"transport of 2-oxoglutarate [extraorganism;cytosol](passive)",
"metabolites":{
"cpd00024_C":1.0,
"cpd00024_E":-1.0
},
"lower_bound":-1000.0,
"upper_bound":1000.0,
"gene_reaction_rule":"ACIAD1651"
},
{
"id":"TRANS_RXNIM_5792",
"name":"transport of cis,cis-muconate [extraorganism;cytosol](secondary symport)",
"metabolites":{
"cpd00067_C":1.0,
"cpd00067_P":-1.0,
"cpd01630_C":1.0,
"cpd01630_E":-1.0
},
"lower_bound":-1000.0,
"upper_bound":1000.0,
"gene_reaction_rule":"ACIAD1681 or ACIAD1694"
},
{
"id":"TRANS_RXN_WATER",
"name":"transport of h2o [extraorganism;cytosol](passive)",
"metabolites":{
"cpd00001_C":1.0,
"cpd00001_E":-1.0
},
"lower_bound":-1000.0,
"upper_bound":1000.0,
"gene_reaction_rule":""
},
{
"id":"EXF_INOSINE_E",
"name":"exchange flux of inosine",
"metabolites":{
"cpd00246_E":-1.0
},
"lower_bound":-0.0,
"upper_bound":1000.0,
"gene_reaction_rule":""
},
{
"id":"DAHPSYN_RXN",
"name":"2-dehydro-3-deoxyphosphoheptonate aldolase",
"metabolites":{
"cpd00001_C":-1.0,
"cpd00009_C":1.0,
"cpd00061_C":-1.0,
"cpd00236_C":-1.0,
"cpd02857_C":1.0
},
"lower_bound":0.0,
"upper_bound":1000.0,
"gene_reaction_rule":"ACIAD2330 or ACIAD1878"
},
{
"id":"C8_0_COA_DEHYDROGENASE_RXN",
"name":"acyl-coa dehydrogenase (c8:0)",
"metabolites":{
"cpd00015_C":-1.0,
"cpd00982_C":1.0,
"cpd01335_C":-1.0,
"cpd03130_C":1.0
},
"lower_bound":0.0,
"upper_bound":1000.0,
"gene_reaction_rule":"ACIAD0452 or ACIAD3262 or ACIAD2537 or ACIAD2277 or ACIAD1886 or ACIAD1821 or ACIAD1819 or ACIAD1700 or ACIAD1693 or ACIAD1607 or ACIAD1413 or ACIAD0623 or ACIAD0624 or ACIAD0428 or ACIAD0427 or ACIAD1723 or ACIAD2226 or ACIAD3191 or ACIAD1566"
},
{
"id":"TRANS_RXNIM_5923",
"name":"transport of tricarballylate [extraorganism;cytosol](secondary symport)",
"metabolites":{
"cpd00067_C":1.0,
"cpd00067_P":-1.0,
"cpd16654_C":1.0,
"cpd16654_E":-1.0
},
"lower_bound":-1000.0,
"upper_bound":1000.0,
"gene_reaction_rule":"ACIAD1541"
},
{
"id":"CPD_1301_ACTIVATION_RXN",
"name":"5-methyltetrahydropteroyltri-l-glutamate synthesis",
"metabolites":{
"cpd00001_C":1.0,
"cpd00003_C":1.0,
"cpd00004_C":-1.0,
"cpd00033_C":1.0,
"cpd00054_C":-1.0,
"cpd02555_C":-1.0,
"cpd02738_C":1.0
},
"lower_bound":-1000.0,
"upper_bound":1000.0,
"gene_reaction_rule":"ACIAD2255 and ACIAD2283"
},
{
"id":"C8_0_2ENOYLACP_REDUCTASE_RXN",
"name":"2enoyl-acp reductase (c8:0)",
"metabolites":{
"C8_0_2ENOYLACP_C":-1.0,
"C8_0_ACP_C":1.0,
"cpd00003_C":1.0,
"cpd00004_C":-1.0
},
"lower_bound":0.0,
"upper_bound":1000.0,
"gene_reaction_rule":"ACIAD3116"
},
{
"id":"C18_1_C9_COA_DEHYDROGENASE_RXN",
"name":"acyl-coa dehydrogenase (c18:1(c9))",
"metabolites":{
"cpd00015_C":-1.0,
"cpd00982_C":1.0,
"cpd16638_C":-1.0,
"cpd31963_C":1.0
},
"lower_bound":0.0,
"upper_bound":1000.0,
"gene_reaction_rule":"ACIAD0452 or ACIAD3262 or ACIAD2537 or ACIAD2277 or ACIAD1886 or ACIAD1821 or ACIAD1819 or ACIAD1700 or ACIAD1693 or ACIAD1607 or ACIAD1413 or ACIAD0623 or ACIAD0624 or ACIAD0428 or ACIAD0427 or ACIAD1723 or ACIAD2226 or ACIAD3191 or ACIAD1566"
},
{
"id":"XANPRIBOSYLTRAN_RXN",
"name":"xanthine-guanine phosphoribosyltransferase",
"metabolites":{
"cpd00012_C":1.0,
"cpd00103_C":-1.0,
"cpd00309_C":-1.0,
"cpd00497_C":1.0
},
"lower_bound":0.0,
"upper_bound":1000.0,
"gene_reaction_rule":"ACIAD3164"
},
{
"id":"2_ISOPROPYLMALATESYN_RXN",
"name":"2-isopropylmalate synthase",
"metabolites":{
"cpd00001_C":-1.0,
"cpd00010_C":1.0,
"cpd00022_C":-1.0,
"cpd00123_C":-1.0,
"cpd01646_C":1.0
},
"lower_bound":-1000.0,
"upper_bound":1000.0,
"gene_reaction_rule":"ACIAD0530"
},
{
"id":"ISOCIT_CLEAV_RXN",
"name":"isocitrate lyase",
"metabolites":{
"cpd00036_C":1.0,
"cpd00040_C":1.0,
"cpd00260_C":-1.0
},
"lower_bound":0.0,
"upper_bound":1000.0,
"gene_reaction_rule":"ACIAD1084"
},
{
"id":"MALATE_DEH_RXN",
"name":"malate dehydrogenase",
"metabolites":{
"cpd00003_C":-1.0,
"cpd00004_C":1.0,
"cpd00032_C":1.0,
"cpd00130_C":-1.0
},
"lower_bound":-1000.0,
"upper_bound":1000.0,
"gene_reaction_rule":"ACIAD3155"
},
{
"id":"BENZOATE_CIS_DIOL_DEHYDROGENASE_RXN",
"name":"cis-1,2-dihydroxycyclohexa-3,5-diene-1-carboxylate dehydrogenase",
"metabolites":{
"cpd00003_C":-1.0,
"cpd00004_C":1.0,
"cpd00011_C":1.0,
"cpd00077_C":1.0,
"cpd02821_C":-1.0
},
"lower_bound":0.0,
"upper_bound":1000.0,
"gene_reaction_rule":"ACIAD1439"
},
{
"id":"C14_0_COA_DEHYDROGENASE_RXN",
"name":"acyl-coa dehydrogenase (c14:0)",
"metabolites":{
"C14_0_2ENOYLCOA_C":1.0,
"cpd00015_C":-1.0,
"cpd00982_C":1.0,
"cpd01695_C":-1.0
},
"lower_bound":0.0,
"upper_bound":1000.0,
"gene_reaction_rule":"ACIAD0452 or ACIAD3262 or ACIAD2537 or ACIAD2277 or ACIAD1886 or ACIAD1821 or ACIAD1819 or ACIAD1700 or ACIAD1693 or ACIAD1607 or ACIAD1413 or ACIAD0623 or ACIAD0624 or ACIAD0428 or ACIAD0427 or ACIAD1723 or ACIAD2226 or ACIAD3191 or ACIAD1566"
},
{
"id":"ALCOHOL_DEHYDROG_RXN",
"name":"ethanol dehydroganase",
"metabolites":{
"cpd00003_C":1.0,
"cpd00004_C":-1.0,
"cpd00071_C":-1.0,
"cpd00363_C":1.0
},
"lower_bound":-1000.0,
"upper_bound":1000.0,
"gene_reaction_rule":"ACIAD3616 or ACIAD2929 or ACIAD2015 or ACIAD3339 or ACIAD1950"
},
{
"id":"TRANS_RXN_COUMARATE",
"name":"transport of 4-coumarate [extraorganism;cytosol](secondary symport)",
"metabolites":{
"cpd00067_C":1.0,
"cpd00067_P":-1.0,
"cpd00604_C":1.0,
"cpd00604_E":-1.0
},
"lower_bound":-1000.0,
"upper_bound":1000.0,
"gene_reaction_rule":"ACIAD1709"
},
{
"id":"ABC_13_RXN",
"name":"transport of l-glutamate [extraorganism;cytosol](primary active atp)",
"metabolites":{
"cpd00001_C":-1.0,
"cpd00002_C":-1.0,
"cpd00008_C":1.0,
"cpd00009_C":1.0,
"cpd00023_C":1.0,
"cpd00023_E":-1.0
},
"lower_bound":0.0,
"upper_bound":1000.0,
"gene_reaction_rule":"ACIAD2056 and ACIAD2058 and ACIAD2059 and ACIAD2061"
},
{
"id":"ALANINE_TRNA_LIGASE_RXN",
"name":"alanine--trna ligase",
"metabolites":{
"ALA_TRNAS_C":-1.0,
"CHARGED_ALA_TRNAS_C":1.0,
"cpd00002_C":-1.0,
"cpd00012_C":1.0,
"cpd00018_C":1.0,
"cpd00035_C":-1.0
},
"lower_bound":0.0,
"upper_bound":1000.0,
"gene_reaction_rule":"ACIAD1253"
},
{
"id":"C18_1_C9_ACP_HYDROLASE_RXN",
"name":"acyl-acp hydrolase (c18:1(c9))",
"metabolites":{
"ACP_C":1.0,
"C18_1_C9__ACP_C":-1.0,
"cpd00001_C":-1.0,
"cpd00536_C":1.0
},
"lower_bound":0.0,
"upper_bound":1000.0,
"gene_reaction_rule":""
},
{
"id":"TRANS_RXN_THYMIDINE",
"name":"transport of thymidine [extraorganism;cytosol](secondary symport)",
"metabolites":{
"cpd00067_C":1.0,
"cpd00067_P":-1.0,
"cpd00184_C":1.0,
"cpd00184_E":-1.0
},
"lower_bound":0.0,
"upper_bound":1000.0,
"gene_reaction_rule":"ACIAD2389"
},
{
"id":"TRANS_RXNIM_5876_BIS",
"name":"transport of d-glucarate [extraorganism;cytosol](passive)",
"metabolites":{
"cpd00571_C":1.0,
"cpd00571_E":-1.0
},
"lower_bound":-1000.0,
"upper_bound":1000.0,
"gene_reaction_rule":"ACIAD0127"
},
{
"id":"HISTALDEHYD_RXN",
"name":"histidinol dehydrogenase",
"metabolites":{
"cpd00001_C":-1.0,
"cpd00003_C":-1.0,
"cpd00004_C":1.0,
"cpd00119_C":1.0,
"cpd01324_C":-1.0
},
"lower_bound":0.0,
"upper_bound":1000.0,
"gene_reaction_rule":"ACIAD0663"
},
{
"id":"3_ISOPROPYLMALDEHYDROG_RXN",
"name":"3-isopropylmalate dehydrogenase",
"metabolites":{
"cpd00003_C":-1.0,
"cpd00004_C":1.0,
"cpd00011_C":1.0,
"cpd00200_C":1.0,
"cpd02693_C":-1.0
},
"lower_bound":0.0,
"upper_bound":1000.0,
"gene_reaction_rule":"ACIAD0469"
},
{
"id":"C18_0_2ENOYLCOA_HYDRATASE_RXN",
"name":"enoyl-coa hydratase (c18:0)",
"metabolites":{
"cpd00001_C":-1.0,
"cpd14936_C":1.0,
"cpd14937_C":-1.0
},
"lower_bound":0.0,
"upper_bound":1000.0,
"gene_reaction_rule":"ACIAD1692 or ACIAD1726 or ACIAD1820 or ACIAD1829 or ACIAD2351 or ACIAD3073 or ACIAD1609 or ACIAD0335 or ACIAD1608 or ACIAD1701 or ACIAD1565 or ACIAD2416"
},
{
"id":"C18_0_3OXOCOA_THIOLASE_RXN",
"name":"3oxoacyl-coa thiolase (c18:0)",
"metabolites":{
"cpd00010_C":-1.0,
"cpd00022_C":1.0,
"cpd00134_C":1.0,
"cpd14935_C":-1.0
},
"lower_bound":-1000.0,
"upper_bound":1000.0,
"gene_reaction_rule":"ACIAD0334 or ACIAD1450 or ACIAD1706 or ACIAD1977 or ACIAD1689 or ACIAD2988 or ACIAD1569"
},
{
"id":"EXF_PI_E",
"name":"exchange flux of phosphate",
"metabolites":{
"cpd00009_E":-1.0
},
"lower_bound":-1000.0,
"upper_bound":1000.0,
"gene_reaction_rule":""
},
{
"id":"PRTRANS_RXN",
"name":"anthranilate phosphoribosyltransferase",
"metabolites":{
"cpd00012_C":1.0,
"cpd00093_C":-1.0,
"cpd00103_C":-1.0,
"cpd02642_C":1.0
},
"lower_bound":-1000.0,
"upper_bound":1000.0,
"gene_reaction_rule":"ACIAD2462"
},
{
"id":"C16_0_3OXOACP_REDUCTASE_RXN",
"name":"3oxoacyl-acp reductase (c16:0)",
"metabolites":{
"C16_0_3HYDROXYACP_C":1.0,
"C16_0_3OXOACP_C":-1.0,
"cpd00005_C":-1.0,
"cpd00006_C":1.0
},
"lower_bound":0.0,
"upper_bound":1000.0,
"gene_reaction_rule":"ACIAD0871"
},
{
"id":"EXF_GLYCOLLATE_E",
"name":"exchange flux of glycolate",
"metabolites":{
"cpd00139_E":-1.0
},
"lower_bound":-0.0,
"upper_bound":1000.0,
"gene_reaction_rule":""
},
{
"id":"DHBDEHYD_RXN",
"name":"2,3-dihydro-2,3-dihydroxybenzoate dehydrogenase",
"metabolites":{
"cpd00003_C":-1.0,
"cpd00004_C":1.0,
"cpd00168_C":1.0,
"cpd02566_C":-1.0
},
"lower_bound":0.0,
"upper_bound":1000.0,
"gene_reaction_rule":"ACIAD2774"
},
{
"id":"AFATTYACID_WAX_ASSEMBLY_RXN",
"name":"generic fatty acid of wax assembly",
"metabolites":{
"AFATTYACID_WAX_C":1.0,
"cpd00214_C":-0.24,
"cpd00536_C":-0.445,
"cpd34461_C":-0.315
},
"lower_bound":-1000.0,
"upper_bound":1000.0,
"gene_reaction_rule":""
},
{
"id":"TRANS_RXNIM_5736",
"name":"transport of cytosine [extraorganism;cytosol](secondary symport)",
"metabolites":{
"cpd00067_C":1.0,
"cpd00067_P":-1.0,
"cpd00307_C":1.0,
"cpd00307_E":-1.0
},
"lower_bound":0.0,
"upper_bound":1000.0,
"gene_reaction_rule":"ACIAD2541"
},
{
"id":"2_OCTAPRENYL_6_METHOXYPHENOL_HYDROX_RXN",
"name":"2-octaprenyl-6-methoxyphenol hydroxylase",
"metabolites":{
"cpd00001_C":1.0,
"cpd00007_C":-1.0,
"cpd03446_C":-1.0,
"cpd03447_C":1.0
},
"lower_bound":0.0,
"upper_bound":1000.0,
"gene_reaction_rule":"ACIAD1127"
},
{
"id":"EXF_DEOXYADENOSINE_E",
"name":"exchange flux of deoxyadenosine",
"metabolites":{
"cpd00438_E":-1.0
},
"lower_bound":-0.0,
"upper_bound":1000.0,
"gene_reaction_rule":""
},
{
"id":"C5_0_3OXOCOA_THIOLASE_RXN",
"name":"3oxoacyl-coa thiolase (c5:0)",
"metabolites":{
"cpd00010_C":-1.0,
"cpd00022_C":1.0,
"cpd00086_C":1.0,
"cpd23921_C":-1.0
},
"lower_bound":-1000.0,
"upper_bound":1000.0,
"gene_reaction_rule":"ACIAD0334 or ACIAD1450 or ACIAD1706 or ACIAD1977 or ACIAD1689 or ACIAD2988 or ACIAD1569"
},
{
"id":"PROPKIN_RXN",
"name":"propionate kinase",
"metabolites":{
"cpd00002_C":-1.0,
"cpd00008_C":1.0,
"cpd00141_C":-1.0,
"cpd01844_C":1.0
},
"lower_bound":-1000.0,
"upper_bound":1000.0,
"gene_reaction_rule":"ACIAD0541"
},
{
"id":"ALDHDEHYDROG_RXN",
"name":"aminoacetaldehyde dehydrogenase",
"metabolites":{
"cpd00001_C":-1.0,
"cpd00003_C":-1.0,
"cpd00004_C":1.0,
"cpd00033_C":1.0,
"cpd04122_C":-1.0
},
"lower_bound":-1000.0,
"upper_bound":1000.0,
"gene_reaction_rule":"ACIAD2018 or ACIAD2542"
},
{
"id":"C12_0_COA_DEHYDROGENASE_RXN",
"name":"acyl-coa dehydrogenase (c12:0)",
"metabolites":{
"cpd00015_C":-1.0,
"cpd00982_C":1.0,
"cpd01260_C":-1.0,
"cpd02060_C":1.0
},
"lower_bound":0.0,
"upper_bound":1000.0,
"gene_reaction_rule":"ACIAD0452 or ACIAD3262 or ACIAD2537 or ACIAD2277 or ACIAD1886 or ACIAD1821 or ACIAD1819 or ACIAD1700 or ACIAD1693 or ACIAD1607 or ACIAD1413 or ACIAD0623 or ACIAD0624 or ACIAD0428 or ACIAD0427 or ACIAD1723 or ACIAD2226 or ACIAD3191 or ACIAD1566"
},
{
"id":"TRANS_RXN_122",
"name":"transport of l-glutamate [extraorganism;cytosol](secondary symport)",
"metabolites":{
"cpd00023_C":1.0,
"cpd00023_E":-1.0,
"cpd00971_C":1.0,
"cpd00971_E":-1.0
},
"lower_bound":0.0,
"upper_bound":1000.0,
"gene_reaction_rule":"ACIAD0804"
},
{
"id":"DUMP_NUCLEOTIDASE_RXN",
"name":"dump nucleotidase",
"metabolites":{
"cpd00001_C":-1.0,
"cpd00009_C":1.0,
"cpd00299_C":-1.0,
"cpd00412_C":1.0
},
"lower_bound":0.0,
"upper_bound":1000.0,
"gene_reaction_rule":"ACIAD0015 or ACIAD1227"
},
{
"id":"MUCONATE_CYCLOISOMERASE_RXN",
"name":"muconate cycloisomerase",
"metabolites":{
"cpd01630_C":1.0,
"cpd02536_C":-1.0
},
"lower_bound":-1000.0,
"upper_bound":1000.0,
"gene_reaction_rule":"ACIAD1446"
},
{
"id":"C8_0_3HYDROXYACP_DEHYDRATASE_RXN",
"name":"3hydroxyacyl-acp dehydratase (c8:0)",
"metabolites":{
"C8_0_2ENOYLACP_C":1.0,
"C8_0_3HYDROXYACP_C":-1.0,
"cpd00001_C":1.0
},
"lower_bound":0.0,
"upper_bound":1000.0,
"gene_reaction_rule":"ACIAD1381"
},
{
"id":"SULFITE_REDUCT_RXN",
"name":"sulfite reductase (nadph)",
"metabolites":{
"cpd00001_C":3.0,
"cpd00005_C":-3.0,
"cpd00006_C":3.0,
"cpd00081_C":-1.0,
"cpd00239_C":1.0
},
"lower_bound":-1000.0,
"upper_bound":1000.0,
"gene_reaction_rule":"ACIAD2982 and ACIAD2981"
},
{
"id":"NICOTINAMID_RXN",
"name":"nicotinamidase",
"metabolites":{
"cpd00001_C":-1.0,
"cpd00013_C":1.0,
"cpd00133_C":-1.0,
"cpd00218_C":1.0
},
"lower_bound":0.0,
"upper_bound":1000.0,
"gene_reaction_rule":"ACIAD3584 or ACIAD1618"
},
{
"id":"C10_0_CARBOXY2ENOYLCOA_HYDRATASE_RXN",
"name":"dicarboxylate enoyl-coa hydratase (c10:0)",
"metabolites":{
"cpd00001_C":-1.0,
"cpd16604_C":-1.0,
"cpd16605_C":1.0
},
"lower_bound":0.0,
"upper_bound":1000.0,
"gene_reaction_rule":"ACIAD1692"
},
{
"id":"GLYC3PDEHYDROG_RXN",
"name":"glycerol-3-phosphate dehydrogenase",
"metabolites":{
"cpd00015_C":-1.0,
"cpd00080_C":-1.0,
"cpd00095_C":1.0,
"cpd00982_C":1.0
},
"lower_bound":0.0,
"upper_bound":1000.0,
"gene_reaction_rule":"ACIAD2844"
},
{
"id":"TRANS_RXN_GLUTARATE",
"name":"transport of glutarate [extraorganism;cytosol](secondary symport)",
"metabolites":{
"cpd00067_C":1.0,
"cpd00067_P":-1.0,
"cpd00379_C":1.0,
"cpd00379_E":-1.0
},
"lower_bound":-1000.0,
"upper_bound":1000.0,
"gene_reaction_rule":"ACIAD1694"
},
{
"id":"C12_0_COATRANSFERASE_RXN",
"name":"acyl coa transferase (c12:0)",
"metabolites":{
"cpd00002_C":-1.0,
"cpd00010_C":-1.0,
"cpd00012_C":1.0,
"cpd00018_C":1.0,
"cpd01260_C":1.0,
"cpd01741_C":-1.0
},
"lower_bound":0.0,
"upper_bound":1000.0,
"gene_reaction_rule":"ACIAD1818 or ACIAD0235 or ACIAD1724 or ACIAD2106"
},
{
"id":"PSEUDOURIDYLATE_SYNTHASE_RXN",
"name":"pseudouridylate synthase",
"metabolites":{
"cpd00001_C":1.0,
"cpd00092_C":-1.0,
"cpd00101_C":-1.0,
"cpd00859_C":1.0
},
"lower_bound":0.0,
"upper_bound":1000.0,
"gene_reaction_rule":"ACIAD0474 or ACIAD0715 or ACIAD2897 or ACIAD3307 or ACIAD0861 or ACIAD1189 or ACIAD1203 or ACIAD2377 or ACIAD2380 or ACIAD0442"
},
{
"id":"EXF_LYS_E",
"name":"exchange flux of l-lysine",
"metabolites":{
"cpd00039_E":-1.0
},
"lower_bound":-0.0,
"upper_bound":1000.0,
"gene_reaction_rule":""
},
{
"id":"SHIKIMATE_5_DEHYDROGENASE_RXN",
"name":"shikimate 5-dehydrogenase",
"metabolites":{
"cpd00005_C":-1.0,
"cpd00006_C":1.0,
"cpd00383_C":1.0,
"cpd01716_C":-1.0
},
"lower_bound":-1000.0,
"upper_bound":1000.0,
"gene_reaction_rule":"ACIAD0418"
},
{
"id":"C18_1_C9_3HYDROXYCOA_DEHYDROGENASE_RXN",
"name":"hydroxyacyl-coa dehydrogenase (c18:1(c9))",
"metabolites":{
"cpd00003_C":-1.0,
"cpd00004_C":1.0,
"cpd32253_C":1.0,
"cpd34075_C":-1.0
},
"lower_bound":0.0,
"upper_bound":1000.0,
"gene_reaction_rule":"ACIAD0335 or ACIAD1565 or ACIAD2416"
},
{
"id":"PHOSPHOGLUCMUT_RXN",
"name":"phosphoglucomutase",
"metabolites":{
"cpd00079_C":1.0,
"cpd00089_C":-1.0
},
"lower_bound":-1000.0,
"upper_bound":1000.0,
"gene_reaction_rule":"ACIAD0104"
},
{
"id":"C6_1_C4_3HYDROXYCOA_DEHYDROGENASE_RXN",
"name":"hydroxyacyl-coa dehydrogenase (c6:1(c4))",
"metabolites":{
"cpd00003_C":-1.0,
"cpd00004_C":1.0,
"cpd16646_C":-1.0,
"cpd16647_C":1.0
},
"lower_bound":0.0,
"upper_bound":1000.0,
"gene_reaction_rule":"ACIAD0335 or ACIAD1565 or ACIAD2416"
},
{
"id":"C16_1_C7_3OXOACP_SYNTHASE_RXN",
"name":"3oxoacyl-acp synthase (c16:1(c7))",
"metabolites":{
"ACP_C":1.0,
"C14_1_C5__ACP_C":-1.0,
"C16_1_C7__3OXOACP_C":1.0,
"MALONYL_ACP_C":-1.0,
"cpd00011_C":1.0
},
"lower_bound":0.0,
"upper_bound":1000.0,
"gene_reaction_rule":"ACIAD0580 or ACIAD0879 or ACIAD0583 or ACIAD2101"
},
{
"id":"ACETATEKIN_RXN",
"name":"acetate kinase",
"metabolites":{
"cpd00002_C":-1.0,
"cpd00008_C":1.0,
"cpd00029_C":-1.0,
"cpd00196_C":1.0
},
"lower_bound":-1000.0,
"upper_bound":1000.0,
"gene_reaction_rule":"ACIAD0541"
},
{
"id":"DTDPKIN_RXN",
"name":"dtdp kinase",
"metabolites":{
"cpd00002_C":-1.0,
"cpd00008_C":1.0,
"cpd00297_C":-1.0,
"cpd00357_C":1.0
},
"lower_bound":-1000.0,
"upper_bound":1000.0,
"gene_reaction_rule":"ACIAD0556"
},
{
"id":"RXNIM_6173",
"name":"4-hydroxyphenyl-beta-hydroxypropionyl-coa lyase",
"metabolites":{
"cpd00022_C":1.0,
"cpd00484_C":1.0,
"cpd23287_C":-1.0
},
"lower_bound":-1000.0,
"upper_bound":1000.0,
"gene_reaction_rule":"ACIAD1726"
},
{
"id":"EXF_GLUCONATE_E",
"name":"exchange flux of gluconate",
"metabolites":{
"cpd00222_E":-1.0
},
"lower_bound":-0.0,
"upper_bound":1000.0,
"gene_reaction_rule":""
},
{
"id":"PHOSACETYLTRANS_RXN",
"name":"phosphate acetyltransferase",
"metabolites":{
"cpd00009_C":1.0,
"cpd00010_C":-1.0,
"cpd00022_C":1.0,
"cpd00196_C":-1.0
},
"lower_bound":-1000.0,
"upper_bound":1000.0,
"gene_reaction_rule":"ACIAD2287 or ACIAD0540"
},
{
"id":"ASPARTATEKIN_RXN",
"name":"aspartate kinase",
"metabolites":{
"cpd00002_C":-1.0,
"cpd00008_C":1.0,
"cpd00041_C":-1.0,
"cpd01977_C":1.0
},
"lower_bound":0.0,
"upper_bound":1000.0,
"gene_reaction_rule":"ACIAD1252"
},
{
"id":"C14_0_COA_THIOLASE_RXN",
"name":"acyl-coa thiolase (c14:0)",
"metabolites":{
"cpd00001_C":-1.0,
"cpd00010_C":1.0,
"cpd01695_C":-1.0,
"cpd03847_C":1.0
},
"lower_bound":0.0,
"upper_bound":1000.0,
"gene_reaction_rule":"ACIAD0341 or ACIAD3231 or ACIAD1023"
},
{
"id":"TRANS_RXN_BUTANEDIOL",
"name":"transport of 2,3-butanediol [extraorganism;cytosol](passive)",
"metabolites":{
"cpd01947_C":1.0,
"cpd01947_E":-1.0
},
"lower_bound":0.0,
"upper_bound":1000.0,
"gene_reaction_rule":""
},
{
"id":"TRANS_RXNIM_5803",
"name":"transport of benzoate [periplasm;cytosol](secondary symport) irreversible",
"metabolites":{
"cpd00067_C":1.0,
"cpd00067_P":-1.0,
"cpd00153_C":1.0,
"cpd00153_P":-1.0
},
"lower_bound":0.0,
"upper_bound":1000.0,
"gene_reaction_rule":"ACIAD1440 or ACIAD1982"
},
{
"id":"RXN0_743",
"name":"phosphoribosylaminoimidazole carboxylase, mutase subunit",
"metabolites":{
"cpd02893_C":1.0,
"cpd11310_C":-1.0
},
"lower_bound":0.0,
"upper_bound":1000.0,
"gene_reaction_rule":"ACIAD3659"
},
{
"id":"FUMARYLACETOACETASE_RXN",
"name":"fumarylacetoacetase",
"metabolites":{
"cpd00001_C":-1.0,
"cpd00106_C":1.0,
"cpd00142_C":1.0,
"cpd00763_C":-1.0
},
"lower_bound":-1000.0,
"upper_bound":1000.0,
"gene_reaction_rule":"ACIAD1809"
},
{
"id":"C16_0_3HYDROXYCOA_DEHYDROGENASE_RXN",
"name":"hydroxyacyl-coa dehydrogenase (c16:0)",
"metabolites":{
"cpd00003_C":-1.0,
"cpd00004_C":1.0,
"cpd03113_C":-1.0,
"cpd03114_C":1.0
},
"lower_bound":0.0,
"upper_bound":1000.0,
"gene_reaction_rule":"ACIAD0335 or ACIAD1565 or ACIAD2416"
},
{
"id":"H2NEOPTERINP3PYROPHOSPHOHYDRO_RXN",
"name":"dihydroneopterin triphosphate pyrophosphatase",
"metabolites":{
"cpd00001_C":-1.0,
"cpd00012_C":1.0,
"cpd02978_C":-1.0,
"cpd03521_C":1.0
},
"lower_bound":0.0,
"upper_bound":1000.0,
"gene_reaction_rule":""
},
{
"id":"1_ACYLGLYCEROL_3_P_ACYLTRANSFER_PL_RXN",
"name":"glycerol-3-phosphate acyltransferase",
"metabolites":{
"ACP_C":1.0,
"ACYL_ACP_PL_C":-1.0,
"ACYL_SN_GLYCEROL_3P_PL_C":-1.0,
"cpd11422_C":1.0
},
"lower_bound":0.0,
"upper_bound":1000.0,
"gene_reaction_rule":"ACIAD3232"
},
{
"id":"DSERDEAM_RXN",
"name":"d-serine dehydratase",
"metabolites":{
"cpd00013_C":1.0,
"cpd00020_C":1.0,
"cpd00550_C":-1.0
},
"lower_bound":0.0,
"upper_bound":1000.0,
"gene_reaction_rule":"ACIAD1048"
},
{
"id":"EXF_L_ALPHA_ALANINE_E",
"name":"exchange flux of l-alanine",
"metabolites":{
"cpd00035_E":-1.0
},
"lower_bound":-0.0,
"upper_bound":1000.0,
"gene_reaction_rule":""
},
{
"id":"THIAZOLSYN3_RXN",
"name":"hydroxyethylthiazole kinase",
"metabolites":{
"cpd00002_C":-1.0,
"cpd00008_C":1.0,
"cpd02636_C":-1.0,
"cpd02654_C":1.0
},
"lower_bound":-1000.0,
"upper_bound":1000.0,
"gene_reaction_rule":"ACIAD2011"
},
{
"id":"C6_0_3HYDROXYCOA_DEHYDROGENASE_RXN",
"name":"hydroxyacyl-coa dehydrogenase (c6:0)",
"metabolites":{
"cpd00003_C":-1.0,
"cpd00004_C":1.0,
"cpd03122_C":-1.0,
"cpd03123_C":1.0
},
"lower_bound":0.0,
"upper_bound":1000.0,
"gene_reaction_rule":"ACIAD0335 or ACIAD1565 or ACIAD2416"
},
{
"id":"EXF_DEOXYGUANOSINE_E",
"name":"exchange flux of deoxyguanosine",
"metabolites":{
"cpd00277_E":-1.0
},
"lower_bound":-0.0,
"upper_bound":1000.0,
"gene_reaction_rule":""
},
{
"id":"PREPHENATE_TRANSAMINE_RXN",
"name":"l-arogenate aminotransferase",
"metabolites":{
"cpd00023_C":-1.0,
"cpd00024_C":1.0,
"cpd00219_C":-1.0,
"cpd00616_C":1.0
},
"lower_bound":-1000.0,
"upper_bound":1000.0,
"gene_reaction_rule":"ACIAD0112"
},
{
"id":"C6_0_COA_DEHYDROGENASE_RXN",
"name":"acyl-coa dehydrogenase (c6:0)",
"metabolites":{
"cpd00015_C":-1.0,
"cpd00982_C":1.0,
"cpd03124_C":-1.0,
"cpd03125_C":1.0
},
"lower_bound":0.0,
"upper_bound":1000.0,
"gene_reaction_rule":"ACIAD0452 or ACIAD3262 or ACIAD2537 or ACIAD2277 or ACIAD1886 or ACIAD1821 or ACIAD1819 or ACIAD1700 or ACIAD1693 or ACIAD1607 or ACIAD1413 or ACIAD0623 or ACIAD0624 or ACIAD0428 or ACIAD0427 or ACIAD1723 or ACIAD2226 or ACIAD3191 or ACIAD1566"
},
{
"id":"TRANS_RXN_GLUCONATE",
"name":"transport of gluconate [extraorganism;cytosol](passive)",
"metabolites":{
"cpd00222_C":1.0,
"cpd00222_E":-1.0
},
"lower_bound":0.0,
"upper_bound":1000.0,
"gene_reaction_rule":""
},
{
"id":"HOMOCYSMETB12_RXN",
"name":"5-methyltetrahydrofolate--homocysteine s-methyltransferase",
"metabolites":{
"cpd00060_C":1.0,
"cpd00087_C":1.0,
"cpd00135_C":-1.0,
"cpd00345_C":-1.0
},
"lower_bound":0.0,
"upper_bound":1000.0,
"gene_reaction_rule":"ACIAD1045"
},
{
"id":"C14_1_C5_2ENOYLACP_REDUCTASE_RXN",
"name":"2enoyl-acp reductase (c14:1(c5))",
"metabolites":{
"C14_1_C5__2ENOYLACP_C":-1.0,
"C14_1_C5__ACP_C":1.0,
"cpd00003_C":1.0,
"cpd00004_C":-1.0
},
"lower_bound":0.0,
"upper_bound":1000.0,
"gene_reaction_rule":"ACIAD3116"
},
{
"id":"TRANS_RXNIM_5847",
"name":"transport of l-alanine [extraorganism;cytosol](secondary symport)",
"metabolites":{
"cpd00035_C":1.0,
"cpd00035_E":-1.0,
"cpd00971_C":1.0,
"cpd00971_E":-1.0
},
"lower_bound":0.0,
"upper_bound":1000.0,
"gene_reaction_rule":"ACIAD0805"
},
{
"id":"2PGADEHYDRAT_RXN",
"name":"phosphopyruvate hydratase",
"metabolites":{
"cpd00001_C":1.0,
"cpd00061_C":1.0,
"cpd00482_C":-1.0
},
"lower_bound":-1000.0,
"upper_bound":1000.0,
"gene_reaction_rule":"ACIAD2001"
},
{
"id":"RXNIM_6188",
"name":"protocatechualdehyde dehydrogenase",
"metabolites":{
"cpd00001_C":-1.0,
"cpd00003_C":-1.0,
"cpd00004_C":1.0,
"cpd00197_C":1.0,
"cpd16507_C":-1.0
},
"lower_bound":-1000.0,
"upper_bound":1000.0,
"gene_reaction_rule":"ACIAD1725"
},
{
"id":"EXF_CHLOROGENATE_E",
"name":"exchange flux of chlorogenate",
"metabolites":{
"cpd00634_E":-1.0
},
"lower_bound":-0.0,
"upper_bound":1000.0,
"gene_reaction_rule":""
},
{
"id":"RXNIM_6204",
"name":"tricarballylate dehydrogenase",
"metabolites":{
"cpd00331_C":1.0,
"cpd15560_C":-1.0,
"cpd15561_C":1.0,
"cpd16654_C":-1.0
},
"lower_bound":0.0,
"upper_bound":1000.0,
"gene_reaction_rule":"ACIAD1537 and ACIAD1536"
},
{
"id":"TRANS_RXN_201",
"name":"transport of citrate [extraorganism;cytosol](secondary symport)",
"metabolites":{
"cpd00067_C":1.0,
"cpd00067_P":-1.0,
"cpd00137_C":1.0,
"cpd00137_E":-1.0
},
"lower_bound":0.0,
"upper_bound":1000.0,
"gene_reaction_rule":"ACIAD1074 or ACIAD1931 or ACIAD2818"
},
{
"id":"P_PANTOCYSLIG_RXN",
"name":"phosphopantothenate--cysteine ligase",
"metabolites":{
"cpd00012_C":1.0,
"cpd00046_C":1.0,
"cpd00052_C":-1.0,
"cpd00084_C":-1.0,
"cpd02201_C":-1.0,
"cpd02666_C":1.0
},
"lower_bound":-1000.0,
"upper_bound":1000.0,
"gene_reaction_rule":"ACIAD3125"
},
{
"id":"TRANS_RXNIM_5901",
"name":"transport of fructose [extraorganism;cytosol](secondary antiport)",
"metabolites":{
"cpd00020_C":1.0,
"cpd00061_C":-1.0,
"cpd00212_E":-1.0,
"cpd01850_C":1.0
},
"lower_bound":0.0,
"upper_bound":1000.0,
"gene_reaction_rule":"ACIAD1990 and ACIAD1993 and ACIAD3058"
},
{
"id":"TRANS_RXN_DEOXYGUANOSINE",
"name":"transport of deoxyguanosine [extraorganism;cytosol](secondary symport)",
"metabolites":{
"cpd00067_C":1.0,
"cpd00067_P":-1.0,
"cpd00277_C":1.0,
"cpd00277_E":-1.0
},
"lower_bound":0.0,
"upper_bound":1000.0,
"gene_reaction_rule":"ACIAD2389"
},
{
"id":"ABC_70_RXN",
"name":"transport of so42- [extraorganism;cytosol](primary active atp)",
"metabolites":{
"cpd00001_C":-1.0,
"cpd00002_C":-1.0,
"cpd00008_C":1.0,
"cpd00009_C":1.0,
"cpd00048_C":1.0,
"cpd00048_E":-1.0
},
"lower_bound":0.0,
"upper_bound":1000.0,
"gene_reaction_rule":"(ACIAD3369 or ACIAD2591) and ACIAD2594 and ACIAD2595 and ACIAD2596"
},
{
"id":"EXF_NICOTINAMIDE_NUCLEOTIDE_E",
"name":"exchange flux of nicotinamide nucleotide",
"metabolites":{
"cpd00355_E":-1.0
},
"lower_bound":-0.0,
"upper_bound":1000.0,
"gene_reaction_rule":""
},
{
"id":"GAPOXNPHOSPHN_RXN",
"name":"glyceraldehyde 3-phosphate dehydrogenase (phosphorylating)",
"metabolites":{
"cpd00003_C":-1.0,
"cpd00004_C":1.0,
"cpd00009_C":-1.0,
"cpd00102_C":-1.0,
"cpd00203_C":1.0
},
"lower_bound":-1000.0,
"upper_bound":1000.0,
"gene_reaction_rule":"ACIAD2565"
},
{
"id":"R223_RXN",
"name":"acyl coa transferase (c8:0)",
"metabolites":{
"cpd00002_C":-1.0,
"cpd00010_C":-1.0,
"cpd00012_C":1.0,
"cpd00018_C":1.0,
"cpd01335_C":1.0,
"cpd03846_C":-1.0
},
"lower_bound":0.0,
"upper_bound":1000.0,
"gene_reaction_rule":"ACIAD1818 or ACIAD0235 or ACIAD1724 or ACIAD2106"
},
{
"id":"ADENYLYLSULFATE_REDUCTASE_RXN",
"name":"adenylylsulfate reductase",
"metabolites":{
"APS_C":-1.0,
"cpd00018_C":1.0,
"cpd00081_C":1.0,
"cpd11420_C":1.0,
"cpd11421_C":-1.0
},
"lower_bound":0.0,
"upper_bound":1000.0,
"gene_reaction_rule":"ACIAD0833"
},
{
"id":"TRANS_RXN_SO3_OUT",
"name":"transport of so3-2 [cytosol;extraorganism](passive)",
"metabolites":{
"cpd00081_C":-1.0,
"cpd00081_E":1.0
},
"lower_bound":0.0,
"upper_bound":1000.0,
"gene_reaction_rule":""
},
{
"id":"3_CH3_2_OXOBUTANOATE_OH_CH3_XFER_RXN",
"name":"3-methyl-2-oxobutanoate hydroxymethyltransferase",
"metabolites":{
"cpd00001_C":-1.0,
"cpd00087_C":1.0,
"cpd00123_C":-1.0,
"cpd00125_C":-1.0,
"cpd00712_C":1.0
},
"lower_bound":0.0,
"upper_bound":1000.0,
"gene_reaction_rule":"ACIAD3061"
},
{
"id":"UPPSYN_RXN",
"name":"undecaprenyl pyrophosphate synthetase",
"metabolites":{
"cpd00012_C":8.0,
"cpd00113_C":-8.0,
"cpd00350_C":-1.0,
"cpd02229_C":1.0
},
"lower_bound":0.0,
"upper_bound":1000.0,
"gene_reaction_rule":"ACIAD1374"
},
{
"id":"GLUTATHIONE_REDUCT_NADPH_RXN",
"name":"glutathione reductase (nadph)",
"metabolites":{
"cpd00005_C":-1.0,
"cpd00006_C":1.0,
"cpd00042_C":2.0,
"cpd00067_C":-1.0,
"cpd00111_C":-1.0
},
"lower_bound":-1000.0,
"upper_bound":1000.0,
"gene_reaction_rule":""
},
{
"id":"2_4_2_12_RXN",
"name":"nicotinamide phosphoribosyltransferase",
"metabolites":{
"cpd00012_C":-1.0,
"cpd00103_C":1.0,
"cpd00133_C":1.0,
"cpd00355_C":-1.0
},
"lower_bound":-1000.0,
"upper_bound":1000.0,
"gene_reaction_rule":"ACIAD0963"
},
{
"id":"GLYCERALDEHYDE_DEHYDRO_RXN",
"name":"aldehyde dehydrogenase (nad+)",
"metabolites":{
"cpd00001_C":-1.0,
"cpd00003_C":-1.0,
"cpd00004_C":1.0,
"cpd00223_C":1.0,
"cpd00448_C":-1.0
},
"lower_bound":-1000.0,
"upper_bound":1000.0,
"gene_reaction_rule":"ACIAD2018 or ACIAD2542"
},
{
"id":"1_2_1_27_RXN",
"name":"methylmalonate-semialdehyde dehydrogenase (acylating)",
"metabolites":{
"cpd00003_C":-1.0,
"cpd00004_C":1.0,
"cpd00010_C":-1.0,
"cpd00011_C":1.0,
"cpd00086_C":1.0,
"cpd00287_C":-1.0
},
"lower_bound":0.0,
"upper_bound":1000.0,
"gene_reaction_rule":"ACIAD1604"
},
{
"id":"TRANS_RXNIM_5812_BIS",
"name":"transport of l-ornithine [extraorganism;cytosol](secondary antiport)",
"metabolites":{
"cpd00064_C":1.0,
"cpd00064_E":-1.0,
"cpd00067_C":-1.0,
"cpd00067_P":1.0
},
"lower_bound":-1000.0,
"upper_bound":1000.0,
"gene_reaction_rule":"ACIAD1417"
},
{
"id":"ACETOOHBUTSYN_RXN",
"name":"acetolactate synthase",
"metabolites":{
"cpd00011_C":1.0,
"cpd00020_C":-1.0,
"cpd00094_C":-1.0,
"cpd00498_C":1.0
},
"lower_bound":0.0,
"upper_bound":1000.0,
"gene_reaction_rule":"(ACIAD0999 or ACIAD3104) and ACIAD3103"
},
{
"id":"GABATRANSAM_RXN",
"name":"4-aminobutyrate aminotransferase",
"metabolites":{
"cpd00023_C":1.0,
"cpd00024_C":-1.0,
"cpd00199_C":1.0,
"cpd00281_C":-1.0
},
"lower_bound":-1000.0,
"upper_bound":1000.0,
"gene_reaction_rule":"ACIAD3446"
},
{
"id":"C18_0_COA_THIOLASE_RXN",
"name":"acyl-coa thiolase (c18:0)",
"metabolites":{
"cpd00001_C":-1.0,
"cpd00010_C":1.0,
"cpd00327_C":-1.0,
"cpd01080_C":1.0
},
"lower_bound":0.0,
"upper_bound":1000.0,
"gene_reaction_rule":"ACIAD0341 or ACIAD3231 or ACIAD1023"
},
{
"id":"AICARSYN_RXN",
"name":"adenylosuccinate lyase",
"metabolites":{
"cpd00106_C":1.0,
"cpd02851_C":1.0,
"cpd02921_C":-1.0
},
"lower_bound":-1000.0,
"upper_bound":1000.0,
"gene_reaction_rule":"ACIAD1219"
},
{
"id":"MALATE_DEHYDROGENASE_ACCEPTOR_RXN",
"name":"malate dehydrogenase (acceptor)",
"metabolites":{
"cpd00032_C":1.0,
"cpd00130_C":-1.0,
"cpd15560_C":-1.0,
"cpd15561_C":1.0
},
"lower_bound":0.0,
"upper_bound":1000.0,
"gene_reaction_rule":"ACIAD1007"
},
{
"id":"C10_0_3HYDROXYCOA_DEHYDROGENASE_RXN",
"name":"hydroxyacyl-coa dehydrogenase (c10:0)",
"metabolites":{
"cpd00003_C":-1.0,
"cpd00004_C":1.0,
"cpd03118_C":-1.0,
"cpd03119_C":1.0
},
"lower_bound":0.0,
"upper_bound":1000.0,
"gene_reaction_rule":"ACIAD0335 or ACIAD1565 or ACIAD2416"
},
{
"id":"RXN0_742",
"name":"phosphoribosylaminoimidazole carboxylase, atpase subunit",
"metabolites":{
"cpd00001_C":-1.0,
"cpd00002_C":-1.0,
"cpd00008_C":1.0,
"cpd00009_C":1.0,
"cpd00011_C":-1.0,
"cpd02140_C":-1.0,
"cpd11310_C":1.0
},
"lower_bound":0.0,
"upper_bound":1000.0,
"gene_reaction_rule":"ACIAD3658"
},
{
"id":"OHMETHYLBILANESYN_RXN",
"name":"hydroxymethylbilane synthase",
"metabolites":{
"cpd00001_C":-1.0,
"cpd00013_C":4.0,
"cpd00689_C":-4.0,
"cpd00755_C":1.0
},
"lower_bound":0.0,
"upper_bound":1000.0,
"gene_reaction_rule":"ACIAD0286"
},
{
"id":"ACETALD_DEHYDROG_RXN",
"name":"acetaldehyde dehydrogenase (acetylating)",
"metabolites":{
"cpd00003_C":-1.0,
"cpd00004_C":1.0,
"cpd00010_C":-1.0,
"cpd00022_C":1.0,
"cpd00071_C":-1.0
},
"lower_bound":-1000.0,
"upper_bound":1000.0,
"gene_reaction_rule":"ACIAD2018"
},
{
"id":"FUMHYDR_RXN",
"name":"fumarate hydratase",
"metabolites":{
"cpd00001_C":-1.0,
"cpd00106_C":-1.0,
"cpd00130_C":1.0
},
"lower_bound":-1000.0,
"upper_bound":1000.0,
"gene_reaction_rule":"ACIAD1890 or ACIAD0538"
},
{
"id":"GPH_RXN",
"name":"phosphoglycolate phosphatase",
"metabolites":{
"cpd00001_C":-1.0,
"cpd00009_C":1.0,
"cpd00139_C":1.0,
"cpd00727_C":-1.0
},
"lower_bound":0.0,
"upper_bound":1000.0,
"gene_reaction_rule":"ACIAD0043 or ACIAD0296 or ACIAD0443"
},
{
"id":"XMPXAN_RXN",
"name":"xmp nucleotidase",
"metabolites":{
"cpd00001_C":-1.0,
"cpd00009_C":1.0,
"cpd00497_C":-1.0,
"cpd01217_C":1.0
},
"lower_bound":0.0,
"upper_bound":1000.0,
"gene_reaction_rule":"ACIAD0015 or ACIAD1227"
},
{
"id":"EXF_SUBERATE_E",
"name":"exchange flux of suberate",
"metabolites":{
"cpd05193_E":-1.0
},
"lower_bound":-0.0,
"upper_bound":1000.0,
"gene_reaction_rule":""
},
{
"id":"C16_0_2ENOYLACP_REDUCTASE_RXN",
"name":"2enoyl-acp reductase (c16:0)",
"metabolites":{
"C16_0_2ENOYLACP_C":-1.0,
"C16_0_ACP_C":1.0,
"cpd00003_C":1.0,
"cpd00004_C":-1.0
},
"lower_bound":0.0,
"upper_bound":1000.0,
"gene_reaction_rule":"ACIAD3116"
},
{
"id":"C14_1_C5_3HYDROXYACP_DEHYDRATASE_RXN",
"name":"3hydroxyacyl-acp dehydratase (c14:1(c5))",
"metabolites":{
"C14_1_C5__2ENOYLACP_C":1.0,
"C14_1_C5__3HYDROXYACP_C":-1.0,
"cpd00001_C":1.0
},
"lower_bound":0.0,
"upper_bound":1000.0,
"gene_reaction_rule":"ACIAD1381"
},
{
"id":"PHOSPHASERDECARB_RXN",
"name":"phosphatidylserine decarboxylase",
"metabolites":{
"L_1_PHOSPHATIDYL_ETHANOLAMINE_C":1.0,
"L_1_PHOSPHATIDYL_SERINE_C":-1.0,
"cpd00011_C":1.0
},
"lower_bound":0.0,
"upper_bound":1000.0,
"gene_reaction_rule":"ACIAD3560"
},
{
"id":"DIAMINOPIMDECARB_RXN",
"name":"diaminopimelate decarboxylase",
"metabolites":{
"cpd00011_C":1.0,
"cpd00039_C":1.0,
"cpd00516_C":-1.0
},
"lower_bound":0.0,
"upper_bound":1000.0,
"gene_reaction_rule":"ACIAD2660"
},
{
"id":"SERINE_TRNA_LIGASE_RXN",
"name":"serine--trna ligase",
"metabolites":{
"CHARGED_SER_TRNAS_C":1.0,
"SER_TRNAS_C":-1.0,
"cpd00002_C":-1.0,
"cpd00012_C":1.0,
"cpd00018_C":1.0,
"cpd00054_C":-1.0
},
"lower_bound":0.0,
"upper_bound":1000.0,
"gene_reaction_rule":"ACIAD2935"
},
{
"id":"TYROSINE_TRNA_LIGASE_RXN",
"name":"tyrosine--trna ligase",
"metabolites":{
"CHARGED_TYR_TRNAS_C":1.0,
"TYR_TRNAS_C":-1.0,
"cpd00002_C":-1.0,
"cpd00012_C":1.0,
"cpd00018_C":1.0,
"cpd00069_C":-1.0
},
"lower_bound":0.0,
"upper_bound":1000.0,
"gene_reaction_rule":"ACIAD0013"
},
{
"id":"ADCLY_RXN",
"name":"4-amino-4-deoxychorismate lyase",
"metabolites":{
"cpd00020_C":1.0,
"cpd00443_C":1.0,
"cpd08210_C":-1.0
},
"lower_bound":-1000.0,
"upper_bound":1000.0,
"gene_reaction_rule":"ACIAD2590"
},
{
"id":"RXN0_2044",
"name":"3-hydroxyadipyl-coa dehydrogenase",
"metabolites":{
"cpd00003_C":-1.0,
"cpd00004_C":1.0,
"cpd01507_C":1.0,
"cpd09844_C":-1.0
},
"lower_bound":0.0,
"upper_bound":1000.0,
"gene_reaction_rule":"ACIAD1690"
},
{
"id":"EXF_C14_0_E",
"name":"exchange flux of tetradecanoate",
"metabolites":{
"cpd03847_E":-1.0
},
"lower_bound":-0.0,
"upper_bound":1000.0,
"gene_reaction_rule":""
},
{
"id":"UDP_NACMURALA_GLU_LIG_RXN",
"name":"udp-n-acetylmuramoylalanine--d-glutamate ligase",
"metabolites":{
"cpd00002_C":-1.0,
"cpd00008_C":1.0,
"cpd00009_C":1.0,
"cpd00186_C":-1.0,
"cpd00525_C":1.0,
"cpd00890_C":-1.0
},
"lower_bound":0.0,
"upper_bound":1000.0,
"gene_reaction_rule":"ACIAD0270"
},
{
"id":"RIBOFLAVINKIN_RXN",
"name":"riboflavin kinase",
"metabolites":{
"cpd00002_C":-1.0,
"cpd00008_C":1.0,
"cpd00050_C":1.0,
"cpd00220_C":-1.0
},
"lower_bound":-1000.0,
"upper_bound":1000.0,
"gene_reaction_rule":"ACIAD0023"
},
{
"id":"GLUTAMIN_RXN",
"name":"glutaminase",
"metabolites":{
"cpd00001_C":-1.0,
"cpd00013_C":1.0,
"cpd00023_C":1.0,
"cpd00053_C":-1.0
},
"lower_bound":0.0,
"upper_bound":1000.0,
"gene_reaction_rule":"ACIAD1040 or ACIAD2088"
},
{
"id":"ADENOSINE_NUCLEOSIDASE_RXN",
"name":"adenosine nucleosidase",
"metabolites":{
"cpd00001_C":-1.0,
"cpd00105_C":1.0,
"cpd00128_C":1.0,
"cpd00182_C":-1.0
},
"lower_bound":0.0,
"upper_bound":1000.0,
"gene_reaction_rule":"ACIAD2386"
},
{
"id":"TRANS_RXN_FORMATE",
"name":"transport of formate [extraorganism;cytosol](secondary symport)",
"metabolites":{
"cpd00047_C":1.0,
"cpd00047_E":-1.0,
"cpd00067_C":1.0,
"cpd00067_P":-1.0
},
"lower_bound":-1000.0,
"upper_bound":1000.0,
"gene_reaction_rule":"ACIAD2100"
},
{
"id":"EXF_FRU_E",
"name":"exchange flux of fructose",
"metabolites":{
"cpd00212_E":-1.0
},
"lower_bound":-0.0,
"upper_bound":1000.0,
"gene_reaction_rule":""
},
{
"id":"TRANS_RXN_CPD_3623",
"name":"transport of sebacate [extraorganism;cytosol](secondary symport)",
"metabolites":{
"cpd00067_C":1.0,
"cpd00067_P":-1.0,
"cpd05192_C":1.0,
"cpd05192_E":-1.0
},
"lower_bound":-1000.0,
"upper_bound":1000.0,
"gene_reaction_rule":"ACIAD1694"
},
{
"id":"C14_1_C5_COATRANSFERASE_RXN",
"name":"acyl coa transferase (c14:1(c5))",
"metabolites":{
"cpd00002_C":-1.0,
"cpd00010_C":-1.0,
"cpd00012_C":1.0,
"cpd00018_C":1.0,
"cpd16609_C":-1.0,
"cpd16617_C":1.0
},
"lower_bound":0.0,
"upper_bound":1000.0,
"gene_reaction_rule":"ACIAD1818 or ACIAD0235 or ACIAD1724 or ACIAD2106"
},
{
"id":"ADENPRIBOSYLTRAN_RXN",
"name":"adenine phosphoribosyltransferase",
"metabolites":{
"cpd00012_C":1.0,
"cpd00018_C":1.0,
"cpd00103_C":-1.0,
"cpd00128_C":-1.0
},
"lower_bound":0.0,
"upper_bound":1000.0,
"gene_reaction_rule":"ACIAD3164"
},
{
"id":"PHOSNACMURPENTATRANS_RXN",
"name":"phospho-n-acetylmuramoyl-pentapeptide transferase",
"metabolites":{
"cpd00091_C":1.0,
"cpd00286_C":-1.0,
"cpd02932_C":-1.0,
"cpd03494_C":1.0
},
"lower_bound":0.0,
"upper_bound":1000.0,
"gene_reaction_rule":"ACIAD3363"
},
{
"id":"OROTPDECARB_RXN",
"name":"orotidine-5'-phosphate decarboxylase",
"metabolites":{
"cpd00011_C":1.0,
"cpd00091_C":1.0,
"cpd00810_C":-1.0
},
"lower_bound":0.0,
"upper_bound":1000.0,
"gene_reaction_rule":"ACIAD2341"
},
{
"id":"UNBAL_S_ADENOSYL_4_METHYLTHIO_2_OXOBUTANOATE",
"name":"unbalance flux of s-adenosyl-4-methylthio-2-oxobutanoate",
"metabolites":{
"cpd02701_C":-1.0
},
"lower_bound":0.0,
"upper_bound":1000.0,
"gene_reaction_rule":""
},
{
"id":"EXF_BUTYRIC_ACID_E",
"name":"exchange flux of butanoate",
"metabolites":{
"cpd00211_E":-1.0
},
"lower_bound":-0.0,
"upper_bound":1000.0,
"gene_reaction_rule":""
},
{
"id":"LEUCINE_TRNA_LIGASE_RXN",
"name":"leucine--trna ligase",
"metabolites":{
"CHARGED_LEU_TRNAS_C":1.0,
"LEU_TRNAS_C":-1.0,
"cpd00002_C":-1.0,
"cpd00012_C":1.0,
"cpd00018_C":1.0,
"cpd00107_C":-1.0
},
"lower_bound":0.0,
"upper_bound":1000.0,
"gene_reaction_rule":"ACIAD3106"
},
{
"id":"PHENYLALANINE_TRNA_LIGASE_RXN",
"name":"phenylalanine--trna ligase",
"metabolites":{
"CHARGED_PHE_TRNAS_C":1.0,
"PHE_TRNAS_C":-1.0,
"cpd00002_C":-1.0,
"cpd00012_C":1.0,
"cpd00018_C":1.0,
"cpd00066_C":-1.0
},
"lower_bound":0.0,
"upper_bound":1000.0,
"gene_reaction_rule":"ACIAD3041 and ACIAD3042"
},
{
"id":"C6_2_c2_c4_COATRANSFERASE_RXN",
"name":"acyl coa transferase (c6:2(c2,c4))",
"metabolites":{
"C6_1_C4__2ENOYLCOA_C":1.0,
"cpd00002_C":-1.0,
"cpd00010_C":-1.0,
"cpd00012_C":1.0,
"cpd00018_C":1.0,
"cpd16648_C":-1.0
},
"lower_bound":0.0,
"upper_bound":1000.0,
"gene_reaction_rule":"ACIAD1818 or ACIAD0235 or ACIAD1724 or ACIAD2106"
},
{
"id":"C18_1_C9_3OXOACP_REDUCTASE_RXN",
"name":"3oxoacyl-acp reductase (c18:1(c9))",
"metabolites":{
"C18_1_C9__3HYDROXYACP_C":1.0,
"C18_1_C9__3OXOACP_C":-1.0,
"cpd00005_C":-1.0,
"cpd00006_C":1.0
},
"lower_bound":0.0,
"upper_bound":1000.0,
"gene_reaction_rule":"ACIAD0871"
},
{
"id":"ASPARTASE_RXN",
"name":"aspartate ammonia-lyase",
"metabolites":{
"cpd00013_C":1.0,
"cpd00041_C":-1.0,
"cpd00106_C":1.0
},
"lower_bound":0.0,
"upper_bound":1000.0,
"gene_reaction_rule":"ACIAD1744"
},
{
"id":"TRANS_RXNIM_5805",
"name":"transport of benzoate [periplasm;cytosol](secondary symport) reversible",
"metabolites":{
"cpd00067_C":1.0,
"cpd00067_P":-1.0,
"cpd00153_C":1.0,
"cpd00153_P":-1.0
},
"lower_bound":-1000.0,
"upper_bound":1000.0,
"gene_reaction_rule":"ACIAD1434 or ACIAD1815"
},
{
"id":"TRANS_RXN_PI",
"name":"transport of phosphate [extraorganism;cytosol](secondary symport)",
"metabolites":{
"cpd00009_C":1.0,
"cpd00009_E":-1.0,
"cpd00067_C":1.0,
"cpd00067_P":-1.0
},
"lower_bound":0.0,
"upper_bound":1000.0,
"gene_reaction_rule":"ACIAD3641"
},
{
"id":"TRANS_RXNIM_5899",
"name":"transport of l-aspartate [extraorganism;cytosol](primary active atp)",
"metabolites":{
"cpd00001_C":-1.0,
"cpd00002_C":-1.0,
"cpd00008_C":1.0,
"cpd00009_C":1.0,
"cpd00041_C":1.0,
"cpd00041_E":-1.0
},
"lower_bound":0.0,
"upper_bound":1000.0,
"gene_reaction_rule":"ACIAD2056 and ACIAD2058 and ACIAD2059 and ACIAD2061"
},
{
"id":"EXF_GUANOSINE_E",
"name":"exchange flux of guanosine",
"metabolites":{
"cpd00311_E":-1.0
},
"lower_bound":-0.0,
"upper_bound":1000.0,
"gene_reaction_rule":""
},
{
"id":"SUCCSEMIALDDEHYDROG_RXN",
"name":"succinate-semialdehyde dehydrogenase (nad(p)+)",
"metabolites":{
"cpd00001_C":-1.0,
"cpd00005_C":1.0,
"cpd00006_C":-1.0,
"cpd00036_C":1.0,
"cpd00199_C":-1.0
},
"lower_bound":-1000.0,
"upper_bound":1000.0,
"gene_reaction_rule":"ACIAD0960 or ACIAD2539 or ACIAD3445"
},
{
"id":"NICOTINATEPRIBOSYLTRANS_RXN",
"name":"nicotinate phosphoribosyltransferase",
"metabolites":{
"NICOTINATE_NUCLEOTIDE_C":1.0,
"cpd00001_C":-1.0,
"cpd00002_C":-1.0,
"cpd00008_C":1.0,
"cpd00009_C":1.0,
"cpd00012_C":1.0,
"cpd00103_C":-1.0,
"cpd00218_C":-1.0
},
"lower_bound":0.0,
"upper_bound":1000.0,
"gene_reaction_rule":"ACIAD3562"
},
{
"id":"ISOLEUCINE_TRNA_LIGASE_RXN",
"name":"isoleucine--trna ligase",
"metabolites":{
"CHARGED_ILE_TRNAS_C":1.0,
"ILE_TRNAS_C":-1.0,
"cpd00002_C":-1.0,
"cpd00012_C":1.0,
"cpd00018_C":1.0,
"cpd00322_C":-1.0
},
"lower_bound":0.0,
"upper_bound":1000.0,
"gene_reaction_rule":"ACIAD0022"
},
{
"id":"LUMAZINESYN_RXN",
"name":"lumazine synthase",
"metabolites":{
"cpd00001_C":2.0,
"cpd00009_C":1.0,
"cpd02656_C":1.0,
"cpd02882_C":-1.0,
"cpd11225_C":-1.0
},
"lower_bound":0.0,
"upper_bound":1000.0,
"gene_reaction_rule":"ACIAD3571"
},
{
"id":"TRANS_RXN_CPD_148",
"name":"transport of octane [extraorganism;cytosol](passive)",
"metabolites":{
"cpd00998_C":1.0,
"cpd00998_E":-1.0
},
"lower_bound":0.0,
"upper_bound":1000.0,
"gene_reaction_rule":""
},
{
"id":"EXF_FE2_E",
"name":"exchange flux of fe+2",
"metabolites":{
"cpd00021_E":-1.0
},
"lower_bound":-1000.0,
"upper_bound":1000.0,
"gene_reaction_rule":""
},
{
"id":"TRANS_RXN_242A_BIS",
"name":"transport of homoserine lactone [cytosol;extraorganism](secondary antiport)",
"metabolites":{
"cpd00067_C":1.0,
"cpd00067_P":-1.0,
"cpd01873_C":-1.0,
"cpd01873_E":1.0
},
"lower_bound":0.0,
"upper_bound":1000.0,
"gene_reaction_rule":"ACIAD0173"
},
{
"id":"CYSTATHIONINE_BETA_LYASE_RXN",
"name":"cystathionine beta-lyase",
"metabolites":{
"cpd00001_C":-1.0,
"cpd00013_C":1.0,
"cpd00020_C":1.0,
"cpd00135_C":1.0,
"cpd00424_C":-1.0
},
"lower_bound":-1000.0,
"upper_bound":1000.0,
"gene_reaction_rule":"ACIAD3222"
},
{
"id":"RXN_2",
"name":"vanillate o-demethylase oxygenase",
"metabolites":{
"cpd00001_C":1.0,
"cpd00003_C":1.0,
"cpd00004_C":-1.0,
"cpd00007_C":-1.0,
"cpd00055_C":1.0,
"cpd00197_C":1.0,
"cpd04073_C":-1.0
},
"lower_bound":0.0,
"upper_bound":1000.0,
"gene_reaction_rule":"ACIAD0979 and ACIAD0980"
},
{
"id":"GLUTRACE_RXN",
"name":"glutamate racemase",
"metabolites":{
"cpd00023_C":-1.0,
"cpd00186_C":1.0
},
"lower_bound":-1000.0,
"upper_bound":1000.0,
"gene_reaction_rule":"ACIAD3257"
},
{
"id":"PHOSPHASERSYN_RXN",
"name":"phosphatidylserine synthase",
"metabolites":{
"CDPDIACYLGLYCEROL_C":-1.0,
"L_1_PHOSPHATIDYL_SERINE_C":1.0,
"cpd00046_C":1.0,
"cpd00054_C":-1.0
},
"lower_bound":0.0,
"upper_bound":1000.0,
"gene_reaction_rule":"ACIAD3149"
},
{
"id":"HOMOCYSMET_RXN",
"name":"5-methyltetrahydropteroyltriglutamate--homocysteine s-methyltransferase",
"metabolites":{
"cpd00060_C":1.0,
"cpd00135_C":-1.0,
"cpd02555_C":1.0,
"cpd02738_C":-1.0
},
"lower_bound":0.0,
"upper_bound":1000.0,
"gene_reaction_rule":"ACIAD3523"
},
{
"id":"WAX_ESTER_SYNTHASE_RXN",
"name":"wax ester synthase",
"metabolites":{
"ACYL_COA_WAX_C":-1.0,
"ACYL_OH_WAX_C":-1.0,
"WAX_ESTER_C":1.0,
"cpd00010_C":1.0
},
"lower_bound":0.0,
"upper_bound":1000.0,
"gene_reaction_rule":"ACIAD0832"
},
{
"id":"C10_0_CARBOXY3HYDROXYCOA_DEHYDROGENASE_RXN",
"name":"dicarboxylate hydroxyacyl-coa dehydrogenase (c10:0)",
"metabolites":{
"cpd00003_C":-1.0,
"cpd00004_C":1.0,
"cpd16605_C":-1.0,
"cpd16606_C":1.0
},
"lower_bound":0.0,
"upper_bound":1000.0,
"gene_reaction_rule":"ACIAD1690"
},
{
"id":"2_METHYLCITRATE_SYNTHASE_RXN",
"name":"2-methylcitrate synthase",
"metabolites":{
"cpd00001_C":-1.0,
"cpd00010_C":1.0,
"cpd00032_C":-1.0,
"cpd00086_C":-1.0,
"cpd01501_C":1.0
},
"lower_bound":0.0,
"upper_bound":1000.0,
"gene_reaction_rule":"ACIAD2756"
},
{
"id":"TRANS_RXN_162",
"name":"transport of l-glutamate [extraorganism;cytosol](secondary symport)",
"metabolites":{
"cpd00023_C":1.0,
"cpd00023_E":-1.0,
"cpd00067_C":1.0,
"cpd00067_P":-1.0
},
"lower_bound":0.0,
"upper_bound":1000.0,
"gene_reaction_rule":"ACIAD1339 or ACIAD0319 or ACIAD0481 or ACIAD3230"
},
{
"id":"PROLINE_TRNA_LIGASE_RXN",
"name":"proline--trna ligase",
"metabolites":{
"CHARGED_PRO_TRNAS_C":1.0,
"PRO_TRNAS_C":-1.0,
"cpd00002_C":-1.0,
"cpd00012_C":1.0,
"cpd00018_C":1.0,
"cpd00129_C":-1.0
},
"lower_bound":0.0,
"upper_bound":1000.0,
"gene_reaction_rule":"ACIAD0782"
},
{
"id":"TRANS_RXNIM_5747",
"name":"transport of betaine [extraorganism;cytosol](passive)",
"metabolites":{
"cpd00540_C":1.0,
"cpd00540_E":-1.0
},
"lower_bound":-1000.0,
"upper_bound":1000.0,
"gene_reaction_rule":"ACIAD2280 or ACIAD1036"
},
{
"id":"EXF_C16_1_C7_E",
"name":"exchange flux of cis-hexadec-7-enoate",
"metabolites":{
"cpd34461_E":-1.0
},
"lower_bound":-0.0,
"upper_bound":1000.0,
"gene_reaction_rule":""
},
{
"id":"TRANS_RXNIM_5753",
"name":"transport of d-methionine [extraorganism;cytosol](primary active atp)",
"metabolites":{
"cpd00001_C":-1.0,
"cpd00002_C":-1.0,
"cpd00008_C":1.0,
"cpd00009_C":1.0,
"cpd00637_C":1.0,
"cpd00637_E":-1.0
},
"lower_bound":0.0,
"upper_bound":1000.0,
"gene_reaction_rule":"(ACIAD2068 and ACIAD2069 and ACIAD2070) or (ACIAD2449 and ACIAD2450 and ACIAD2451)"
},
{
"id":"EXF_MAL_E",
"name":"exchange flux of malate",
"metabolites":{
"cpd00130_E":-1.0
},
"lower_bound":-0.0,
"upper_bound":1000.0,
"gene_reaction_rule":""
},
{
"id":"TRANS_RXN_WATER_PERIP",
"name":"transport of h2o [extraorganism;periplasm](passive)",
"metabolites":{
"cpd00001_E":-1.0,
"cpd00001_P":1.0
},
"lower_bound":-1000.0,
"upper_bound":1000.0,
"gene_reaction_rule":""
},
{
"id":"TRANS_RXN_4_HYDROXY_BENZYL_ALCOHOL",
"name":"transport of 4-hydroxybenzyl alcohol [extraorganism;cytosol](passive)",
"metabolites":{
"cpd15378_C":1.0,
"cpd15378_E":-1.0
},
"lower_bound":-1000.0,
"upper_bound":1000.0,
"gene_reaction_rule":""
},
{
"id":"SARCOX_RXN",
"name":"sarcosine oxidase",
"metabolites":{
"cpd00001_C":-1.0,
"cpd00007_C":-1.0,
"cpd00025_C":1.0,
"cpd00033_C":1.0,
"cpd00055_C":1.0,
"cpd00183_C":-1.0
},
"lower_bound":0.0,
"upper_bound":1000.0,
"gene_reaction_rule":"ACIAD2549 and ACIAD2550 and ACIAD2551 and ACIAD2552"
},
{
"id":"C18_1_C9_COATRANSFERASE_RXN",
"name":"acyl coa transferase (c18:1(c9))",
"metabolites":{
"cpd00002_C":-1.0,
"cpd00010_C":-1.0,
"cpd00012_C":1.0,
"cpd00018_C":1.0,
"cpd00536_C":-1.0,
"cpd16638_C":1.0
},
"lower_bound":0.0,
"upper_bound":1000.0,
"gene_reaction_rule":"ACIAD1818 or ACIAD0235 or ACIAD1724 or ACIAD2106"
},
{
"id":"DALADALALIG_RXN",
"name":"d-alanine--d-alanine ligase",
"metabolites":{
"cpd00002_C":-1.0,
"cpd00008_C":1.0,
"cpd00009_C":1.0,
"cpd00117_C":-2.0,
"cpd00731_C":1.0
},
"lower_bound":-1000.0,
"upper_bound":1000.0,
"gene_reaction_rule":"ACIAD3515"
},
{
"id":"DTDPDEHYDRHAMEPIM_RXN",
"name":"dtdp-4-dehydrorhamnose 3,5-epimerase",
"metabolites":{
"cpd00521_C":-1.0,
"cpd00522_C":1.0
},
"lower_bound":-1000.0,
"upper_bound":1000.0,
"gene_reaction_rule":"ACIAD0080"
},
{
"id":"2_NITROPROPANE_DIOXYGENASE_RXN",
"name":"2-nitropropane dioxygenase",
"metabolites":{
"cpd00007_C":-0.5,
"cpd00075_C":1.0,
"cpd00178_C":1.0,
"cpd01433_C":-1.0
},
"lower_bound":0.0,
"upper_bound":1000.0,
"gene_reaction_rule":"ACIAD1570"
},
{
"id":"SUCCGLUALDDEHYD_RXN",
"name":"succinylglutamic semialdehyde dehydrogenase",
"metabolites":{
"cpd00001_C":-1.0,
"cpd00003_C":-1.0,
"cpd00004_C":1.0,
"cpd00067_C":1.0,
"cpd03525_C":1.0,
"cpd03526_C":-1.0
},
"lower_bound":-1000.0,
"upper_bound":1000.0,
"gene_reaction_rule":"ACIAD1287"
},
{
"id":"EXF_CPD_3623_E",
"name":"exchange flux of sebacate",
"metabolites":{
"cpd05192_E":-1.0
},
"lower_bound":-0.0,
"upper_bound":1000.0,
"gene_reaction_rule":""
},
{
"id":"TRANS_RXNIM_5742",
"name":"transport of l-methionine [extraorganism;cytosol](primary active atp)",
"metabolites":{
"cpd00001_C":-1.0,
"cpd00002_C":-1.0,
"cpd00008_C":1.0,
"cpd00009_C":1.0,
"cpd00060_C":1.0,
"cpd00060_E":-1.0
},
"lower_bound":0.0,
"upper_bound":1000.0,
"gene_reaction_rule":"ACIAD2449 and ACIAD2450 and ACIAD2451"
},
{
"id":"C18_1_C9_3OXOACP_SYNTHASE_RXN",
"name":"3oxoacyl-acp synthase (c18:1(c9))",
"metabolites":{
"ACP_C":1.0,
"C16_1_C7__ACP_C":-1.0,
"C18_1_C9__3OXOACP_C":1.0,
"MALONYL_ACP_C":-1.0,
"cpd00011_C":1.0
},
"lower_bound":0.0,
"upper_bound":1000.0,
"gene_reaction_rule":"ACIAD0580 or ACIAD0879 or ACIAD0583 or ACIAD2101"
},
{
"id":"TRANS_RXN_TAURINE",
"name":"transport of taurine [extraorganism;cytosol](primary active atp)",
"metabolites":{
"cpd00001_C":-1.0,
"cpd00002_C":-1.0,
"cpd00008_C":1.0,
"cpd00009_C":1.0,
"cpd00210_C":1.0,
"cpd00210_E":-1.0
},
"lower_bound":0.0,
"upper_bound":1000.0,
"gene_reaction_rule":"(ACIAD1914 and ACIAD1463 and ACIAD1465 and ACIAD1464) or (ACIAD1523 and ACIAD1524 and ACIAD1525 and ACIAD1526) or (ACIAD1586 and ACIAD1587 and ACIAD1593 and ACIAD1601)"
},
{
"id":"EXF_CHOLINE_E",
"name":"exchange flux of choline",
"metabolites":{
"cpd00098_E":-1.0
},
"lower_bound":-0.0,
"upper_bound":1000.0,
"gene_reaction_rule":""
},
{
"id":"TRANS_RXN_L_ASPARTATE",
"name":"transport of l-aspartate [extraorganism;cytosol](secondary symport)",
"metabolites":{
"cpd00041_C":1.0,
"cpd00041_E":-1.0,
"cpd00067_C":1.0,
"cpd00067_P":-1.0
},
"lower_bound":0.0,
"upper_bound":1000.0,
"gene_reaction_rule":"ACIAD1026 or ACIAD2227 or ACIAD1191 or ACIAD1866"
},
{
"id":"PRAISOM_RXN",
"name":"phosphoribosylanthranilate isomerase",
"metabolites":{
"cpd00956_C":1.0,
"cpd02642_C":-1.0
},
"lower_bound":0.0,
"upper_bound":1000.0,
"gene_reaction_rule":"ACIAD0635"
},
{
"id":"C16_0_3OXOACP_SYNTHASE_RXN",
"name":"3oxoacyl-acp synthase (c16:0)",
"metabolites":{
"ACP_C":1.0,
"C14_0_ACP_C":-1.0,
"C16_0_3OXOACP_C":1.0,
"MALONYL_ACP_C":-1.0,
"cpd00011_C":1.0
},
"lower_bound":0.0,
"upper_bound":1000.0,
"gene_reaction_rule":"ACIAD0580 or ACIAD0879 or ACIAD0583 or ACIAD2101"
},
{
"id":"BHBDCLOS_RXN",
"name":"3-hydroxyacyl-coa dehydrogenase",
"metabolites":{
"cpd00003_C":-1.0,
"cpd00004_C":1.0,
"cpd00279_C":1.0,
"cpd00842_C":-1.0
},
"lower_bound":0.0,
"upper_bound":1000.0,
"gene_reaction_rule":"ACIAD0335 or ACIAD1565 or ACIAD2416"
},
{
"id":"GMP_SYN_GLUT_RXN",
"name":"gmp synthase (glutamine-hydrolysing)",
"metabolites":{
"cpd00001_C":-1.0,
"cpd00002_C":-1.0,
"cpd00012_C":1.0,
"cpd00018_C":1.0,
"cpd00023_C":1.0,
"cpd00053_C":-1.0,
"cpd00126_C":1.0,
"cpd00497_C":-1.0
},
"lower_bound":-1000.0,
"upper_bound":1000.0,
"gene_reaction_rule":"ACIAD0151"
},
{
"id":"C18_0_3HYDROXYACP_DEHYDRATASE_RXN",
"name":"3hydroxyacyl-acp dehydratase (c18:0)",
"metabolites":{
"C18_0_2ENOYLACP_C":1.0,
"C18_0_3HYDROXYACP_C":-1.0,
"cpd00001_C":1.0
},
"lower_bound":0.0,
"upper_bound":1000.0,
"gene_reaction_rule":"ACIAD1381"
},
{
"id":"BENZYL_ALC_DEHYDROGENASE_RXN",
"name":"benzyl alcohol dehydrogenase",
"metabolites":{
"cpd00003_C":-1.0,
"cpd00004_C":1.0,
"cpd00225_C":1.0,
"cpd00435_C":-1.0
},
"lower_bound":-1000.0,
"upper_bound":1000.0,
"gene_reaction_rule":"ACIAD1429 or ACIAD1578"
},
{
"id":"RXN0_1461",
"name":"coproporphyrinogen oxidase",
"metabolites":{
"cpd00007_C":-2.0,
"cpd00011_C":2.0,
"cpd00025_C":2.0,
"cpd00791_C":1.0,
"cpd02083_C":-1.0
},
"lower_bound":0.0,
"upper_bound":1000.0,
"gene_reaction_rule":"ACIAD3250"
},
{
"id":"TRANS_RXN_CPDIM_6115",
"name":"transport of 4-hydroxybenzylacetate [extraorganism;cytosol](passive)",
"metabolites":{
"cpd16658_C":1.0,
"cpd16658_E":-1.0
},
"lower_bound":-1000.0,
"upper_bound":1000.0,
"gene_reaction_rule":""
},
{
"id":"TRANS_RXN_CPDIM_6116",
"name":"transport of 3,4-dihydroxyphenylpropionate [extraorganism;cytosol](passive)",
"metabolites":{
"cpd07333_C":1.0,
"cpd07333_E":-1.0
},
"lower_bound":-1000.0,
"upper_bound":1000.0,
"gene_reaction_rule":""
},
{
"id":"EXF_C12_0_E",
"name":"exchange flux of dodecanoate",
"metabolites":{
"cpd01741_E":-1.0
},
"lower_bound":-0.0,
"upper_bound":1000.0,
"gene_reaction_rule":""
},
{
"id":"HISTAMINOTRANS_RXN",
"name":"histidinol-phosphate aminotransferase",
"metabolites":{
"cpd00023_C":-1.0,
"cpd00024_C":1.0,
"cpd00807_C":1.0,
"cpd00930_C":-1.0
},
"lower_bound":-1000.0,
"upper_bound":1000.0,
"gene_reaction_rule":"ACIAD0664"
},
{
"id":"TRANS_RXN_CPDIM_6112",
"name":"transport of ethylsalicylate [extraorganism;cytosol](passive)",
"metabolites":{
"cpd16657_C":1.0,
"cpd16657_E":-1.0
},
"lower_bound":-1000.0,
"upper_bound":1000.0,
"gene_reaction_rule":""
},
{
"id":"ASPARAGHYD_RXN",
"name":"asparaginase",
"metabolites":{
"cpd00001_C":-1.0,
"cpd00013_C":1.0,
"cpd00041_C":1.0,
"cpd00132_C":-1.0
},
"lower_bound":0.0,
"upper_bound":1000.0,
"gene_reaction_rule":"ACIAD2088"
},
{
"id":"FAD_UBIQUINONE_RXN",
"name":"fad oxidation",
"metabolites":{
"cpd00015_C":1.0,
"cpd00982_C":-1.0,
"cpd15560_C":-1.0,
"cpd15561_C":1.0
},
"lower_bound":0.0,
"upper_bound":1000.0,
"gene_reaction_rule":""
},
{
"id":"PQQ_OXID_RXN",
"name":"pyrroloquinoline-quinone oxidation",
"metabolites":{
"cpd00067_P":2.0,
"cpd00097_P":1.0,
"cpd00986_P":-1.0
},
"lower_bound":0.0,
"upper_bound":1000.0,
"gene_reaction_rule":""
},
{
"id":"COFACTORSYN_RXN",
"name":"cofactor assembly",
"metabolites":{
"COFACTOR_W_C":0.61,
"cpd00003_C":-0.07,
"cpd00006_C":-0.07,
"cpd00010_C":-0.08,
"cpd00015_C":-0.07,
"cpd00016_C":-0.08,
"cpd00017_C":-0.08,
"cpd00028_C":-0.08,
"cpd00050_C":-0.07,
"cpd00056_C":-0.08,
"cpd00087_C":-0.08,
"cpd00557_C":-0.08,
"cpd02229_C":-0.08,
"cpd15560_C":-0.08
},
"lower_bound":0.0,
"upper_bound":1000.0,
"gene_reaction_rule":""
},
{
"id":"GLUTDEHYD_RXN",
"name":"glutamate dehydrogenase (nadp+)",
"metabolites":{
"cpd00001_C":-1.0,
"cpd00005_C":1.0,
"cpd00006_C":-1.0,
"cpd00013_C":1.0,
"cpd00023_C":-1.0,
"cpd00024_C":1.0
},
"lower_bound":-1000.0,
"upper_bound":1000.0,
"gene_reaction_rule":"ACIAD1110 or ACIAD2680"
},
{
"id":"EXF_MALONATE_E",
"name":"exchange flux of malonate",
"metabolites":{
"cpd00308_E":-1.0
},
"lower_bound":-0.0,
"upper_bound":1000.0,
"gene_reaction_rule":""
},
{
"id":"C6_0_COA_THIOLASE_RXN",
"name":"acyl-coa thiolase (c6:0)",
"metabolites":{
"cpd00001_C":-1.0,
"cpd00010_C":1.0,
"cpd01113_C":1.0,
"cpd03124_C":-1.0
},
"lower_bound":0.0,
"upper_bound":1000.0,
"gene_reaction_rule":"ACIAD0341 or ACIAD3231 or ACIAD1023"
},
{
"id":"ACETATE_COA_LIGASE_RXN",
"name":"acetate--coa ligase",
"metabolites":{
"cpd00002_C":-1.0,
"cpd00010_C":-1.0,
"cpd00012_C":1.0,
"cpd00018_C":1.0,
"cpd00022_C":1.0,
"cpd00029_C":-1.0
},
"lower_bound":0.0,
"upper_bound":1000.0,
"gene_reaction_rule":"ACIAD1606 or ACIAD1611 or ACIAD3475"
},
{
"id":"RXN0_361",
"name":"cytidine hydrolase",
"metabolites":{
"cpd00001_C":-1.0,
"cpd00105_C":1.0,
"cpd00307_C":1.0,
"cpd00367_C":-1.0
},
"lower_bound":0.0,
"upper_bound":1000.0,
"gene_reaction_rule":"ACIAD2386"
},
{
"id":"RXN0_366",
"name":"guanosine hydrolase",
"metabolites":{
"cpd00001_C":-1.0,
"cpd00105_C":1.0,
"cpd00207_C":1.0,
"cpd00311_C":-1.0
},
"lower_bound":0.0,
"upper_bound":1000.0,
"gene_reaction_rule":"ACIAD2386"
},
{
"id":"DCTP_DEAM_RXN",
"name":"dctp deaminase",
"metabolites":{
"cpd00001_C":-1.0,
"cpd00013_C":1.0,
"cpd00356_C":-1.0,
"cpd00358_C":1.0
},
"lower_bound":0.0,
"upper_bound":1000.0,
"gene_reaction_rule":"ACIAD0776"
},
{
"id":"ACETOINDEHYDROG_RXN",
"name":"diacetyl reductase",
"metabolites":{
"cpd00003_C":-1.0,
"cpd00004_C":1.0,
"cpd00361_C":-1.0,
"cpd00551_C":1.0
},
"lower_bound":-1000.0,
"upper_bound":1000.0,
"gene_reaction_rule":"ACIAD1022"
},
{
"id":"PGPPHOSPHA_RXN",
"name":"phosphatidylglycerophosphatase",
"metabolites":{
"L_1_PHOSPHATIDYL_GLYCEROL_C":1.0,
"L_1_PHOSPHATIDYL_GLYCEROL_P_C":-1.0,
"cpd00001_C":-1.0,
"cpd00009_C":1.0
},
"lower_bound":0.0,
"upper_bound":1000.0,
"gene_reaction_rule":"ACIAD3574 or ACIAD2942 or ACIAD0554"
},
{
"id":"ARYLSULFAT_RXN",
"name":"phenylsulfatase",
"metabolites":{
"cpd00001_C":-1.0,
"cpd00048_C":1.0,
"cpd00127_C":1.0,
"cpd01471_C":-1.0
},
"lower_bound":0.0,
"upper_bound":1000.0,
"gene_reaction_rule":"ACIAD1598"
},
{
"id":"UDPNACETYLGLUCOSAMACYLTRANS_RXN",
"name":"acyl-[acyl-carrier-protein]--udp-n-acetylglucosamine o-acyltransferase",
"metabolites":{
"3_OHACYL_ACP_LPS_C":-1.0,
"ACP_C":1.0,
"cpd00037_C":-1.0,
"cpd02886_C":1.0
},
"lower_bound":-1000.0,
"upper_bound":1000.0,
"gene_reaction_rule":"ACIAD1382"
},
{
"id":"TRANS_RXN_27_BIS",
"name":"transport of 3-dehydro-shikimate [periplasm;cytosol](secondary symport)",
"metabolites":{
"cpd00067_C":1.0,
"cpd00067_P":-1.0,
"cpd01716_C":1.0,
"cpd01716_P":-1.0
},
"lower_bound":0.0,
"upper_bound":1000.0,
"gene_reaction_rule":"ACIAD0223"
},
{
"id":"TRANS_RXN_118",
"name":"transport of l-proline [extraorganism;cytosol](secondary symport)",
"metabolites":{
"cpd00129_C":1.0,
"cpd00129_E":-1.0,
"cpd00971_C":1.0,
"cpd00971_E":-1.0
},
"lower_bound":0.0,
"upper_bound":1000.0,
"gene_reaction_rule":"ACIAD1649"
},
{
"id":"ERYTH4PDEHYDROG_RXN",
"name":"d-erythrose 4-phosphate dehydrogenase",
"metabolites":{
"cpd00001_C":-1.0,
"cpd00003_C":-1.0,
"cpd00004_C":1.0,
"cpd00236_C":-1.0,
"cpd02147_C":1.0
},
"lower_bound":-1000.0,
"upper_bound":1000.0,
"gene_reaction_rule":"ACIAD1255"
},
{
"id":"C5_0_CARBOXY2ENOYLCOA_HYDRATASE_RXN",
"name":"dicarboxylate enoyl-coa hydratase (c5:0)",
"metabolites":{
"cpd00001_C":-1.0,
"cpd01599_C":-1.0,
"cpd16643_C":1.0
},
"lower_bound":0.0,
"upper_bound":1000.0,
"gene_reaction_rule":"ACIAD1692"
},
{
"id":"PROTEINSYN_RXN",
"name":"protein synthesis reaction",
"metabolites":{
"ALA_TRNAS_C":0.13,
"ARG_TRNAS_C":0.03,
"ASN_TRNAS_C":0.04,
"ASP_TRNAS_C":0.04,
"CHARGED_ALA_TRNAS_C":-0.13,
"CHARGED_ARG_TRNAS_C":-0.03,
"CHARGED_ASN_TRNAS_C":-0.04,
"CHARGED_ASP_TRNAS_C":-0.04,
"CHARGED_CYS_TRNAS_C":-0.02,
"CHARGED_GLN_TRNAS_C":-0.04,
"CHARGED_GLT_TRNAS_C":-0.04,
"CHARGED_GLY_TRNAS_C":-0.2,
"CHARGED_HIS_TRNAS_C":-0.01,
"CHARGED_ILE_TRNAS_C":-0.05,
"CHARGED_LEU_TRNAS_C":-0.07,
"CHARGED_LYS_TRNAS_C":-0.05,
"CHARGED_MET_TRNAS_C":-0.02,
"CHARGED_PHE_TRNAS_C":-0.02,
"CHARGED_PRO_TRNAS_C":-0.04,
"CHARGED_SER_TRNAS_C":-0.04,
"CHARGED_THR_TRNAS_C":-0.05,
"CHARGED_TRP_TRNAS_C":-0.01,
"CHARGED_TYR_TRNAS_C":-0.02,
"CHARGED_VAL_TRNAS_C":-0.08,
"CYS_TRNAS_C":0.02,
"GLN_TRNAS_C":0.04,
"GLT_TRNAS_C":0.04,
"GLY_TRNAS_C":0.2,
"HIS_TRNAS_C":0.01,
"ILE_TRNAS_C":0.05,
"LEU_TRNAS_C":0.07,
"LYS_TRNAS_C":0.05,
"MET_TRNAS_C":0.02,
"PHE_TRNAS_C":0.02,
"PROTEIN_W_C":0.0988,
"PRO_TRNAS_C":0.04,
"SER_TRNAS_C":0.04,
"THR_TRNAS_C":0.05,
"TRP_TRNAS_C":0.01,
"TYR_TRNAS_C":0.02,
"VAL_TRNAS_C":0.08,
"cpd00002_C":-4.2,
"cpd00008_C":4.2,
"cpd00009_C":4.2
},
"lower_bound":0.0,
"upper_bound":1000.0,
"gene_reaction_rule":""
},
{
"id":"TRANS_RXN_58",
"name":"transport of l-lysine [extraorganism;cytosol](secondary symport)",
"metabolites":{
"cpd00039_C":1.0,
"cpd00039_E":-1.0,
"cpd00067_C":1.0,
"cpd00067_P":-1.0
},
"lower_bound":0.0,
"upper_bound":1000.0,
"gene_reaction_rule":"ACIAD0700"
},
{
"id":"GTP_CYCLOHYDRO_I_RXN",
"name":"gtp cyclohydrolase i",
"metabolites":{
"cpd00001_C":-2.0,
"cpd00038_C":-1.0,
"cpd00047_C":1.0,
"cpd02978_C":1.0
},
"lower_bound":-1000.0,
"upper_bound":1000.0,
"gene_reaction_rule":"ACIAD2470"
},
{
"id":"_DEOXY_ADENYLATE_KINASE_RXN",
"name":"(deoxy)adenylate kinase",
"metabolites":{
"cpd00002_C":-1.0,
"cpd00008_C":1.0,
"cpd00177_C":1.0,
"cpd00294_C":-1.0
},
"lower_bound":-1000.0,
"upper_bound":1000.0,
"gene_reaction_rule":"ACIAD1105"
},
{
"id":"UROGENDECARBOX_RXN",
"name":"uroporphyrinogen decarboxylase",
"metabolites":{
"cpd00011_C":4.0,
"cpd00774_C":-1.0,
"cpd02083_C":1.0
},
"lower_bound":0.0,
"upper_bound":1000.0,
"gene_reaction_rule":"ACIAD2474"
},
{
"id":"GLYOXII_RXN",
"name":"hydroxyacylglutathione hydrolase",
"metabolites":{
"cpd00001_C":-1.0,
"cpd00042_C":1.0,
"cpd00221_C":1.0,
"cpd02182_C":-1.0
},
"lower_bound":-1000.0,
"upper_bound":1000.0,
"gene_reaction_rule":"ACIAD2452"
},
{
"id":"C6_0_3OXOCOA_THIOLASE_RXN",
"name":"3oxoacyl-coa thiolase (c6:0)",
"metabolites":{
"cpd00010_C":-1.0,
"cpd00022_C":1.0,
"cpd00120_C":1.0,
"cpd03123_C":-1.0
},
"lower_bound":-1000.0,
"upper_bound":1000.0,
"gene_reaction_rule":"ACIAD0334 or ACIAD1450 or ACIAD1706 or ACIAD1977 or ACIAD1689 or ACIAD2988 or ACIAD1569"
},
{
"id":"RXN0_383",
"name":"ctp pyrophosphorylase",
"metabolites":{
"cpd00001_C":-1.0,
"cpd00012_C":1.0,
"cpd00046_C":1.0,
"cpd00052_C":-1.0
},
"lower_bound":0.0,
"upper_bound":1000.0,
"gene_reaction_rule":"ACIAD3065"
},
{
"id":"RXN0_384",
"name":"datp pyrophosphorylase",
"metabolites":{
"cpd00001_C":-1.0,
"cpd00012_C":1.0,
"cpd00115_C":-1.0,
"cpd00294_C":1.0
},
"lower_bound":0.0,
"upper_bound":1000.0,
"gene_reaction_rule":"ACIAD3065"
},
{
"id":"RXN0_385",
"name":"dgtp pyrophosphorylase",
"metabolites":{
"cpd00001_C":-1.0,
"cpd00012_C":1.0,
"cpd00241_C":-1.0,
"cpd00296_C":1.0
},
"lower_bound":0.0,
"upper_bound":1000.0,
"gene_reaction_rule":"ACIAD3065"
},
{
"id":"TRANS_RXN_57",
"name":"transport of 4-aminobutyrate [extraorganism;cytosol](secondary symport)",
"metabolites":{
"cpd00067_C":1.0,
"cpd00067_P":-1.0,
"cpd00281_C":1.0,
"cpd00281_E":-1.0
},
"lower_bound":0.0,
"upper_bound":1000.0,
"gene_reaction_rule":"ACIAD3448"
},
{
"id":"HOMOSERKIN_RXN",
"name":"homoserine kinase",
"metabolites":{
"cpd00002_C":-1.0,
"cpd00008_C":1.0,
"cpd00227_C":-1.0,
"cpd00809_C":1.0
},
"lower_bound":-1000.0,
"upper_bound":1000.0,
"gene_reaction_rule":"ACIAD0834 or ACIAD1457 or ACIAD3403"
},
{
"id":"ALDOXIME_DEHYDRAT_RXN",
"name":"propionaldoxime aldoxime dehydratase",
"metabolites":{
"cpd00001_C":1.0,
"cpd16666_C":-1.0,
"cpd25464_C":1.0
},
"lower_bound":0.0,
"upper_bound":1000.0,
"gene_reaction_rule":"ACIAD1620"
},
{
"id":"1_5_1_29_RXN",
"name":"fmn reductase",
"metabolites":{
"cpd00003_C":-1.0,
"cpd00004_C":1.0,
"cpd00050_C":1.0,
"cpd01270_C":-1.0
},
"lower_bound":-1000.0,
"upper_bound":1000.0,
"gene_reaction_rule":"ACIAD3470"
},
{
"id":"1PFRUCTPHOSN_RXN",
"name":"1-phosphofructokinase",
"metabolites":{
"cpd00002_C":-1.0,
"cpd00008_C":1.0,
"cpd00290_C":1.0,
"cpd01850_C":-1.0
},
"lower_bound":0.0,
"upper_bound":1000.0,
"gene_reaction_rule":"ACIAD1992"
},
{
"id":"3_OHACYL_ACP_LPS_ASSEMBLY_RXN",
"name":"generic hydroxyacyl-acp of lps assembly",
"metabolites":{
"3_OHACYL_ACP_LPS_C":1.0,
"3_OHLAUROYL_ACP_C":-0.56,
"3_OHMYRISTOYL_ACP_C":-0.44
},
"lower_bound":-1000.0,
"upper_bound":1000.0,
"gene_reaction_rule":""
},
{
"id":"1_6_1_2_RXN",
"name":"nad(p)(+) transhydrogenase (ab-specific)",
"metabolites":{
"cpd00003_C":1.0,
"cpd00004_C":-1.0,
"cpd00005_C":1.0,
"cpd00006_C":-1.0,
"cpd00067_C":2.0,
"cpd00067_P":-2.0
},
"lower_bound":0.0,
"upper_bound":1000.0,
"gene_reaction_rule":"ACIAD3078 and ACIAD3079 and ACIAD3080"
},
{
"id":"TRANS_RXN_CPD_592",
"name":"transport of 4-guanidinobutanoate [extraorganism;cytosol](passive)",
"metabolites":{
"cpd00762_C":1.0,
"cpd00762_E":-1.0
},
"lower_bound":-1000.0,
"upper_bound":1000.0,
"gene_reaction_rule":""
},
{
"id":"C8_0_3HYDROXYCOA_DEHYDROGENASE_RXN",
"name":"hydroxyacyl-coa dehydrogenase (c8:0)",
"metabolites":{
"cpd00003_C":-1.0,
"cpd00004_C":1.0,
"cpd03120_C":-1.0,
"cpd03121_C":1.0
},
"lower_bound":0.0,
"upper_bound":1000.0,
"gene_reaction_rule":"ACIAD0335 or ACIAD1565 or ACIAD2416"
},
{
"id":"GLYCEROL_3_P_ACYLTRANSFER_PL_RXN",
"name":"glycerol-3-phosphate acyltransferase",
"metabolites":{
"ACP_C":1.0,
"ACYL_ACP_PL_C":-1.0,
"ACYL_SN_GLYCEROL_3P_PL_C":1.0,
"cpd00080_C":-1.0
},
"lower_bound":0.0,
"upper_bound":1000.0,
"gene_reaction_rule":"ACIAD3232"
},
{
"id":"2_6_1_76_RXN",
"name":"diaminobutyrate-2-oxoglutarate transaminase",
"metabolites":{
"cpd00023_C":-1.0,
"cpd00024_C":1.0,
"cpd00346_C":-1.0,
"cpd02095_C":1.0
},
"lower_bound":-1000.0,
"upper_bound":1000.0,
"gene_reaction_rule":"ACIAD1210"
},
{
"id":"RXN_2881",
"name":"formaldehyde thf binding",
"metabolites":{
"cpd00001_C":1.0,
"cpd00055_C":-1.0,
"cpd00087_C":-1.0,
"cpd00125_C":1.0
},
"lower_bound":0.0,
"upper_bound":1000.0,
"gene_reaction_rule":"SPONT"
},
{
"id":"TRANS_RXN_ETHANESULFONATE",
"name":"transport of ethane sulfonate [extraorganism;cytosol](primary active atp)",
"metabolites":{
"cpd00001_C":-1.0,
"cpd00002_C":-1.0,
"cpd00008_C":1.0,
"cpd00009_C":1.0,
"cpd11579_C":1.0,
"cpd11579_E":-1.0
},
"lower_bound":0.0,
"upper_bound":1000.0,
"gene_reaction_rule":"(ACIAD0034 and ACIAD0035 and ACIAD0037 and ACIAD0038) or (ACIAD1523 and ACIAD1524 and ACIAD1525 and ACIAD1526) or (ACIAD1586 and ACIAD1587 and ACIAD1593 and ACIAD1601)"
},
{
"id":"ASPARTATE_RACEMASE_RXN",
"name":"aspartate racemase",
"metabolites":{
"cpd00041_C":-1.0,
"cpd00320_C":1.0
},
"lower_bound":-1000.0,
"upper_bound":1000.0,
"gene_reaction_rule":"ACIAD0318"
},
{
"id":"3_HYDROXYISOBUTYRATE_DEHYDROGENASE_RXN",
"name":"3-hydroxyisobutyrate dehydrogenase",
"metabolites":{
"cpd00003_C":-1.0,
"cpd00004_C":1.0,
"cpd00287_C":1.0,
"cpd00876_C":-1.0
},
"lower_bound":-1000.0,
"upper_bound":1000.0,
"gene_reaction_rule":"ACIAD0032 or ACIAD2316 or ACIAD1605 or ACIAD3074"
},
{
"id":"TRANS_RXN_C18_1_C9_",
"name":"transport of cis-octadec-9-enoate [extraorganism;cytosol](secondary symport)",
"metabolites":{
"cpd00067_C":1.0,
"cpd00067_P":-1.0,
"cpd00536_C":1.0,
"cpd00536_E":-1.0
},
"lower_bound":-1000.0,
"upper_bound":1000.0,
"gene_reaction_rule":"ACIAD0835 or ACIAD1427 or ACIAD2788 or ACIAD3392"
},
{
"id":"EXF_ETOH_E",
"name":"exchange flux of ethanol",
"metabolites":{
"cpd00363_E":-1.0
},
"lower_bound":-0.0,
"upper_bound":1000.0,
"gene_reaction_rule":""
},
{
"id":"EXF_URIDINE_E",
"name":"exchange flux of uridine",
"metabolites":{
"cpd00249_E":-1.0
},
"lower_bound":-0.0,
"upper_bound":1000.0,
"gene_reaction_rule":""
},
{
"id":"ADENOSYLHOMOCYSTEINE_NUCLEOSIDASE_RXN",
"name":"adenosylhomocysteine nucleosidase",
"metabolites":{
"cpd00001_C":-1.0,
"cpd00019_C":-1.0,
"cpd00128_C":1.0,
"cpd02227_C":1.0
},
"lower_bound":-1000.0,
"upper_bound":1000.0,
"gene_reaction_rule":"ACIAD0773"
},
{
"id":"C14_0_3OXOACP_SYNTHASE_RXN",
"name":"3oxoacyl-acp synthase (c14:0)",
"metabolites":{
"ACP_C":1.0,
"C12_0_ACP_C":-1.0,
"C14_0_3OXOACP_C":1.0,
"MALONYL_ACP_C":-1.0,
"cpd00011_C":1.0
},
"lower_bound":0.0,
"upper_bound":1000.0,
"gene_reaction_rule":"ACIAD0580 or ACIAD0879 or ACIAD0583 or ACIAD2101"
},
{
"id":"PYRIMSYN1_RXN",
"name":"4-amino-2-methyl-5-phosphomethylpyrimidine synthetase",
"metabolites":{
"cpd00001_C":-1.0,
"cpd00047_C":2.0,
"cpd02140_C":-1.0,
"cpd02775_C":1.0
},
"lower_bound":-1000.0,
"upper_bound":1000.0,
"gene_reaction_rule":"ACIAD0276"
},
{
"id":"TRANS_RXNIM_5839",
"name":"transport of l-lysine [cytosol;extraorganism](secondary antiport)",
"metabolites":{
"cpd00039_C":-1.0,
"cpd00039_E":1.0,
"cpd00067_C":1.0,
"cpd00067_P":-1.0
},
"lower_bound":0.0,
"upper_bound":1000.0,
"gene_reaction_rule":"ACIAD0966"
},
{
"id":"NADH_DEHYDROG_A_RXN",
"name":"nadh dehydrogenase (ubiquinone)",
"metabolites":{
"cpd00003_C":1.0,
"cpd00004_C":-1.0,
"cpd00067_C":-5.0,
"cpd00067_P":4.0,
"cpd15560_C":-1.0,
"cpd15561_C":1.0
},
"lower_bound":0.0,
"upper_bound":1000.0,
"gene_reaction_rule":"ACIAD0743 and ACIAD0742 and ACIAD0741 and ACIAD0740 and ACIAD0739 and ACIAD0738 and ACIAD0737 and ACIAD0736 and ACIAD0735 and ACIAD0734 and ACIAD0733 and ACIAD0731 and ACIAD0730"
},
{
"id":"ACSERLY_RXN",
"name":"cysteine synthase",
"metabolites":{
"cpd00029_C":1.0,
"cpd00084_C":1.0,
"cpd00239_C":-1.0,
"cpd00722_C":-1.0
},
"lower_bound":-1000.0,
"upper_bound":1000.0,
"gene_reaction_rule":"ACIAD1662 or ACIAD3071"
},
{
"id":"EXF_CPD_6501_E",
"name":"exchange flux of benzylacetate",
"metabolites":{
"cpd11193_E":-1.0
},
"lower_bound":-0.0,
"upper_bound":1000.0,
"gene_reaction_rule":""
},
{
"id":"C6_1_C4_3OXOCOA_THIOLASE_RXN",
"name":"3oxoacyl-coa thiolase (c6:1(c4))",
"metabolites":{
"cpd00010_C":-1.0,
"cpd00022_C":1.0,
"cpd00650_C":1.0,
"cpd16647_C":-1.0
},
"lower_bound":-1000.0,
"upper_bound":1000.0,
"gene_reaction_rule":"ACIAD0334 or ACIAD1450 or ACIAD1706 or ACIAD1977 or ACIAD1689 or ACIAD2988 or ACIAD1569"
},
{
"id":"C16_1_C7_3HYDROXYCOA_DEHYDROGENASE_RXN",
"name":"hydroxyacyl-coa dehydrogenase (c16:1(c7))",
"metabolites":{
"cpd00003_C":-1.0,
"cpd00004_C":1.0,
"cpd32368_C":1.0,
"cpd32607_C":-1.0
},
"lower_bound":0.0,
"upper_bound":1000.0,
"gene_reaction_rule":"ACIAD0335 or ACIAD1565 or ACIAD2416"
},
{
"id":"TRANS_RXNIM_5839_BIS",
"name":"transport of l-arginine [cytosol;extraorganism](secondary antiport)",
"metabolites":{
"cpd00051_C":-1.0,
"cpd00051_E":1.0,
"cpd00067_C":1.0,
"cpd00067_P":-1.0
},
"lower_bound":0.0,
"upper_bound":1000.0,
"gene_reaction_rule":"ACIAD0966"
},
{
"id":"EXF_SHIKIMATE_E",
"name":"exchange flux of shikimate",
"metabolites":{
"cpd00383_E":-1.0
},
"lower_bound":-0.0,
"upper_bound":1000.0,
"gene_reaction_rule":""
},
{
"id":"OPPSYN_RXN",
"name":"octaprenyl pyrophosphate synthase",
"metabolites":{
"cpd00012_C":5.0,
"cpd00113_C":-5.0,
"cpd00350_C":-1.0,
"cpd02557_C":1.0
},
"lower_bound":0.0,
"upper_bound":1000.0,
"gene_reaction_rule":"ACIAD2940"
},
{
"id":"C18_1_C9_ALDEHYDE_REDUCTASE_RXN",
"name":"long chain aldehyde reductase (c18:1(c9))",
"metabolites":{
"cpd00003_C":1.0,
"cpd00004_C":-1.0,
"cpd16639_C":1.0,
"cpd23520_C":-1.0
},
"lower_bound":-1000.0,
"upper_bound":1000.0,
"gene_reaction_rule":""
},
{
"id":"C14_0_3HYDROXYACP_DEHYDRATASE_RXN",
"name":"3hydroxyacyl-acp dehydratase (c14:0)",
"metabolites":{
"3_OHMYRISTOYL_ACP_C":-1.0,
"cpd00001_C":1.0,
"cpd03127_C":1.0
},
"lower_bound":0.0,
"upper_bound":1000.0,
"gene_reaction_rule":"ACIAD1381"
},
{
"id":"SHIKIMATE_KINASE_RXN",
"name":"shikimate-kinase",
"metabolites":{
"cpd00002_C":-1.0,
"cpd00008_C":1.0,
"cpd00383_C":-1.0,
"cpd02030_C":1.0
},
"lower_bound":0.0,
"upper_bound":1000.0,
"gene_reaction_rule":"ACIAD3354"
},
{
"id":"TRANS_RXN_CPDIM_6114",
"name":"transport of salicylacetate [extraorganism;cytosol](passive)",
"metabolites":{
"cpd01006_C":1.0,
"cpd01006_E":-1.0
},
"lower_bound":-1000.0,
"upper_bound":1000.0,
"gene_reaction_rule":""
},
{
"id":"RXN0_949",
"name":"lipoate synthase",
"metabolites":{
"C8_0_ACP_C":-1.0,
"LIPOYL_ACP_C":1.0,
"cpd00017_C":-2.0,
"cpd00060_C":2.0,
"cpd03091_C":2.0
},
"lower_bound":0.0,
"upper_bound":1000.0,
"gene_reaction_rule":"ACIAD1015 or ACIAD2275"
},
{
"id":"GROWTH_RXN",
"name":"overall biomass assembly",
"metabolites":{
"AFATTYACID_FREE_W_C":-0.002,
"COFACTOR_W_C":-0.032,
"DNA_W_C":-0.032,
"LPS_W_C":-0.003,
"PEPTIDO_W_E":-0.028,
"PL_W_C":-0.041,
"POLYSACCH_FREE_W_C":-0.021,
"PROTEIN_W_C":-0.633,
"RNA_W_C":-0.2,
"TRIACYLGLYCEROL_W_C":-0.002,
"WAX_ESTER_W_C":-0.006,
"cpd00001_C":-40.0,
"cpd00002_C":-40.0,
"cpd00008_C":40.0,
"cpd00009_C":40.0
},
"lower_bound":0.0,
"upper_bound":1000.0,
"gene_reaction_rule":"",
"objective_coefficient":1.0
},
{
"id":"TRANS_RXN_27",
"name":"transport of shikimate [periplasm;cytosol](secondary symport)",
"metabolites":{
"cpd00067_C":1.0,
"cpd00067_P":-1.0,
"cpd00383_C":1.0,
"cpd00383_P":-1.0
},
"lower_bound":0.0,
"upper_bound":1000.0,
"gene_reaction_rule":"ACIAD0223"
},
{
"id":"ARGININE_N_SUCCINYLTRANSFERASE_RXN",
"name":"arginine n-succinyltransferase",
"metabolites":{
"cpd00010_C":1.0,
"cpd00051_C":-1.0,
"cpd00078_C":-1.0,
"cpd02103_C":1.0
},
"lower_bound":-1000.0,
"upper_bound":1000.0,
"gene_reaction_rule":"ACIAD1286"
},
{
"id":"RIBULP3EPIM_RXN",
"name":"ribulose phosphate 3-epimerase",
"metabolites":{
"cpd00171_C":-1.0,
"cpd00198_C":1.0
},
"lower_bound":-1000.0,
"upper_bound":1000.0,
"gene_reaction_rule":"ACIAD0685"
},
{
"id":"TRANS_RXN_ADIPATE",
"name":"transport of adipate [extraorganism;cytosol](secondary symport)",
"metabolites":{
"cpd00067_C":1.0,
"cpd00067_P":-1.0,
"cpd03642_C":1.0,
"cpd03642_E":-1.0
},
"lower_bound":-1000.0,
"upper_bound":1000.0,
"gene_reaction_rule":"ACIAD1694"
},
{
"id":"THREONINE_TRNA_LIGASE_RXN",
"name":"threonine--trna ligase",
"metabolites":{
"CHARGED_THR_TRNAS_C":1.0,
"THR_TRNAS_C":-1.0,
"cpd00002_C":-1.0,
"cpd00012_C":1.0,
"cpd00018_C":1.0,
"cpd00161_C":-1.0
},
"lower_bound":0.0,
"upper_bound":1000.0,
"gene_reaction_rule":"ACIAD3055"
},
{
"id":"TREHALOSEPHOSPHA_RXN",
"name":"trehalose-phosphatase",
"metabolites":{
"cpd00001_C":-1.0,
"cpd00009_C":1.0,
"cpd00523_C":-1.0,
"cpd00794_C":1.0
},
"lower_bound":0.0,
"upper_bound":1000.0,
"gene_reaction_rule":"ACIAD1775"
},
{
"id":"RXNIM_6212",
"name":"4-hydroxy-benzyl-alcohol dehydrogenase",
"metabolites":{
"cpd00003_C":-1.0,
"cpd00004_C":1.0,
"cpd00484_C":1.0,
"cpd15378_C":-1.0
},
"lower_bound":-1000.0,
"upper_bound":1000.0,
"gene_reaction_rule":"ACIAD1429 or ACIAD1578"
},
{
"id":"H2PTERIDINEPYROPHOSPHOKIN_RXN",
"name":"2-amino-4-hydroxy-6-hydroxymethyldihydropteridine pyrophosphokinase",
"metabolites":{
"cpd00002_C":-1.0,
"cpd00018_C":1.0,
"cpd00954_C":-1.0,
"cpd02920_C":1.0
},
"lower_bound":-1000.0,
"upper_bound":1000.0,
"gene_reaction_rule":"ACIAD3062"
},
{
"id":"RXNIM_6210",
"name":"3,4-dihydroxyphenylpropionate coa ligase",
"metabolites":{
"cpd00002_C":-1.0,
"cpd00010_C":-1.0,
"cpd00012_C":1.0,
"cpd00018_C":1.0,
"cpd07333_C":-1.0,
"cpd35716_C":1.0
},
"lower_bound":0.0,
"upper_bound":1000.0,
"gene_reaction_rule":"ACIAD1724"
},
{
"id":"RXNIM_6211",
"name":"3,4-dihydroxyphenylpropionyl-coa dehydrogenase",
"metabolites":{
"cpd00015_C":-1.0,
"cpd00270_C":1.0,
"cpd00982_C":1.0,
"cpd35716_C":-1.0
},
"lower_bound":-1000.0,
"upper_bound":1000.0,
"gene_reaction_rule":"ACIAD1723"
},
{
"id":"C10_0_CARBOXY_COATRANSFERASE_RXN",
"name":"dicarboxylate acyl coa transferase (c10:0)",
"metabolites":{
"cpd00002_C":-1.0,
"cpd00010_C":-1.0,
"cpd00012_C":1.0,
"cpd00018_C":1.0,
"cpd05192_C":-1.0,
"cpd16607_C":1.0
},
"lower_bound":0.0,
"upper_bound":1000.0,
"gene_reaction_rule":"ACIAD1695 and ACIAD1696"
},
{
"id":"ALANINE_AMINOTRANSFERASE_RXN",
"name":"ALANINE-AMINOTRANSFERASE-RXN",
"metabolites":{
"cpd00020_C":-1.0,
"cpd00023_C":-1.0,
"cpd00024_C":1.0,
"cpd00035_C":1.0
},
"lower_bound":-1000.0,
"upper_bound":1000.0,
"gene_reaction_rule":"ACIAD2080 or ACIAD2087"
},
{
"id":"DXPREDISOM_RXN",
"name":"1-deoxy-d-xylulose reductoisomerase",
"metabolites":{
"cpd00005_C":1.0,
"cpd00006_C":-1.0,
"cpd08286_C":-1.0,
"cpd08289_C":1.0
},
"lower_bound":-1000.0,
"upper_bound":1000.0,
"gene_reaction_rule":"ACIAD1376"
},
{
"id":"4_HYDROXYBENZOATE_3_MONOOXYGENASE_RXN",
"name":"4-hydroxybenzoate 3-monooxygenase",
"metabolites":{
"cpd00001_C":1.0,
"cpd00005_C":-1.0,
"cpd00006_C":1.0,
"cpd00007_C":-1.0,
"cpd00136_C":-1.0,
"cpd00197_C":1.0
},
"lower_bound":0.0,
"upper_bound":1000.0,
"gene_reaction_rule":"ACIAD1719"
},
{
"id":"ACETYLGLUTKIN_RXN",
"name":"acetylglutamate kinase",
"metabolites":{
"cpd00002_C":-1.0,
"cpd00008_C":1.0,
"cpd00477_C":-1.0,
"cpd02552_C":1.0
},
"lower_bound":-1000.0,
"upper_bound":1000.0,
"gene_reaction_rule":"ACIAD0903"
},
{
"id":"3_6_1_41_RXN",
"name":"bis(5'-nucleosyl)-tetraphosphatase (symmetrical)",
"metabolites":{
"cpd00001_C":-1.0,
"cpd00008_C":2.0,
"cpd00924_C":-1.0
},
"lower_bound":0.0,
"upper_bound":1000.0,
"gene_reaction_rule":"ACIAD3008"
},
{
"id":"4_COUMARATE_COA_LIGASE_RXN",
"name":"4-coumarate--coa ligase",
"metabolites":{
"cpd00002_C":-1.0,
"cpd00010_C":-1.0,
"cpd00012_C":1.0,
"cpd00018_C":1.0,
"cpd00192_C":1.0,
"cpd00604_C":-1.0
},
"lower_bound":0.0,
"upper_bound":1000.0,
"gene_reaction_rule":"ACIAD1724"
},
{
"id":"2_METHYLCITRATE_DEHYDRATASE_RXN",
"name":"2-methylcitrate dehydratase",
"metabolites":{
"cpd00001_C":1.0,
"cpd01501_C":-1.0,
"cpd02597_C":1.0
},
"lower_bound":-1000.0,
"upper_bound":1000.0,
"gene_reaction_rule":"ACIAD2755"
},
{
"id":"1_14_12_1_RXN",
"name":"anthranilate 1,2-dioxygenase (deaminating, decarboxylating)",
"metabolites":{
"cpd00005_C":-1.0,
"cpd00006_C":1.0,
"cpd00007_C":-1.0,
"cpd00011_C":1.0,
"cpd00013_C":1.0,
"cpd00077_C":1.0,
"cpd00093_C":-1.0
},
"lower_bound":0.0,
"upper_bound":1000.0,
"gene_reaction_rule":"ACIAD2669 and ACIAD2670 and ACIAD2671"
},
{
"id":"TRANS_RXN_DEOXYCYTIDINE",
"name":"transport of deoxycytidine [extraorganism;cytosol](secondary symport)",
"metabolites":{
"cpd00067_C":1.0,
"cpd00067_P":-1.0,
"cpd00654_C":1.0,
"cpd00654_E":-1.0
},
"lower_bound":0.0,
"upper_bound":1000.0,
"gene_reaction_rule":"ACIAD2389"
},
{
"id":"C18_1_C9_COA_REDUCTASE_RXN",
"name":"fatty acyl-coa reductase (c18:1(c9))",
"metabolites":{
"cpd00005_C":-1.0,
"cpd00006_C":1.0,
"cpd00010_C":1.0,
"cpd16638_C":-1.0,
"cpd23520_C":1.0
},
"lower_bound":-1000.0,
"upper_bound":1000.0,
"gene_reaction_rule":"ACIAD3383"
},
{
"id":"TRANS_RXN_CPD_110",
"name":"transport of salicylate [extraorganism;cytosol](passive)",
"metabolites":{
"cpd00599_C":1.0,
"cpd00599_E":-1.0
},
"lower_bound":0.0,
"upper_bound":1000.0,
"gene_reaction_rule":""
},
{
"id":"NICONUCADENYLYLTRAN_RXN",
"name":"nicotinate-nucleotide adenylyltransferase",
"metabolites":{
"NICOTINATE_NUCLEOTIDE_C":-1.0,
"cpd00002_C":-1.0,
"cpd00012_C":1.0,
"cpd00638_C":1.0
},
"lower_bound":0.0,
"upper_bound":1000.0,
"gene_reaction_rule":"ACIAD2606"
},
{
"id":"L_GLN_FRUCT_6_P_AMINOTRANS_RXN",
"name":"glucosamine--fructose-6-phosphate aminotransferase (isomerizing)",
"metabolites":{
"cpd00023_C":1.0,
"cpd00053_C":-1.0,
"cpd00288_C":1.0,
"cpd15394_C":-1.0
},
"lower_bound":0.0,
"upper_bound":1000.0,
"gene_reaction_rule":"ACIAD3576"
},
{
"id":"FADSYN_RXN",
"name":"fmn adenylyltransferase",
"metabolites":{
"cpd00002_C":-1.0,
"cpd00012_C":1.0,
"cpd00015_C":1.0,
"cpd00050_C":-1.0
},
"lower_bound":-1000.0,
"upper_bound":1000.0,
"gene_reaction_rule":"ACIAD0023"
},
{
"id":"EXF_3_KETOBUTYRATE_E",
"name":"exchange flux of acetoacetate",
"metabolites":{
"cpd00142_E":-1.0
},
"lower_bound":-0.0,
"upper_bound":1000.0,
"gene_reaction_rule":""
},
{
"id":"UDPHYDROXYMYRGLUCOSAMNACETYLTRANS_RXN",
"name":"udp-3-o-[3-hydroxylauroyl] glucosamine n-acyltransferase",
"metabolites":{
"3_OHACYL_ACP_LPS_C":-1.0,
"ACP_C":1.0,
"cpd02835_C":1.0,
"cpd03584_C":-1.0
},
"lower_bound":-1000.0,
"upper_bound":1000.0,
"gene_reaction_rule":"ACIAD1380"
},
{
"id":"C4_0_3HYDROXYACP_DEHYDRATASE_RXN",
"name":"3hydroxyacyl-acp dehydratase (c4:0)",
"metabolites":{
"C4_0_2ENOYLACP_C":1.0,
"C4_0_3HYDROXYACP_C":-1.0,
"cpd00001_C":1.0
},
"lower_bound":0.0,
"upper_bound":1000.0,
"gene_reaction_rule":"ACIAD1381"
},
{
"id":"SUCCGLUDESUCC_RXN",
"name":"succinylglutamate desuccinylase",
"metabolites":{
"cpd00001_C":-1.0,
"cpd00023_C":1.0,
"cpd00036_C":1.0,
"cpd03525_C":-1.0
},
"lower_bound":0.0,
"upper_bound":1000.0,
"gene_reaction_rule":"ACIAD1289"
},
{
"id":"UDPREDUCT_RXN",
"name":"udp reductase",
"metabolites":{
"cpd00001_C":1.0,
"cpd00014_C":-1.0,
"cpd00978_C":1.0,
"cpd11420_C":1.0,
"cpd11421_C":-1.0
},
"lower_bound":-1000.0,
"upper_bound":1000.0,
"gene_reaction_rule":"ACIAD0724 and ACIAD0722"
},
{
"id":"ACETOLACTSYN_RXN",
"name":"acetolactate synthase",
"metabolites":{
"cpd00011_C":1.0,
"cpd00020_C":-2.0,
"cpd00668_C":1.0
},
"lower_bound":0.0,
"upper_bound":1000.0,
"gene_reaction_rule":"(ACIAD0999 or ACIAD3104) and ACIAD3103"
},
{
"id":"C18_0_3HYDROXYCOA_DEHYDROGENASE_RXN",
"name":"hydroxyacyl-coa dehydrogenase (c18:0)",
"metabolites":{
"cpd00003_C":-1.0,
"cpd00004_C":1.0,
"cpd14935_C":1.0,
"cpd14936_C":-1.0
},
"lower_bound":0.0,
"upper_bound":1000.0,
"gene_reaction_rule":"ACIAD0335 or ACIAD1565 or ACIAD2416"
},
{
"id":"KDO_8PPHOSPHAT_RXN",
"name":"3-deoxy-manno-octulosonate-8-phosphatase",
"metabolites":{
"cpd00001_C":-1.0,
"cpd00009_C":1.0,
"cpd00875_C":1.0,
"cpd02730_C":-1.0
},
"lower_bound":0.0,
"upper_bound":1000.0,
"gene_reaction_rule":"ACIAD1483"
},
{
"id":"C12_0_2ENOYLACP_ISOMERASE_RXN",
"name":"2enoyl-acp isomerase (c12:0)",
"metabolites":{
"C12_0_2ENOYLACP_C":-1.0,
"C12_1_C3__ACP_C":1.0
},
"lower_bound":-1000.0,
"upper_bound":1000.0,
"gene_reaction_rule":""
},
{
"id":"3_HYDROXYBUTYRATE_DEHYDROGENASE_RXN",
"name":"3-hydroxybutyrate dehydrogenase",
"metabolites":{
"cpd00003_C":-1.0,
"cpd00004_C":1.0,
"cpd00142_C":1.0,
"cpd00797_C":-1.0
},
"lower_bound":-1000.0,
"upper_bound":1000.0,
"gene_reaction_rule":"ACIAD2509"
},
{
"id":"AIRS_RXN",
"name":"phosphoribosylformylglycinamidine cyclo-ligase",
"metabolites":{
"cpd00002_C":-1.0,
"cpd00008_C":1.0,
"cpd00009_C":1.0,
"cpd02140_C":1.0,
"cpd02826_C":-1.0
},
"lower_bound":0.0,
"upper_bound":1000.0,
"gene_reaction_rule":"ACIAD2634"
},
{
"id":"C10_0_CARBOXY3OXOCOA_THIOLASE_RXN",
"name":"dicarboxylate 3oxoacyl-coa thiolase (c10:0)",
"metabolites":{
"cpd00010_C":-1.0,
"cpd00022_C":1.0,
"cpd16606_C":-1.0,
"cpd16652_C":1.0
},
"lower_bound":-1000.0,
"upper_bound":1000.0,
"gene_reaction_rule":"ACIAD1689"
},
{
"id":"3_DEHYDROQUINATE_SYNTHASE_RXN",
"name":"3-dehydroquinate synthase",
"metabolites":{
"cpd00009_C":1.0,
"cpd00699_C":1.0,
"cpd02857_C":-1.0
},
"lower_bound":0.0,
"upper_bound":1000.0,
"gene_reaction_rule":"ACIAD3353"
},
{
"id":"PEPCARBOX_RXN",
"name":"phosphoenolpyruvate carboxylase",
"metabolites":{
"cpd00001_C":-1.0,
"cpd00009_C":1.0,
"cpd00011_C":-1.0,
"cpd00032_C":1.0,
"cpd00061_C":-1.0
},
"lower_bound":0.0,
"upper_bound":1000.0,
"gene_reaction_rule":"ACIAD3627"
},
{
"id":"ACYL_ACP_PL_ASSEMBLY_RXN",
"name":"generic acyl acp of phospholipids assembly",
"metabolites":{
"ACYL_ACP_PL_C":1.0,
"C16_0_ACP_C":-0.1,
"C16_1_C7__ACP_C":-0.2,
"C18_0_ACP_C":-0.1,
"C18_1_C9__ACP_C":-0.6
},
"lower_bound":-1000.0,
"upper_bound":1000.0,
"gene_reaction_rule":""
},
{
"id":"AFATTYACID_TAG_ASSEMBLY_RXN",
"name":"generic fatty acid of triacylglycerol assembly",
"metabolites":{
"AFATTYACID_TAG_C":1.0,
"cpd00214_C":-0.717,
"cpd00536_C":-0.149,
"cpd01080_C":-0.06,
"cpd03847_C":-0.022,
"cpd34461_C":-0.052
},
"lower_bound":-1000.0,
"upper_bound":1000.0,
"gene_reaction_rule":""
},
{
"id":"EXF_L_ORNITHINE_E",
"name":"exchange flux of l-ornithine",
"metabolites":{
"cpd00064_E":-1.0
},
"lower_bound":-0.0,
"upper_bound":1000.0,
"gene_reaction_rule":""
},
{
"id":"C16_0_COA_DEHYDROGENASE_RXN",
"name":"acyl-coa dehydrogenase (c16:0)",
"metabolites":{
"cpd00015_C":-1.0,
"cpd00134_C":-1.0,
"cpd00982_C":1.0,
"cpd03126_C":1.0
},
"lower_bound":0.0,
"upper_bound":1000.0,
"gene_reaction_rule":"ACIAD0452 or ACIAD3262 or ACIAD2537 or ACIAD2277 or ACIAD1886 or ACIAD1821 or ACIAD1819 or ACIAD1700 or ACIAD1693 or ACIAD1607 or ACIAD1413 or ACIAD0623 or ACIAD0624 or ACIAD0428 or ACIAD0427 or ACIAD1723 or ACIAD2226 or ACIAD3191 or ACIAD1566"
},
{
"id":"CHD_RXN",
"name":"choline dehydrogenase",
"metabolites":{
"cpd00015_C":-1.0,
"cpd00098_C":-1.0,
"cpd00447_C":1.0,
"cpd00982_C":1.0
},
"lower_bound":-1000.0,
"upper_bound":1000.0,
"gene_reaction_rule":"ACIAD1008"
},
{
"id":"FABAUNSATDEHYDR_RXN",
"name":"3hydroxyacyl-acp dehydratase (c10:0)",
"metabolites":{
"C10_0_2ENOYLACP_C":1.0,
"C10_0_3HYDROXYACP_C":-1.0,
"cpd00001_C":1.0
},
"lower_bound":0.0,
"upper_bound":1000.0,
"gene_reaction_rule":"ACIAD1381"
},
{
"id":"NITRATREDUCT_RXN",
"name":"nitrate reductase",
"metabolites":{
"cpd00001_C":1.0,
"cpd00075_C":1.0,
"cpd00209_C":-1.0,
"cpd15560_C":1.0,
"cpd15561_C":-1.0
},
"lower_bound":0.0,
"upper_bound":1000.0,
"gene_reaction_rule":"ACIAD1908 and ACIAD1909"
},
{
"id":"C10_0_COA_DEHYDROGENASE_RXN",
"name":"acyl-coa dehydrogenase (c10:0)",
"metabolites":{
"cpd00015_C":-1.0,
"cpd00982_C":1.0,
"cpd03128_C":-1.0,
"cpd03129_C":1.0
},
"lower_bound":0.0,
"upper_bound":1000.0,
"gene_reaction_rule":"ACIAD0452 or ACIAD3262 or ACIAD2537 or ACIAD2277 or ACIAD1886 or ACIAD1821 or ACIAD1819 or ACIAD1700 or ACIAD1693 or ACIAD1607 or ACIAD1413 or ACIAD0623 or ACIAD0624 or ACIAD0428 or ACIAD0427 or ACIAD1723 or ACIAD2226 or ACIAD3191 or ACIAD1566"
},
{
"id":"EXF_ANTHRANILATE_E",
"name":"exchange flux of 2-aminobenzoate",
"metabolites":{
"cpd00093_E":-1.0
},
"lower_bound":-0.0,
"upper_bound":1000.0,
"gene_reaction_rule":""
},
{
"id":"EXF_D_GLUCARATE_E",
"name":"exchange flux of d-glucarate",
"metabolites":{
"cpd00571_E":-1.0
},
"lower_bound":-0.0,
"upper_bound":1000.0,
"gene_reaction_rule":""
},
{
"id":"DCMP_NUCLEOTIDASE_RXN",
"name":"dcmp nucleotidase",
"metabolites":{
"cpd00001_C":-1.0,
"cpd00009_C":1.0,
"cpd00206_C":-1.0,
"cpd00654_C":1.0
},
"lower_bound":0.0,
"upper_bound":1000.0,
"gene_reaction_rule":"ACIAD0015 or ACIAD1227"
},
{
"id":"TRANS_RXN_104_BIS",
"name":"transport of d-lactate [extraorganism;cytosol](secondary symport)",
"metabolites":{
"cpd00067_C":1.0,
"cpd00067_P":-1.0,
"cpd00221_C":1.0,
"cpd00221_E":-1.0
},
"lower_bound":-1000.0,
"upper_bound":1000.0,
"gene_reaction_rule":"ACIAD0106"
},
{
"id":"SERINE_O_ACETTRAN_RXN",
"name":"serine o-acetyltransferase",
"metabolites":{
"cpd00010_C":1.0,
"cpd00022_C":-1.0,
"cpd00054_C":-1.0,
"cpd00722_C":1.0
},
"lower_bound":-1000.0,
"upper_bound":1000.0,
"gene_reaction_rule":"ACIAD2091 or ACIAD1584 or ACIAD1763"
},
{
"id":"C5_0_2ENOYLCOA_HYDRATASE_RXN",
"name":"enoyl-coa hydratase (c5:0)",
"metabolites":{
"cpd00001_C":-1.0,
"cpd01613_C":-1.0,
"cpd23920_C":1.0
},
"lower_bound":0.0,
"upper_bound":1000.0,
"gene_reaction_rule":"ACIAD1692 or ACIAD1726 or ACIAD1820 or ACIAD1829 or ACIAD2351 or ACIAD3073 or ACIAD1609 or ACIAD0335 or ACIAD1608 or ACIAD1701 or ACIAD1565 or ACIAD2416"
},
{
"id":"EXF_NA_E",
"name":"exchange flux of na+",
"metabolites":{
"cpd00971_E":-1.0
},
"lower_bound":-1000.0,
"upper_bound":1000.0,
"gene_reaction_rule":""
},
{
"id":"7KAPSYN_RXN",
"name":"8-amino-7-oxononanoate synthase",
"metabolites":{
"cpd00010_C":1.0,
"cpd00011_C":1.0,
"cpd00035_C":-1.0,
"cpd00782_C":-1.0,
"cpd00800_C":1.0
},
"lower_bound":0.0,
"upper_bound":1000.0,
"gene_reaction_rule":"ACIAD0857"
},
{
"id":"VALINE_TRNA_LIGASE_RXN",
"name":"valine--trna ligase",
"metabolites":{
"CHARGED_VAL_TRNAS_C":1.0,
"VAL_TRNAS_C":-1.0,
"cpd00002_C":-1.0,
"cpd00012_C":1.0,
"cpd00018_C":1.0,
"cpd00156_C":-1.0
},
"lower_bound":0.0,
"upper_bound":1000.0,
"gene_reaction_rule":"ACIAD2950"
},
{
"id":"RXN_1126",
"name":"caffeate coa ligase",
"metabolites":{
"cpd00002_C":-1.0,
"cpd00010_C":-1.0,
"cpd00012_C":1.0,
"cpd00018_C":1.0,
"cpd00270_C":1.0,
"cpd00881_C":-1.0
},
"lower_bound":0.0,
"upper_bound":1000.0,
"gene_reaction_rule":"ACIAD1724"
},
{
"id":"TRANS_RXN_CARBON_DIOXIDE",
"name":"transport of co2 [extraorganism;cytosol](passive)",
"metabolites":{
"cpd00011_C":1.0,
"cpd00011_E":-1.0
},
"lower_bound":-1000.0,
"upper_bound":1000.0,
"gene_reaction_rule":""
},
{
"id":"EXF_NITRATE_E",
"name":"exchange flux of no3-",
"metabolites":{
"cpd00209_E":-1.0
},
"lower_bound":-0.0,
"upper_bound":1000.0,
"gene_reaction_rule":""
},
{
"id":"C6_0_2ENOYLCOA_HYDRATASE_RXN",
"name":"enoyl-coa hydratase (c6:0)",
"metabolites":{
"cpd00001_C":-1.0,
"cpd03122_C":1.0,
"cpd03125_C":-1.0
},
"lower_bound":0.0,
"upper_bound":1000.0,
"gene_reaction_rule":"ACIAD1692 or ACIAD1726 or ACIAD1820 or ACIAD1829 or ACIAD2351 or ACIAD3073 or ACIAD1609 or ACIAD0335 or ACIAD1608 or ACIAD1701 or ACIAD1565 or ACIAD2416"
},
{
"id":"C16_0_3HYDROXYACP_DEHYDRATASE_RXN",
"name":"3hydroxyacyl-acp dehydratase (c16:0)",
"metabolites":{
"C16_0_2ENOYLACP_C":1.0,
"C16_0_3HYDROXYACP_C":-1.0,
"cpd00001_C":1.0
},
"lower_bound":0.0,
"upper_bound":1000.0,
"gene_reaction_rule":"ACIAD1381"
},
{
"id":"EXF_ETHANESULFONATE_E",
"name":"exchange flux of ethane sulfonate",
"metabolites":{
"cpd11579_E":-1.0
},
"lower_bound":-0.0,
"upper_bound":1000.0,
"gene_reaction_rule":""
},
{
"id":"EXF_WATER_E",
"name":"exchange flux of h2o",
"metabolites":{
"cpd00001_E":-1.0
},
"lower_bound":-1000.0,
"upper_bound":1000.0,
"gene_reaction_rule":""
},
{
"id":"HOMOSERINE_O_ACETYLTRANSFERASE_RXN",
"name":"homoserine o-acetyltransferase",
"metabolites":{
"cpd00010_C":1.0,
"cpd00022_C":-1.0,
"cpd00227_C":-1.0,
"cpd00790_C":1.0
},
"lower_bound":-1000.0,
"upper_bound":1000.0,
"gene_reaction_rule":"ACIAD0529"
},
{
"id":"IMIDPHOSDEHYD_RXN",
"name":"imidazoleglycerol-phosphate dehydratase",
"metabolites":{
"cpd00001_C":1.0,
"cpd00930_C":1.0,
"cpd02843_C":-1.0
},
"lower_bound":0.0,
"upper_bound":1000.0,
"gene_reaction_rule":"ACIAD3395"
},
{
"id":"ACYLACYLTRAN_RXN",
"name":"acyl acyltransferase",
"metabolites":{
"ACP_C":2.0,
"ACYL_ACP_LPS_C":-2.0,
"cpd03586_C":-1.0,
"cpd03587_C":1.0
},
"lower_bound":0.0,
"upper_bound":1000.0,
"gene_reaction_rule":"ACIAD2638 or ACIAD0484"
},
{
"id":"TRANS_RXN_151",
"name":"transport of so42- [extraorganism;cytosol](secondary symport)",
"metabolites":{
"cpd00048_C":1.0,
"cpd00048_E":-1.0,
"cpd00067_C":1.0,
"cpd00067_P":-1.0
},
"lower_bound":0.0,
"upper_bound":1000.0,
"gene_reaction_rule":"ACIAD3384 or ACIAD1241"
},
{
"id":"THI_P_KIN_RXN",
"name":"thiamine-phosphate kinase",
"metabolites":{
"cpd00002_C":-1.0,
"cpd00008_C":1.0,
"cpd00056_C":1.0,
"cpd00793_C":-1.0
},
"lower_bound":-1000.0,
"upper_bound":1000.0,
"gene_reaction_rule":"ACIAD3573"
},
{
"id":"C6_0_CARBOXYCOA_DEHYDROGENASE_RXN",
"name":"dicarboxylate acyl-coa dehydrogenase (c6:0)",
"metabolites":{
"cpd00015_C":-1.0,
"cpd00982_C":1.0,
"cpd09842_C":-1.0,
"cpd28230_C":1.0
},
"lower_bound":0.0,
"upper_bound":1000.0,
"gene_reaction_rule":"ACIAD1693"
},
{
"id":"EXF_PROPIONATE_E",
"name":"exchange flux of propionate",
"metabolites":{
"cpd00141_E":-1.0
},
"lower_bound":-0.0,
"upper_bound":1000.0,
"gene_reaction_rule":""
},
{
"id":"PYRUVDEH_RXN",
"name":"pyruvate dehydrogenase",
"metabolites":{
"cpd00003_C":-1.0,
"cpd00004_C":1.0,
"cpd00010_C":-1.0,
"cpd00011_C":1.0,
"cpd00020_C":-1.0,
"cpd00022_C":1.0
},
"lower_bound":0.0,
"upper_bound":1000.0,
"gene_reaction_rule":"ACIAD3507 and ACIAD3506 and ACIAD2874"
},
{
"id":"C14_0_COATRANSFERASE_RXN",
"name":"acyl coa transferase (c14:0)",
"metabolites":{
"cpd00002_C":-1.0,
"cpd00010_C":-1.0,
"cpd00012_C":1.0,
"cpd00018_C":1.0,
"cpd01695_C":1.0,
"cpd03847_C":-1.0
},
"lower_bound":0.0,
"upper_bound":1000.0,
"gene_reaction_rule":"ACIAD1818 or ACIAD0235 or ACIAD1724 or ACIAD2106"
},
{
"id":"C16_0_COA_THIOLASE_RXN",
"name":"acyl-coa thiolase (c16:0)",
"metabolites":{
"cpd00001_C":-1.0,
"cpd00010_C":1.0,
"cpd00134_C":-1.0,
"cpd00214_C":1.0
},
"lower_bound":0.0,
"upper_bound":1000.0,
"gene_reaction_rule":"ACIAD0341 or ACIAD3231 or ACIAD1023"
},
{
"id":"DIHYDRONEOPTERIN_MONO_P_DEPHOS_RXN",
"name":"dihydroneopterin monophosphate dephosphorylase",
"metabolites":{
"cpd00001_C":-1.0,
"cpd00009_C":1.0,
"cpd02961_C":1.0,
"cpd03521_C":-1.0
},
"lower_bound":0.0,
"upper_bound":1000.0,
"gene_reaction_rule":""
},
{
"id":"EXF_D_GALACTARATE_E",
"name":"exchange flux of d-galactarate",
"metabolites":{
"cpd00652_E":-1.0
},
"lower_bound":-0.0,
"upper_bound":1000.0,
"gene_reaction_rule":""
},
{
"id":"EXF_CPD_3571_E",
"name":"exchange flux of tricarballylate",
"metabolites":{
"cpd16654_E":-1.0
},
"lower_bound":-0.0,
"upper_bound":1000.0,
"gene_reaction_rule":""
},
{
"id":"EXF_ADIPATE_E",
"name":"exchange flux of adipate",
"metabolites":{
"cpd03642_E":-1.0
},
"lower_bound":-0.0,
"upper_bound":1000.0,
"gene_reaction_rule":""
},
{
"id":"TRANS_RXN_BUTYRIC_ACID",
"name":"transport of butanoate [extraorganism;cytosol](passive)",
"metabolites":{
"cpd00211_C":1.0,
"cpd00211_E":-1.0
},
"lower_bound":0.0,
"upper_bound":1000.0,
"gene_reaction_rule":""
},
{
"id":"2OXOGLUTARATEDEH_RXN",
"name":"2-oxogluterate dehydrogenase",
"metabolites":{
"cpd00003_C":-1.0,
"cpd00004_C":1.0,
"cpd00010_C":-1.0,
"cpd00011_C":1.0,
"cpd00024_C":-1.0,
"cpd00078_C":1.0
},
"lower_bound":0.0,
"upper_bound":1000.0,
"gene_reaction_rule":"ACIAD2876 and ACIAD2875 and ACIAD2874"
},
{
"id":"TRANS_RXN_GLYCEROL_OUT",
"name":"transport of glycerol [cytosol;extraorganism](passive)",
"metabolites":{
"cpd00100_C":-1.0,
"cpd00100_E":1.0
},
"lower_bound":0.0,
"upper_bound":1000.0,
"gene_reaction_rule":""
},
{
"id":"UDPKIN_RXN",
"name":"udp kinase",
"metabolites":{
"cpd00002_C":-1.0,
"cpd00008_C":1.0,
"cpd00014_C":-1.0,
"cpd00062_C":1.0
},
"lower_bound":-1000.0,
"upper_bound":1000.0,
"gene_reaction_rule":"ACIAD0556"
},
{
"id":"ADPREDUCT_RXN",
"name":"adp reductase",
"metabolites":{
"cpd00001_C":1.0,
"cpd00008_C":-1.0,
"cpd00177_C":1.0,
"cpd11420_C":1.0,
"cpd11421_C":-1.0
},
"lower_bound":0.0,
"upper_bound":1000.0,
"gene_reaction_rule":"ACIAD0724 and ACIAD0722"
},
{
"id":"C16_1_C7_COA_REDUCTASE_RXN",
"name":"fatty acyl-coa reductase (c16:1(c7))",
"metabolites":{
"cpd00005_C":-1.0,
"cpd00006_C":1.0,
"cpd00010_C":1.0,
"cpd16626_C":1.0,
"cpd16627_C":-1.0
},
"lower_bound":-1000.0,
"upper_bound":1000.0,
"gene_reaction_rule":"ACIAD3383"
},
{
"id":"C16_1_C7_COA_THIOLASE_RXN",
"name":"acyl-coa thiolase (c16:1(c7))",
"metabolites":{
"cpd00001_C":-1.0,
"cpd00010_C":1.0,
"cpd16627_C":-1.0,
"cpd34461_C":1.0
},
"lower_bound":0.0,
"upper_bound":1000.0,
"gene_reaction_rule":"ACIAD0341 or ACIAD3231 or ACIAD1023"
},
{
"id":"TRANS_RXNIM_5876",
"name":"transport of d-galactarate [extraorganism;cytosol](passive)",
"metabolites":{
"cpd00652_C":1.0,
"cpd00652_E":-1.0
},
"lower_bound":-1000.0,
"upper_bound":1000.0,
"gene_reaction_rule":"ACIAD0127"
},
{
"id":"TRANS_RXNIM_5874",
"name":"transport of xanthine [extraorganism;cytosol](secondary symport)",
"metabolites":{
"cpd00067_C":1.0,
"cpd00067_P":-1.0,
"cpd00309_C":1.0,
"cpd00309_E":-1.0
},
"lower_bound":0.0,
"upper_bound":1000.0,
"gene_reaction_rule":"ACIAD0031 or ACIAD0136 or ACIAD0702 or ACIAD0762 or ACIAD1243"
},
{
"id":"EXF_BENZYL_ALCOHOL_E",
"name":"exchange flux of benzyl alcohol",
"metabolites":{
"cpd00435_E":-1.0
},
"lower_bound":-0.0,
"upper_bound":1000.0,
"gene_reaction_rule":""
},
{
"id":"PANTOTHENATE_KIN_RXN",
"name":"pantothenate kinase",
"metabolites":{
"cpd00002_C":-1.0,
"cpd00008_C":1.0,
"cpd00644_C":-1.0,
"cpd02201_C":1.0
},
"lower_bound":-1000.0,
"upper_bound":1000.0,
"gene_reaction_rule":"ACIAD0840"
},
{
"id":"HISTIDINE_TRNA_LIGASE_RXN",
"name":"histidine--trna ligase",
"metabolites":{
"CHARGED_HIS_TRNAS_C":1.0,
"HIS_TRNAS_C":-1.0,
"cpd00002_C":-1.0,
"cpd00012_C":1.0,
"cpd00018_C":1.0,
"cpd00119_C":-1.0
},
"lower_bound":0.0,
"upper_bound":1000.0,
"gene_reaction_rule":"ACIAD0562"
},
{
"id":"TRYPSYN_RXN",
"name":"tryptophan synthase",
"metabolites":{
"cpd00001_C":1.0,
"cpd00054_C":-1.0,
"cpd00065_C":1.0,
"cpd00102_C":1.0,
"cpd02210_C":-1.0
},
"lower_bound":0.0,
"upper_bound":1000.0,
"gene_reaction_rule":"ACIAD0642 and ACIAD0636"
},
{
"id":"GMKALT_RXN",
"name":"dgmp kinase",
"metabolites":{
"cpd00002_C":-1.0,
"cpd00008_C":1.0,
"cpd00295_C":1.0,
"cpd00296_C":-1.0
},
"lower_bound":0.0,
"upper_bound":1000.0,
"gene_reaction_rule":"ACIAD3324"
},
{
"id":"ASPARTATE_SEMIALDEHYDE_DEHYDROGENASE_RXN",
"name":"aspartate-semialdehyde dehydrogenase",
"metabolites":{
"cpd00005_C":-1.0,
"cpd00006_C":1.0,
"cpd00009_C":1.0,
"cpd00346_C":1.0,
"cpd01977_C":-1.0
},
"lower_bound":-1000.0,
"upper_bound":1000.0,
"gene_reaction_rule":"ACIAD0479"
},
{
"id":"KDOTRANS_RXN",
"name":"3-deoxy-d-manno-2-octulosonate transferase",
"metabolites":{
"cpd00046_C":1.0,
"cpd02546_C":-1.0,
"cpd02993_C":-1.0,
"cpd03585_C":1.0
},
"lower_bound":0.0,
"upper_bound":1000.0,
"gene_reaction_rule":"ACIAD3479"
},
{
"id":"TRANS_RXN_105",
"name":"transport of glycolate [extraorganism;cytosol](secondary symport)",
"metabolites":{
"cpd00067_C":1.0,
"cpd00067_P":-1.0,
"cpd00139_C":1.0,
"cpd00139_E":-1.0
},
"lower_bound":-1000.0,
"upper_bound":1000.0,
"gene_reaction_rule":"ACIAD0106"
},
{
"id":"FORMYLTHFDEFORMYL_RXN",
"name":"formyltetrahydrofolate deformylase",
"metabolites":{
"cpd00001_C":-1.0,
"cpd00047_C":1.0,
"cpd00087_C":1.0,
"cpd00201_C":-1.0
},
"lower_bound":0.0,
"upper_bound":1000.0,
"gene_reaction_rule":"ACIAD2554 or ACIAD0505"
},
{
"id":"TRANS_RXNIM_5720",
"name":"transport of nh4+ [extraorganism;cytosol](passive)",
"metabolites":{
"cpd00013_C":1.0,
"cpd00013_E":-1.0
},
"lower_bound":-1000.0,
"upper_bound":1000.0,
"gene_reaction_rule":"ACIAD2958 or ACIAD0245 or ACIAD2534"
},
{
"id":"GTP_CYCLOHYDRO_II_RXN",
"name":"gtp cyclohydrolase ii",
"metabolites":{
"cpd00001_C":-3.0,
"cpd00012_C":1.0,
"cpd00038_C":-1.0,
"cpd00047_C":1.0,
"cpd00957_C":1.0
},
"lower_bound":-1000.0,
"upper_bound":1000.0,
"gene_reaction_rule":"ACIAD3249"
},
{
"id":"C18_0_ACP_HYDROLASE_RXN",
"name":"acyl-acp hydrolase (c18:0)",
"metabolites":{
"ACP_C":1.0,
"C18_0_ACP_C":-1.0,
"cpd00001_C":-1.0,
"cpd01080_C":1.0
},
"lower_bound":0.0,
"upper_bound":1000.0,
"gene_reaction_rule":""
},
{
"id":"ACETOOHBUTREDUCTOISOM_RXN",
"name":"ketol-acid reductoisomerase",
"metabolites":{
"cpd00005_C":-1.0,
"cpd00006_C":1.0,
"cpd00498_C":-1.0,
"cpd02535_C":1.0
},
"lower_bound":-1000.0,
"upper_bound":1000.0,
"gene_reaction_rule":"ACIAD3102"
},
{
"id":"EXF_GLY_E",
"name":"exchange flux of glycine",
"metabolites":{
"cpd00033_E":-1.0
},
"lower_bound":-0.0,
"upper_bound":1000.0,
"gene_reaction_rule":""
},
{
"id":"EXF_SUC_E",
"name":"exchange flux of succinate",
"metabolites":{
"cpd00036_E":-1.0
},
"lower_bound":-0.0,
"upper_bound":1000.0,
"gene_reaction_rule":""
},
{
"id":"BENZOATE_DIOXYGENASE_NON_DECARBOX_RXN",
"name":"benzoate 1,2-dioxygenase",
"metabolites":{
"cpd00003_C":1.0,
"cpd00004_C":-1.0,
"cpd00007_C":-1.0,
"cpd00153_C":-1.0,
"cpd02821_C":1.0
},
"lower_bound":0.0,
"upper_bound":1000.0,
"gene_reaction_rule":"ACIAD1436 and ACIAD1437 and ACIAD1438"
},
{
"id":"C6_0_CARBOXY_COATRANSFERASE_RXN",
"name":"dicarboxylate acyl coa transferase (c6:0)",
"metabolites":{
"cpd00002_C":-1.0,
"cpd00010_C":-1.0,
"cpd00012_C":1.0,
"cpd00018_C":1.0,
"cpd03642_C":-1.0,
"cpd09842_C":1.0
},
"lower_bound":0.0,
"upper_bound":1000.0,
"gene_reaction_rule":"ACIAD1695 and ACIAD1696"
},
{
"id":"ORNCARBAMTRANSFER_RXN",
"name":"ornithine carbamoyltransferase",
"metabolites":{
"cpd00009_C":1.0,
"cpd00064_C":-1.0,
"cpd00146_C":-1.0,
"cpd00274_C":1.0
},
"lower_bound":-1000.0,
"upper_bound":1000.0,
"gene_reaction_rule":"ACIAD1366"
},
{
"id":"EXF_DEOXYURIDINE_E",
"name":"exchange flux of deoxyuridine",
"metabolites":{
"cpd00412_E":-1.0
},
"lower_bound":-0.0,
"upper_bound":1000.0,
"gene_reaction_rule":""
},
{
"id":"6_2_1_34_RXN",
"name":"trans-feruloyl-coa synthetase",
"metabolites":{
"cpd00002_C":-1.0,
"cpd00010_C":-1.0,
"cpd00012_C":1.0,
"cpd00018_C":1.0,
"cpd00321_C":1.0,
"cpd01059_C":-1.0
},
"lower_bound":-1000.0,
"upper_bound":1000.0,
"gene_reaction_rule":"ACIAD1724"
},
{
"id":"EXF_HOMO_SER_E",
"name":"exchange flux of homoserine",
"metabolites":{
"cpd00227_E":-1.0
},
"lower_bound":-0.0,
"upper_bound":1000.0,
"gene_reaction_rule":""
},
{
"id":"THYMIDYLATESYN_RXN",
"name":"thymidylate synthase",
"metabolites":{
"cpd00125_C":-1.0,
"cpd00298_C":1.0,
"cpd00299_C":-1.0,
"cpd00330_C":1.0
},
"lower_bound":-1000.0,
"upper_bound":1000.0,
"gene_reaction_rule":"ACIAD0515"
},
{
"id":"GLYC3PDEHYDROGBIOSYN_RXN",
"name":"glycerol-3-phosphate dehydrogenase (nad(p)+)",
"metabolites":{
"cpd00005_C":1.0,
"cpd00006_C":-1.0,
"cpd00080_C":-1.0,
"cpd00095_C":1.0
},
"lower_bound":-1000.0,
"upper_bound":1000.0,
"gene_reaction_rule":"ACIAD1317"
},
{
"id":"TRANS_RXN_CYTIDINE",
"name":"transport of cytidine [extraorganism;cytosol](secondary symport)",
"metabolites":{
"cpd00067_C":1.0,
"cpd00067_P":-1.0,
"cpd00367_C":1.0,
"cpd00367_E":-1.0
},
"lower_bound":0.0,
"upper_bound":1000.0,
"gene_reaction_rule":"ACIAD2389"
},
{
"id":"1_2_1_65_RXN",
"name":"salicylaldehyde dehydrogenase",
"metabolites":{
"cpd00001_C":-1.0,
"cpd00003_C":-1.0,
"cpd00004_C":1.0,
"cpd00599_C":1.0,
"cpd03709_C":-1.0
},
"lower_bound":-1000.0,
"upper_bound":1000.0,
"gene_reaction_rule":"ACIAD1430 or ACIAD1725"
},
{
"id":"EXF_HOMOSERINE_LACTONE_E",
"name":"exchange flux of homoserine lactone",
"metabolites":{
"cpd01873_E":-1.0
},
"lower_bound":-0.0,
"upper_bound":1000.0,
"gene_reaction_rule":""
},
{
"id":"RXN0_901",
"name":"xanthine dehydrogenase",
"metabolites":{
"cpd00001_C":-1.0,
"cpd00003_C":-1.0,
"cpd00004_C":1.0,
"cpd00300_C":1.0,
"cpd00309_C":-1.0
},
"lower_bound":-1000.0,
"upper_bound":1000.0,
"gene_reaction_rule":"ACIAD2466 and ACIAD2467 and ACIAD2468"
},
{
"id":"PRPPSYN_RXN",
"name":"ribose-phosphate pyrophosphokinase",
"metabolites":{
"cpd00002_C":-1.0,
"cpd00018_C":1.0,
"cpd00101_C":-1.0,
"cpd00103_C":1.0
},
"lower_bound":-1000.0,
"upper_bound":1000.0,
"gene_reaction_rule":"ACIAD2907"
},
{
"id":"PROTOPORGENOXI_RXN",
"name":"protoporphyrinogen oxidase",
"metabolites":{
"cpd00001_C":2.0,
"cpd00007_C":-1.0,
"cpd00791_C":-1.0,
"cpd01476_C":1.0
},
"lower_bound":0.0,
"upper_bound":1000.0,
"gene_reaction_rule":""
},
{
"id":"TRANS_RXN_C14_0",
"name":"transport of tetradecanoate [extraorganism;cytosol](secondary symport)",
"metabolites":{
"cpd00067_C":1.0,
"cpd00067_P":-1.0,
"cpd03847_C":1.0,
"cpd03847_E":-1.0
},
"lower_bound":-1000.0,
"upper_bound":1000.0,
"gene_reaction_rule":"ACIAD0835 or ACIAD1427 or ACIAD2788 or ACIAD3392"
},
{
"id":"C6_0_3HYDROXYACP_DEHYDRATASE_RXN",
"name":"3hydroxyacyl-acp dehydratase (c6:0)",
"metabolites":{
"C6_0_2ENOYLACP_C":1.0,
"C6_0_3HYDROXYACP_C":-1.0,
"cpd00001_C":1.0
},
"lower_bound":0.0,
"upper_bound":1000.0,
"gene_reaction_rule":"ACIAD1381"
},
{
"id":"H2NEOPTERINALDOL_RXN",
"name":"dihydroneopterin aldolase",
"metabolites":{
"cpd00229_C":1.0,
"cpd00954_C":1.0,
"cpd02961_C":-1.0
},
"lower_bound":0.0,
"upper_bound":1000.0,
"gene_reaction_rule":"ACIAD2408"
},
{
"id":"TRANS_RXN_PRO",
"name":"transport of l-proline [extraorganism;cytosol](secondary symport)",
"metabolites":{
"cpd00067_C":1.0,
"cpd00067_P":-1.0,
"cpd00129_C":1.0,
"cpd00129_E":-1.0
},
"lower_bound":0.0,
"upper_bound":1000.0,
"gene_reaction_rule":"ACIAD1168"
},
{
"id":"C5_0_COA_DEHYDROGENASE_RXN",
"name":"acyl-coa dehydrogenase (c5:0)",
"metabolites":{
"cpd00015_C":-1.0,
"cpd00660_C":-1.0,
"cpd00982_C":1.0,
"cpd01613_C":1.0
},
"lower_bound":0.0,
"upper_bound":1000.0,
```
